# Effector membrane translocation biosensors reveal G protein and βarrestin coupling profiles of 100 therapeutically relevant GPCRs

**DOI:** 10.1101/2020.04.20.052027

**Authors:** Charlotte Avet, Arturo Mancini, Billy Breton, Christian Le Gouill, Alexander S. Hauser, Claire Normand, Hiroyuki Kobayashi, Florence Gross, Mireille Hogue, Viktoriya Lukasheva, Stéphane St-Onge, Marilyn Carrier, Madeleine Héroux, Sandra Morissette, Eric Fauman, Jean-Philippe Fortin, Stephan Schann, Xavier Leroy, David E. Gloriam, Michel Bouvier

## Abstract

The recognition that individual GPCRs can activate multiple signaling pathways has raised the possibility of developing drugs selectively targeting therapeutically relevant ones. This requires tools to determine which G proteins and βarrestins are activated by a given receptor. Here, we present a set of BRET sensors monitoring the activation of the 12 G protein subtypes based on the translocation of their effectors to the plasma membrane (EMTA). Unlike most of the existing detection systems, EMTA does not require modification of receptors or G proteins (except for G_s_). EMTA was found to be suitable for the detection of constitutive activity, inverse agonism, biased signaling and polypharmacology. Profiling of 100 therapeutically relevant human GPCRs resulted in 1,500 pathway-specific concentration-response curves and revealed a great diversity of coupling profiles ranging from exquisite selectivity to broad promiscuity. Overall, this work describes unique resources for studying the complexities underlying GPCR signaling and pharmacology.

## Introduction

G protein-coupled receptors (GPCRs) play crucial roles in the regulation of a wide variety of physiological processes and represent one-third of clinically prescribed drugs (Hauser, Attwood, Rask-Andersen, Schioth, & Gloriam, 2017). Classically, GPCR-mediated signal transduction was believed to rely on linear signaling pathways whereby a given GPCR selectively activates a single G protein family, defined by the nature of its Gα subunit (Oldham & Hamm, 2008). Gα proteins are divided into four major families (G_s_, G_i/o_, G_q/11_, and G_12/13_) encoded by 16 human genes. Once activated, these proteins each trigger different downstream effectors yielding different biological outcomes. It has now become evident that many GPCRs can couple to more than one G protein family and that ligands can selectively promote the activation of different subsets of these pathways (Namkung et al., 2018; Quoyer et al., 2013). These observations extended the concept of ligand-biased signaling, which was first established for ligand-directed selectivity between βarrestin and G protein (Azzi et al., 2003; Wei et al., 2003), to functional selectivity between G proteins. Ligand-directed functional selectivity represents a promising avenue for GPCRs drug discovery since it offers the opportunity of activating pathways important for therapeutic efficacy while minimizing activation of pathways responsible for undesirable side effects (Galandrin, Oligny-Longpre, & Bouvier, 2007; Kenakin, 2019).

To fully explore the potential of functional selectivity, it is essential to have an exhaustive description of the signaling partners that can be activated by a given receptor, providing receptor- and ligand-specific signaling signatures. Currently, few assays allow for an exhaustive pathway-specific analysis of GPCR signaling; these include BRET-based G protein activation sensors platforms (Gales et al., 2005; Masuho et al., 2015; Maziarz et al., 2020; Mende et al., 2018; Olsen et al., 2020) and the TGF-α shedding assay (Inoue et al., 2019). However, several of these platforms require modification of G protein subunits that may create functional distortions. Moreover, these assays may detect non-productive conformational rearrangements of the G protein heterotrimer as was recently reported for G_12_ (Okashah et al., 2020).

Here, we describe unique sensors that do not require modification of receptors or G proteins (except for G_s_) for interrogating the signaling profiles of GPCRs. The platform includes 15 pathway-selective enhanced bystander bioluminescence resonance energy transfer (ebBRET) biosensors monitoring the translocation of downstream effectors to the plasma membrane for G_i/o_, G_q/11_ and G_12/13_, the dissociation of the Gα subunit from the plasma membrane for G_s_ and the recruitment of βarrestin to the plasma membrane. Overall, the new ebBRET-based **<u>E</u>**ffector **<u>M</u>**embrane **<u>T</u>**ranslocation **<u>A</u>**ssays, named EMTA, provide a readily accessible large scale and comprehensive platform to study constitutive and ligand-directed GPCR signaling. The signaling signatures of 100 GPCRs using the EMTA platform also provides a rich source of information to explore the principles underlying receptor/G protein/βarrestin coupling selectivity relationships. It thus provides a unique set of tools that is complementary to previously described platforms and existing datasets, and offers a map of the coupling potentials for individual GPCR that will stimulate future studies investigating the relevance of these couplings in different physiological systems.

## Results

### ebBRET-based G protein effector membrane translocation assay (EMTA) allows detection of each Gα protein subunit activation

To detect the activation of Gα subtypes, we created an EMTA biosensor platform based on ebBRET (Namkung et al., 2016) (**Figure 1A**). The biosensors at the heart of EMTA consist of sub-domains of the G protein-effector proteins p63-RhoGEF, Rap1GAP and PDZ-RhoGEF that selectively interact with activated G_q/11_, G_i/o_ or G_12/13_, respectively. These domains were fused at their C-terminus to *Renilla* luciferase (RlucII) and co-expressed with different unmodified receptor and Gα protein subtypes. Upon GPCR activation, the energy donor-fused effectors translocate to the plasma membrane to bind activated Gα proteins, bringing RlucII in close proximity to the energy acceptor, *Renilla* green fluorescent protein, targeted to the plasma membrane through a CAAX motif (rGFP-CAAX), thus leading to an increase in ebBRET. The same plasma membrane translocation principle is used to measure βarrestin recruitment (Namkung et al., 2016) (**Figure 1B,** top). Because no selective soluble downstream effector of G_s_ exists, the assay was modified taking advantage of Gα_s_ dissociation from the plasma membrane following its activation (Wedegaertner, Bourne, & von Zastrow, 1996). In this configuration, the RlucII is directly fused to Gα_s_ (Carr et al., 2014). Its activation upon GPCR stimulation leads to its dissociation from the plasma membrane (Martin & Lambert, 2016), resulting in a reduction in ebBRET (**Figure 1B,** bottom).

**Figure 1.**
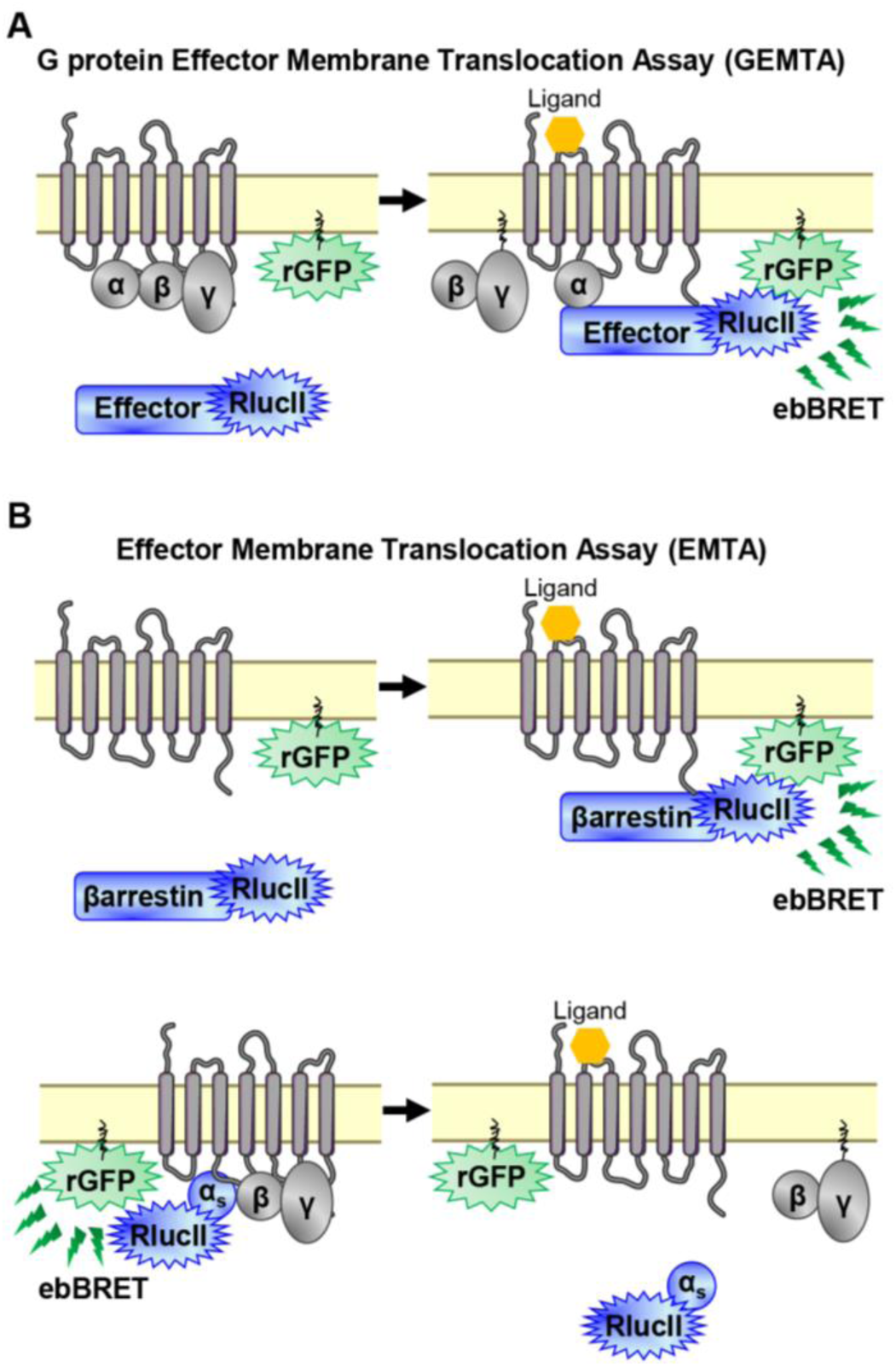
EMTA ebBRET platform to monitor G protein activation and βarrestin recruitment. (**A**) Schematic of the G protein Effector Membrane Translocation Assay (GEMTA) to monitor Gα protein activation. Upon receptor activation, RlucII-tagged effector proteins (Effector-RlucII) translocate towards and interact with active Gα subunits from each G protein family, leading to increased ebBRET. (**B**) Principle of the Effector Membrane Translocation Assay (EMTA) monitoring βarrestin recruitment to the plasma membrane (*top*) and Gα_s_ activation (*bottom*). *Top*; upon receptor activation, RlucII-tagged βarrestins (βarrestin-RlucII) translocate to the plasma membrane, thus increasing ebBRET with rGFP-CAAX. *Bottom*; Internalization of activated RlucII-tagged Gα_s_ (Gα_s_-RlucII) following receptor stimulation decreases ebBRET with the membrane-anchored rGFP-CAAX.

The sensitivity and selectivity of the newly created G protein EMTA biosensors, were validated using prototypical GPCRs known to activate specific Gα subtypes. The responses were monitored upon heterologous expression of specific Gα subunits belonging to G_i/o_, G_q/11_ or G_12/13_ families in the absence or presence of pharmacological inhibitors and using engineered cells lacking selected Gα subtypes. The dopamine D_2_ receptor was used to validate the ability of the G_i/o_ binding domain of Rap1GAP (Jordan, Carey, Stork, & Iyengar, 1999; Meng, Glick, Polakis, & Casey, 1999) to selectively detect G_i/o_ activation. The dopamine-promoted increase in ebBRET between Rap1GAP-RlucII and rGFP-CAAX in the presence of Gα_i/o_ subunits was not affected by the G_q/11_-selective inhibitor UBO-QIC (a.k.a., FR900359 (Schrage et al., 2015); **Figure 2A**, left), whereas the Gα_i/o_ family inhibitor, pertussis toxin (PTX), completely blocked the response for all members of Gα_i/o_ family except for Gα_z_, known to be insensitive to PTX (Casey, Fong, Simon, & Gilman, 1990) (**Figure 2A**, right). Gonadotropin-releasing hormone (GnRH) stimulation of the GnRH receptor (GnRHR), used as a prototypical G_q/11_-coupled receptor, promoted ebBRET between the RlucII-fused G_q/11_ binding domain of p63-RhoGEF (p63-RhoGEF-RlucII) (Lutz et al., 2007; Rojas et al., 2007) and rGFP-CAAX. The ebBRET increase observed in the presence of different Gα_q/11_ subunits was not significantly (p=0.077, 0.0636 and 0.073 for G_q_, G_11_ and G_14_, respectively) affected by PTX (**Figure 2B**, right), whereas UBO-QIC completely blocked the response for all members of Gα_q/11_ family except for Gα_15_, known to be insensitive to UBO-QIC (Schrage et al., 2015) (**Figure 2B**, left). These two G protein specific EMTA were sensitive enough to detect responses elicited by endogenous G proteins since deletion of G_i/o_ (ΔG_i/o_) or G_q/11_ (ΔG_q/11_) subtypes completely abolished the responses induced by D_2_ or GnRHR activation in the absence of heterologously expressed G proteins (**Figure S1I**). It should however be noted that relying on endogenous proteins does not allow the identification of specific members of G_i/o_ (i.e.: G_i1_, G_i2_, G_i3_, G_oA_, G_oB_ or G_z_) or G_q/11_ (i.e.: G_q_, G_11_, G_14_ or G_15_) families.

**Figure 2.**
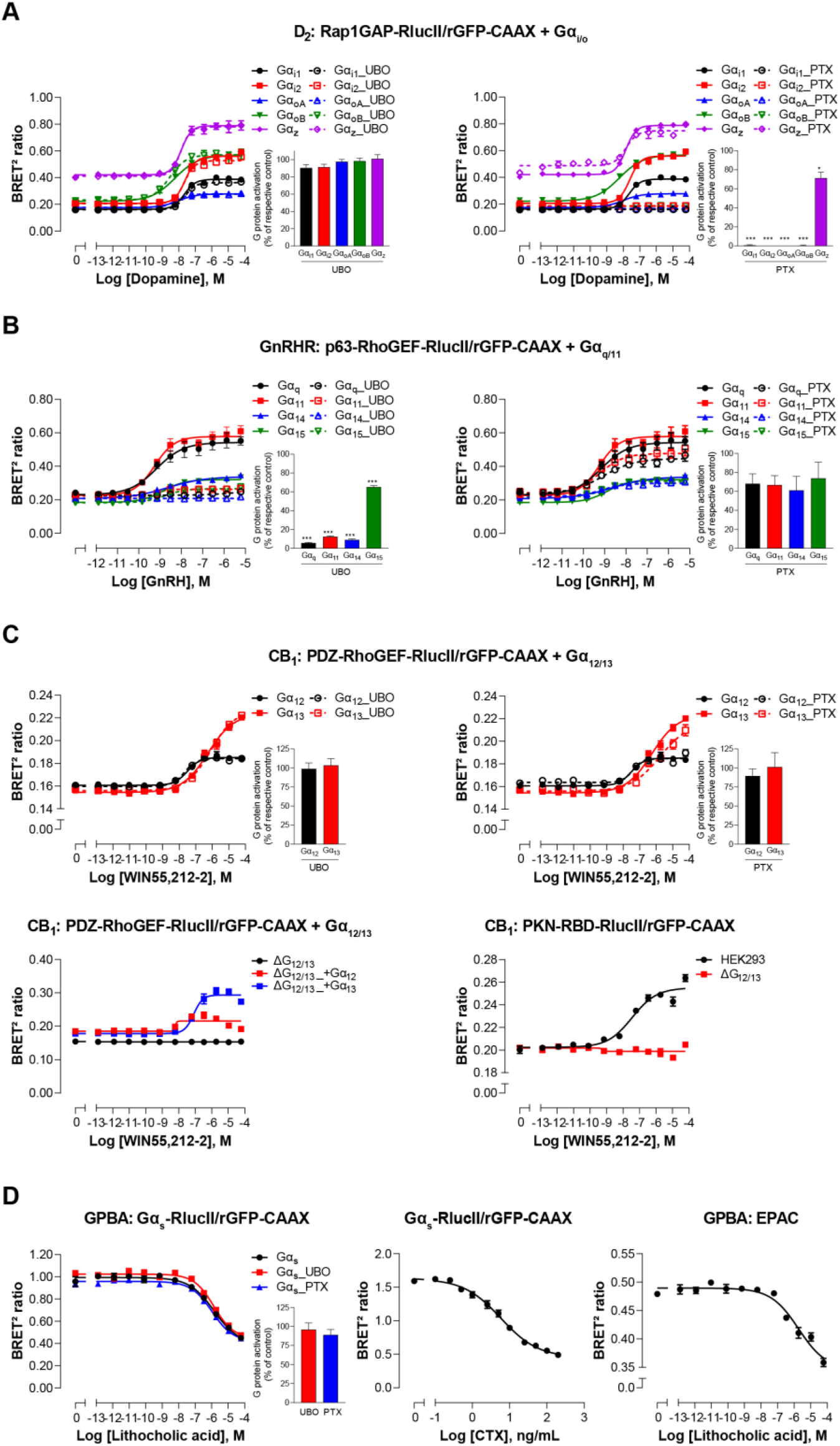
Validation of EMTA ebBRET-based platform to monitor Gα protein activation. (**A**) Pharmacological validation of the Gα_i/o_ activation sensor. HEK293 cells were transfected with the D_2_ receptor and the Gα_i/o_ family-specific sensor, along with each Gα_i/o_ subunit. Dose-response curve using the Gα_i/o_ activation sensor, in the presence or absence of UBO-QIC (*left*) or PTX (*right*) inhibitors. *Insets*; E_max_ values determined from dose-response curves of inhibitor-pretreated cells. (**B**) Pharmacological validation of the Gα_q/11_ activation sensor. HEK293 cells were transfected with the GnRH receptor and the Gα_q/11_ family- specific sensors, along with each Gα_q/11_ subunit. Dose-response curve using Gα_q/11_ activation sensor, in the presence or absence of UBO-QIC (*left*) or PTX (*right*) inhibitors. *Insets*; E_max_ values determined from dose- response curves of inhibitor-pretreated cells. (**C**) Validation of the Gα_12/13_ activation sensor. Cells were transfected with the CB_1_ receptor and one of the Gα_12/13_ activation sensors, along with the Gα_12_ or Gα_13_ subunits. Dose-response curves of HEK293 cells (*top*) or the parental and devoid of G_12/13_ (ΔG_12/13_) HEK293 cells (*bottom*) using the PDZ-RhoGEF-RlucII/rGFP-CAAX sensors (*top and bottom left*) or PKN-RBD- RlucII/rGFP-CAAX (*bottom right*), pretreated or not with UBO-QIC or PTX (*top*). (**D**) Pharmacological validation of the Gα_s_ activation sensor. HEK293 cells were transfected with the GPBA receptor and the Gα_s_ activation (*left and central*) or the EPAC (*right*) sensors. *Left:* Dose-response curves using the Gα_s_ activation sensor in the presence or absence of UBO-QIC or PTX, inhibitors of Gα_q_ or Gα_i/o_, respectively. *Central*: Dose- response activation of the Gα_s_ sensor using CTX, a Gα_s_ activator. *Right*: Dose-response curve using the EPAC sensor. *Inset*; E_max_ values determined from dose-response curves of inhibitors-pretreated cells. Data are expressed as BRET ratio for the dose-response curves or expressed in % of respective control cells (E_max_ graphs) and are means ± SEM of 3 (**A**-**C**) or 4 (**D**) independent experiments performed in one replicate. Unpaired t-test (**A**-**D**): *p < 0.05 and ***p < 0.001 compared to control cells.

The selectivity of the G_12/13_ binding domain of PDZ-RhoGEF (Fukuhara, Chikumi, & Gutkind, 2001) was confirmed using the cannabinoid receptor type 1 (CB_1_). The ebBRET between PDZ-RhoGEF-RlucII and rGFP-CAAX in the presence of Gα_12_ or Gα_13_ promoted by the cannabinoid agonist WIN-55,212-2 was not affected by UBO-QIC (**Figure 2C**, top left), nor PTX (**Figure 2C**, top right). Given the lack of selective G_12/13_ pharmacological inhibitor, we used HEK293 cells genetically deleted for Gα_12_ and Gα_13_ proteins (ΔG_12/13_) to further confirm the response selectivity. As expected, PDZ-RhoGEF-RlucII/rGFP-CAAX ebBRET was observed only following reintroduction of either Gα_12_ (ΔG_12/13__+G_12_) or Gα_13_ (ΔG_12/13__+G_13_) (**Figure 2C**, bottom left). The G_12/13_ coupling of CB_1_ was further confirmed by monitoring the recruitment of PKN to the plasma membrane (**Figure 2C**, bottom right) in agreement with previous reports (Inoue et al., 2019).

To further assess the selectivity of each EMTA biosensor, we took advantage of the fact that the endothelin-1 receptor (ET_A_) can activate G_q/11_, G_i/o_ and G_12/13_ family members. As shown in **Figure S2,** only over-expression of the Gα family members corresponding to their selective effectors (Rap1GAP for G_i/o_, p63-RhoGEF for G_q/11_ and PDZ-RhoGEF for G_12/13_) significantly increased the recruitment of the effector-RlucII to the plasma membrane. A recent study (Chandan N.R., 2021) showed that G_i/o_ can also activate full length PDZ-RhoGEF. Although the domain of PDZ-RhoGEF required for this activation has not been identified yet, the selectivity of our PDZ-RhoGEF sensor for G_12/13_ *vs.* all other G protein families most likely results from the fact that we used a truncated version of PDZ-RhoGEF that only contains the G_12/13_ binding domain and lacks the PDZ domain involved in protein-protein interaction, the actin-binding domain and the DH/PH domains involved in GEF activity and RhoA activation (Aittaleb, Boguth, & Tesmer, 2010).

It should be noted that in the heterologous expression configuration, competition with endogenous G proteins did not occur to a significant extent since the potencies of the responses to a given G protein subtype were not affected by genetic deletion of the different G protein family members (**Figure S1** and **Supplementary Table 1A**). Similarly, overexpression of G proteins, GPCRs or effectors-RlucII did not affect the potencies of the responses observed (**Figure S3** and **Supplementary Table 1B-D**), indicating that, in our experimental conditions, overexpression of the different components of EMTA sensors must likely not bias the coupling response. In addition to spectrometric assessment of coupling selectivity (above) and activation kinetics (**Figure S4**), EMTA allows to image the real-time recruitment of the G protein effectors to the plasma membrane (**Videos 1-3**) thus providing spatiotemporal resolution for the imaging detection of Gα_i/o_, Gα_q/11,14,15_ and Gα_12/13_ activation.

The sensitivity of the EMTA platform is illustrated by a direct side-by-side comparison of the signals detected with EMTA *vs.* BRET assays based on Gαβγ dissociation (Gαβγ) (Gales et al., 2005; Gales et al., 2006; Olsen et al., 2020), that reveals a significantly larger assay windows for EMTA for the 6 Gα subunits tested for 8 selected receptors, (**Figure S5**).

For the Gα_s_ translocation biosensor, the bile acid receptor (GPBA) was chosen for validation (Kawamata et al., 2003). As expected, lithocholic acid stimulation resulted in a concentration-dependent decrease in ebBRET between Gα_s_-RlucII and rGFP-CAAX (**Figure 2D**, left). Cholera toxin (CTX), which directly activates Gα_s_ (De Haan & Hirst, 2004), led to a decrease in ebBRET (**Figure 2D**, center), confirming that loss of Gα_s_ plasma membrane localization results from its activation. The potency of lithocholic acid to promote G_s_ dissociation from the plasma membrane was well in line with its potency to increase cAMP production as assessed using a BRET²-based EPAC biosensor (Leduc et al., 2009) (**Figure 2D**, right). The G_s_-plasma membrane dissociation ebBRET signal was not affected by UBO-QIC or PTX (**Figure 2D**, left), confirming the selectivity of the biosensor.

### Signaling signatures of one hundred therapeutically relevant receptors reveals distinct G protein and βarrestin selectivity profiles

We used EMTA to assess the signaling signature of a panel of 100 human GPCRs that are either already the target of clinically used drugs (74 receptors), considered for pre- or clinical drug development (6 receptors), or pathophysiologically relevant (**Supplementary Table S2A**). To establish the coupling potentials for each receptor, we quantified its ability to activate 15 pathways: Gα_s_, Gα_i1_, Gα_i2_, Gα_oA_, Gα_oB_, Gα_z_, Gα_12_, Gα_13_, Gα_q_, Gα_11_, Gα_14_, Gα_15_ and βarrestin 2 as well as βarrestin 1 and 2 in presence of GRK2 (**Supplementary File 1**).

E_max_ and pEC_50_ values were determined (**Supplementary Table 2**) and, based on the pre- determined threshold criteria (Emax ≥ mean of vehicle-stimulated + 2*SD; see Methods), a ‘yes or no’ agonist-dependent activation was assigned to each signaling pathway and summarized using radial graph representations (**Figure S7**). To assess whether endogenous receptors could contribute to the observed responses, assays were also carried out in cells not transfected with the studied receptor (**Figure S8**). When an agonist-promoted response was observed in non-transfected parental HEK293 cells, this response was not considered as a receptor-specific response (see Methods).

To compare the signaling profiles across all receptors and pathways and to overcome differences in receptor expression levels and individual biosensor dynamic windows, we first min-max normalized E_max_ and pEC_50_ values (between 0 and 1) across receptors as a function of a reference receptor yielding the largest response for a given pathway (**Figure 3A**, left). Then, these values were again min-max normalized (between 0 and 1) for the same receptor across pathways, using the pathway with the largest response for this receptor as the reference (**Figure 3A**, right; see description in Methods). Such double normalization allows direct comparison of the coupling efficiency to different G proteins for a given receptor and across receptors for a given G protein. This coupling efficiency is summarized as heatmaps (**Figure 3B**) that reveals a high diversity of signaling profiles. The selectivity toward the different G protein families varies considerably among GPCRs (**Figure 4**). In our dataset, which is the first using unmodified GPCRs and Gα proteins (except for G_s_), 29% of the receptors coupled to only one family, whereas others displayed more promiscuity by coupling to 2, 3 or 4 families (36%, 25% and 10% respectively). Receptors coupling to a single G protein family favored the members of the G_i/o_ family. Indeed, 27% of the receptors coupling to G_i/o_ only activated this subtype family in comparison to 0, 2.4 and 9.1% for receptors activating G_12/13_, G_q/11_ and G_s_, respectively, thus displaying more promiscuous coupling. A detailed comparative analysis of the selectivity profiles that we observed using the EMTA sensors with that of the chimeric G protein-based assay developed by Inoue *et al*. (Inoue et al., 2019) and the IUPHAR/BPS Guide to Pharmacology database (GtP; https://www.guidetopharmacology.org/) is presented in the accompanying paper (Hauser et al., 2021). **Supplementary Table 2C** allows a direct comparison of the relative potency determined using EMTA for both the new and the already known (i.e.: identified in GtP database) couplings. As can be seen in the table, although in many cases the potency for the novel couplings is lower, this is not a universal finding since for some receptors, the pEC_50_s for the new couplings are similar (ex: G_12_ for CB_1_; G_13_ for serotonin 5-HT_2C_; G_12/13_ for adenosine 2A (A_2A_) and prostaglandin E1 (EP_1_) receptors; G_i/o_ for corticotropin-releasing hormone receptor 1 (CRFR1), ET_A_ and G-protein-coupled receptor 39 (GPR39)) or higher (ex: G_z_ for serotonin 5-HT_2B_; G_15_ for adenosine 3 (A_3_) and melanocortin 3 (MC3R) receptors; G_12_ for bradykinin 2 (B_2_), cholecystokinin A (CCK_1_), chemokine receptor 6 (CCR6) and ET_A_ receptors; G_12/13_ for CRFR1 and GPR68) than those for the canonical ones. Interestingly, in many instances the potency for the newly uncovered couplings are similar to those for βarrestins, which is generally lower than for their canonical G proteins, a finding consistent with the role of βarrestins in signaling arrest at the plasma membrane. The potency differences observed for the activation of different G protein subtypes by a given receptor may lead to preferential activation of some pathways over others. This relative selectivity is likely to be influenced by tissue-dependent G protein subtype expression levels. The physiological consequences of such selectivity remain to be investigated.

**Figure 3.**
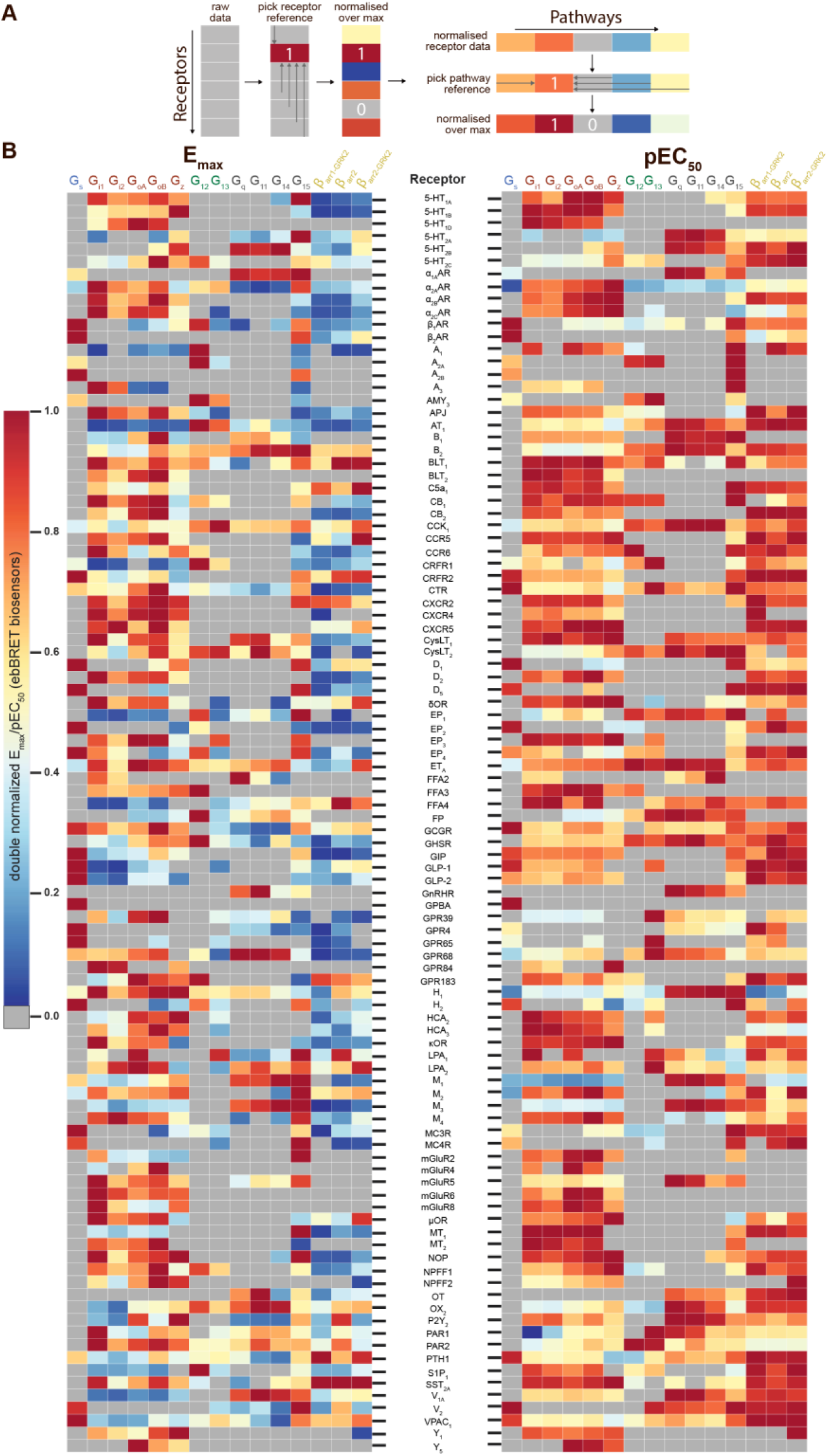
Heatmaps illustrating the diversity of receptor-specific signaling signatures detected with the EMTA ebBRET platform. (**A**) First, values within each pathway were normalized relative to the maximal response observed across all receptors (max = 1; *left*). These values were then normalized across pathways for the same receptor, with the highest-ranking pathway serving as the reference (max = 1; *right*). (**B**) Heatmap representation of double normalized E_max_ (*left*) and pEC_50_ (*right*) data. Empty cells (grey) indicate no detected coupling. IUPHAR receptor names are displayed.

**Figure 4.**
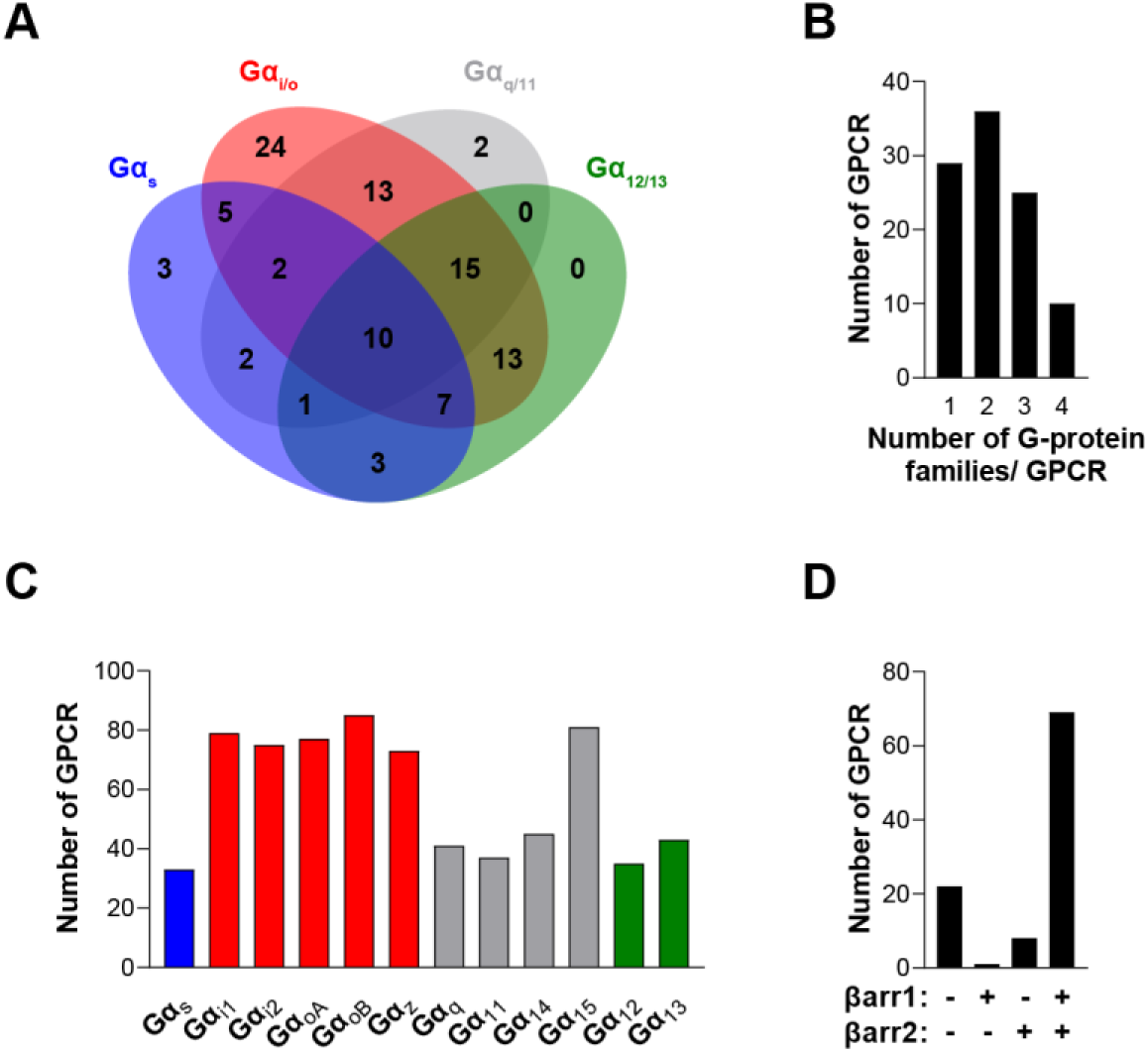
The EMTA ebBRET platform has a unique ability to uncover coupling selectivity between G protein families. (**A**) Venn diagram showing the numbers of receptors coupled to each G protein family in the EMTA ebBRET biosensor assay. (**B**) Evaluation of receptors coupling promiscuity: number of receptors that couple to members of 1, 2, 3 or 4 G protein families. (**C**) Determination of G protein subunit coupling frequency: number of receptors that activate each Gα subunit. (**D**) Proportion of receptors recruiting βarrestins: number of receptors that do not recruit (-/-) or that recruit either (+/- or -/+) or both (+/+) βarrestin isotypes. All data are based on double normalized E_max_ values from Figure 3.

When examining the frequency of coupling for each Gα subunit family (**Figure 4C**), the G_i/o_ family members were the most commonly activated, with 89% of the tested receptors activating a G_i/o_ family member. In contrast, only 33%, 49% and 45% of the receptors activate G_s,_ G_12/13_ or G_q/11_ (excluding Gα_15_) family members, respectively. Not surprisingly, and consistent with its reported promiscuous coupling, Gα_15_ was found to be activated by 81% of the receptors. For some receptors, we also observed preferential coupling of distinct members within a subtype family (**Figure S7**). For instance, 33% of G_i/o_-coupled receptors can couple to only a subpopulation of the family (**Figure S10A**). For the G_q/11_ family, only 44% activate all family members with 45% activating only Gα_15_ and 11% engaging only 2 or 3 members of the family. A matrix expressing the % of receptors engaging a specific Gα subtype that also activated another subtype, is illustrated in **Figure S10B.** When considering individual families, considerable variation within the G_i/o_ family was observed. The greatest similarities were observed between Gα_oB_ and either Gα_oA_ or Gα_z_, and the lowest between Gα_i1_ and Gα_z_. A striking example of intra-family coupling selectivity is the serotonin 5-HT_2B_ that activates only Gα_oB_ and Gα_z_ and GPR65 that selectively activates Gα_oB_. Similarly, when considering the ligand-promoted responses above our threshold criteria (see Methods), histamine H_2_ and MC3R receptors show preferred activation of Gα_oB_ and Gα_z_, whereas the prostaglandin F (FP) and neuropeptide Y5 (Y_5_) receptors preferentially activate Gα_oB_, Gα_oA_ and Gα_z_. Even when all members of a given family are found to be activated, some receptors activate specific family members with greater potencies (**Supplementary Table 2C**).

When considering βarrestin recruitment, our analysis shows that 22% of receptors did not recruit βarrestin1 or 2, even in the presence of overexpressed GRK2 (**Figure 4D**). Among the receptors able to recruit βarrestins, only a very small number selectively recruited βarrestin1 (1.3%) or βarrestin2 (6.4%), most of them recruiting both βarrestins in the presence of GRK2 (92.3%) (**Figure 4D**). Overexpression of GRK2 potentiated the recruitment of βarrestin2 for 68% of receptors highlighting the importance of GRK2 expression level in determining βarrestin activation (**Supplementary File 1** and **Supplementary Table 2**).

### Comparison with Previous Datasets Reveals Commonalities and Crucial Differences

We compared the signaling profiles obtained here with those presented by Inoue *et al*. (Inoue et al., 2019) and the GtP dataset. Of note, this comparison only considers the final reported couplings that in the Inoue’s study were based on the criteria of positive coupling if LogRAi ≥ -1 and negative coupling if LogRAi ≤ -1, and is influenced by the different cut-offs and normalization used in the two studies. A comparison of couplings using common Emax standard deviation cut-off, quantitative normalization and aggregation of G proteins into families is provided in the accompanying paper (Hauser et al., 2021). As can be seen in **Supplementary Table 3A**, among the 70 receptors common to both studies, less couplings were detected in our study than reported in Inoue *et al*. for Gα_s_ (21 *vs.* 28), Gα_i1_ (54 *vs.* 56), Gα_q_ (31 *vs.* 34) and Gα_14_ (36 *vs.* 40). In contrast, more receptors activating Gα_12_ (29 *vs.* 23), Gα_o_ (59 *vs.* 41), Gα_13_ (30 *vs.* 15), Gα_z_ (52 *vs.* 37) and Gα_15_ (62 *vs.* 15) were detected in our study. When comparing with data collected in GtP, that reports couplings grouped for G protein families (*i.e.*: G_s_, G_i/o_, G_q/11_ or G_12/13_) and not at the single G protein subtype level, we detected less couplings than what was reported in GtP for Gα_s_ (32 *vs.* 37), but more for Gα_i/o_ (89 *vs.* 69), Gα_q/11_ (81 *vs.* 48) and Gα_12/13_ (47 *vs.* 10), among the 99 receptors common to both datasets (**Supplementary Table 3B**).

Altogether, the comparative analysis reveals 64% and 69% identity of couplings between the EMTA and Inoue’s or GtP datasets, respectively. Each dataset reporting unique couplings and missing couplings found in the other two datasets. The reasons for these differences are plausibly due to intrinsic differences in the assays used. For instance, for G_12/13_ and G_15_ specifically, the difference with the GtP dataset most likely results from the fact that in most cases G_12/13_ or G_15_ activation were determined indirectly since, until their recent description (G_12/13_: (Quoyer et al., 2013; Schrage et al., 2015); G_15_:(Inoue et al., 2019; Olsen et al., 2020)), no robust readily available assay existed to monitor the activation of these G proteins.

### Validation of newly identified G_12/13_ and G_15_ couplings

Given the overrepresentation of both G_12/13_ and G_15_ couplings, obtained with the EMTA assays *vs.* those reported by Inoue *et al*. and the GtP datasets, the validity of the EMTA assay to detect real productive couplings, was confirmed using orthogonal assays for selective examples not reported in the two other datasets. For G_12/13_, we used the PKN- based BRET biosensor detecting Rho activation downstream of either G_12/13_ or G_q/11_ (Namkung et al., 2018) and the MyrPB-Ezrin-based BRET biosensor detecting the activation of Ezrin downstream of G_12/13_ (Leguay et al., 2021), both in the absence of heterologously expressed G proteins. Ligand stimulation of FP and CysLT_2_ receptors led to Rho and ezrin activation (**Figure S9A**), that were insensitive to the G_q/11_ inhibitor YM- 254890, confirming that these receptors activate Gα_12/13_.

For newly identified G_15_ couplings we took advantage of the lack of Gα_15_ in HEK293 cells and assessed the impact of Gα_15_ heterologous expression on receptor-mediated calcium responses (**Figure S9B**). For prostaglandin E2 (EP_2_) and κ-opioid (κOR) receptors, which couple to G_15_ but no other G_q/11_ members, expression of Gα_15_ significantly increased the PGE2- and Dynorphin A- promoted calcium responses. For α_2A_ adrenergic (α_2A_AR) and vasopressin 2 (V_2_) receptors that couple other G_q/11_ family members, treatment with YM- 254890 completely abolished the agonist-promoted calcium response in the absence of Gα_15_. In contrast, the calcium response evoked by α_2A_AR and V_2_ agonists following Gα_15_ expression was completely insensitive to YM-254890 (**Figure S9B**), confirming that these receptors can activate this YM-254890-insensitive G protein subtype (Takasaki et al., 2004).

### EMTA platform detects constitutive receptor activity and biased signaling

We went on to assess the ability of the EMTA platform to detect receptor constitutive activity. Transfection of increasing amounts of adenosine A_1_ receptor (A_1_) led to a receptor-dependent increase in basal ebBRET of the Gα_i2_-activation sensor (**Figure 5A**, left), reflecting A_1_ constitutive activity. The A_1_ inverse agonist DPCPX (Lu et al., 2014) dose-dependently decreased the constitutive A_1_-mediated activation of Gα_i2_ (**Figure 5A**, left), indicating that EMTA can detect inverse agonism. Although we can not exclude that the high basal activity resulted from activation by adenosine in the cell culture medium, the fact that high basal activity was observed for A_1_ but not A_3_, despite a similar potency of adenosine to activate these two receptors subtypes (see **Figure S11A**), supports the notion that the increased basal activity reflects A_1_ constitutive activity.

**Figure 5.**
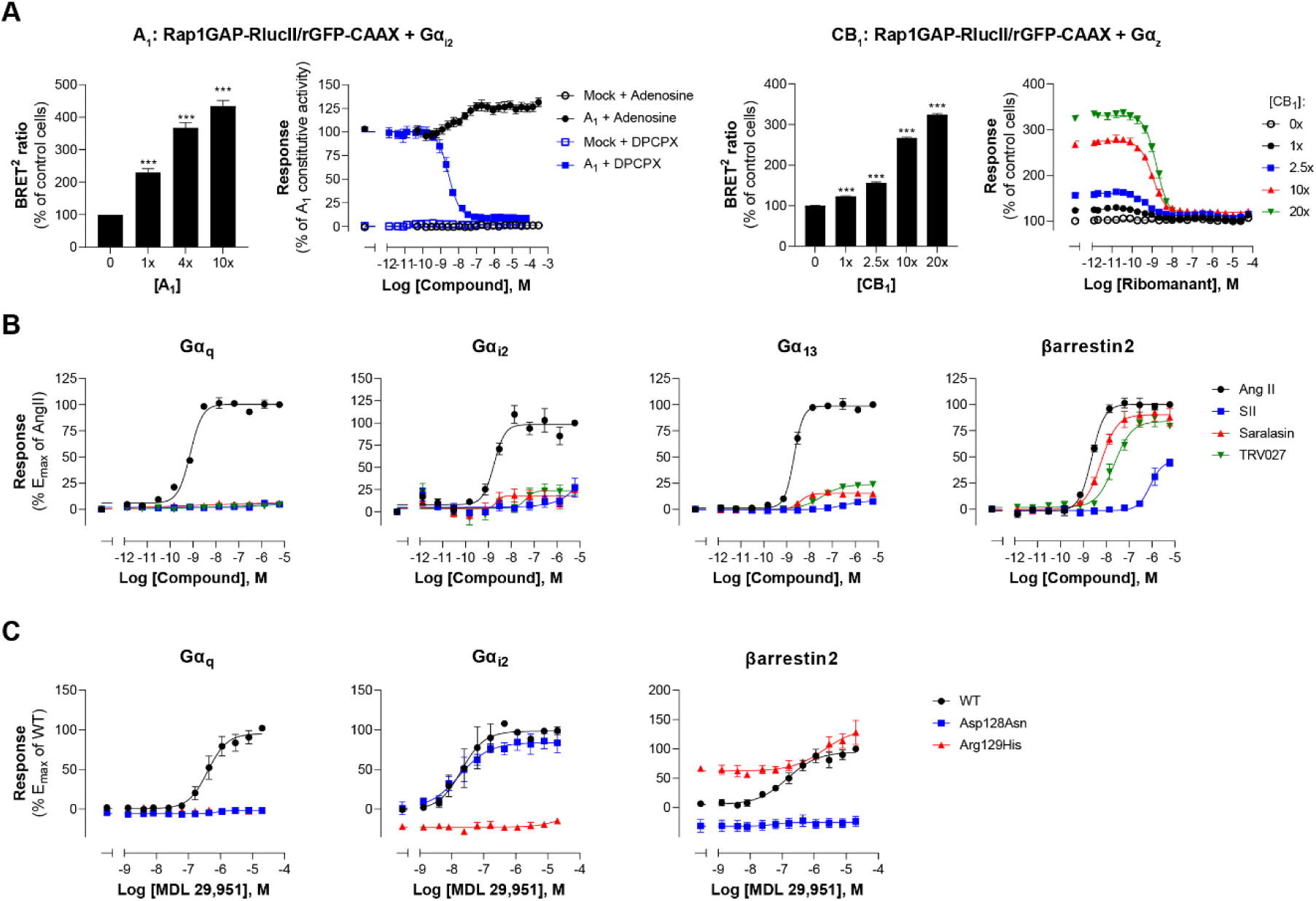
Multiple applications using the EMTA ebBRET platform. (**A**) Inverse agonist activity detection. *Left*: Gα_i2_ activation in HEK293 cells transfected with the Rap1GAP-RlucII/rGFP-CAAX sensors with untagged Gα_i2_ and increasing amount of A_1_ receptor plasmid. Data are expressed in % of response obtained in control cells (0 ng of A_1_) and are means ± SEM of 4-6 independent experiments performed in two replicates. One Way ANOVA test: ***p < 0.001 compared to control cells. HEK293 cells expressing the Gα_i2_ activation sensor and control (Mock) or A_1_ receptor plasmid were stimulated (10 min) with increasing concentrations of the indicated compound. Data are expressed in % of constitutive response obtained in vehicle-treated A_1_ transfected cells and are means ± SEM of 4-6 independent experiments performed in one replicate. *Right:* Gα_z_ activation in HEK293 cells transfected with the Rap1GAP-RlucII/rGFP-CAAX sensors with untagged Gα_z_ and increasing amount of CB_1_ receptor plasmid. Data are expressed in % of response obtained in control cells (0 ng of CB_1_) and are means ± SEM of 4 independent experiments performed in one replicate. One Way ANOVA test: ***p < 0.001 compared to control cells. HEK293 cells expressing the Gα_z_ activation sensor and increasing amount of CB_1_ receptor plasmid were directly stimulated (10 min) with increasing concentrations of the CB_1_ inverse agonist rimonabant. Data are expressed as % of the response obtained in control cells (0 ng of CB_1_) treated with vehicle and are means ± SEM of 4 independent experiments performed in one replicate. (**B**) Ligand-biased detection. Concentration-response curves of AT_1_ for the endogenous ligand (Angiotensin II, AngII) and biased agonists [Sar1-Ile4-Ile8] AngII (SII), saralasin or TRV027. G-protein and βarrestin2 signaling activity were assessed by EMTA platform. Data are expressed in % of maximal response elicited by AngII and are means ± SEM of 3-6 independent experiments performed in one replicate. (**C**) Functional selectivity of naturally occurring receptor variants. Concentration-response curves for WT or E/DRY motif Asp128Asn and Arg129His variants of GPR17 upon agonist stimulation in HEK293 cells co-expressing the indicated EMTA biosensor. Data are expressed in % of maximal response elicited by WT receptor and are means ± SEM of 3 independent experiments performed in one replicate.

To further confirm that the platform can adequately detect inverse agonism, a second receptor for which no endogenous ligand should be present in the media, the CB_1_ receptor, was used. As illustrated in **Figure 5A** (right), increase CB_1_ expression led to a ligand-independent constitutive activation of G_z_, that could be completely blocked by the CB_1_ inverse agonist rimonabant.

EMTA also faithfully detected biased signaling. Indeed, as previously reported (Namkung et al., 2018; Wei et al., 2003), angiotensin analogs such as SII, saralasin or TRV027 displayed biased signaling by promoting efficient βarrestin2 recruitment but marginal or no Gα_q_, Gα_i2_ or Gα_13_ activation as compared to angiotensin II that activated all G proteins and βarrestin2 (**Figure 5B**). The platform was also used to identify biased-signaling resulting from single nucleotide polymorphisms. As shown in **Figure 5C**, two naturally occurring variants of human GPR17 (isoform 2) localised in the TM3 E/DRY motif resulted in altered functional selectivity profiles. Whereas the Asp128Asn variant displayed WT-like activity on Gα_i2_, it lost the ability to activate Gα_q_ and βarrestin2. In contrast, variant Arg129His at the neighboring position resulted in an increased constitutive βarrestin2 recruitment and a loss of Gα_i2_ and Gα_q_ protein signaling.

### Combining G_z_ and G_15_ biosensors for safety panels and systems pharmacology

The G protein coupling profiles obtained for the 100 GPCRs revealed that 95% of receptors activate either Gα_z_ (73%) or Gα_15_ (81%). Measuring activation of both pathways simultaneously provides an almost universal sensor applicable to screening. Combining the two sensors (Rap1GAP-RlucII/p63-RhoGEF-RlucII/rGFP-CAAX) in the same cells allowed to detect ligand concentration-dependent activation of a safety panel of 24 GPCRs, that are well established as contributors to clinical adverse drug reactions (Bowes et al., 2012) (**Figure S12**). Indeed, the G_z_/G_15_ sensor captured the activation of receptors largely or uniquely coupled to either Gα_z_ (e.g., CB_2_) or Gα_15_ (e.g., A_2A_ and A_2B_), as well as receptors coupled (to varying degrees) to both pathways. The usefulness of the G_z_/G_15_ combined sensor to detect off-target ligand activity is illustrated in **Figure 6A**. Most ligands tested were specific for their primary target(s). However, certain ligands displayed functional cross-reactivity with GPCRs other than their cognate targets. These included the activation of the α_2A_AR by dopamine and serotonin, the D_2_ by noradrenaline and serotonin, and of the CB_1_ and CB_2_ receptors by acetylcholine (**Figures 6B-C**). The activation of D_2_ by noradrenaline and serotonin was confirmed by the ability of the D_2_-family selective antagonist eticlopride to block the dopamine-, serotonin- and noradrenaline- promoted responses detected using the combined G_z_/G_15_ or the G_i2_- and G_oB_-selective sensors and βarrestin2 sensor (**Figure 6B**, top). Similarly, use of the α_2_AR selective antagonist, WB4101, allowed to confirm that dopamine can activate Gα_i2_, Gα_oB_ and βarrestin2 through the α_2A_AR (**Figure 6B**, bottom). Such pleiotropic activation of different monoaminergic receptors by catecholamines and serotonin has been previously observed (Roth, Sheffler, & Kroeze, 2004; Sanchez-Soto et al., 2016; Sunahara et al., 1991). Direct activation of the α_2A_AR by dopamine was confirmed by showing that treatment with the D_2_-family receptor selective antagonist eticlopride had negligible effect on dopamine-mediated activation of Gα_i2_ and Gα_oB_ in cells heterologously expressing α_2A_AR, confirming that the response did not result from the activation of endogenously expressed dopamine receptor. In contrast, eticlopride blocked the activation of Gα_i2_ and Gα_oB_ in cells heterologously expressing D_2_ (**Figure S13**).

**Figure 6.**
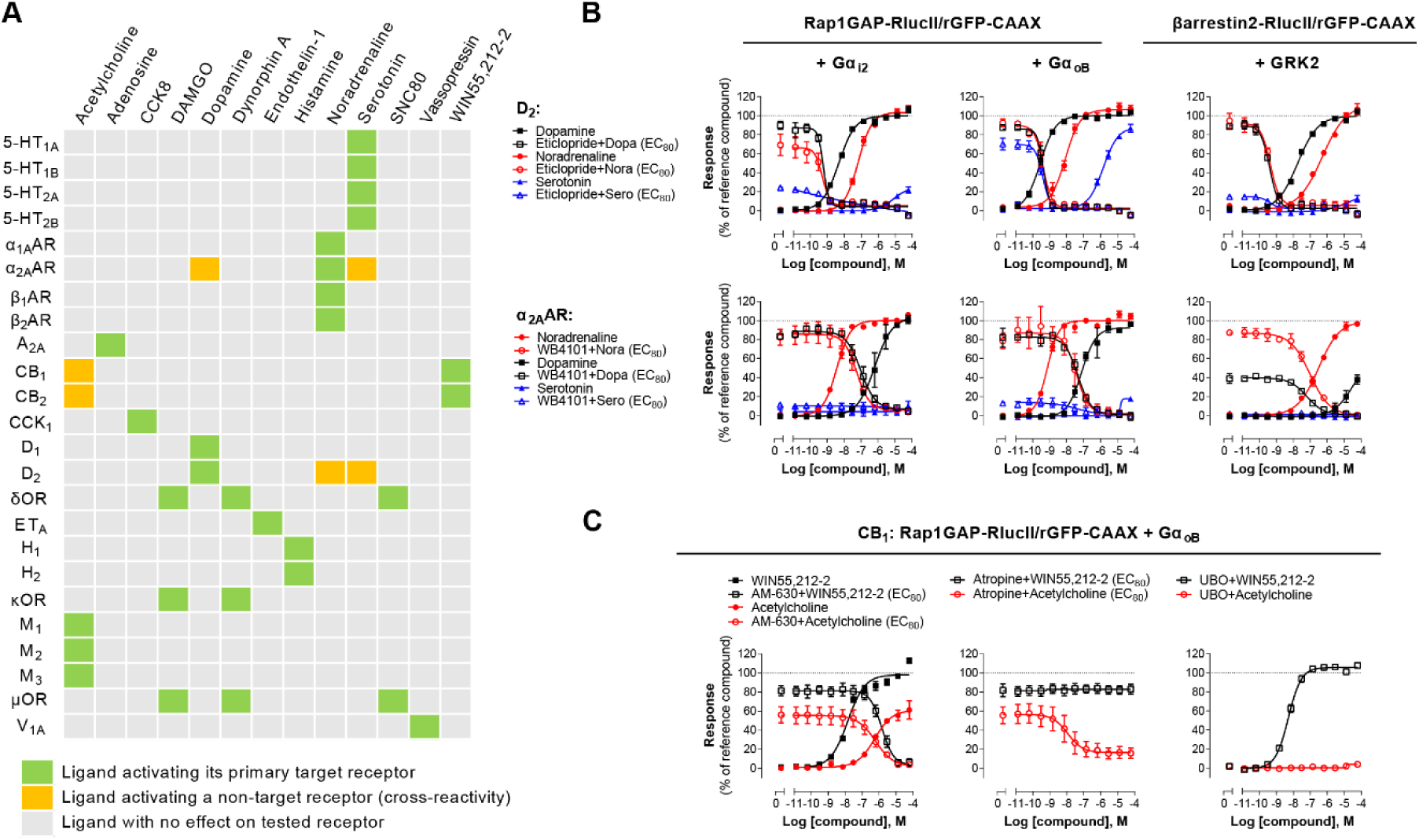
Detection of direct and indirect (*trans*) mechanisms of ligand polypharmacology using the G_z_/G_15_ biosensor. (**A**) Test of the G_z_/G_15_ biosensor on a safety target panel. ebBRET signal was measured before and after stimulation with the indicated ligand in HEK293 cells transfected with the combined G_z_/G_15_ biosensor and one of the 24 receptors listed. (**B**) Cross-activation of D_2_ and α_2A_AR by others natural ligands. For the agonist mode read, HEK293 cells expressing D_2_ or α_2A_AR and either the Gα_i2_, Gα_oB_, or the βarrestin2+GRK2 sensors were stimulated with increasing concentrations of the indicated ligand. For the antagonist mode read, cells were pretreated with increasing concentrations of the selective D_2_ antagonist eticlopride or the selective α_2A_AR antagonist WB4101 before stimulation with an EC_80_ of the indicated ligand. Data are means ± SEM from 3-4 independent experiments performed in one replicate and expressed in % of the response elicited by dopamine or noradrenaline for D_2_ and α_2A_AR expressing cells, respectively. (**C**) Indirect (*trans*) activation of CB_1_ by acetylcholine. For the agonist mode read, HEK293 cells expressing CB_1_ and the Rap1GAP-RlucII/rGFP-CAAX sensors with untagged Gα_oB_ were stimulated with increasing concentrations of the indicated ligand. For the antagonist mode read, same cells were pretreated or not with increasing concentrations of the CB inverse agonist AM-630 (*left*) or the cholinergic antagonist atropine (*central*) before stimulation with an EC_80_ of the indicated ligand. To evaluate the contribution of G_q/11_- coupled receptor, cells were pretreated with the Gα_q_ inhibitor UBO-QIC and then stimulated with increasing concentrations of the indicated ligand (*right*). Data are means ± SEM from 3-5 independent experiments performed in one replicate and expressed in % of the response elicited by WIN55,212-2.

These cross-reactivity may be direct (i.e., via direct binding of a ligand to its non-cognate receptor) as suggested above, or indirect (e.g., “trans”, via ligand activation of its canonical receptor, leading to subsequent secretion of factors that activate the non-canonical target). One such example of trans-activation is provided by the activation of cannabinoid CB_1_ and CB_2_ receptors by acetylcholine (detected by the G_z/15_ and confirmed with the G_oB_ sensors; **Figure 6A**, **C**). Indeed, the activation was completely inhibited by both the CB inverse agonist AM-630 and by the cholinergic antagonist atropine (**Figure 6C**, left). Yet the response evoked by the CB selective agonist WIN55,212 2 was not blocked by atropine (**Figure 6C**, center). Gα_oB_ activation by acetylcholine did not result from direct activation of endogenous muscarinic receptors since no Gα_oB_ response was observed in parental cells (**Figure S8**). Given that the M_3_ muscarinic receptor, which is endogenously expressed at relatively high levels in HEK293 cells (Atwood, Lopez, Wager-Miller, Mackie, & Straiker, 2011), is strongly coupled to the G_q/11_, CB_1_-expressing cells were pretreated with G_q/11/14_ inhibitor UBO-QIC prior to stimulation with acetylcholine. UBO-QIC pre-treatment blocked acetylcholine- but not WIN55,212-2-mediated Gα_oB_ activation (**Figure 6C**, right). These results demonstrate that CB_1_ activation by acetylcholine is indirect and potentially involves the secretion of an endogenous CBR ligand following activation of G_q/11_ by endogenous muscarinic acetylcholine receptors. The combined G_z_/G_15_ sensor is therefore a useful tool to identify interplay between receptors and to explore systems pharmacology resulting from such cross-talks.

## Discussion

This study describes the development and validation of a genetically encoded ebBRET-based biosensor platform allowing live-cell mapping of GPCR-G protein coupling preferences covering 12 heterotrimeric G proteins. The novel EMTA biosensors were combined with previously described ebBRET-based βarrestin trafficking sensors (Namkung et al., 2016), providing an unprecedented description of GPCR signaling partner couplings. In addition to providing a resource to study GPCR functional selectivity (see companion paper (Hauser et al., 2021)), the sensors provide versatile and readily usable tools to study, on a large-scale, pharmacological processes such as constitutive activity, inverse agonism, ligand-biased signaling, and signaling cross-talk.

Our EMTA-based biosensor platform offers several advantages relative to other available approaches. First, EMTA provides direct real-time measurement of proximal signaling events following GPCR activation (i.e., Gα protein activation and βarrestin recruitment) and resulting in lower level of amplification than those of assays relying on enzymatic activity of downstream effectors (i.e.: adenylyl cyclase or phospholipase C) or artificial detection systems (i.e.: gene-reporter or TGF-α shedding assays) that measure signal accumulation sometimes following extended incubation times. In addition, measuring proximal activity reduces the risk of cross-talks between pathways that may complicate data interpretation when considering downstream signaling as the readout (Mancini, Frauli, & Breton, 2015).

Second, EMTA uses native untagged GPCRs and G protein subunits (except for G_s_), contrary to protein complementation (Laschet, Dupuis, & Hanson, 2019), FRET/BRET-based Gαβγ dissociation/receptor-G protein interaction (Bunemann, Frank, & Lohse, 2003; Gales et al., 2005; Gales et al., 2006; Hoffmann et al., 2005; Namkung et al., 2018; Olsen et al., 2020) or TGF-α shedding (Inoue et al., 2019) assays. Modifying these core-signaling components could alter responses, complicate interpretation and explain some of the discrepancies observed between the EMTA platform and other approaches used to study G protein activation. Moreover, the ability to work with unmodified receptors and G proteins (except for G_s_) offers numerous advantages. First, it allows for the detection of endogenous GPCR signaling in either generic HEK293 cells (**Figure S8**) or more physiologically relevant cell lines such as induced pluripotent stem cell (iPSC)-derived cardiomyocytes (**Figure 7A**) and promyelocytic HL-60 cells (**Figure 7B**). Further it allows, in cells expressing sufficient endogenous level of the G proteins of interest, to detect activation of both native receptor and G proteins with no need of overexpression (**Figure 7C-D**). This is illustrated by the ability to detect the recruitment of Rap1GAP upon activation of the endogenous G_i/o_ family members by the formyl peptide receptor 2 (FPR2) in HL-60 cells (**Figure 7C**) or protease-activated receptor-2 (PAR2) in HEK293 cells (**Figure 7D**). The ability to detect the activation of endogenous G protein was also illustrated in **Figure S1I**, where the responses elicited by agonist stimulation were lost in cells genetically deleted of the G protein engaged by the studied receptor (i.e.: G_q/11_ or G_i/o_ families). Recently, another BRET-based approach (Maziarz et al., 2020), taking advantage of a synthetic peptide recognizing the GTP-bound form of Gα subunits, also allows the detection of native G protein activation offering alternative means to probe coupling selectivity profiles for both endogenously and heterologously expressed GPCRs.

**Figure 7.**
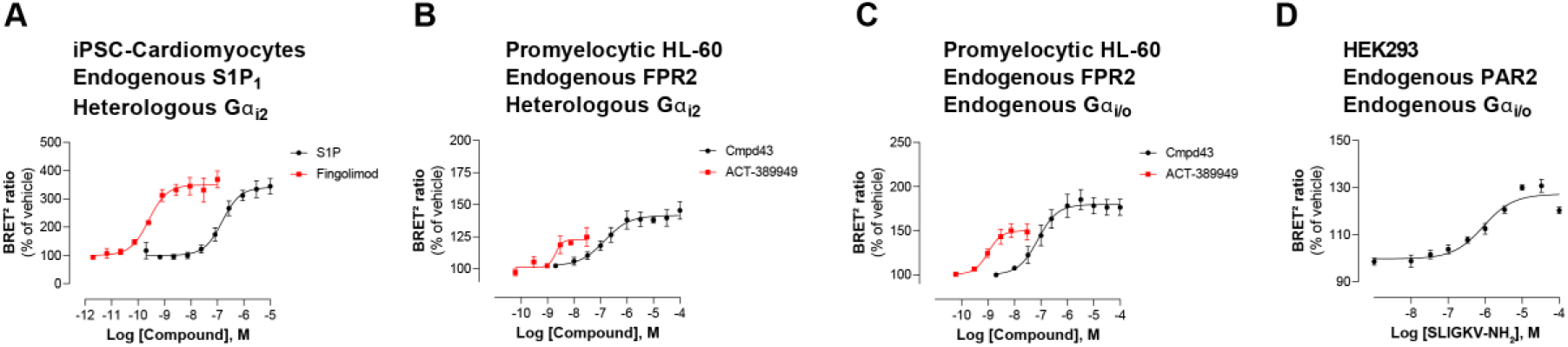
Detection of endogenous receptor- and/or G protein-mediated responses in cells with the EMTA ebBRET platform. Concentration-dependent activation of Gα_i2_ protein by (**A**) endogenous S1P_1_ receptor in iPSC-derived cardiomyocytes transfected with heterologous Gα_i2_, (**B**) endogenous FPR2 in promyelocytic HL-60 cells transfected with heterologous Gα_i2_, (**C**) endogenous FPR2 in promyelocytic HL-60 cells with endogenous G_i/o_ proteins and (**D**) endogenous PAR2 receptor in HEK293 cells with endogenous G_i/o_ proteins. In all cases, cells were co-transfected with the Rap1GAP-RlucII/rGFP-CAAX biosensor. Data are the mean ± SEM of 3-4 independent experiments performed in one replicate and are expressed as BRET^2^ ratio in percentage of response induced by vehicle.

Finally, similarly to BERKY, the EMTA assay platform detects the active form of the Gα subunits rather than the surrogate measurement of Gαβγ dissociation (Gales et al., 2005; Masuho et al., 2015; Maziarz et al., 2020; Mende et al., 2018), which can also detect non-productive binding as recently described for the V_2_ engagement of G_12_ (Okashah et al., 2020).

A potential caveat of EMTA is the use of common downstream effectors for all members of a given G protein family. Indeed, one cannot exclude that distinct members of a given family may display different relative affinities for their common effector. However, such differences are compensated by our data normalization that establishes the maximal response observed for a given subtype as the reference for this pathway (**Figure 3A**), as long as the number of the diversity of receptors including in the analysis is sufficient.

A second potential caveat of EMTA is that, when using heterologously expressed GPCRs and G proteins, some of the responses could result from favorable stoichiometries that may not exist under physiological conditions. It follows that such profiling represents the coupling possibilities of a given GPCR and not necessarily the coupling that will be observed in all cell types. Any couplings observed in such high-throughput studies requires further validation to conclude on their physiological relevance in cells or tissues of interest, and to form hypothesis for futures studies. Because we elected to use unmodified receptors (*i.e.*: not bearing any tags), the expression level of receptors could not be directly monitored. However, the double normalization method developed (see Methods) allows quantitative comparison of coupling preferences across different receptors curtailing the influence of the assay response windows as well as receptor expression levels. Indeed, the double normalization allows ranking the coupling propensity of the receptors first as a function of the receptor which shows the strongest coupling to a specific G protein subtype, and then establishing the maximal response observed for a given G protein subtype as the reference for all G protein activated by a given receptor. In addition, as illustrated using the ET_A_ receptor as example, titrating receptor levels did not influence the pEC_50_ for the activation of the different G protein coupled to this receptor (**Figure S3B** and **Supplementary Table 1C**). Similarly, the pEC_50_ was not affected when titrating the amount of G protein subtype expressed (**Figure S3A** and **Supplementary Table 1B**). As expected, only the amplitude of the response was affected.

It could be argued that overexpressing the G protein effectors (i.e.: p63-RhoGEF, Rap1GAP or PDZ-RhoGEF) used as sensors could influence the couplings observed. This potential caveat is mitigated by the fact that we used truncated part and/or modified versions of these effectors that limit the possibilities of interference with other components of the signaling machinery, and served essentially as a binding detector of the active forms of the G proteins (see Material and Methods). Supporting this notion, titrating the amount of the biosensor effector component did not affect the pEC_50_ of G protein activation (**Figure S3C and Supplementary Table 1D**).

Another limitation of the EMTA platform is the lack of a soluble effector protein selective for activated Gα_s_ thus requiring tagging of the Gα_s_ subunit (**Figure 1B**, bottom) and monitoring its dissociation from the plasma membrane. Yet, our data show that this translocation reflects G_s_ activation state, justifying its use in a G protein activation detection platform.

Finally, because EMTA is able to detect constitutive activity, high receptor expression levels may lead to an elevated basal signal level that may obscure an agonist-promoted response. Such an example can be appreciated for the A_1_ receptor for which the agonist-promoted Gα_i2_ response did not reach the activation threshold criteria because of a very high constitutive activity level (**Figure 5A**). The impact of receptor expression on the constitutive activity and the narrowing on the agonist-promoted response is illustrated for Gα_q_ activation by the 5-HT_2C_ (**Figure S11B**).

A limitation of any large-scale signaling study and drug discovery program is that ligands may elicit responses downstream of receptors other than the one under study. The development of a G_z_/G_15_ quasi-universal biosensor enables efficient screening and detection of such polypharmacology and cross-talk. Using a combination of EMTA and appropriate pharmacological tools, we also proposed a systematic approach to distinguish off-target action of ligands from cross-talk. Interestingly, the cross-talk between the M_3_ and CB receptors detected (**Figure 6**) may have physiological relevance since activation of muscarinic acetylcholine receptors has been shown to enhance the release of endocannabinoids in the hippocampus (Kim, Isokawa, Ledent, & Alger, 2002). The combined G_z_/G_15_ biosensor should be particularly useful for early profiling of compound activity on safety panels and for the design of drugs displaying polypharmacology, an approach that is increasingly considered for the development neuropsychiatric drugs (Roth et al., 2004).

The EMTA platform undoubtedly represents a novel tool-set that could be amenable for high throughput screening of small molecules and biologics across an array of signaling pathways, allowing for the discovery of functionally selective molecules or for GPCR deorphanization campaigns. The ability of the EMTA platform to quantitatively assess -G protein coupling selectivity firmly expands the concept of functional selectivity and potential ligand bias beyond the dichotomic G protein *vs.* βarrestin view and provides plausible functional selectivity profiles that could be tested for their biological and pharmacological outcomes.

## Materials and Methods

### Cells

HEK293 clonal cell line (HEK293SL cells), hereafter referred as HEK293 cells, were a gift from S. Laporte (McGill University, Montreal, Quebec, Canada) and previously described (Namkung et al., 2016). HEK293 cells devoid of functional Gα_s_ (ΔG_s_), Gα_12_ and Gα_13_ (ΔG_12/13_), Gα_q_, Gα_11_, Gα_14_ and Gα_15_ (ΔG_q/11_) and, Gα_i,_ and Gα_o_ (ΔG_i/o_) proteins were a gift from Dr. A. Inoue (Tohoku University, Sendai, Miyagi, Japan) and previously described (Devost et al., 2017; Namkung et al., 2018; Schrage et al., 2015; Stallaert et al., 2017). Cells were maintained in Dulbecco’s Modified Eagle Medium (DMEM, Wisent, Saint-Jean-Baptiste, QC, Canada) supplemented with 10% fetal bovine serum (FBS, Wisent) and 1% antibiotics (100 U/mL penicillin and 100 μg/mL streptomycin (PS); Wisent). HL-60 cells were obtained from ATCC and maintained in RPMI 1640 medium containing L-Glutamine and 25 mM HEPES (Gibco) supplemented with 20% FBS (Wisent) and 1/100 volume PS (Wisent). Differentiation of HL-60 cells into neutrophil-like cells was induced by maintaining the cells in growth medium containing 1.3 % DMSO (Bioshop) during 5 days. Cardiomyocytes derived from induced pluripotent stem cells (iPSCs; iCell Cardiomyocytes) were obtained from FUJIFILM Cellular Dynamics (Madison, WI, USA) and maintained in maintenance medium provided with the cells (special formulation by FujiFilm). Cells were grown at 37°C in 5% CO_2_ and 90% humidity and checked for mycoplasma contamination.

### Plasmids and ebBRET biosensor constructs

Only human GPCRs and human Gα subunits were used in this study. An open reading frame of each full-length GPCR was cloned into pcDNA3.1(+) expression plasmid. Except when otherwise specified, GPCRs sequences were devoid of epitope tags.

Gα_s_-67-RlucII (Carr et al., 2014), Gα_i1_-loop-RlucII and GFP10-Gγ_1_ (Armando et al., 2014), Gα_i2_-loop-RlucII and βarrestin2-RlucII (Quoyer et al., 2013), Gα_oB_-99-RlucII (Mende et al., 2018), Gα_q_-118-RlucII (Breton et al., 2010), Gα_12_-136-RlucII and PKN-RBD-RlucII (Namkung et al., 2018), Gα_13_-130-RlucII (Avet et al., 2020), GFP10-Gγ_2_ (Gales et al., 2006), βarrestin1- RlucII (Zimmerman et al., 2012), rGFP-CAAX (Namkung et al., 2016), EPAC (Leduc et al., 2009), MyrPB-Ezrin-RlucII (Leguay et al., 2021), HA-β_2_AR (Lavoie et al., 2002), signal peptide-Flag-AT_1_ (Goupil et al., 2015) and EAAC-1 (Brabet et al., 1998) were previously described. Full-length, untagged Gα subunits, Gβ_1_ and Gγ_9_ were purchased from cDNA Resource Center. GRK2 was generously provided by Dr. Antonio De Blasi (Istituto Neurologico Mediterraneo Neuromed, Pozzilli, Italy).

To selectively detect G_i/o_ activation, a construct coding for aa 1-442 of Rap1 GTPase-activating protein (comprising a G_i/o_ binding domain) fused to Rluc8, was sequence- optimized, synthetized and subcloned at TopGenetech (St-Laurent, QC, Canada). From this construct, a RlucII tagged version of Rap1GAP (1-442) with a linker sequence (GSAGTGGRAIDIKLPAT) between Rap1GAP and RlucII was created by Gibson assembly in pCDNA3.1_Hygro (+) GFP10-RlucII, replacing GFP10. Three substitutions (i.e., S437A/S439A/S441A) were introduced into the Rap1GAP sequence by PCR-mediated mutagenesis. These putative (S437 and S439) and documented (S441) (McAvoy, Zhou, Greengard, & Nairn, 2009) protein kinase A phosphorylation sites were removed in order to eliminate any G_s_-mediated Rap1GAP recruitment to the plasma-membrane.

To selectively detect G_q/11_ activation, a construct encoding the G_q_ binding domain of the human p63 Rho guanine nucleotide exchange factor (p63RhoGEF; residues: 295-502) tagged with RlucII was done from IMAGE clones (OpenBiosystems; Burlington, ON, Canada) and subcloned by Gibson assembly in pCDNA3.1_Hygro (+) GFP10-RlucII, replacing GFP10. The G_q_ binding domain of p63RhoGEF and RlucII were separated by the peptidic linker ASGSAGTGGRAIDIKLPAT. N-term part containing palmitoylation sites maintaining p63 to plasma membrane and part of its DH domain involved in RhoA binding/activation (Aittaleb et al., 2010; Aittaleb, Nishimura, Linder, & Tesmer, 2011) are absent of the sensor.

To selectively detect G_12/13_ activation, a construct encoding the G_12/13_ binding domain of the human PDZ-RhoGEF (residues: 281-483) tagged with RlucII was done by PCR amplification from IMAGE clones (OpenBiosystems) and subcloned by Gibson assembly in pCDNA3.1_Hygro (+) GFP10-RlucII, replacing GFP10. The peptidic linker GIRLREALKLPAT is present between RlucII and the G_12/13_ binding domain of PDZ-RhoGEF. The sensor is lacking the PDZ domain of PDZ-RhoGEF involved in protein-protein interaction, as well as actin-binding domain and DH/PH domains involved in GEF activity and RhoA activation (Aittaleb et al., 2010).

### Transfection

For BRET experiments, HEK293 cells (1.2 mL at 3.5 × 10^5^ cells per mL) were transfected with a fixed final amount of pre-mixed biosensor-encoding DNA (0.57 μg, adjusted with salmon sperm DNA; Invitrogen) and human receptor DNA. Transfections were performed using a polyethylenimine solution (PEI, 1 mg/mL; Polysciences, Warrington, PA, USA) diluted in NaCl (150 mM, pH 7.0; 3:1 PEI/DNA ratio). Gelatin solution (1%; Sigma-Aldrich, Saint-Louis, Missouri) was used to stabilize DNA/PEI transfection mixes. Following addition of cells to the stabilized DNA/PEI transfection mix, cells were immediately seeded (3.5 × 10^4^ cells/well) into 96-well white microplates (Greiner Bio-one; Monroe, NC, USA) and maintained in culture for the next 48 h in DMEM containing 2% FBS and 1% PS. DMEM medium without L-glutamine (Wisent) was used for transfection of cells with mGluR to avoid receptor activation and desensitization. For Neutrophil-like differentiated HL-60 cells, cells were resuspended in electroporation medium (growth medium containing an extra 15 mM of HEPES pH 7.0) at 25 x 10^6^ cells/mL. Electroporation reactions were prepared by adding 50 µL of DNA mastermix (20 µg total of DNA adjusted with salmon sperm DNA, supplemented with 210 mM NaCl) to 200 µL of cell suspension and transferring into 0.4 cm gap electroporation cuvettes (Bio-Rad). The cells were electroporated at 350 µF/400 V using a Bio-Rad Gene Pulser II electroporation system, washed in electroporation medium, and seeded in 96-well plates at 0.8 x 10^6^ cells/well in 200 µL of growth medium. BRET assays were performed 6 hours post-electroporation. For iPSC Cardiomyocytes, cells were seeded in 96-well plates pretreated with fibronectin (10µg/ml 60 min; Sigma-Aldrich) at 3.5 × 10^4^ cells /well. After 48 h, attached iPSCs cells were transfected with the indicated biosensor components, using TransIT-LT1 reagent (Mirus; Madison, WI, USA), according to manufacturer recommendation. BRET assays were performed 48 hours after transfection.

For Ca^2+^ experiments, cells (3.5 x 10^4^ cells/well) were co-transfected with the indicated receptor, with or without Gα_15_ protein, using PEI and seeded in poly-ornithine coated 96- well clear-bottomed black microplates (Greiner Bio-one) and maintained in culture for the next 48 h.

For BRET-based imagery, cells (4 x 10^5^ cells/dish) were seeded into 35-mm poly-d-lysine-coated glass-bottom culture dishes (Mattek Corporation; Ashland, MA, USA) in 2 ml of fresh medium and incubated at 37°C in 5% CO_2_, 3 day before imaging experiments. Twenty-four hours later, cells were transfected with EMTA ebBRET biosensors and the indicated receptor (i.e., p63-RhoGEF-RlucII/rGFP-CAAX + Gα_q_ and GnRHR, Rap1GAP- RlucII/rGFP-CAAX + Gα_i2_ and D_2_ or PDZ-RhoGEF-RlucII/rGFP-CAAX + Gα_13_ and TPαR) using X-tremeGENE 9 DNA transfection reagent (3:1 reagent/DNA ratio; Roche) diluted in OptiMEM (Gibco) and maintained in culture for the next 48 h in DMEM containing 10% FBS and 1% PS.

### Bioluminescence Resonance Energy Transfer Measurement

Enhanced bystander BRET (ebBRET) was used to monitor the activation of each Gα protein, as well as βarrestin 1 and 2 recruitment to the plasma membrane. Gα_s_ protein activation was measured between the plasma membrane marker rGFP-CAAX and human Gα_s_-RlucII in presence of human Gβ_1_, Gγ_9_ and the tested receptor. Gα_s_ downstream cAMP production was determined using the EPAC biosensor and GPBA receptor. Gα_i/o_ protein family activation was followed using the selective-G_i/o_ effector Rap1GAP-RlucII and rGFP- CAAX along with the human Gα_i1_, Gα_i2_, Gα_oA_, Gα_oB_ or Gα_z_ subunits and the tested receptor. Gα_q/11_ protein family activation was determined using the selective-G_q/11_ effector p63- RhoGEF-RlucII and rGFP-CAAX along with the human Gα_q_, Gα_11_, Gα_14_ or Gα_15/16_ subunits and the tested receptor. Gα_12/13_ protein family activation was monitored using the selective-G_12/13_ effector PDZ-RhoGEF-RlucII and rGFP-CAAX in presence of either Gα_12_ or Gα_13_ and the tested receptor. The expression level of the Gα subunits was monitored by Western blot in HEK293 cells that endogenously expressed Gα_i1_, Gα_i2_, Gα_12_, Gα_13_, Gα_q_, Gα_11_, Gα_14_ and Gαs but not Gα_oA_, Gα_oB_, Gα_z_ and Gα_15_ (**Figure S6**). Gα_12/13_-downstream activation of the Rho pathway was measured using PKN-RBD-RlucII and rGFP-CAAX with the indicated receptor. βarrestin recruitment to the plasma membrane was determined using DNA mix containing rGFP-CAAX and βarrestin1-RlucII with GRK2 or βarrestin2-RlucII alone or with GRK2 and the tested receptor. Glutamate transporters EAAC-1 and EAAT-1 were systematically co-transfected with the mGluR to prevent receptor activation and desensitization by glutamate secreted in the medium by the cells (Brabet et al., 1998). All ligands were also tested for potential activation of endogenous receptors by transfecting the biosensors without receptor DNA. The G_z_/G_15_ biosensor consists of a combination of the following plasmids: rGFP-CAAX, Rap1GAP-RlucII, Gα_z_, p63-RhoGEF-RlucII and Gα_15_. For G protein activation detection using the BRET-based Gαβγ dissociation sensors, cells were co-transfected with untagged Gβ_1_ and Gα_q_-118-RlucII, Gα_12_-136-RlucII or Gα_13_-130- RlucII with GFP10-Gγ_1_, or Gα_i1_-loop-RlucII, Gα_i2_-loop-RlucII or Gα_oB_-99-RlucII with GFP10- Gγ_2_, along with the indicated receptor.

The day of the BRET experiment, cells were incubated in HBSS for 1 h at room temperature (RT). Cells were then co-treated with increasing concentrations of ligand (see **Supplementary Table 2** for details) and the luciferase substrate coelenterazine prolume purple (1 µM, NanoLight Technologies; Pinetop, AZ, USA) for 10 min at RT. Plates were read on a Synergy Neo microplate reader (BioTek Instruments, Inc.; Winooski, VT, USA) equipped with 410 ± 80 nm donor and 515 ± 30 nm acceptor filters or with a Spark microplate reader (Tecan; Männedorf, Switzerland) using the BRET^2^ manufacturer settings. The BRET signal (BRET²) was determined by calculating the ratio of the light intensity emitted by the acceptor over the light intensity emitted by the donor. To validate the specificity of the biosensor responses, cells were pretreated in the absence or presence of either the Gα_q_ inhibitor UBO-QIC (100 nM, 30 min; Institute for Pharmaceutical Biology of the University of Bonn, Germany), the Gα_i/o_ inhibitor PTX (100 ng/mL, 18 h; List Biological Laboratories, Campbell, California, USA) or the Gα_s_ activator CTX (0 to 200 ng/mL, 4h; Sigma-Aldrich) before stimulation with agonist. For Inverse agonist activity detection of A_1_ or CB_1_ receptors, cells were stimulated during 10 min with increasing concentrations of DPCPX or rimonabant, respectively. For ligand-cross receptor activation experiments, cells were pretreated for 10 min with increasing concentrations of antagonists or inverse agonist (eticlopride for D_2_, WB4101 for α_2A_AR, atropine for muscarinic receptors and AM-630 for CB_1_) before a 10 min stimulation with an EC_80_ concentration of the indicated agonist. BRET was measured as described above. For the safety target panel ligand screen using the combined G_z_/G_15_ sensor, basal ebBRET level was first measured 10 min following addition of coelenterazine prolume purple (1 µM) and ebBRET level was measured again following a 10 min stimulation with a single dose of the indicated ligand (1 μM for endothelin-1 and 10 μM for all other ligands). Technical replicates for each receptor were included on the same 96-well plate. For kinetics experiment of ET_A_ activation, basal BRET was measured during 150 sec before cells stimulation with either vehicle (DMSO) or 1 µM of endothelin-1 (at time 0 sec) and BRET signal was recorded each 30 sec during 3570 sec. For the validation of G_12/13_-mediated signal by new identified G_12/13_-coupled receptor using PKN- or Ezrin-based BRET biosensors, cells were pretreated or not with the Gα_q_ inhibitor YM-254890 (1 µM, 30 min; Wako Pure Chemical Industries; Wako Pure Chemical Industries (Fujifilm), Osaka, Japan) before agonist stimulation for 10 min. For G protein activation detection using the BRET-based Gαβγ dissociation sensors, and for titration experiments of either Gα proteins subunit with GEMTA sensors, GPCRs with GEMTA sensors or Effector-RlucII (p63-RhoGEF-RlucII for Gα_q/11_, Rap1GAP-RlucII for Gα_i/o_ or PDZ-RhoGEF-RlucII for Gα_12/13_) from GEMTA sensors, cells were stimulated with increasing concentrations of the indicated agonist in presence of prolume purple for 10 min before BRET measurement. For BRET in iPSC Cardiomyocytes and HL-60 cells, cells were incubated in Tyrode Hepes buffer (137 mM NaCl, 0.9 mM KCl, 1 mM MgCl_2_, 11.9 mM NaHCO_3_, 3.6 mM NaH_2_PO_4_, 25 mM HEPES, 5.5 mM D-Glucose and 1 mM CaCl_2_, pH 7.4) 30 min at RT before to be treated with increasing concentration of agonist for 15 min, using prolume purple (2 µM) as luciferase substrate, and BRET measured.

### BRET Data analyses and coupling efficiency evaluation

All BRET ratios were standardized using the equation below and represented as universal BRET (*u*BRET) values: *u*BRET = ((BRET ratio – A)/(B-A)) * 10 000. Constants A and B correspond to the following values:

A. = pre-established BRET ratio obtained from transfection of negative control (vector coding for RlucII alone);
B. = pre-established BRET ratio obtained from transfection of positive control (vector coding for a GFP10-RlucII fusion protein).

For a given signaling pathway, *u*BRET values at each agonist concentration were normalized as the % of the response obtained in the absence of agonist (vehicle) and concentration-response curves were fitted in GraphPad Prism 8 software using a four-parameter logistic nonlinear regression model. Results are expressed as mean ± SEM of at least three independent experiments.

A ligand-promoted response was considered real when the E_max_ value was ≥ to the mean + 2*SD of the response obtained in vehicle condition and that a pEC_50_ value could be determined in the agonist concentration range used to stimulate the receptor. Consequently, a score of 0 or 1 was assigned to each signaling pathway depending on an agonist’s ability to activate the tested pathway (0= no activation; 1= activation). In the case were responses associated to endogenous receptor were detectable, we considered as “distorted” and exclude all the responses observed in the presence of transfected receptor for which E_max_ was ≤ to 2*mean of the E_max_ value obtained with endogenous receptors or pEC_50_ was ≥ to 2*mean of the pEC_50_ value obtained with endogenous receptors. Consequently, a score of 0 was assigned for these distorted responses in radial graph representation (**Figure S7**) and dose-response curves were placed on a gray background in signaling signature profile panels (**Supplementary File 1**). Whenever transfected receptors produced an increase in E_max_ or a left-shift in pEC_50_ values compared to endogenous receptors, responses were considered ‘true’ and were assigned with a score of 1 for radial graph representation (**Figure S7**) and dose-response curves were placed on a yellow background in signaling signature profile panels to indicate a partial contribution of endogenous receptors (**Supplementary File 1**).

We used a double normalization of E_max_ and pEC_50_ values to compare the signaling efficiency obtained for the 100 GPCRs across all receptors and pathways. E_max_ and pEC_50_ values deduced from concentration-response curves were first normalized between 0 and 1 across receptors by ranking the receptors as a function of the receptor that most efficiently activate a given pathway and then using the activation value for the pathway (including G protein and βarrestin subtypes) that a given receptor most efficiently activate as a reference for the other pathways that can be activated by this receptor. This double normalization can be translated in the following formalized equation:

- STEP1: For each receptor and for each pathway:

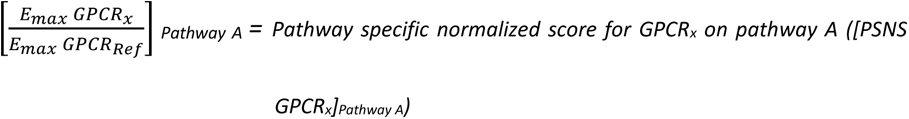

Where: GPCR_x_ is receptor being analyzed, GPCR_Ref_ is the receptor giving greatest E_max_ on pathway A of all receptors studied (i.e., reference receptor for pathway A). A PSNS was determined for every receptor and every pathway coupled to that receptor.

- STEP2: For any given receptor:

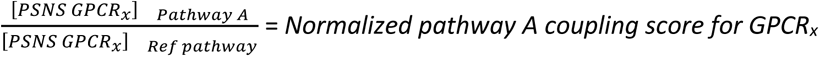

Where: [PSNS GPCR_x_] _Pathway A_ is the pathway specific normalized score for GPCR_x_ on pathway A, and [PSNS GPCR_x_] _Ref pathway_ is the pathway specific normalized score for the pathway giving the highest PSNS for GPCR_x_ (i.e., reference pathway for GPCR_x_).

For the safety target panel ligand screen using the combined G_z_/G_15_ sensor, the fold ligand-induced stimulation was calculated for each receptor by dividing the BRET ratio after ligand addition (measured at 10 minutes post stimulation) by the basal BRET ratio prior to receptor stimulation. Activation thresholds were defined as the mean + 2*SD of the ligand-stimulated response obtained with receptor-null cells expressing only the combined G_z_/G_15_ sensor.

### Ca^2+^ mobilization assay

The day of experiment, cells were incubated with 100 μL of a Ca^2+^-sensitive dye-loading buffer (FLIPR calcium 5 assay kit, Molecular Devices; Sunnyvale, CA, USA) containing 2.5 mM probenecid (Sigma-Aldrich) for 1 h at 37°C in a 5% CO_2_ incubator. During a data run, cells in individual wells were exposed to an EC_80_ concentration of agonist, and fluorescent signals were recorded every 1.5 s for 3 min using the FlexStation II microplate reader (Molecular Devices). For receptors that also activate other G_q/11_ family members, cells were pretreated with the G_q/11_ inhibitor YM-254890 (1 µM, 30 min) before agonist stimulation. Gα_15_ is resistant to inhibition by YM-254890, thus allowing to measure Ca^2+^ responses generated specifically by Gα_15_.

### BRET-based imaging

BRET images were obtained as previously described (Kobayashi, Picard, Schonegge, & Bouvier, 2019). Briefly, the day of imaging experiment, cells were carefully rinsed with HBSS, and images were acquired before and after agonists addition (100 nM for GnRH and U46619, and 1 µM for dopamine) diluted in HBSS in presence of the luciferase substrate coelenterazine prolume purple (20 µM).

Images were recorded using an inverted microscope (Nikon Eclipse Ti-U) equipped with x60 objective lens (Nikon CFI Apochromat TIRF) and EM-CCD camera (Nuvu HNu 512). Measurements were carried out in photon counting mode with EM gain 3,000. Exposure time of each photon counting was 100 ms. Successive 100 frames were acquired alternatively with 480 nm longpass filter (acceptor frames) or without filter (total luminescence frames), and integrated. Image integrations were repeated 5 times and 500 frames of acceptor and total luminescence were used to generate each image.

BRET values were obtained by dividing acceptor counts by total luminescence counts pixelwise. BRET values from 0.0 to 0.5 were allocated to ‘jet’ heatmap array using MATLAB 2019b. Brightness of each pixel was mapped from the signal level of total luminescence image. 0% and 99.9% signal strength were allocated to the lowest and highest brightness to exclude the influence of defective pixels with gamma correction factor of 2.0.

The movies were generated using ImageJ 1.52a. Frame rate is 3 frames/sec, and frame interval is 100 sec. The field of view of the movie is 137 um x 137 um.

### Western blot analysis

Cells were transfected or not with the indicated biosensors mix as previously described and whole-cell extracts were prepared 48 h later. Briefly, cells were washed with ice-cold PBS and lysed in a buffer containing 10 mM Tris buffer (pH 7.4), 100 mM NaCl, 1 mM EDTA, 1 mM EGTA, 0.1% SDS, 1% Triton X-100, 10% Glycerol supplemented with protease inhibitors cocktails (Thermo Fisher Scientific). Cell lysates were centrifuged at 13,000 × g for 30 min at 4°C. Equal amounts of proteins were separated by SDS-PAGE and transferred onto polyvinylidene fluoride membrane. The membranes were blocked in (incubation 1 hr at room temperature in PBS 0.1% Tween-20, 5% BSA) and successively probed with primary antibody and appropriate goat secondary antibodies coupled to horseradish peroxidase (described in table below). Western blots were visualized using enhanced chemiluminescence and detection was performed using a ChemiDoc MP Imaging System (BioRad). Relative densitometry analysis on protein bands was performed using MultiGauge software (Fujifilm). Results were normalized against control bands.

| Target | Dilution | Species | Class | Reference | Manufacturer |
| --- | --- | --- | --- | --- | --- |
| Gαi1 (I-20) | 1:500 | Rabbit | polyclonal | #sc-391 | Santa Cruz |
| Gαi2 (T-19) | 1:500 | Rabbit | polyclonal | #sc-7276 | Santa Cruz |
| Gαo (K-20) | 1:500 | Rabbit | polyclonal | #sc-387 | Santa Cruz |
| Gαz | 1:1,000 | Rabbit | monoclonal | # ab154846 | Abcam |
| Gαs (K-20) | 1:500 | Rabbit | polyclonal | #sc-823 | Santa Cruz |
| Gα12 (S-20) | 1:500 | Rabbit | polyclonal | #sc-409 | Santa Cruz |
| Gα13 (A-20) | 1:500 | Rabbit | polyclonal | #sc-410 | Santa Cruz |
| Gαq (E-17) | 1:500 | Rabbit | polyclonal | #sc-393 | Santa Cruz |
| Gα11 (C-terminal) | 1:500 | Rabbit | Polyclonal | #SAB2109181 | Sigma-Aldrich |
| Gα14 | 1:500 | Rabbit | Polyclonal | #SAB4300771 | Sigma-Aldrich |
| Gα15 | 1:5,000 | Rabbit | Polyclonal | #PA1-29022 | ThermoFisher scientific (Pierce) |
| βactin | 1:5,000 | Mouse | Monoclonal | #A5441 | Sigma-Aldrich |
| Anti-rabbit HRP-coupled | 1:5,000 | Donkey | Polyclonal | #NA934 | GE Healthcare |
| Anti-mouse HRP-coupled | 1:10,000 | Sheep | Polyclonal | #NA931 | GE Healthcare |

### Statistical Analyses

Curve fitting and statistical analyses were performed using GraphPad Prism 9.3 software and methods are described in the legends of the figures. Significance was determined as p < 0.05.

## Acknowledgments

We thank Shane C. Wright for scientific discussion and Monique Lagacé for critical reading of the manuscript. The authors are grateful to the funding from Bristol-Myers Squibb that supported the detection of Gα proteins by endogenous receptors in iPSC cardiomyocytes and promyelocytic HL-60 cells.

## Author contributions

Conceptualization: CA, AM, BB, CLG, XL, MB Methodology: CA, AM, BB, CLG, MB

Investigation: CA, AM, BB, CN, HK, FG, MH, VL, SS-O, MC, MH, SM

Formal Analysis: CA, AM, ASH, DEG, MB Resources: AM, EF, J-PF, SS, XL, MB

Supervision: MH, XL, DEG, MB Funding Acquisition: SS, DEG, MB

Writing: CA, AM, DEG, MB; All coauthors revised the manuscript

## Data and materials availability

Further information and requests for resources and reagents should be directed to and will be fulfilled upon reasonable request by the Lead Contact, Michel Bouvier.

The ebBRET sensors used in the study are protected by patent applications and have been licensed to Domain Therapeutics. Inquiries for potential commercial use should be addressed to. For non-commercial academic use, the sensors can be obtained freely under material transfer agreement upon request to.

Heatmaps in **Figure 3** were generated using custom python scripts. Scripts are available from the co-corresponding author, David E. Gloriam on request.

## Supplementary Information

### Supplementary Figures

**Figure S1.**
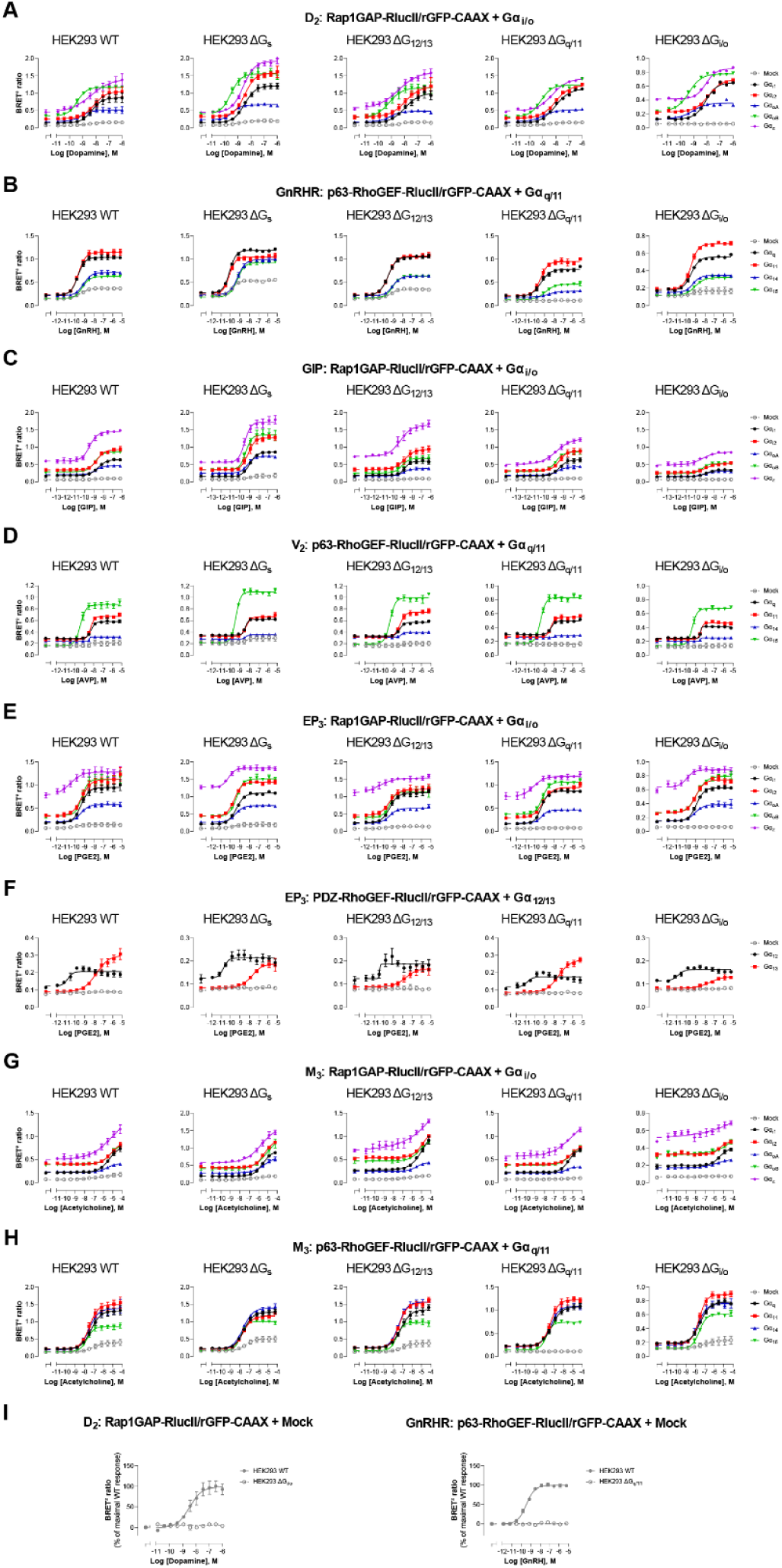
Influence of endogenous G proteins. Dose-response curves elicited in parental (WT) HEK293 cells or devoid of G_s_ (ΔG_s_), G_12/13_ (ΔG_12/13_), G_q/11_ (ΔG_q/11_) or G_i/o_ (ΔG_i/o_) proteins, transfected with the indicated receptor (D_2_, GnRHR, GIP, V_2_, EP_3_ or M_3_) and one of the Gα_i/o_, Gα_q/11_ or Gα_12/13_ activation sensors, along with the indicated Gα subunits. Mock condition corresponded to the response elicited in absence of heterologously expressed Gα subunits (i.e., endogenous G proteins effect). Data are means ± SEM of 3-5 independent experiments performed in one replicate and are expressed as BRET^2^ ratio. Data presented in **I** are the same as in **A**-**B**, but with results expressed as % of maximal response elicited by endogenous G proteins (mock) in WT cells.

**Figure S2.**
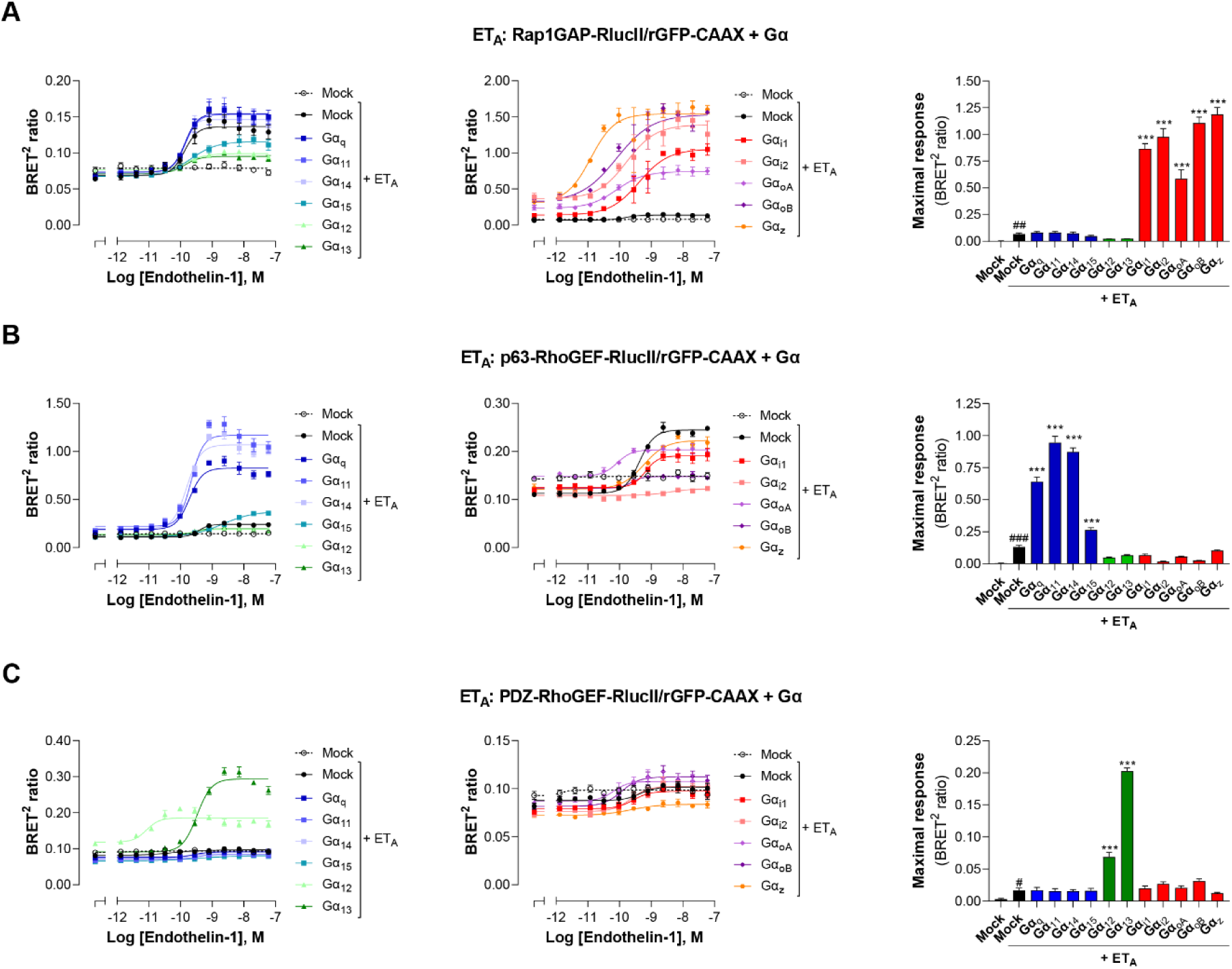
Validation of EMTA ebBRET-based sensors selectivity for each Gα subunit families. HEK293 cells were transfected with the ET_A_ receptor and Gα_i/o_ (A), Gα_q/11_ (B) or Gα_12/13_ (C) activation sensors along with each Gα subunit or control DNA (Mock) as control for response obtained with endogenous Gα proteins. Dose-response curves in response to endothelin-1 are shown (*left and central*), as well as maximal responses obtained with each Gα subunit. Data are means ± SEM of 3 independent experiments performed in one replicate and are expressed as BRET^2^ ratio. Unpaired t-test: ^#^p < 0.05, ^##^p < 0.01 and ^###^p < 0.001 compared to Mock (without receptor) and One-Way ANOVA test: *p < 0.05, **p < 0.01 and ***p < 0.001 compared to Mock + ET_A_.

**Figure S3.**
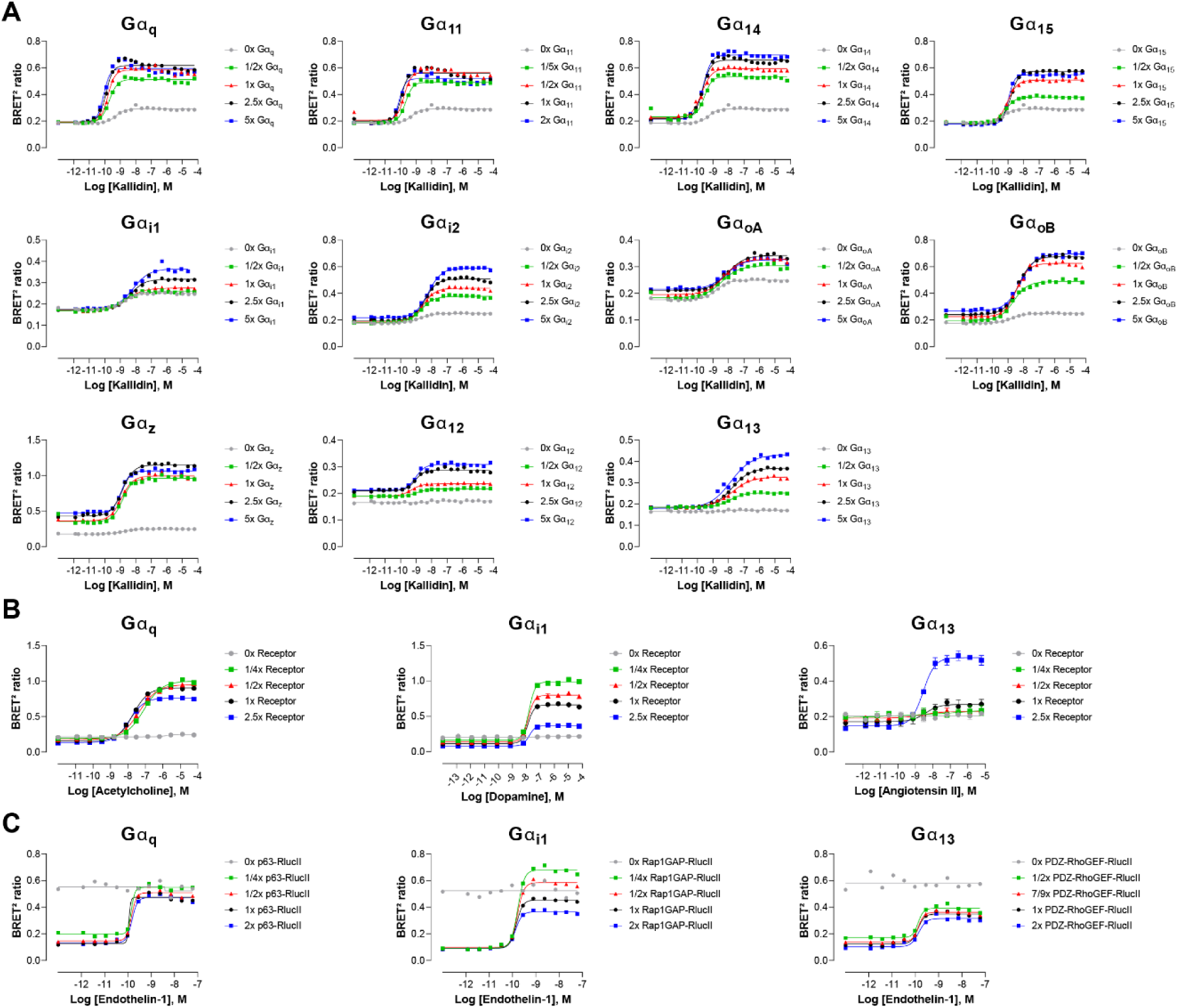
Influence of G protein, GPCR or effector-RlucII level expression. (**A**) Dose-response curves elicited in HEK293 cells transfected with the B_2_ receptor and one of the Gα_q/11_, Gα_i/o_ or Gα_12/13_ activation sensors, along with increasing quantity of the indicated Gα subunits. Data represent a representative experiment (**B**) Dose-response curves elicited in HEK293 cells transfected with increasing quantity of the M_3_, D_2_ or AT_1_ receptors and the Gα_q/11_, Gα_i/o_ or Gα_12/13_ activation sensors, along with the indicated Gα subunits. (**C**) Dose-response curves elicited in HEK293 cells transfected with the ET_A_ receptor and increasing quantity of effector-RlucII (p63-RhoGEF for Gα_q/11_, Rap1GAP for Gα_i/o_ or PDZ-RhoGEF for Gα_12/13_), along with rGFP-CAAX and the indicated Gα subunits. Data are means ± SEM of 3 independent experiments performed in one replicate and are expressed in BRET^2^ ratio.

**Figure S4.**
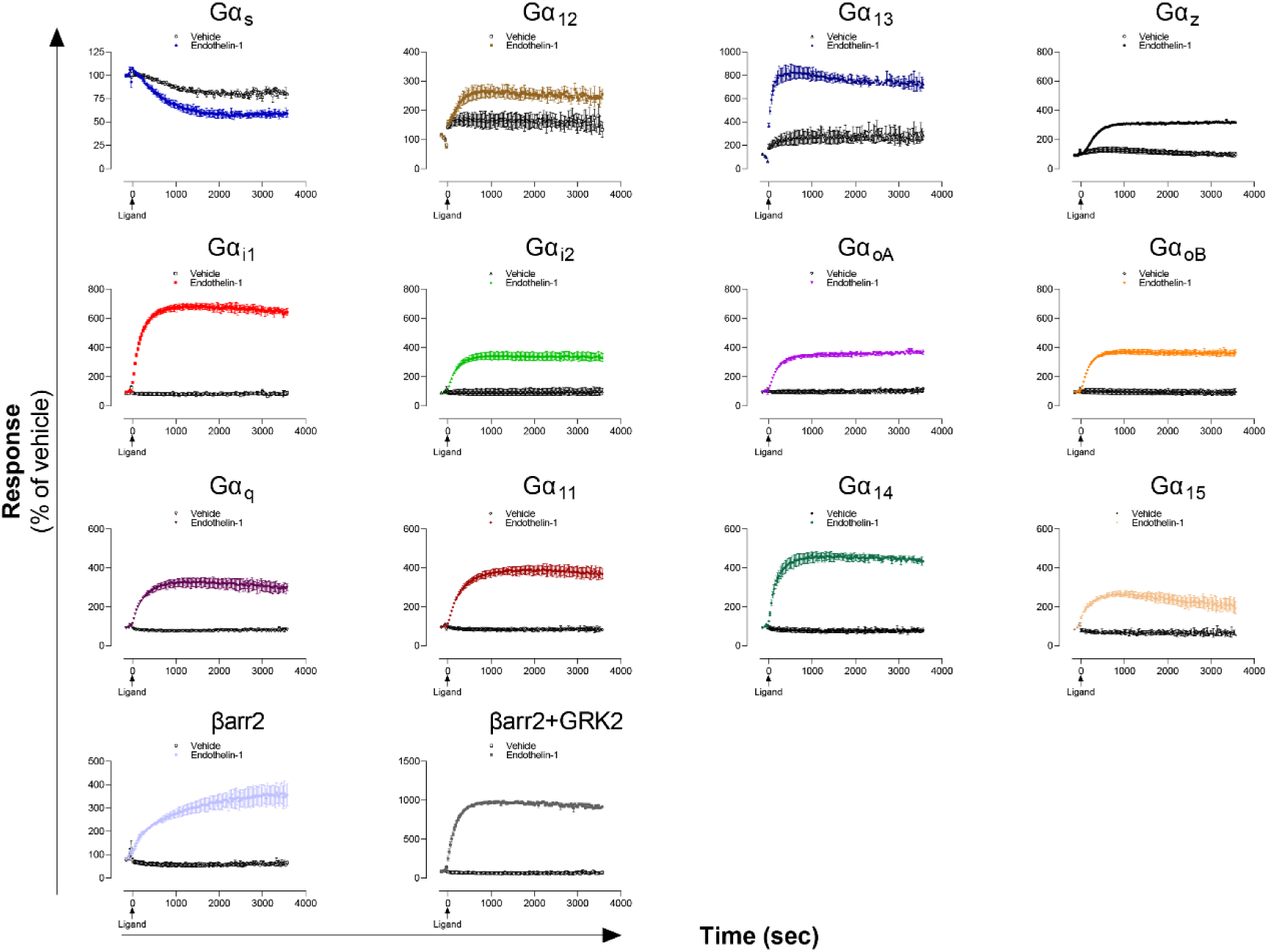
Kinetics of Gα proteins and βarrestins recruitment promoted by the ET_A_ receptor. Kinetics of activation of the indicated pathways following stimulation with vehicle or Endothelin-1 in HEK293 cells expressing the ET_A_ receptor. Data are means ± SD of two replicates of a representative experiment from three independent experiments and are expressed in % of the respective basal response (determined before ligand addition at t=0 sec).

**Figure S5.**
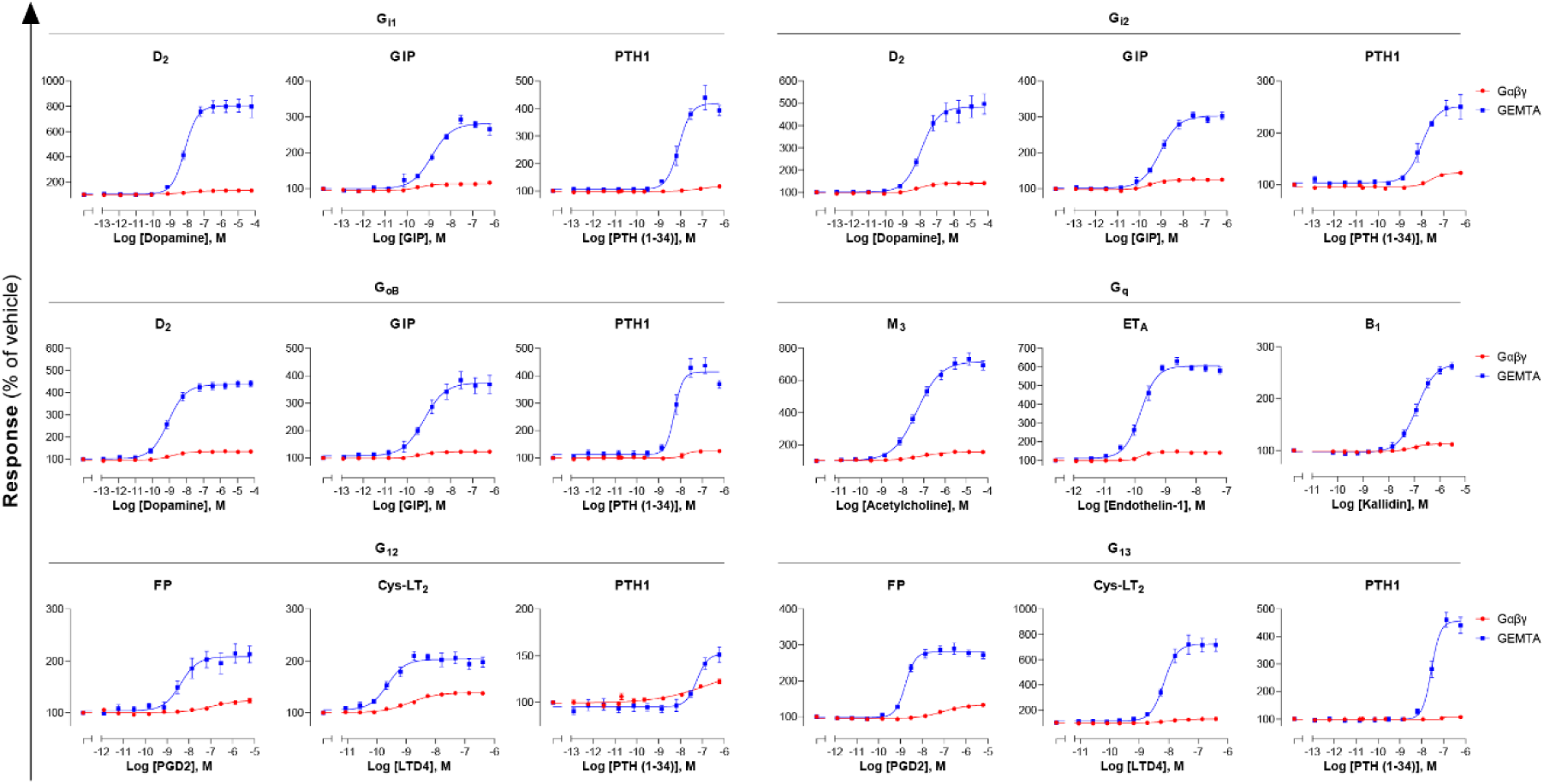
Comparison of EMTA platform and G protein activation BRET assay based on Gαβγ dissociation. Dose-response curves elicited in HEK293 cells transfected with the indicated receptor (D_2_, GIP, PTH1, M_3_, ET_A_, B_1_, FP or Cys-LT_2_) and one of the Gα_i/o_, Gα_q/11_ or Gα_12/13_ EMTA activation sensors, along with the indicated Gα subunits, or the BRET-based Gαβγ dissociation sensors (Gα-RlucII and GFP10-Gγ_1_ for Gα_q_, Gα_12_ and Gα_13_ or GFP10-Gγ_2_ for Gα_i1_, Gα_i2_ and Gα_oB_, with untagged Gβ_1_). Data are means ± SEM from 3-7 independent experiments performed in one replicate and results are expressed in % of the response obtained for cells treated with vehicle.

**Figure S6.**
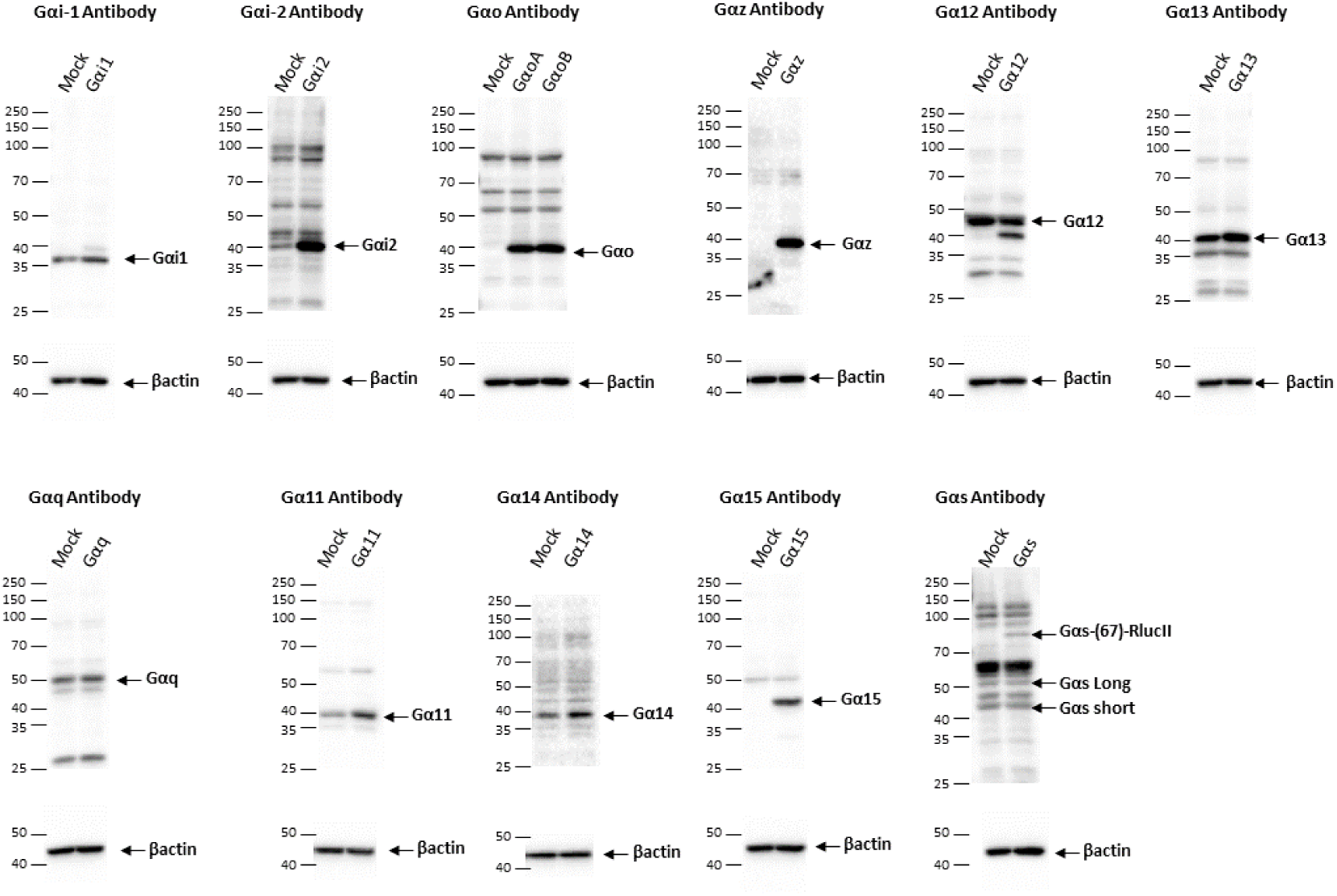
Western blot of G protein level expression in cells transfected with the EMTA ebBRET platform. G protein expression level detection in HEK293 cells transfected with the Gα_i/o_, Gα_12/13_, Gα_q/11_ or Gα_s_ activation sensors along with the indicated Gα protein or control DNA (Mock). Representative immunoblots of 3 independent experiments are shown.

**Figure S7.**
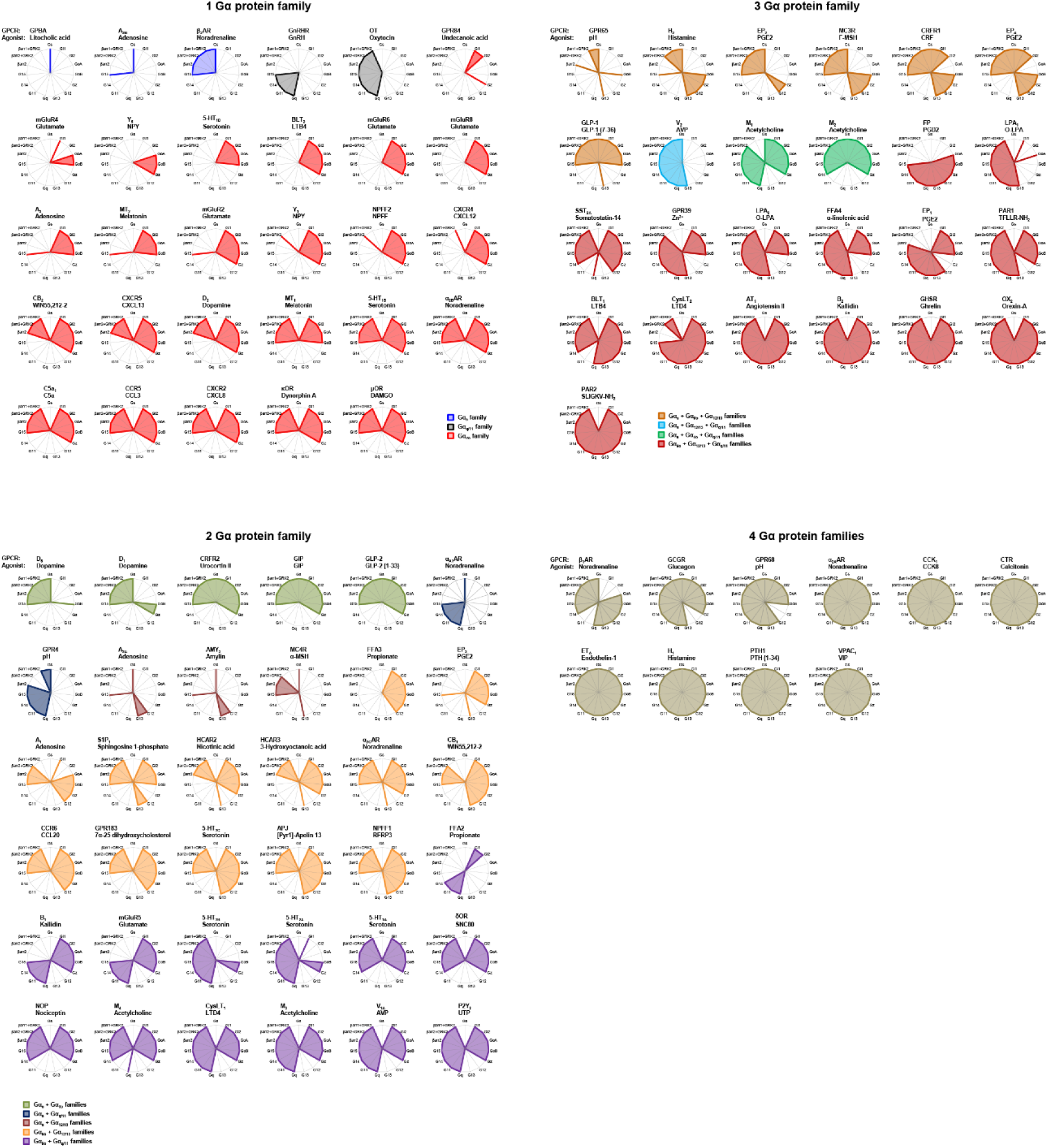
Receptor-specific signaling signatures. E_max_ values derived from concentration-response curves generated on 100 receptors using the 15 ebBRET-based assay are represented as radial graphs. A score of 0 indicates no coupling to a given pathway, whereas a score of 1 indicates a coupling. Receptors are rearranged according to the number of G protein families activated. Gα_15_ has been considered apart from the G_q/11_ family due to its promiscuous nature. See **Supplementary File 1** that shows the dose-response curves of the 100 receptors for the 15 different pathways.

**Figure S8.**
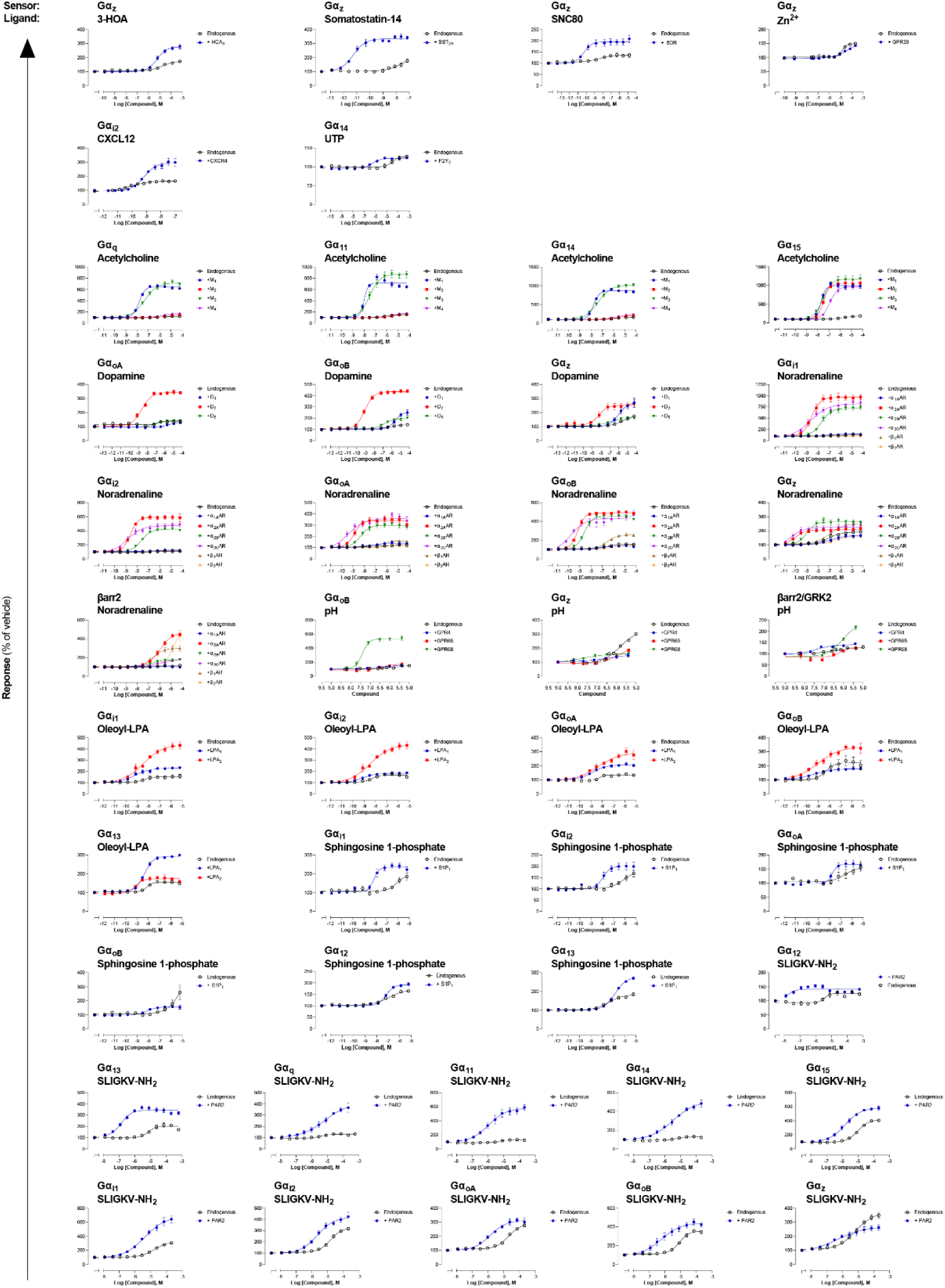
Detection of endogenous receptor-mediated responses with the EMTA ebBRET platform in HEK293 cells. Comparison of concentration-response curves elicited by the indicated ligand for a specific pathway, following the stimulation of HEK293 cells expressing endogenous or heterologously expressed receptors. The data presented refer to the ligands where a signal was detected on non-transfected cells (endogenous expression) (See **Supplementary File 1** for the curves on light gray and yellow background). Data are the mean ± SEM of at least 3 independent experiments performed in one replicate and expressed in % of the response obtained for cells treated with vehicle.

**Figure S9.**
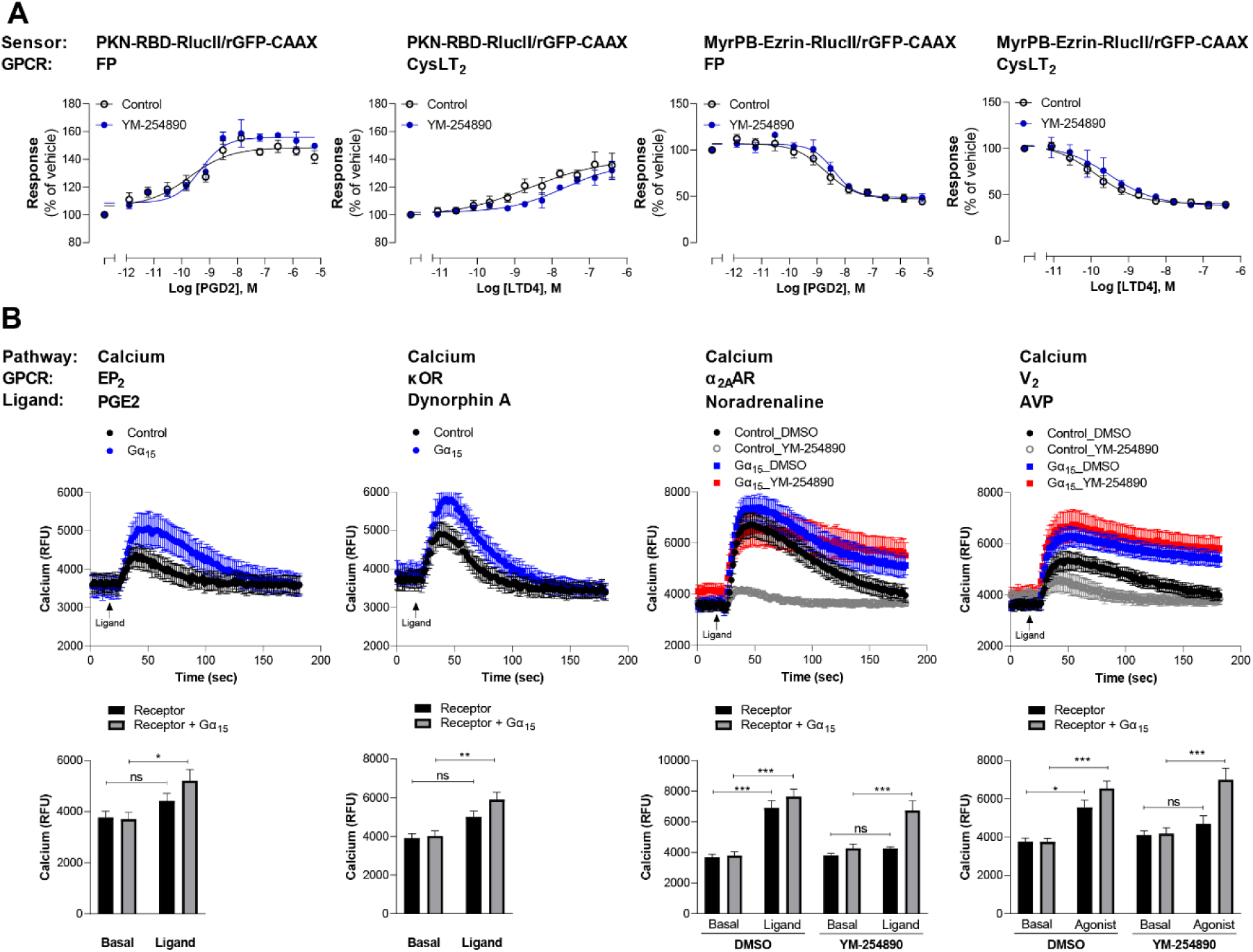
Validation of G_12/13_ and G_15_ signaling for the newly characterized GPCRs. (**A**) Validation of G_12/13_- mediated signal using Rho and Ezrin activation sensors. HEK293 cells expressing FP or CysLT_2_ receptors and the PKN-RBD-RlucII or MyrPB-Ezrin-RlucII/rGFP-CAAX sensors were pretreated or not with the Gα_q_ inhibitor YM-254890 and then stimulated with increasing concentrations of respective ligand. Data are means ± SEM from 3-5 independent experiments performed in one replicate and expressed in % of vehicle treated cells. (B) Validation of Gα_15_-mediated signal by measuring calcium production. *Top*: Kinetics of calcium release induced by the indicated ligand in HEK293 cells expressing the indicated receptor, alone or with Gα_15_ subunit. For receptors that also couple to other G_q/11_ family members, cells were pretreated with DMSO or the Gα_q_ inhibitor YM-254890. *Bottom*: The peak of calcium production obtained from kinetics were compared to the basal level of calcium (determined between 0 and 17 sec). Data are means ± SEM from 5- 7 independent experiments performed in one replicate and expressed in relative fluorescence unit (RFU). Two Way ANOVA test: *p < 0.05, **p < 0.01 and ***p < 0.001 compared to respective basal calcium level. ns: not significant.

**Figure S10.**
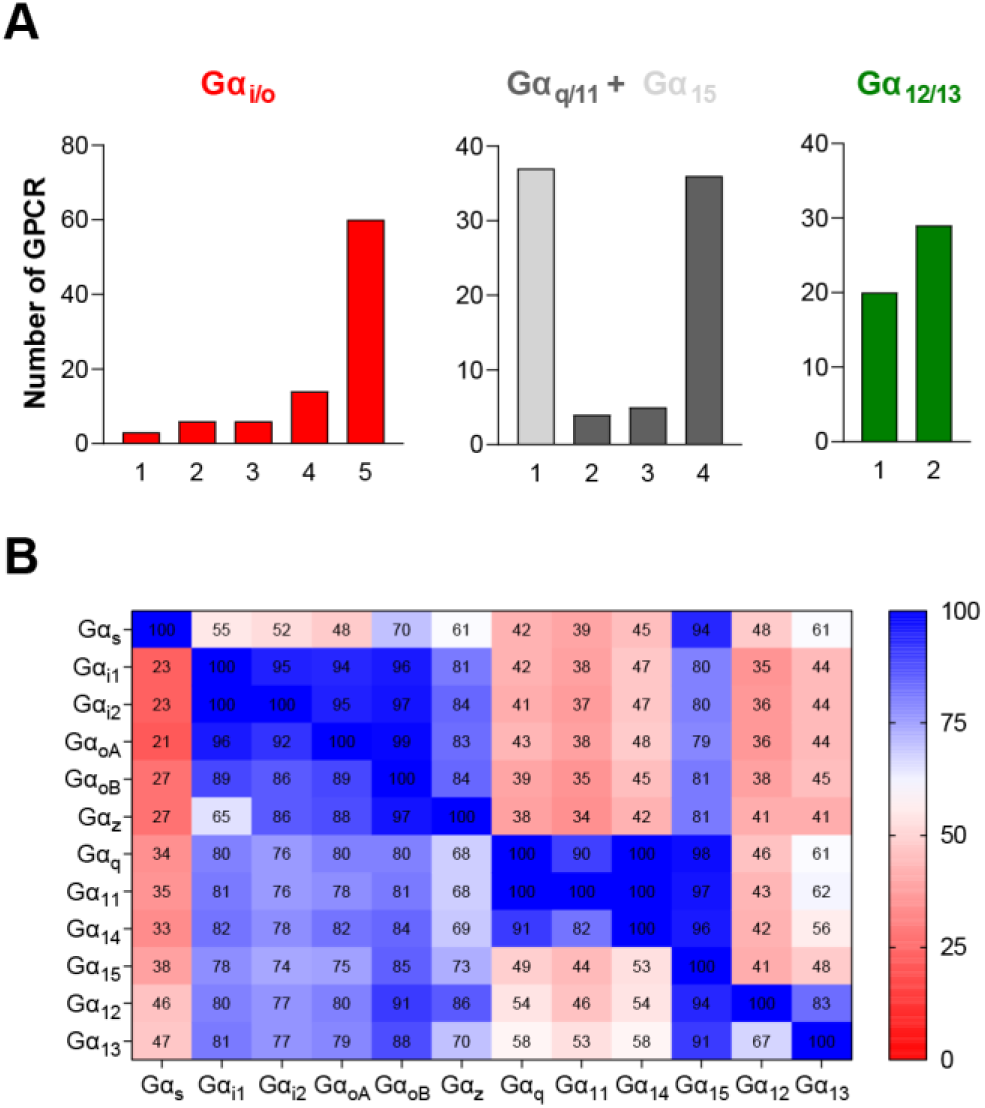
G protein subtypes distribution across the 100 GPCRs profiled with the EMTA ebBRET-based platform. (**A**) Number of receptors that can couple to 1 to 5 of the different subtypes from each G protein family. (**B**) % of receptors activating a specific G protein subtype (Y axis) that also activate another G protein subtype (X axis).

**Figure S11.**
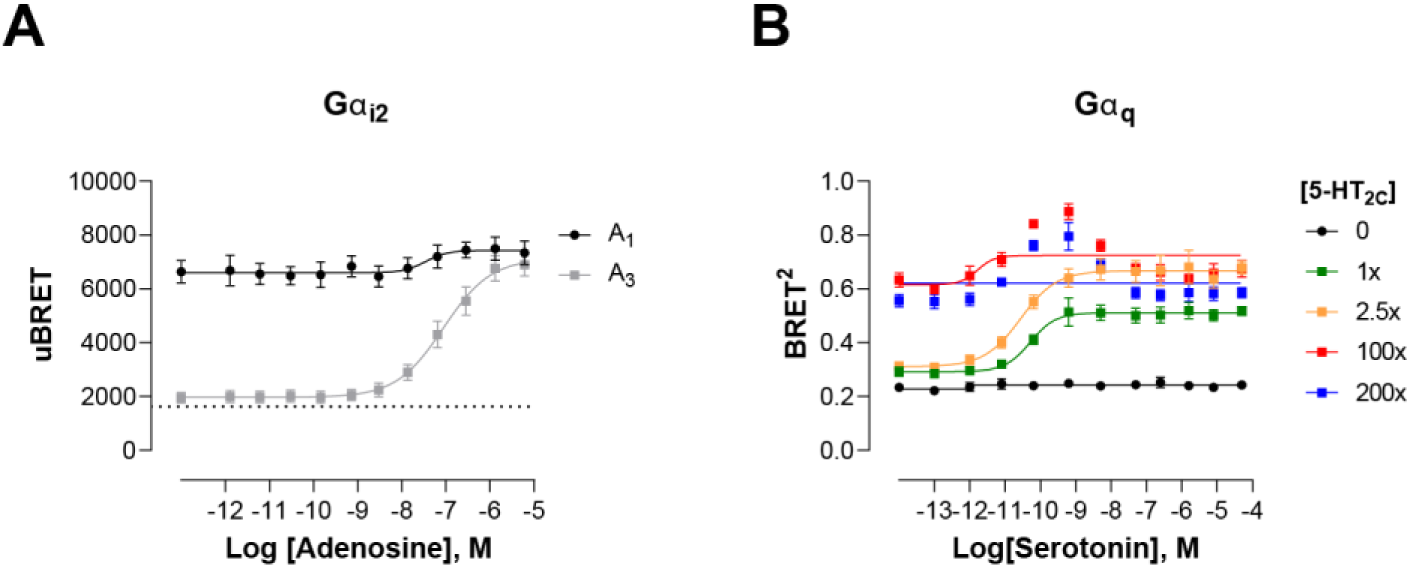
Modulation of ligand-promoted response detected by EMTA ebBRET platform by receptor constitutive activity. (**A**) Concentration-response curves of Gα_i2_ activation elicited by adenosine in HEK293 cells transfected with the Rap1GAP-RlucII/rGFP-CAAX sensors with untagged Gα_i2_ and A_1_ or A_3_ receptors. Basal level of G_i2_ activation detected by the GEMTA sensor in absence of heterologous receptor expression is represented by the interrupted line. Data are expressed as uBRET ratio and are means ± SEM of 4 independent experiments performed in one replicate. (**B**) Concentration-response curves of Gα_q_ activation elicited by serotonin in HEK293 cells transfected with the p63-RlucII/rGFP-CAAX sensors with untagged Gα_q_ and increasing amount of 5-HT_2C_ receptor plasmid. Data are expressed as BRET ratio and are means ± SEM of 4 independent experiments performed in one replicate.

**Figure S12.**
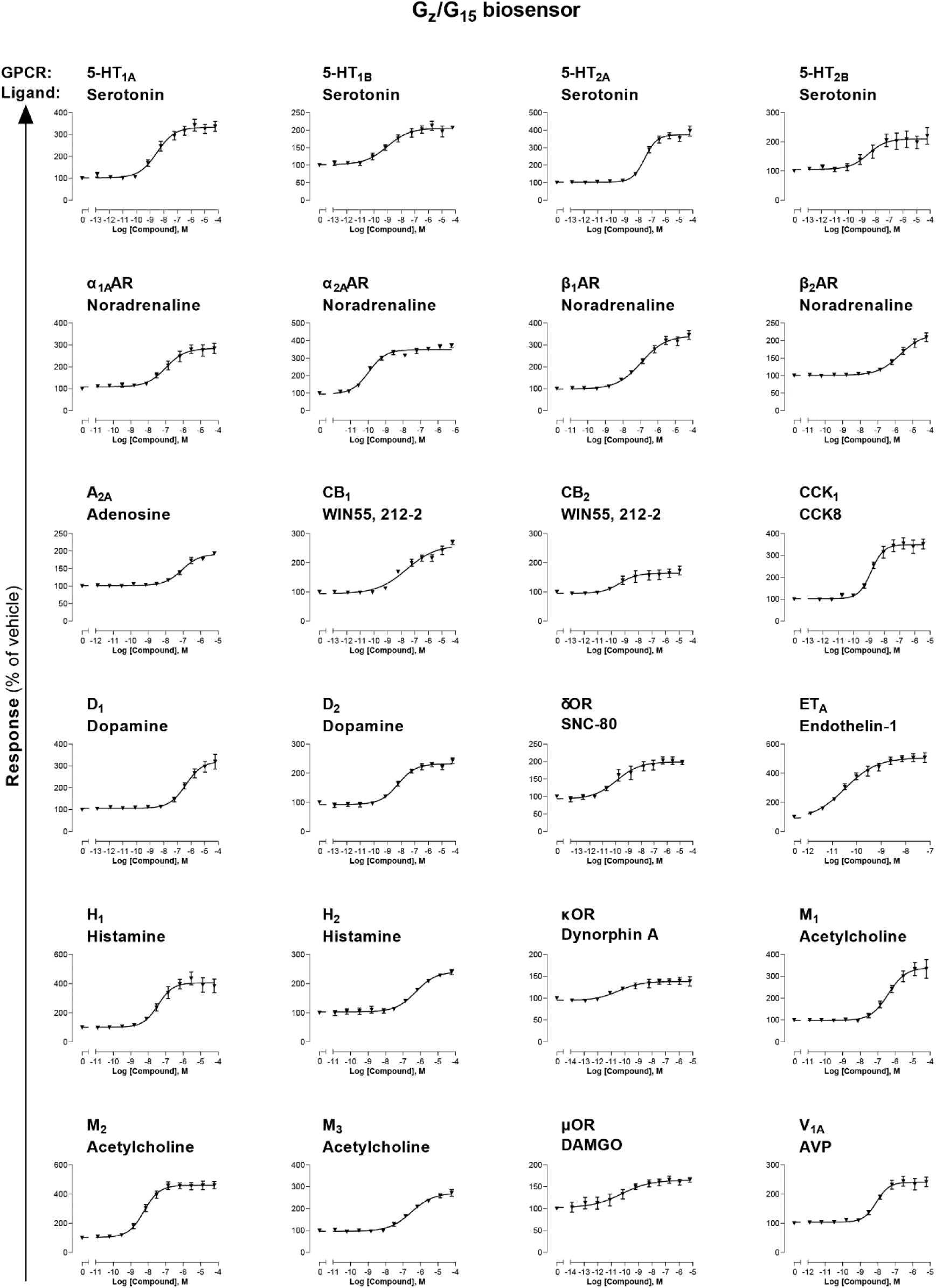
Combined G_z_/G_15_ biosensor. HEK293 cells transfected with the Rap1GAP-RlucII/p63-RhoGEF- RlucII/rGFP-CAAX sensors along with Gα_z_ and Gα_15_ subunits and the indicated untagged receptor were stimulated with increasing concentrations of the indicated ligand. Data are means ± SEM from 3-5 independent experiments performed in one replicate and results are expressed in % of vehicle treated cells.

**Figure S13.**
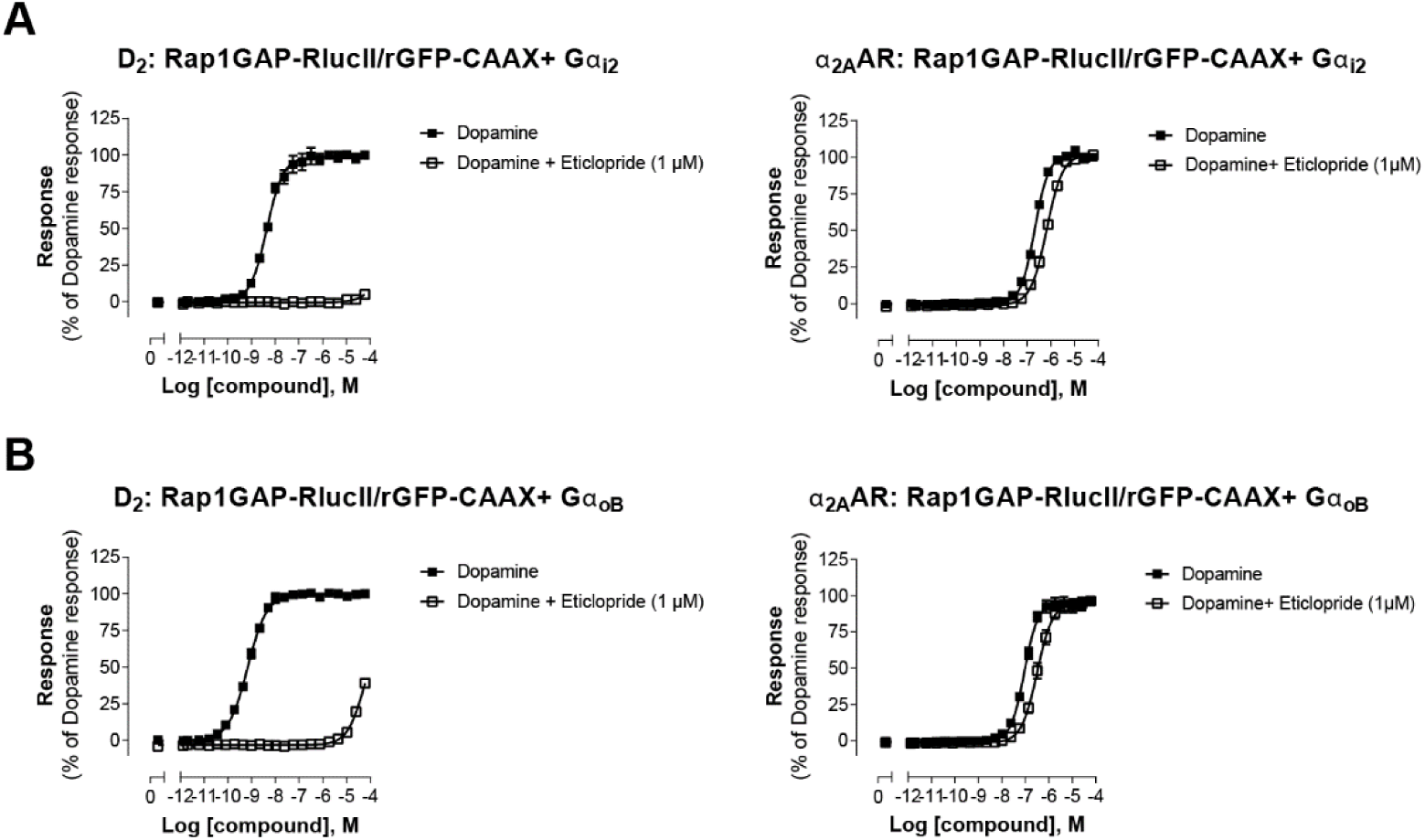
Validation of direct activation of α_2A_AR by Dopamine. HEK293 cells expressing D_2_ or α_2A_AR and the Gα_i2_ (**A**) or the Gα_oB_ (**B**) sensors were pretreated or not with the selective D_2_-family antagonist eticlopride, before stimulation with increasing concentrations of dopamine. Data are means ± SEM from 2- 4 independent experiments performed in one replicate and expressed in % of the response elicited by dopamine.

## Supplementary Materials

**Supplementary File 1. Signaling profiles of 100 therapeutically-relevant human GPCRs using the EMTA ebBRET platform.** Concentration-response curves in HEK293 cells expressing the indicated biosensor after stimulation of heterologously expressed receptor with the indicated ligand. Data are means ± SEM from at least 3 independent experiments and expressed in % of the response obtained for cells treated with vehicle. For ligands that elicited endogenous receptor-mediated responses (curves with light gray and yellow background for responses similar to and better responses than those obtained with the endogenous receptors, respectively), curves from cells expressing endogenous or heterologously expressed receptors are shown in **Figure S8**.

**Supplementary Table 1.** (**A**) **Absolute pEC_50_ values of responses elicited in WT *vs.* Knockout Gα protein background cells.** pEC_50_ values deduced from dose-response curves for various receptor tested in parental (WT) HEK293 cells or devoid of G_s_, G_12/13_, G_q/11_ or G_i/o_ proteins are related to **Figure S1**. (**B**) **Absolute pEC_50_ values of responses elicited in cells transfected with different amounts of Gα proteins.** pEC_50_ values deduced from dose-response curves obtained following Gα subunit titration in HEK293 cells transfected with GEMTA sensors and related to **Figure S3A**. (**C**) **Absolute pEC_50_ values of responses elicited in cells transfected with different amounts of receptors.** pEC_50_ values deduced from dose-response curves obtained following ET_A_ titration in HEK293 cells transfected with GEMTA sensors and related to **Figure S3B.** (**D**) **Absolute pEC_50_ values of responses elicited in cells transfected with different amounts of Effector-RlucII**. pEC_50_ values deduced from dose-response curves obtained following Effector-RlucII titration in HEK293 cells transfected with GEMTA sensors and related to **Figure S3C**.

**Supplementary Table 2. List of tested receptors and ligands, along with the raw E_max_, absolute pEC_50_ and their corresponding double normalized (dnor) values.** The E_max_ (in % of vehicle response) and absolute pEC_50_ values deduced from concentration-response curves for the 100 GPCRs tested as well as the double normalized E_max_ and pEC_50_ values calculated are related to Supplementary File 1 and Figure 3, respectively.

**Supplementary Table 3. Comparison of G protein couplings identified with EMTA platform and other datasets.** Comparison of G protein couplings identified with EMTA platform and TGF-α shedding assay in Inoue *et al*., 2019 (**A**) or reported in GtP database (**B**).

**Video 1. BRET-based imagery of p63-RhoGEF-RlucII recruitment to plasma membrane upon GnRHR activation.** HEK293 cells expressing the p63-RhoGEF-RlucII/rGFP-CAAX sensors with Gα_q_ and GnRHR were stimulated with GnRH. BRET levels (the ratio of the acceptor photon count to the total photon count) are expressed as a color code (lowest being black and purple, and highest being red and white).

**Video 2. BRET-based imagery of Rap1GAP-RlucII recruitment to plasma membrane upon D_2_ activation.** HEK293 cells expressing the Rap1GAP-RlucII/rGFP-CAAX sensors with Gα_i2_ and D_2_ were stimulated with dopamine. BRET levels (the ratio of the acceptor photon count to the total photon count) are expressed as a color code (lowest being black and purple, and highest being red and white).

**Video 3. BRET-based imagery of PDZ-RhoGEF-RlucII recruitment to plasma membrane upon TPαR activation.** HEK293 cells expressing the PDZ-RhoGEF-RlucII/rGFP-CAAX + Gα_13_ and TPαR were stimulated with U46619. BRET levels (the ratio of the acceptor photon count to the total photon count) are expressed as a color code (lowest being black and purple, and highest being red and white).

Since videos cannot be accepted on the bioRvix site, we have generated figures that show the first and last image of each video that have been submitted to the journal (see below).

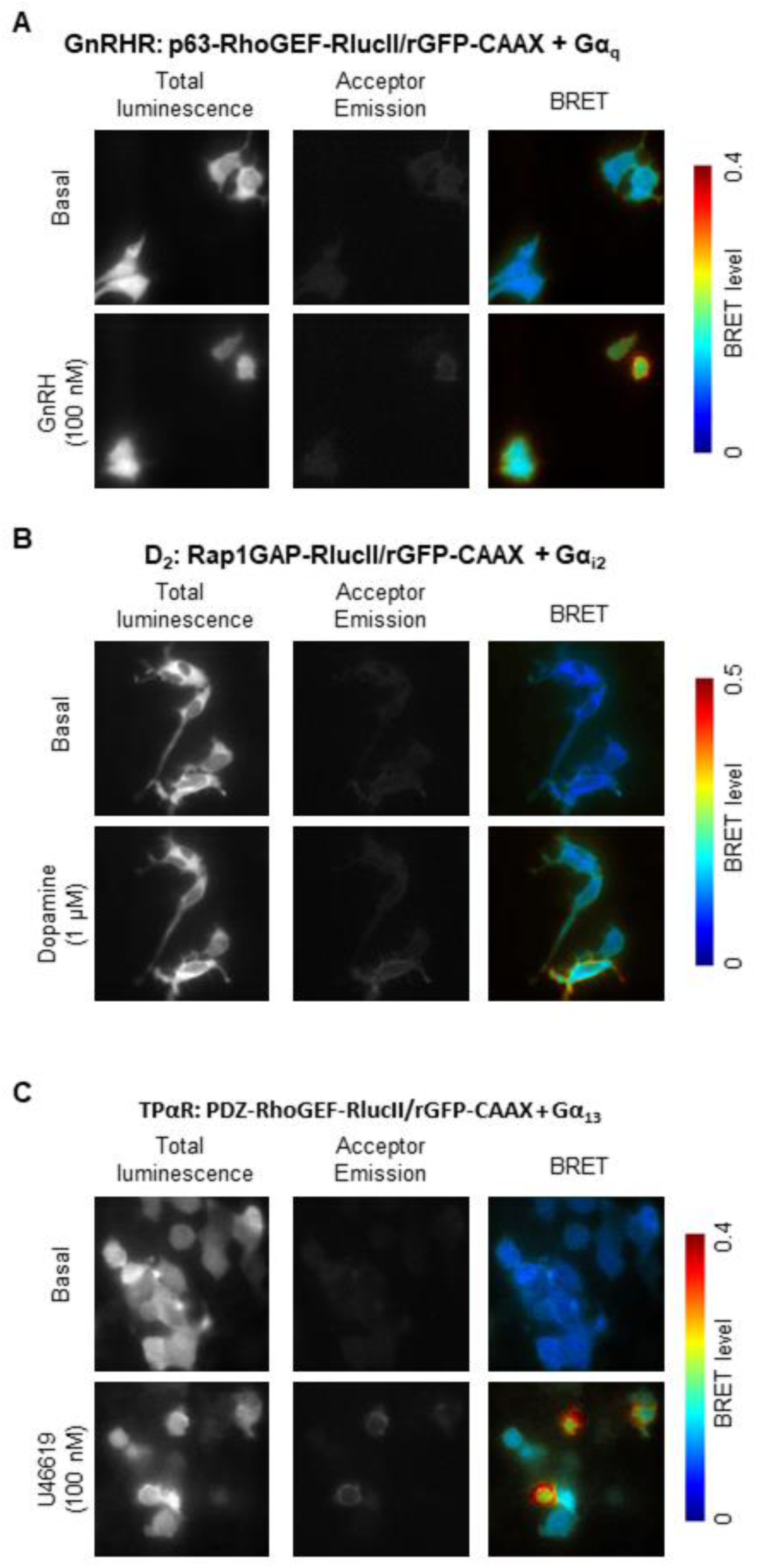
**BRET-based imagery showing sensor recruitment to the plasma membrane upon receptor activation, using the effector membrane translocation assay (EMTA).**

(A) p63-RhoGEF-RlucII recruitment to plasma membrane upon GnRHR activation. HEK293 cells expressing the p63-RhoGEF-RlucII/rGFP-CAAX sensors with Gαq and GnRHR were stimulated with 100 nM GnRH.
(B) Rap1GAP-RlucII recruitment to plasma membrane upon D2 activation. HEK293 cells expressing the Rap1GAP-RlucII/rGFP-CAAX sensors with Gαi2 and D2 were stimulated with 1 µM dopamine.
(C) PDZ-RhoGEF-RlucII recruitment to plasma membrane upon TPαR activation. HEK293 cells expressing the PDZ-RhoGEF-RlucII/rGFP-CAAX + Gα13 and TPαR were stimulated with 100 nM U46619.

Images were obtained in basal condition and 200 sec after stimulation with agonist. In each image, BRET levels (the ratio of the acceptor photon count to the total photon count) are expressed as a heat map color code (lowest being black and purple, and highest being red and white), as shown in the right of the panel.

## References

Aittaleb, M., Boguth, C. A., & Tesmer, J. J. (2010). Structure and function of heterotrimeric G protein-regulated Rho guanine nucleotide exchange factors. Mol Pharmacol, 77(2), 111–125. doi:10.1124/mol.109.061234

Aittaleb, M., Nishimura, A., Linder, M. E., & Tesmer, J. J. (2011). Plasma membrane association of p63 Rho guanine nucleotide exchange factor (p63RhoGEF) is mediated by palmitoylation and is required for basal activity in cells. J Biol Chem, 286(39), 34448–34456. doi:10.1074/jbc.M111.273342

Armando, S., Quoyer, J., Lukashova, V., Maiga, A., Percherancier, Y., Heveker, N., . . . Bouvier, M. (2014). The chemokine CXC4 and CC2 receptors form homo- and heterooligomers that can engage their signaling G-protein effectors and betaarrestin. FASEB J, 28(10), 4509–4523. doi:10.1096/fj.13-242446

Atwood, B. K., Lopez, J., Wager-Miller, J., Mackie, K., & Straiker, A. (2011). Expression of G protein-coupled receptors and related proteins in HEK293, AtT20, BV2, and N18 cell lines as revealed by microarray analysis. BMC Genomics, 12, 14. doi:10.1186/1471-2164-12-14

Avet, C., Sturino, C., Grastilleur, S., Gouill, C. L., Semache, M., Gross, F., . . . Bouvier, M. (2020). The PAR2 inhibitor I-287 selectively targets Galphaq and Galpha12/13 signaling and has anti-inflammatory effects. Commun Biol, 3(1), 719. doi:10.1038/s42003-020-01453-8

Azzi, M., Charest, P. G., Angers, S., Rousseau, G., Kohout, T., Bouvier, M., & Pineyro, G. (2003). Beta-arrestin-mediated activation of MAPK by inverse agonists reveals distinct active conformations for G protein-coupled receptors. Proc Natl Acad Sci U S A, 100(20), 11406–11411. doi:10.1073/pnas.1936664100

Bowes, J., Brown, A. J., Hamon, J., Jarolimek, W., Sridhar, A., Waldron, G., & Whitebread, S. (2012). Reducing safety-related drug attrition: the use of in vitro pharmacological profiling. Nat Rev Drug Discov, 11(12), 909–922. doi:10.1038/nrd3845

Brabet, I., Parmentier, M. L., De Colle, C., Bockaert, J., Acher, F., & Pin, J. P. (1998). Comparative effect of L-CCG-I, DCG-IV and gamma-carboxy-L-glutamate on all cloned metabotropic glutamate receptor subtypes. Neuropharmacology, 37(8), 1043–1051. doi:10.1016/s0028-3908(98)00091-4

Breton, B., Sauvageau, E., Zhou, J., Bonin, H., Le Gouill, C., & Bouvier, M. (2010). Multiplexing of multicolor bioluminescence resonance energy transfer. Biophys J, 99(12), 4037–4046. doi:10.1016/j.bpj.2010.10.025

Bunemann, M., Frank, M., & Lohse, M. J. (2003). Gi protein activation in intact cells involves subunit rearrangement rather than dissociation. Proc Natl Acad Sci U S A, 100(26), 16077–16082. doi:10.1073/pnas.2536719100

Carr, R., 3rd, Du, Y., Quoyer, J., Panettieri, R. A., Jr., Janz, J. M., Bouvier, M., . . . Benovic, J. L. (2014). Development and characterization of pepducins as Gs-biased allosteric agonists. J Biol Chem, 289(52), 35668–35684. doi:10.1074/jbc.M114.618819

Casey, P. J., Fong, H. K., Simon, M. I., & Gilman, A. G. (1990). Gz, a guanine nucleotide- binding protein with unique biochemical properties. J Biol Chem, 265(4), 2383–2390. Retrieved from https://www.ncbi.nlm.nih.gov/pubmed/2105321

Chandan N.R., A. S., SenGupta S., Parent C.A., Smrcka A.V. (2021). Identification of G Protein αi Signaling Partners by Proximity Labeling Reveals a Network of Interactions that Includes PDZ-RhoGEF. bioRxiv preprint. doi:https://doi.org/10.1101/2021.07.15.452545

De Haan, L., & Hirst, T. R. (2004). Cholera toxin: a paradigm for multi-functional engagement of cellular mechanisms (Review). Mol Membr Biol, 21(2), 77–92. doi:10.1080/09687680410001663267

Devost, D., Sleno, R., Petrin, D., Zhang, A., Shinjo, Y., Okde, R., . . . Hebert, T. E. (2017). Conformational Profiling of the AT1 Angiotensin II Receptor Reflects Biased Agonism, G Protein Coupling, and Cellular Context. J Biol Chem, 292(13), 5443–5456. doi:10.1074/jbc.M116.763854

Fukuhara, S., Chikumi, H., & Gutkind, J. S. (2001). RGS-containing RhoGEFs: the missing link between transforming G proteins and Rho? Oncogene, 20(13), 1661–1668. doi:10.1038/sj.onc.1204182

Galandrin, S., Oligny-Longpre, G., & Bouvier, M. (2007). The evasive nature of drug efficacy: implications for drug discovery. Trends Pharmacol Sci, 28(8), 423–430. doi:10.1016/j.tips.2007.06.005

Gales, C., Rebois, R. V., Hogue, M., Trieu, P., Breit, A., Hebert, T. E., & Bouvier, M. (2005). Real-time monitoring of receptor and G-protein interactions in living cells. Nat Methods, 2(3), 177–184. doi:10.1038/nmeth743

Gales, C., Van Durm, J. J., Schaak, S., Pontier, S., Percherancier, Y., Audet, M., . . . Bouvier, M. (2006). Probing the activation-promoted structural rearrangements in preassembled receptor-G protein complexes. Nat Struct Mol Biol, 13(9), 778–786. doi:10.1038/nsmb1134

Goupil, E., Fillion, D., Clement, S., Luo, X., Devost, D., Sleno, R., . . . Hebert, T. E. (2015). Angiotensin II type I and prostaglandin F2alpha receptors cooperatively modulate signaling in vascular smooth muscle cells. J Biol Chem, 290(5), 3137–3148. doi:10.1074/jbc.M114.631119

Hauser, A. S., Attwood, M. M., Rask-Andersen, M., Schioth, H. B., & Gloriam, D. E. (2017). Trends in GPCR drug discovery: new agents, targets and indications. Nat Rev Drug Discov, 16(12), 829–842. doi:10.1038/nrd.2017.178

Hauser, A. S., Avet, C., Normand, C., Mancini, A., Inoue, A., Bouvier, M., & Gloriam, D. E. (2021). GPCR-G protein selectivity - a unified meta-analysis bioRxiv. doi:https://doi.org/10.1101/2021.09.07.459250

Hoffmann, C., Gaietta, G., Bunemann, M., Adams, S. R., Oberdorff-Maass, S., Behr, B., . . . Lohse, M. J. (2005). A FlAsH-based FRET approach to determine G protein-coupled receptor activation in living cells. Nat Methods, 2(3), 171–176. doi:10.1038/nmeth742

Inoue, A., Raimondi, F., Kadji, F. M. N., Singh, G., Kishi, T., Uwamizu, A., . . . Russell, R. B. (2019). Illuminating G-Protein-Coupling Selectivity of GPCRs. Cell, 177(7), 1933–1947 e1925. doi:10.1016/j.cell.2019.04.044

Jordan, J. D., Carey, K. D., Stork, P. J., & Iyengar, R. (1999). Modulation of rap activity by direct interaction of Galpha(o) with Rap1 GTPase-activating protein. J Biol Chem, 274(31), 21507–21510. doi:10.1074/jbc.274.31.21507

Kawamata, Y., Fujii, R., Hosoya, M., Harada, M., Yoshida, H., Miwa, M., . . . Fujino, M. (2003). A G protein-coupled receptor responsive to bile acids. J Biol Chem, 278(11), 9435–9440. doi:10.1074/jbc.M209706200

Kenakin, T. (2019). Biased Receptor Signaling in Drug Discovery. Pharmacol Rev, 71(2), 267–315. doi:10.1124/pr.118.016790

Kim, J., Isokawa, M., Ledent, C., & Alger, B. E. (2002). Activation of muscarinic acetylcholine receptors enhances the release of endogenous cannabinoids in the hippocampus. J Neurosci, 22(23), 10182–10191. Retrieved from https://www.ncbi.nlm.nih.gov/pubmed/12451119

Kobayashi, H., Picard, L. P., Schonegge, A. M., & Bouvier, M. (2019). Bioluminescence resonance energy transfer-based imaging of protein-protein interactions in living cells. Nat Protoc, 14(4), 1084–1107. doi:10.1038/s41596-019-0129-7

Laschet, C., Dupuis, N., & Hanson, J. (2019). A dynamic and screening-compatible nanoluciferase-based complementation assay enables profiling of individual GPCR-G protein interactions. J Biol Chem, 294(11), 4079–4090. doi:10.1074/jbc.RA118.006231

Lavoie, C., Mercier, J. F., Salahpour, A., Umapathy, D., Breit, A., Villeneuve, L. R., . . . Hebert, T. E. (2002). Beta 1/beta 2-adrenergic receptor heterodimerization regulates beta 2-adrenergic receptor internalization and ERK signaling efficacy. J Biol Chem, 277(38), 35402–35410. doi:10.1074/jbc.M204163200

Leduc, M., Breton, B., Gales, C., Le Gouill, C., Bouvier, M., Chemtob, S., & Heveker, N. (2009). Functional selectivity of natural and synthetic prostaglandin EP4 receptor ligands. J Pharmacol Exp Ther, 331(1), 297–307. doi:10.1124/jpet.109.156398

Leguay, K., Decelle, B., He, Y. Y., Pagniez, A., Hogue, M., Kobayashi, H., . . . Carreno, S. (2021). Development of conformational BRET biosensors that monitor ezrin, radixin and moesin activation in real time. J Cell Sci, 134(7). doi:10.1242/jcs.255307

Lu, M., Wang, B., Zhang, C., Zhuang, X., Yuan, M., Wang, H., . . . Li, J. (2014). PQ-69, a novel and selective adenosine A1 receptor antagonist with inverse agonist activity. Purinergic Signal, 10(4), 619–629. doi:10.1007/s11302-014-9424-5

Lutz, S., Shankaranarayanan, A., Coco, C., Ridilla, M., Nance, M. R., Vettel, C., . . . Tesmer, J. J. (2007). Structure of Galphaq-p63RhoGEF-RhoA complex reveals a pathway for the activation of RhoA by GPCRs. Science, 318(5858), 1923–1927. doi:10.1126/science.1147554

Mancini, A., Frauli, M., & Breton, B. (2015). Exploring the Technology Landscape of 7TMR Drug Signaling Profiling. Curr Top Med Chem, 15(24), 2528–2542. doi:10.2174/1568026615666150701113344

Martin, B. R., & Lambert, N. A. (2016). Activated G Protein Galphas Samples Multiple Endomembrane Compartments. J Biol Chem, 291(39), 20295–20302. doi:10.1074/jbc.M116.729731

Masuho, I., Ostrovskaya, O., Kramer, G. M., Jones, C. D., Xie, K., & Martemyanov, K. A. (2015). Distinct profiles of functional discrimination among G proteins determine the actions of G protein-coupled receptors. Sci Signal, 8(405), ra123. doi:10.1126/scisignal.aab4068

Maziarz, M., Park, J. C., Leyme, A., Marivin, A., Garcia-Lopez, A., Patel, P. P., & Garcia- Marcos, M. (2020). Revealing the Activity of Trimeric G-proteins in Live Cells with a Versatile Biosensor Design. Cell, 182(3), 770–785 e716. doi:10.1016/j.cell.2020.06.020

McAvoy, T., Zhou, M. M., Greengard, P., & Nairn, A. C. (2009). Phosphorylation of Rap1GAP, a striatally enriched protein, by protein kinase A controls Rap1 activity and dendritic spine morphology. Proc Natl Acad Sci U S A, 106(9), 3531–3536. doi:10.1073/pnas.0813263106

Mende, F., Hundahl, C., Plouffe, B., Skov, L. J., Sivertsen, B., Madsen, A. N., . . . Holst, B. (2018). Translating biased signaling in the ghrelin receptor system into differential in vivo functions. Proc Natl Acad Sci U S A, 115(43), E10255–E10264. doi:10.1073/pnas.1804003115

Meng, J., Glick, J. L., Polakis, P., & Casey, P. J. (1999). Functional interaction between Galpha(z) and Rap1GAP suggests a novel form of cellular cross-talk. J Biol Chem, 274(51), 36663–36669. doi:10.1074/jbc.274.51.36663

Namkung, Y., Le Gouill, C., Lukashova, V., Kobayashi, H., Hogue, M., Khoury, E., . . . Laporte, S. A. (2016). Monitoring G protein-coupled receptor and beta-arrestin trafficking in live cells using enhanced bystander BRET. Nat Commun, 7, 12178. doi:10.1038/ncomms12178

Namkung, Y., LeGouill, C., Kumar, S., Cao, Y., Teixeira, L. B., Lukasheva, V., . . . Laporte, S. A. (2018). Functional selectivity profiling of the angiotensin II type 1 receptor using pathway-wide BRET signaling sensors. Sci Signal, 11(559). doi:10.1126/scisignal.aat1631

Okashah, N., Wright, S. C., Kawakami, K., Mathiasen, S., Zhou, J., Lu, S., . . . Lambert, N. A. (2020). Agonist-induced formation of unproductive receptor-G12 complexes. Proc Natl Acad Sci U S A, 117(35), 21723–21730. doi:10.1073/pnas.2003787117

Oldham, W. M., & Hamm, H. E. (2008). Heterotrimeric G protein activation by G-protein- coupled receptors. Nat Rev Mol Cell Biol, 9(1), 60–71. doi:10.1038/nrm2299

Olsen, R. H. J., DiBerto, J. F., English, J. G., Glaudin, A. M., Krumm, B. E., Slocum, S. T., . . . Strachan, R. T. (2020). TRUPATH, an open-source biosensor platform for interrogating the GPCR transducerome. Nat Chem Biol, 16(8), 841–849. doi:10.1038/s41589-020-0535-8

Quoyer, J., Janz, J. M., Luo, J., Ren, Y., Armando, S., Lukashova, V., . . . Bouvier, M. (2013). Pepducin targeting the C-X-C chemokine receptor type 4 acts as a biased agonist favoring activation of the inhibitory G protein. Proc Natl Acad Sci U S A, 110(52), E5088–5097. doi:10.1073/pnas.1312515110

Rojas, R. J., Yohe, M. E., Gershburg, S., Kawano, T., Kozasa, T., & Sondek, J. (2007). Galphaq directly activates p63RhoGEF and Trio via a conserved extension of the Dbl homology- associated pleckstrin homology domain. J Biol Chem, 282(40), 29201–29210. doi:10.1074/jbc.M703458200

Roth, B. L., Sheffler, D. J., & Kroeze, W. K. (2004). Magic shotguns versus magic bullets: selectively non-selective drugs for mood disorders and schizophrenia. Nat Rev Drug Discov, 3(4), 353–359. doi:10.1038/nrd1346

Sanchez-Soto, M., Bonifazi, A., Cai, N. S., Ellenberger, M. P., Newman, A. H., Ferre, S., & Yano, H. (2016). Evidence for Noncanonical Neurotransmitter Activation: Norepinephrine as a Dopamine D2-Like Receptor Agonist. Mol Pharmacol, 89(4), 457–466. doi:10.1124/mol.115.101808

Schrage, R., Schmitz, A. L., Gaffal, E., Annala, S., Kehraus, S., Wenzel, D., . . . Kostenis, E. (2015). The experimental power of FR900359 to study Gq-regulated biological processes. Nat Commun, 6, 10156. doi:10.1038/ncomms10156

Stallaert, W., van der Westhuizen, E. T., Schonegge, A. M., Plouffe, B., Hogue, M., Lukashova, V., . . . Bouvier, M. (2017). Purinergic Receptor Transactivation by the beta2-Adrenergic Receptor Increases Intracellular Ca(2+) in Nonexcitable Cells. Mol Pharmacol, 91(5), 533–544. doi:10.1124/mol.116.106419

Sunahara, R. K., Guan, H. C., O’Dowd, B. F., Seeman, P., Laurier, L. G., Ng, G., . . . Niznik, H. A. (1991). Cloning of the gene for a human dopamine D5 receptor with higher affinity for dopamine than D1. Nature, 350(6319), 614–619. doi:10.1038/350614a0

Takasaki, J., Saito, T., Taniguchi, M., Kawasaki, T., Moritani, Y., Hayashi, K., & Kobori, M. (2004). A novel Galphaq/11-selective inhibitor. J Biol Chem, 279(46), 47438–47445. doi:10.1074/jbc.M408846200

Wedegaertner, P. B., Bourne, H. R., & von Zastrow, M. (1996). Activation-induced subcellular redistribution of Gs alpha. Mol Biol Cell, 7(8), 1225–1233. doi:10.1091/mbc.7.8.1225

Wei, H., Ahn, S., Shenoy, S. K., Karnik, S. S., Hunyady, L., Luttrell, L. M., & Lefkowitz, R. J. (2003). Independent beta-arrestin 2 and G protein-mediated pathways for angiotensin II activation of extracellular signal-regulated kinases 1 and 2. Proc Natl Acad Sci U S A, 100(19), 10782–10787. doi:10.1073/pnas.1834556100

Zimmerman, B., Beautrait, A., Aguila, B., Charles, R., Escher, E., Claing, A., . . . Laporte, S. A. (2012). Differential beta-arrestin-dependent conformational signaling and cellular responses revealed by angiotensin analogs. Sci Signal, 5(221), ra33. doi:10.1126/scisignal.2002522

