## Supplementary File 1 for "Effector membrane translocation biosensors reveal G protein and βarrestin coupling profiles of 100 therapeutically relevant GPCRs"

**GPCR: 5-HT$_{1A}$**

**Ligand: Serotonin**

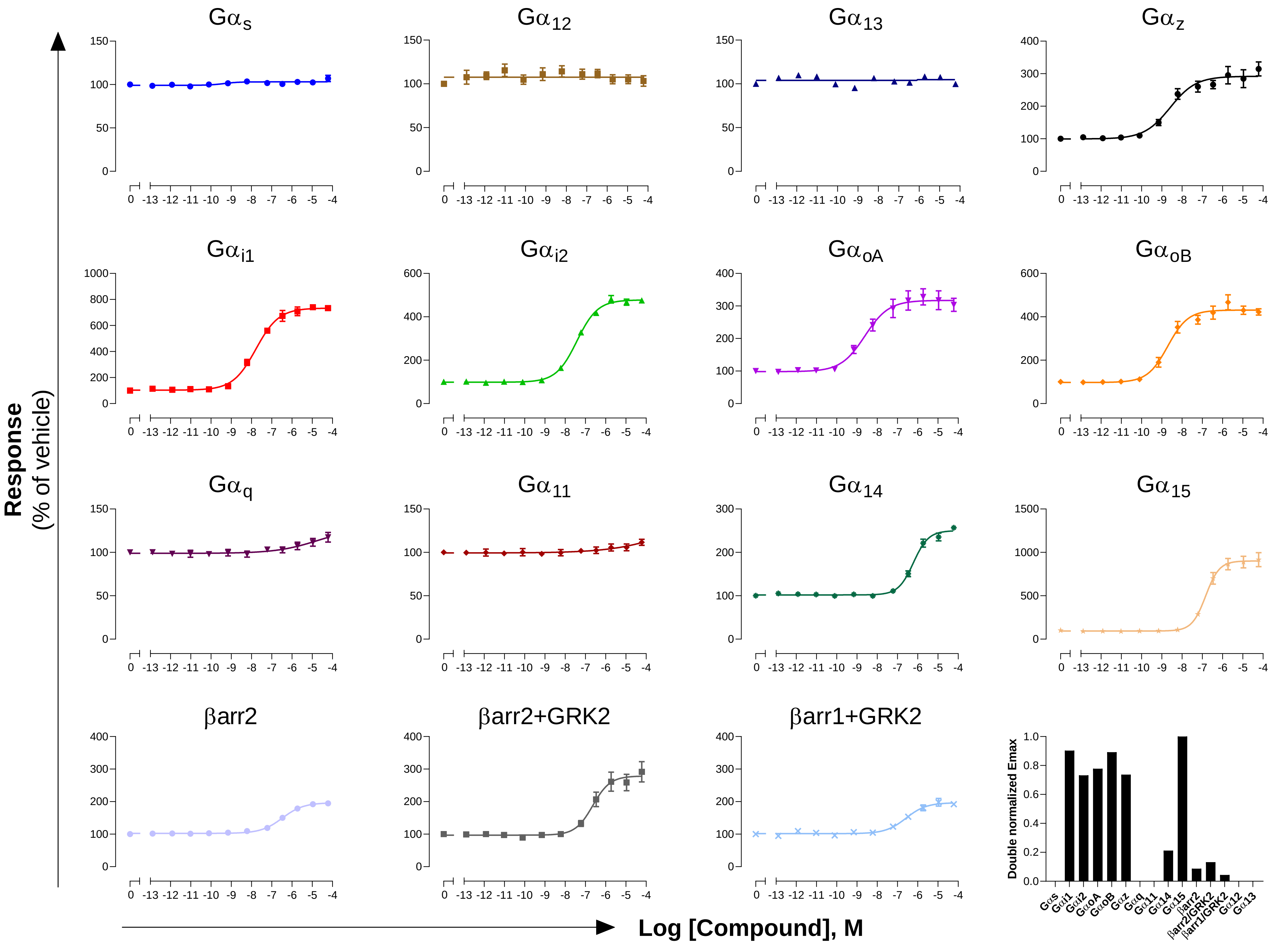

**GPCR: 5-HT$_{1B}$**

**Ligand: Serotonin**

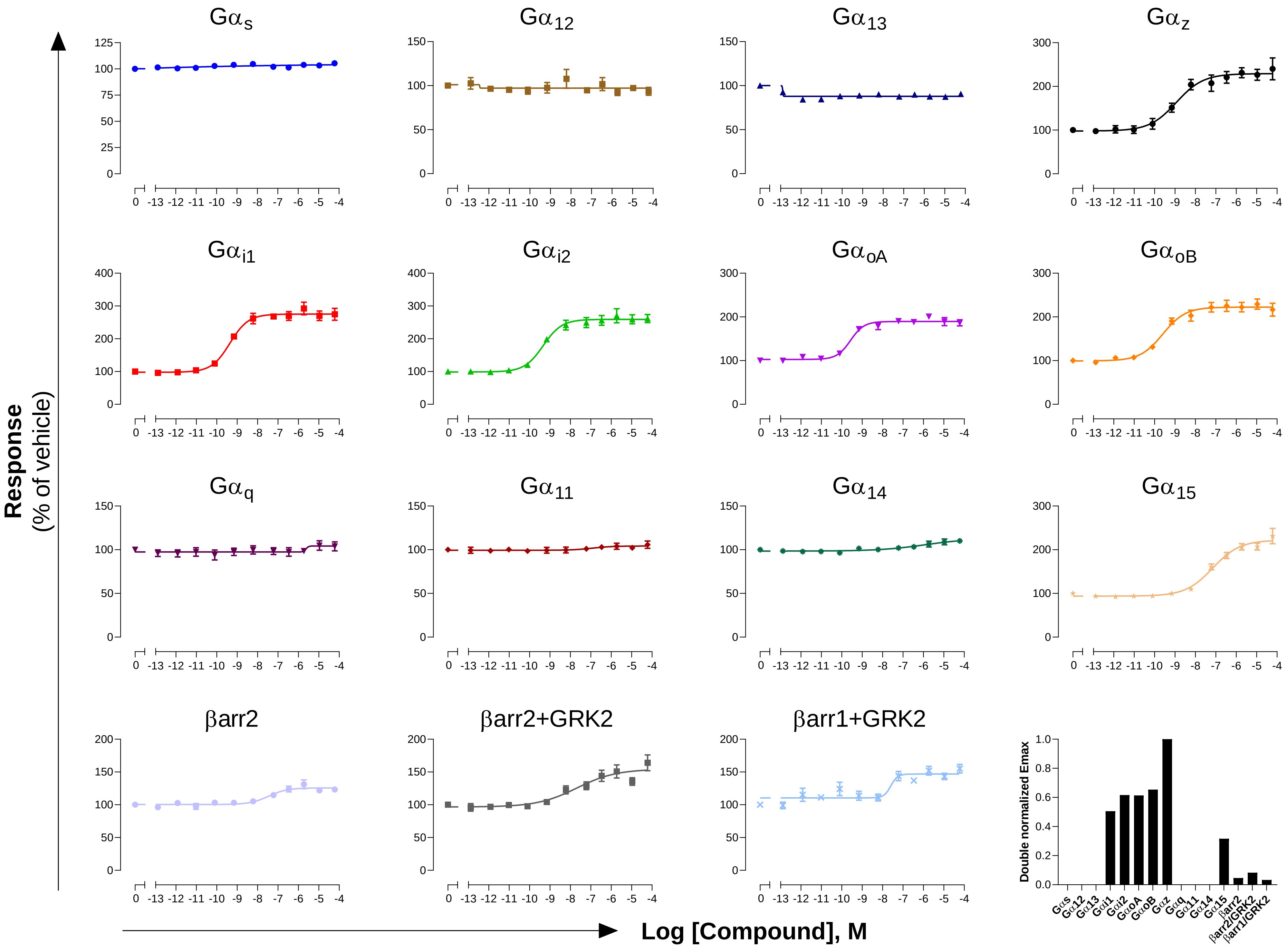

**GPCR: 5-HT$_{1D}$**

**Ligand: Serotonin**

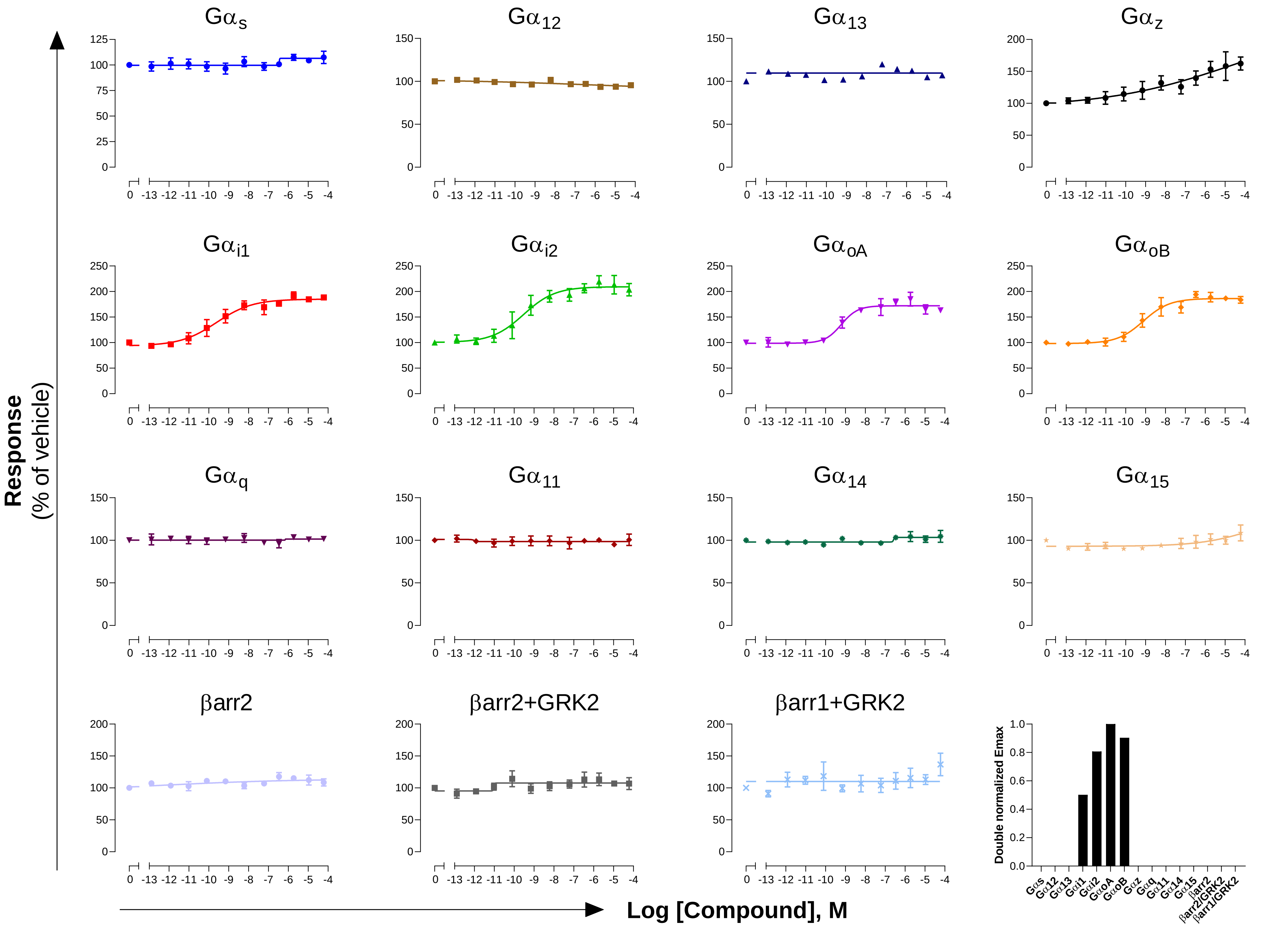

**GPCR: 5-$HT_{2A}$**
**Ligand: Serotonin**

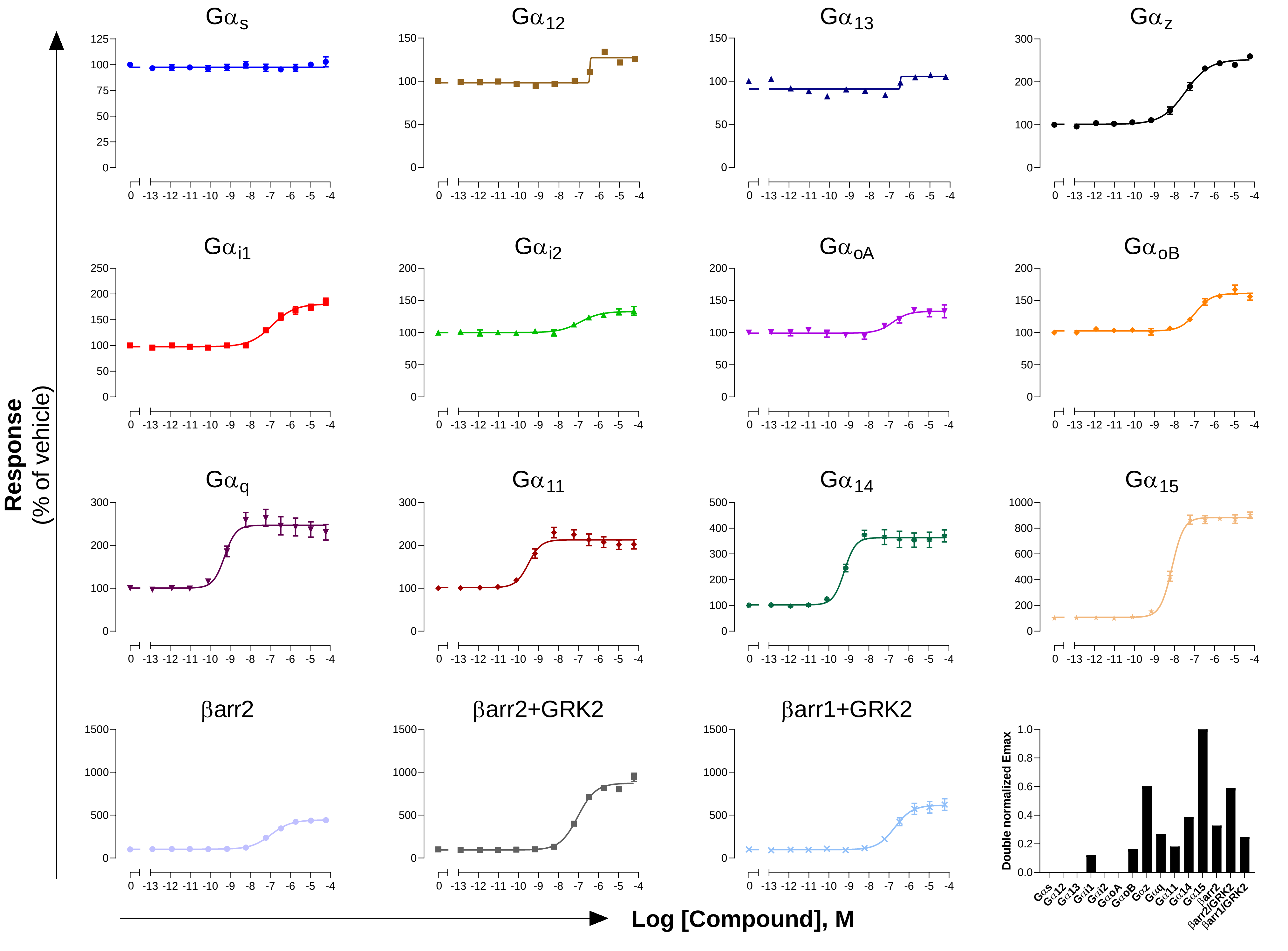

GPCR: 5-$HT_{2B}$

Ligand: Serotonin

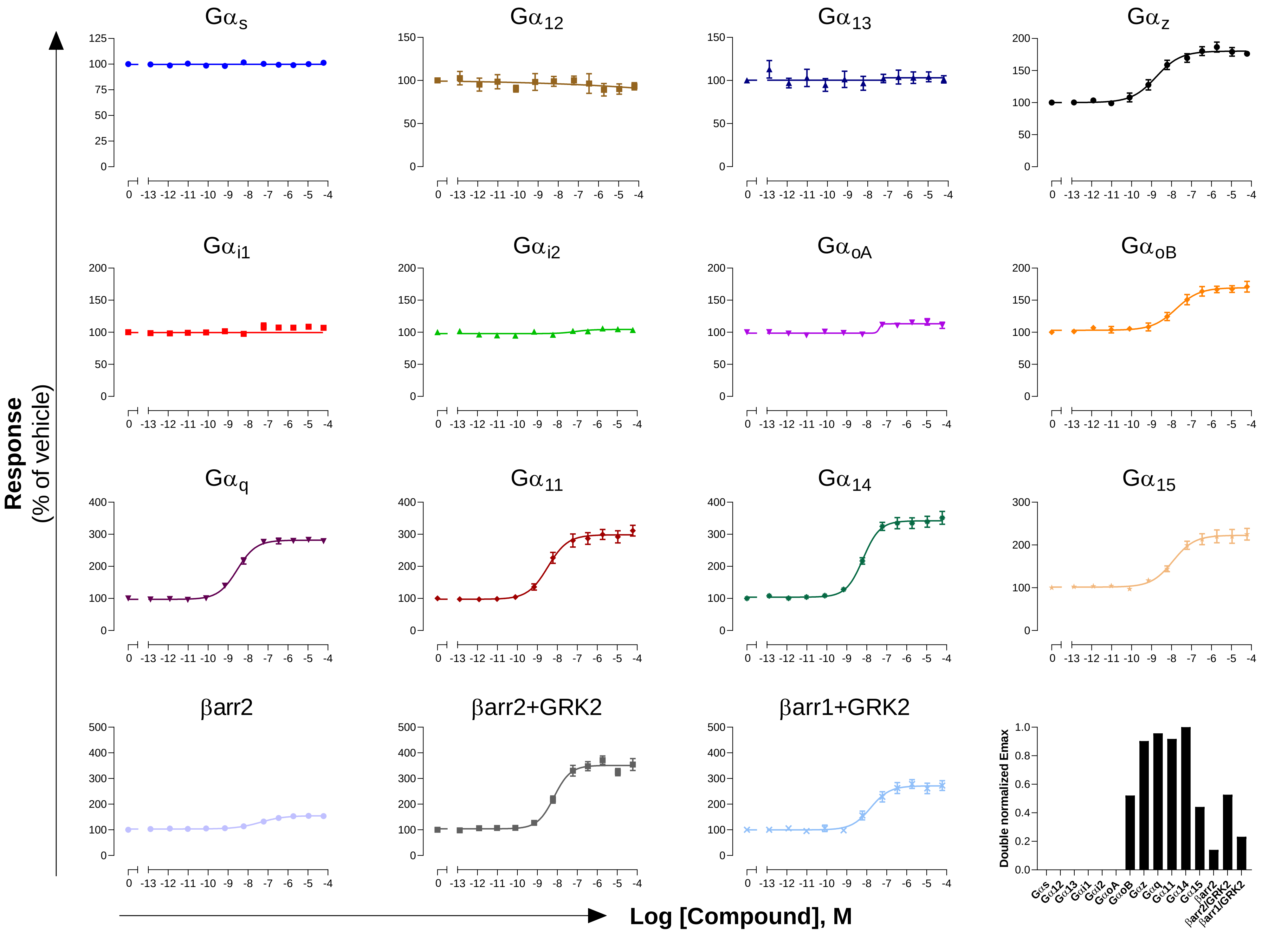

GPCR: 5-HT$_{2C}$

Ligand: Serotonin

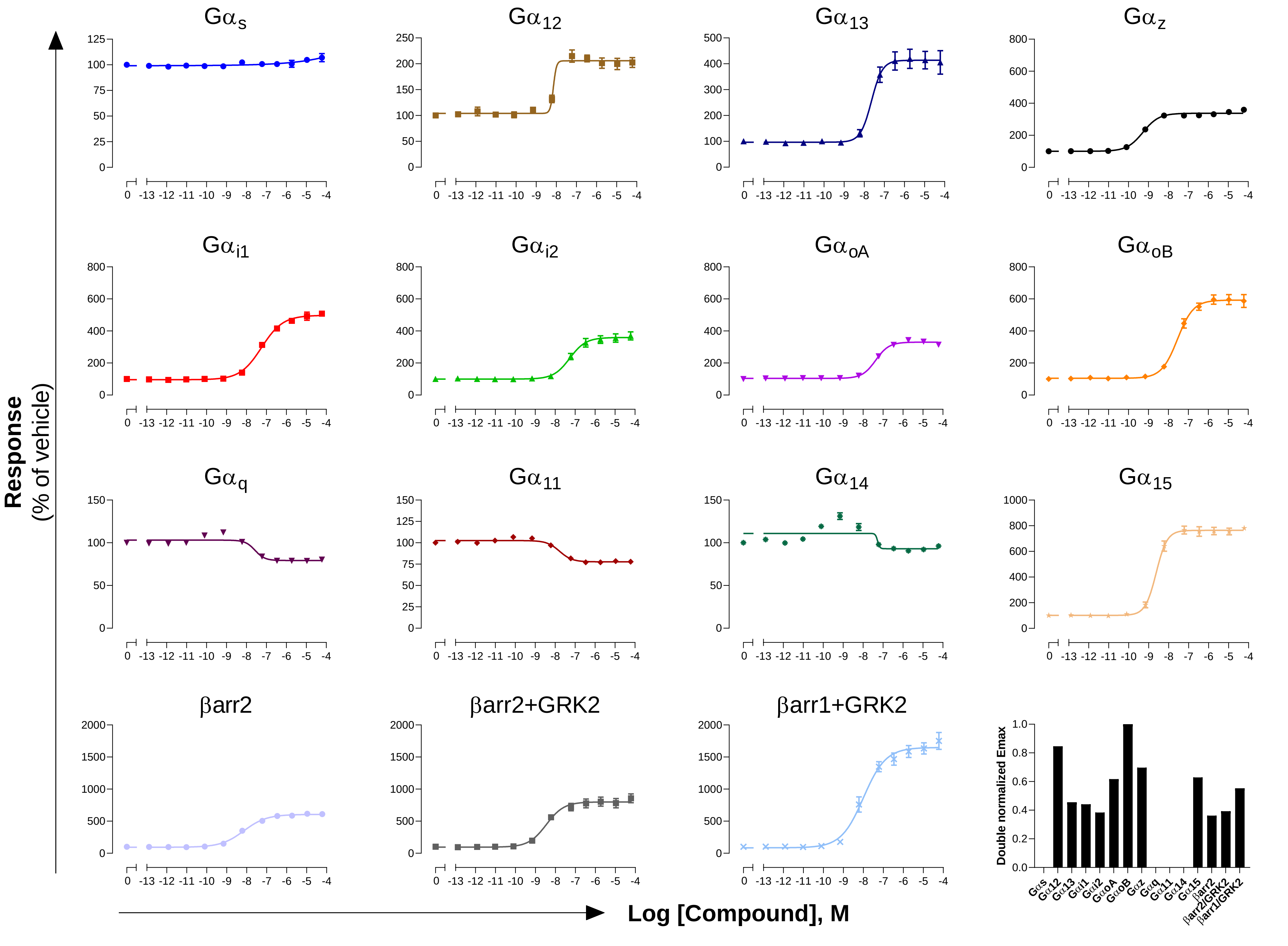

**GPCR: $\alpha_{1A}$AR**

**Ligand: Noradrenaline**

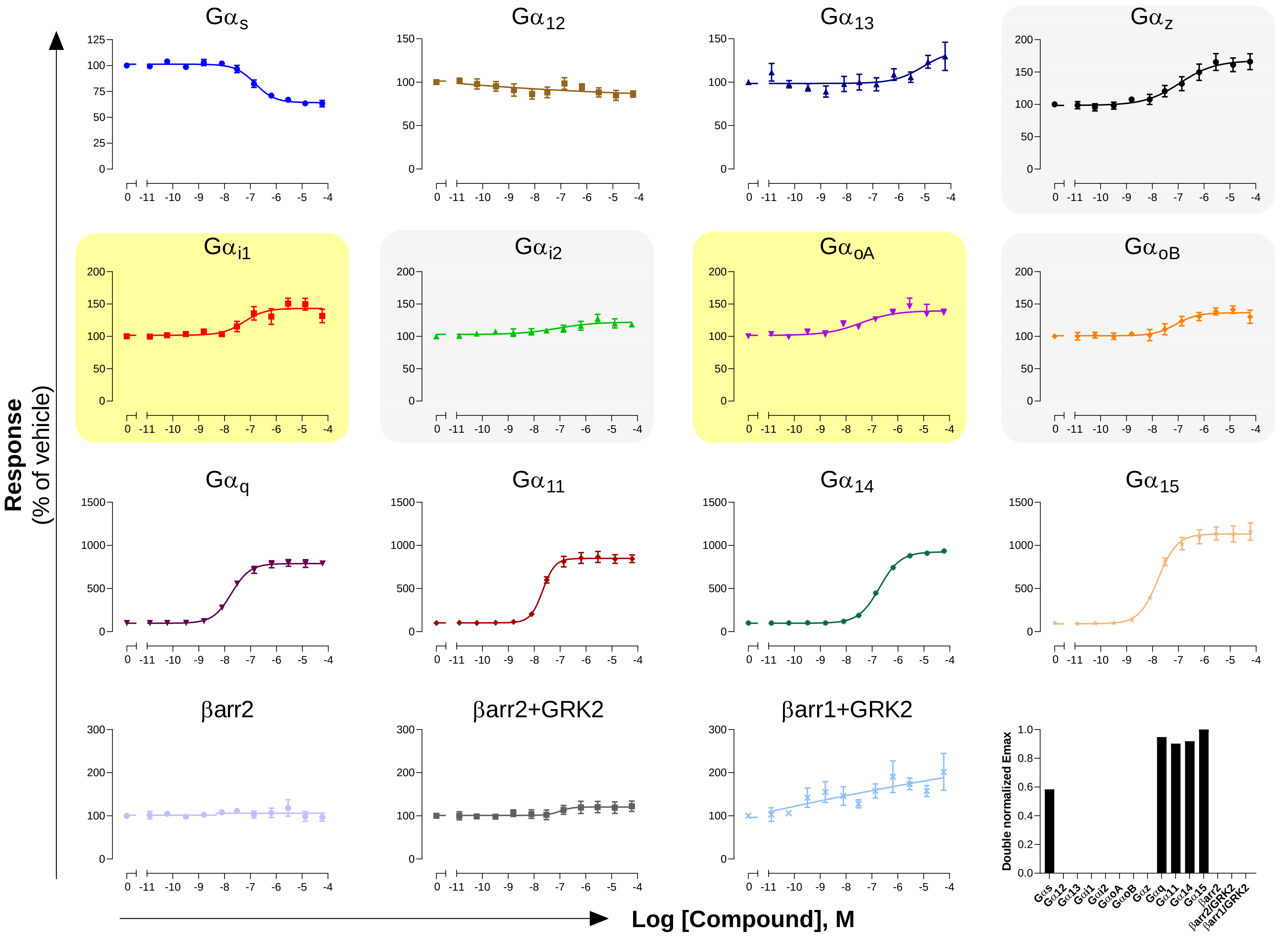

**GPCR: $\alpha_{2A}$AR**

**Ligand: Noradrenaline**

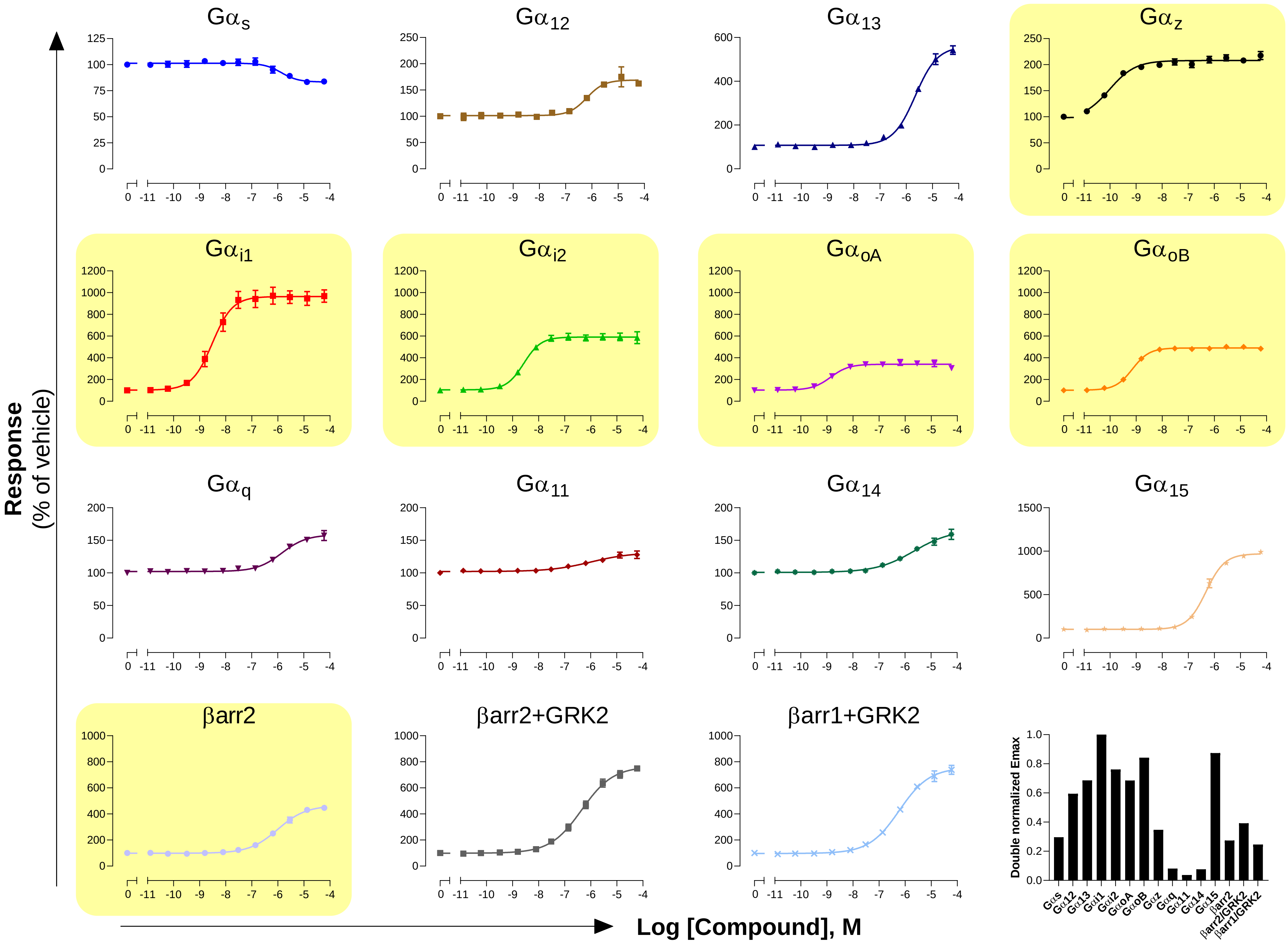

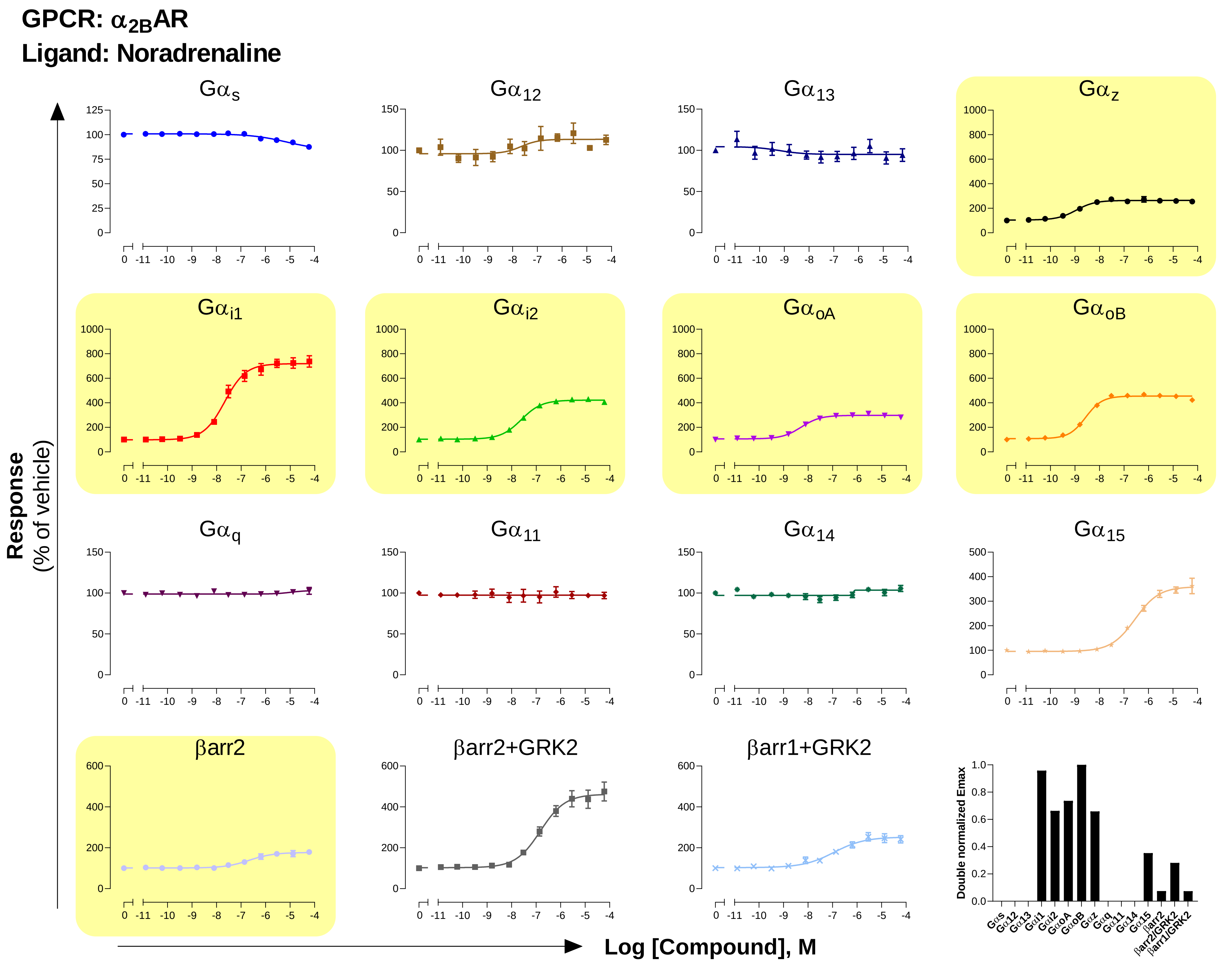

GPCR: α2BAR
Ligand: Noradrenaline
Gαs
Gα12
Gα13
Gαz
Gαi1
Gαi2
GαoA
GαoB
Gαq
Gα11
Gα14
Gα15
βarr2
βarr2+GRK2
βarr1+GRK2
Response
(% of vehicle)
Log [Compound], M
Double normalized Emax
Gαs Gα12 Gα13 Gαi1 Gαi2 GαoA GαoB Gαz Gαq Gα11 Gα14 Gα15 βarr2 βarr2/GRK2 βarr1/GRK2

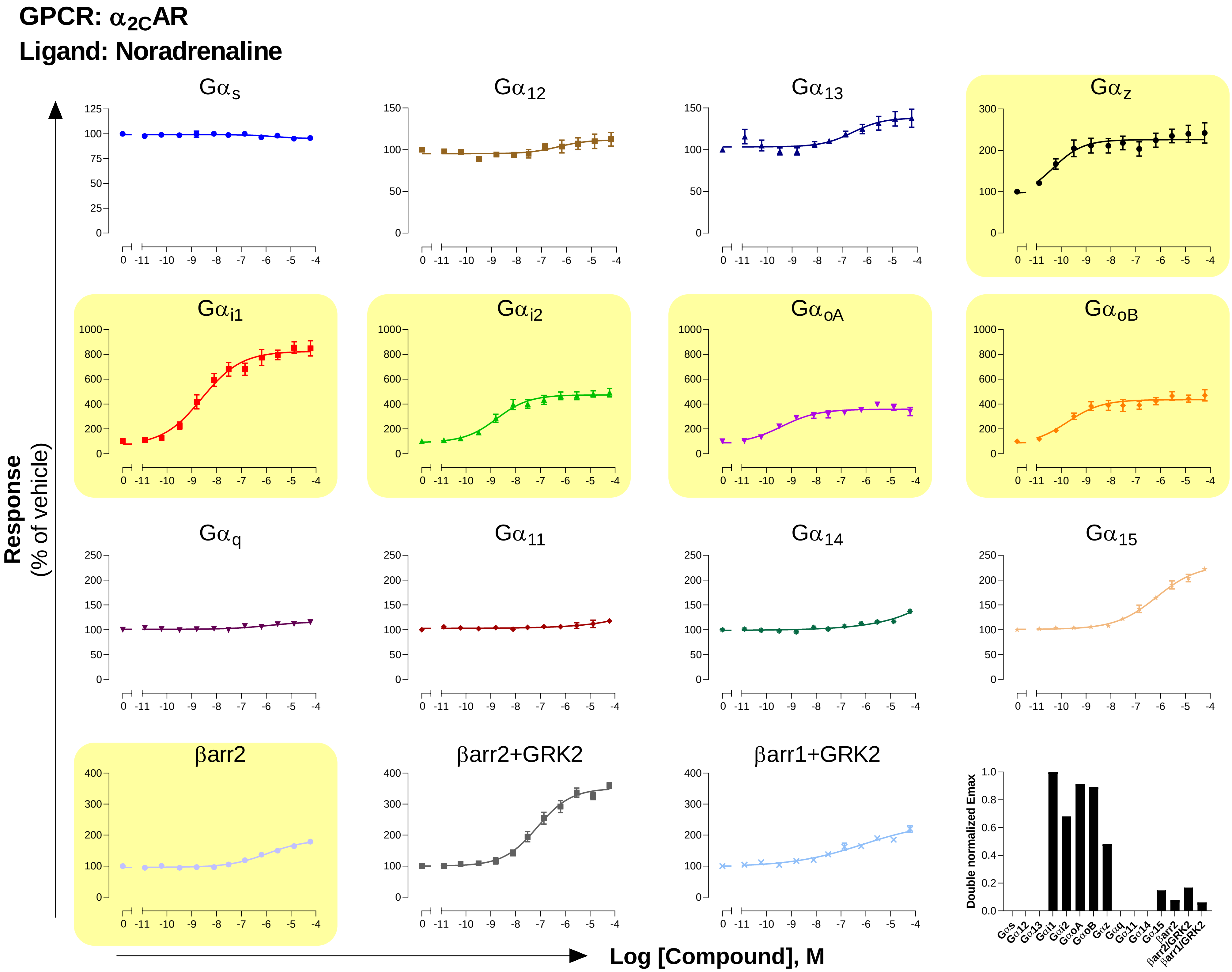

GPCR: α2CAR
Ligand: Noradrenaline
Gαs
Gα12
Gα13
Gαz
Gαi1
Gαi2
GαoA
GαoB
Gαq
Gα11
Gα14
Gα15
βarr2
βarr2+GRK2
βarr1+GRK2
Response
(% of vehicle)
Log [Compound], M
Double normalized Emax

#### GPCR: $\beta_1$AR
#### Ligand: Noradrenaline

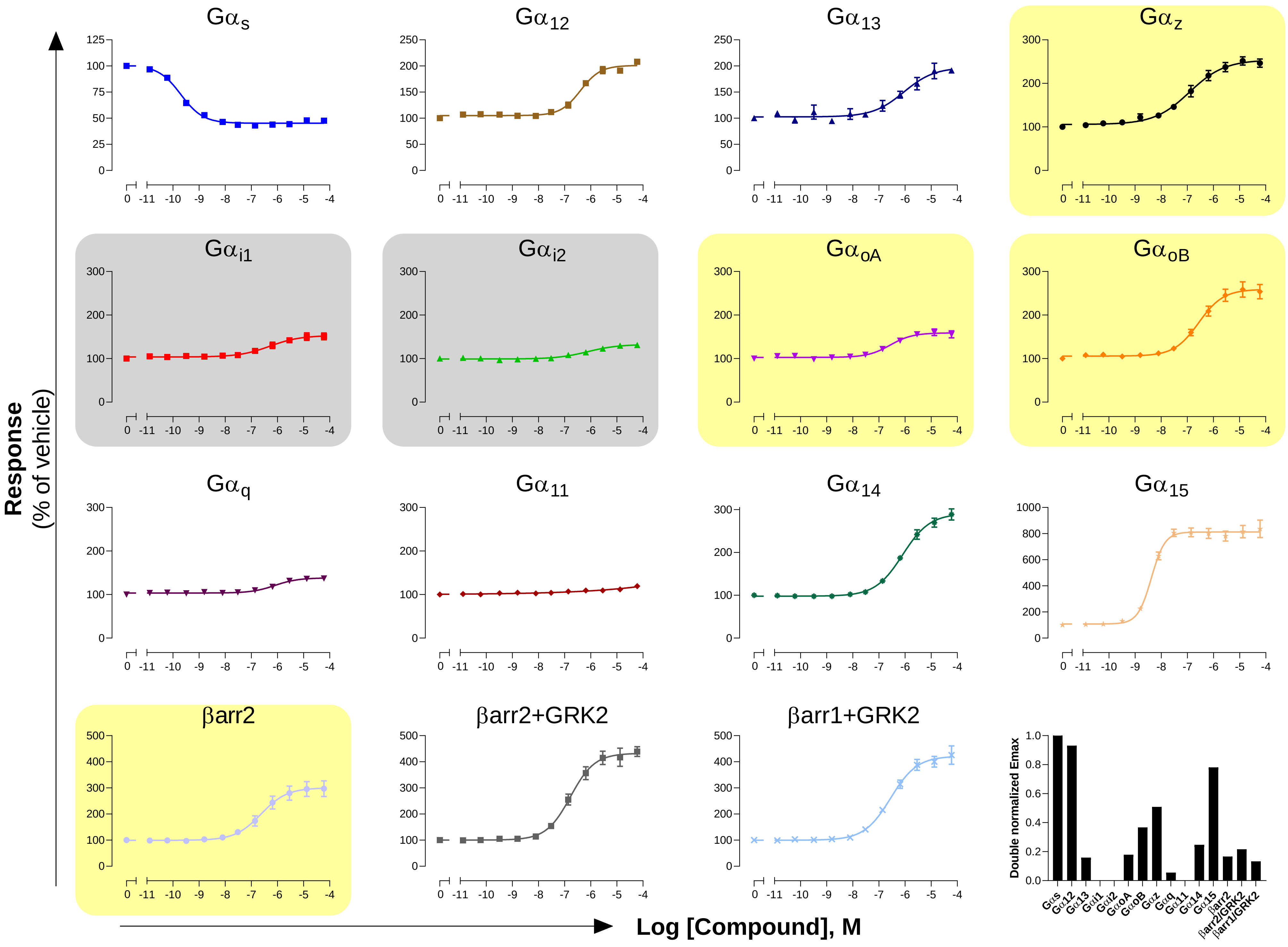

**GPCR: $\beta_2$AR**

**Ligand: Noradrenaline**

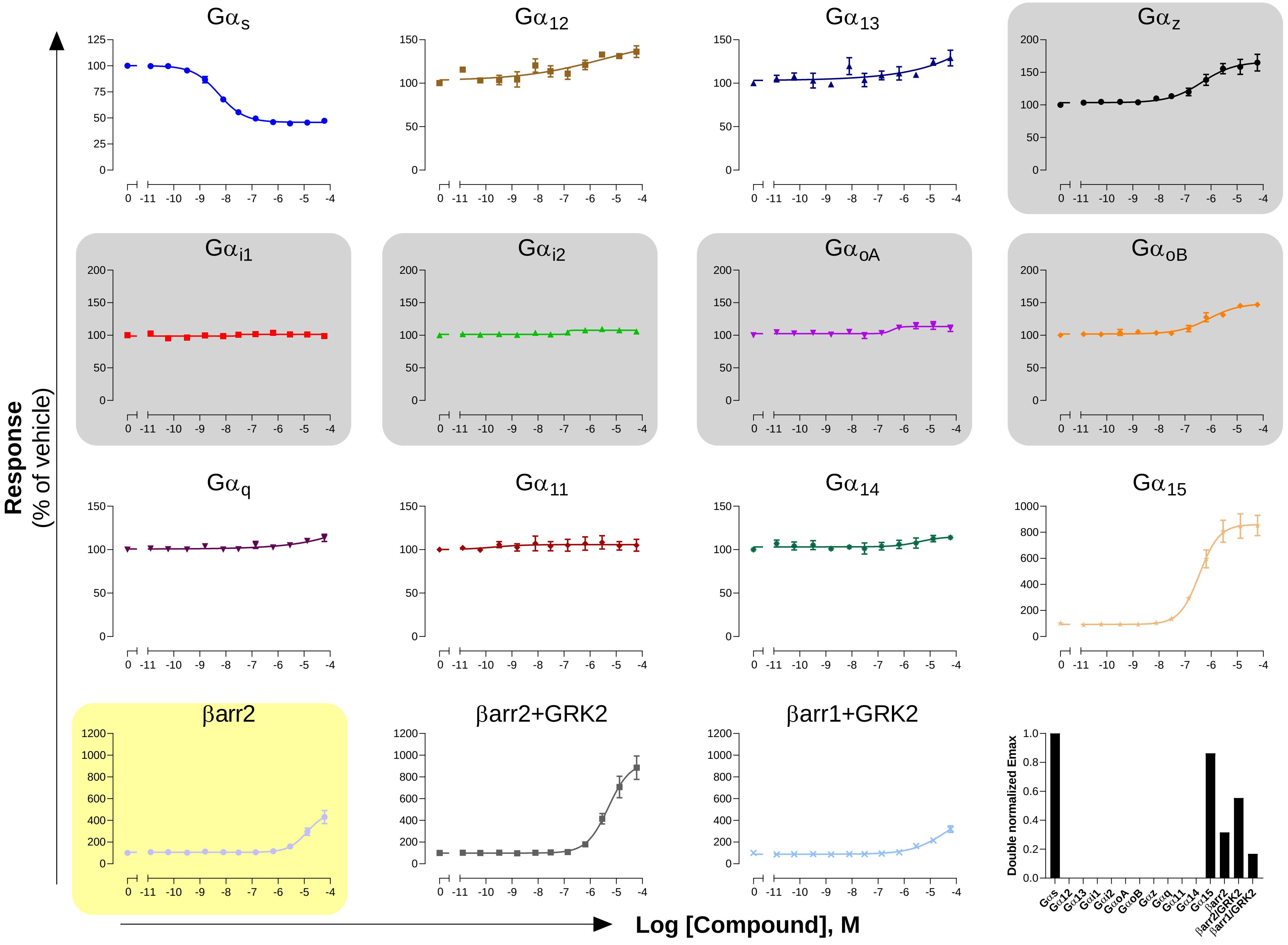

#### GPCR: $A_1$
#### Ligand: Adenosine

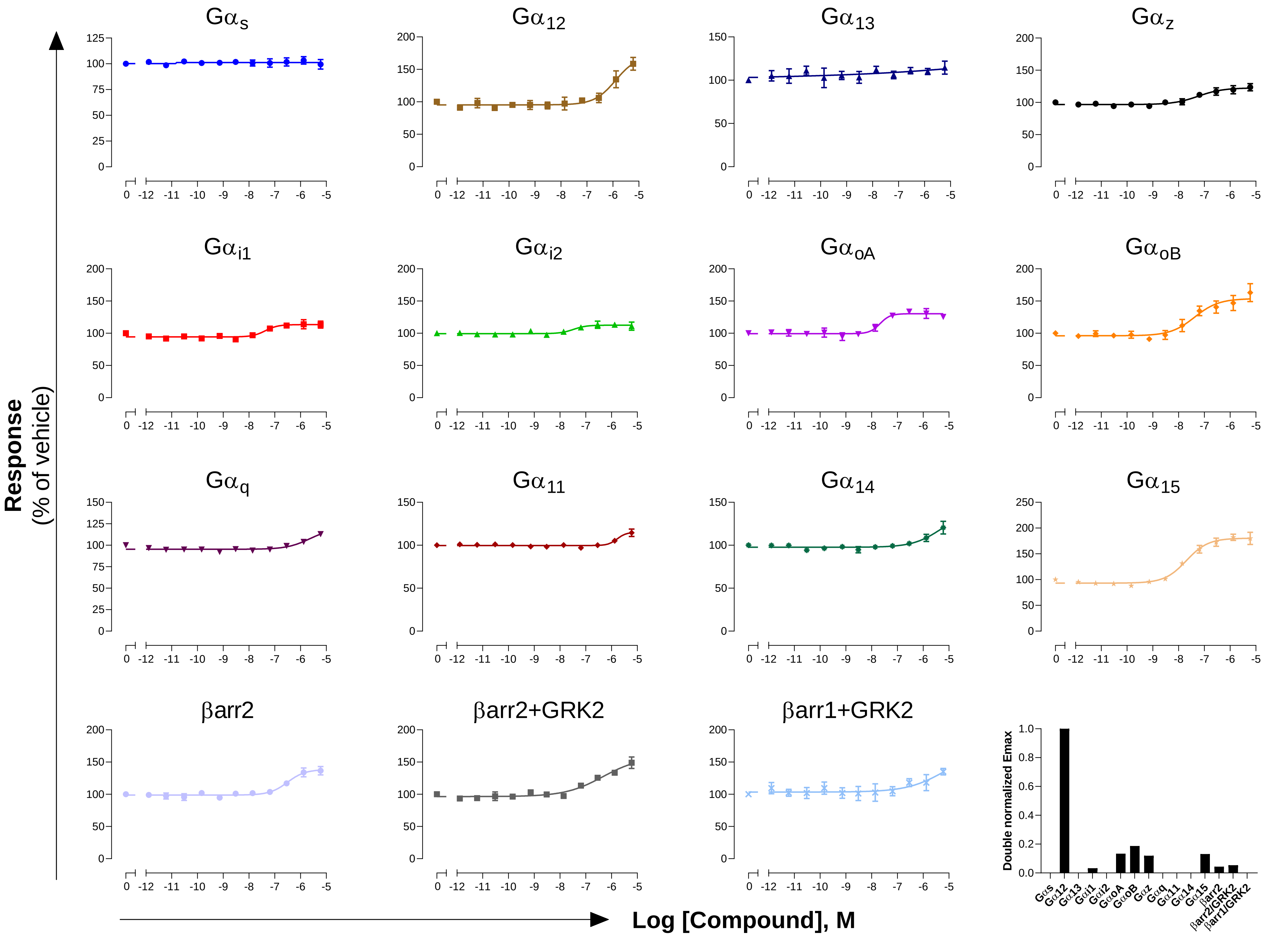

#### GPCR: $A_{2A}$
#### Ligand: Adenosine

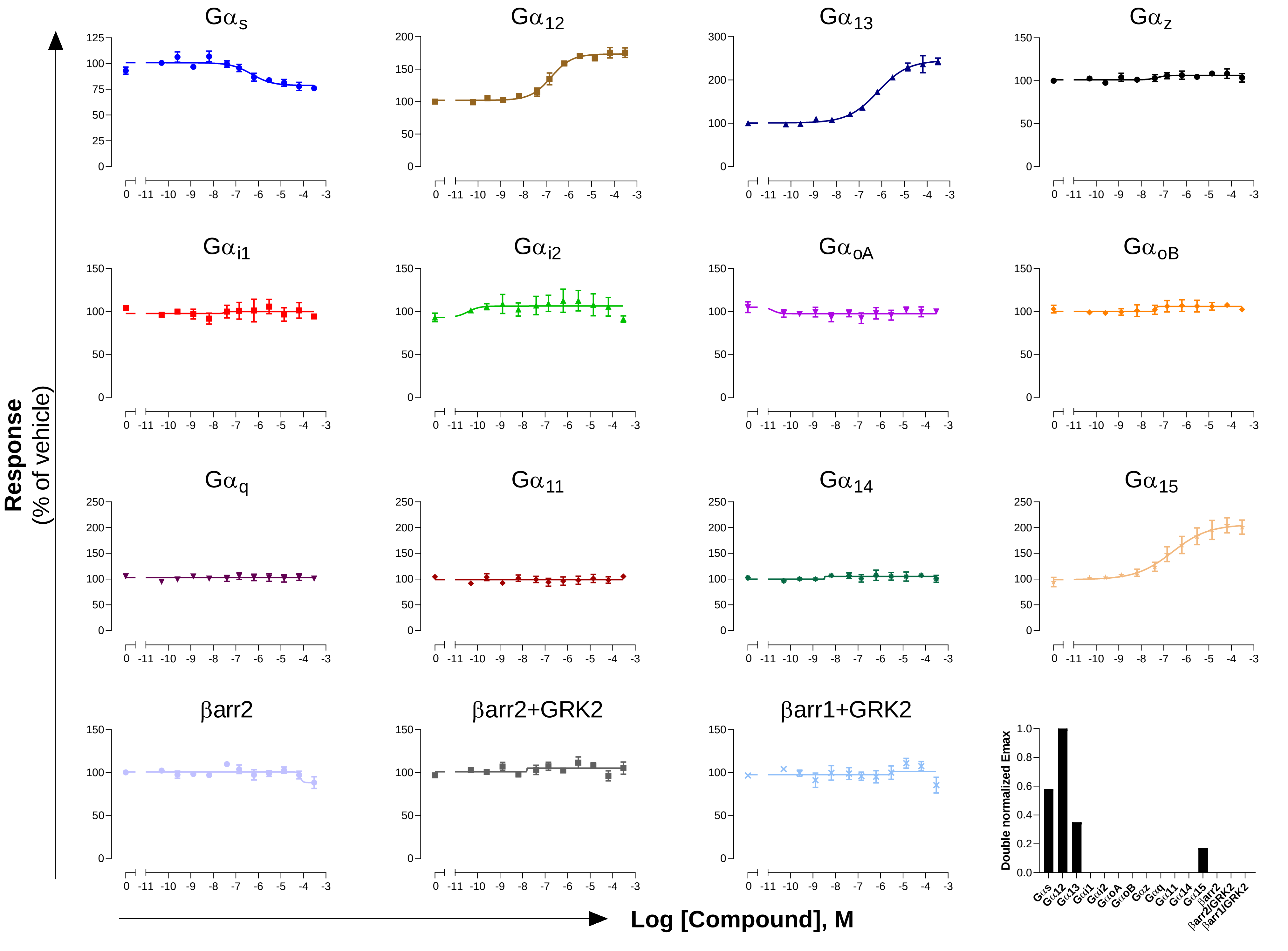

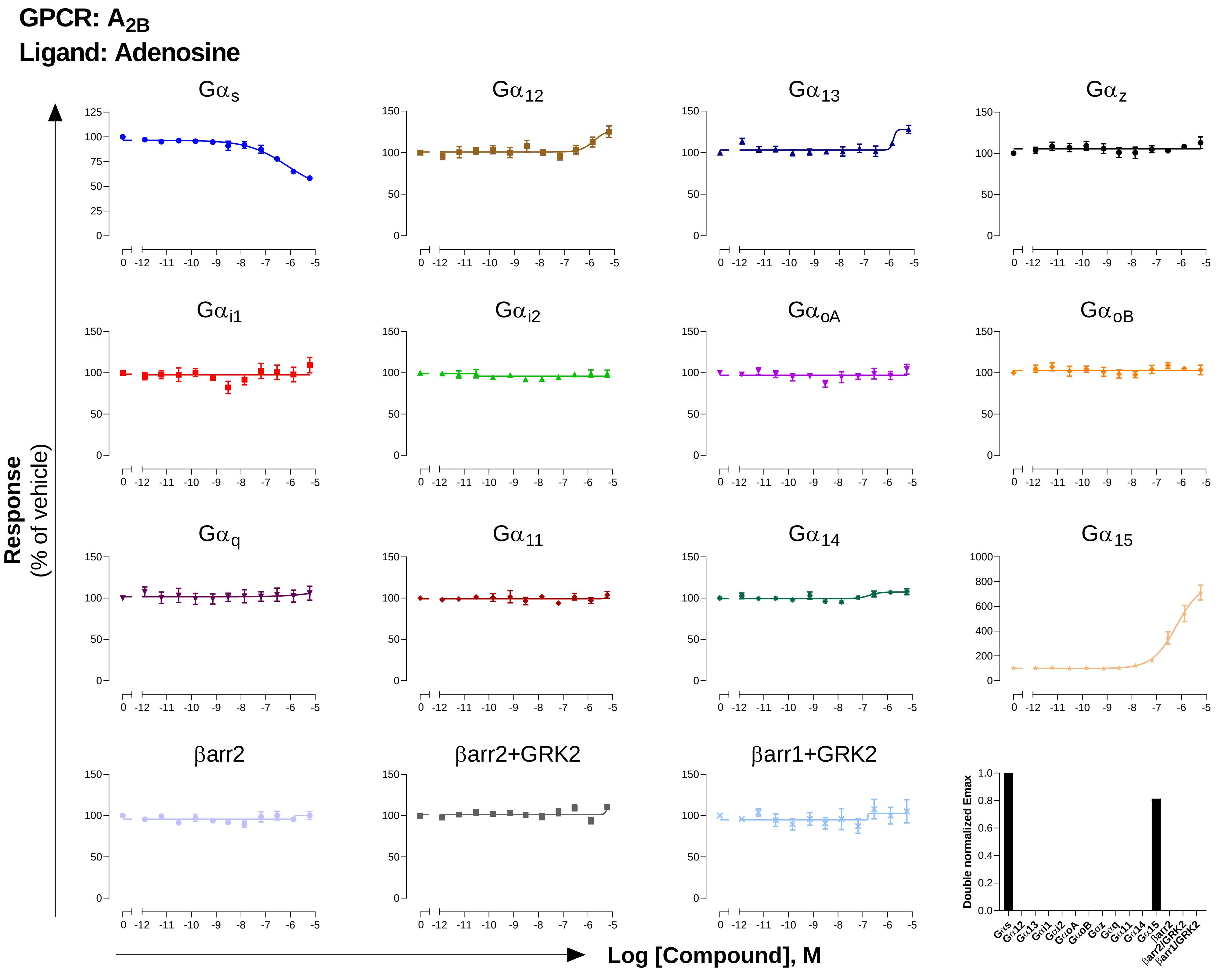
GPCR: $A_{2B}$
Ligand: Adenosine
Gαs
Gα12
Gα13
Gαz
Gαi1
Gαi2
GαoA
GαoB
Gαq
Gα11
Gα14
Gα15
βarr2
βarr2+GRK2
βarr1+GRK2
Response
(% of vehicle)
Log [Compound], M
Double normalized Emax
Gαs Gα12 Gα13 Gαi1 Gαi2 GαoA GαoB Gαz Gαq Gα11 Gα14 Gα15 βarr2 βarr2/GRK2 βarr1/GRK2

**GPCR: $A_3$**

**Ligand: Adenosine**

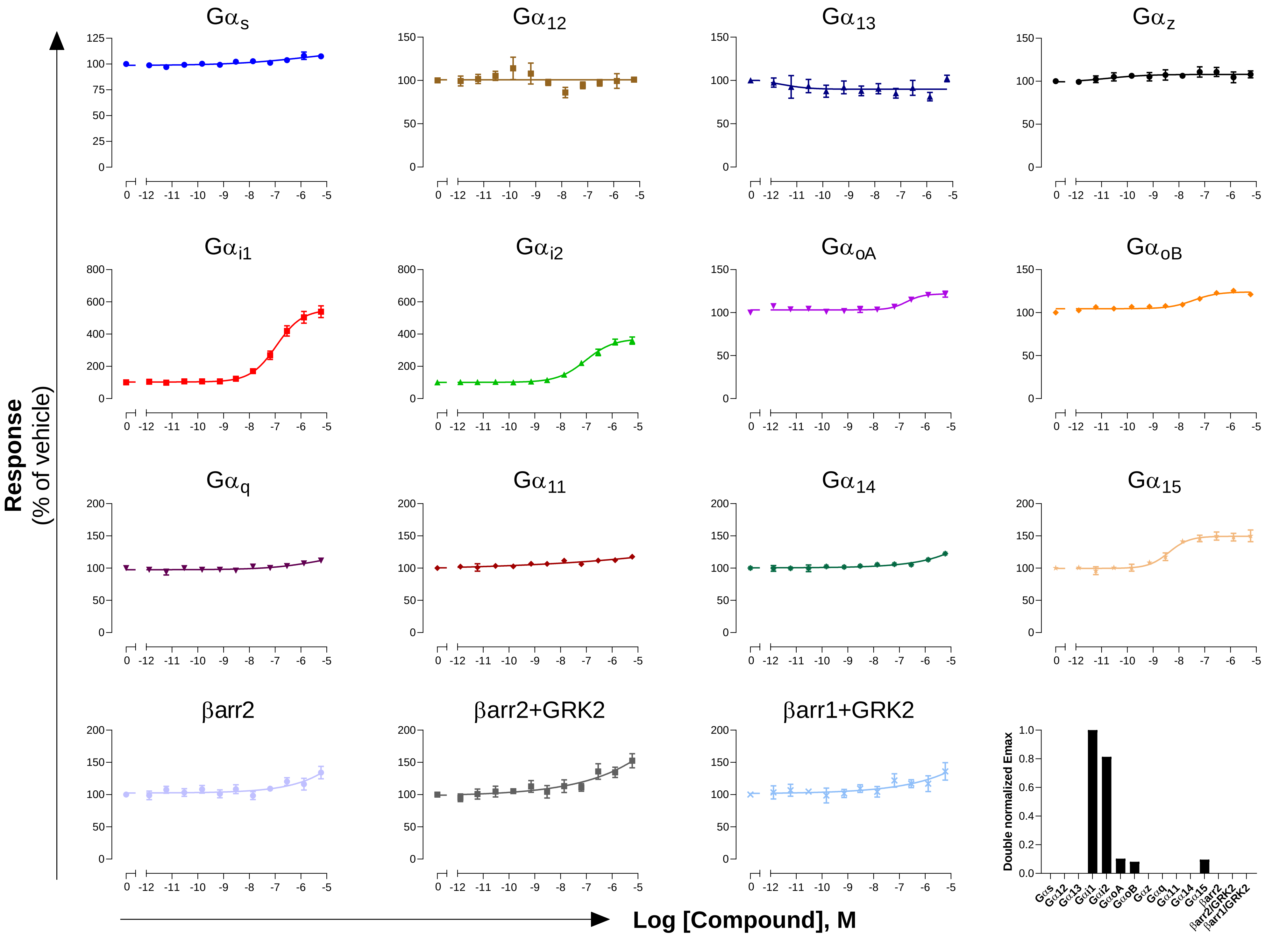

**GPCR: AMY$_3$**

**Ligand: Amylin**

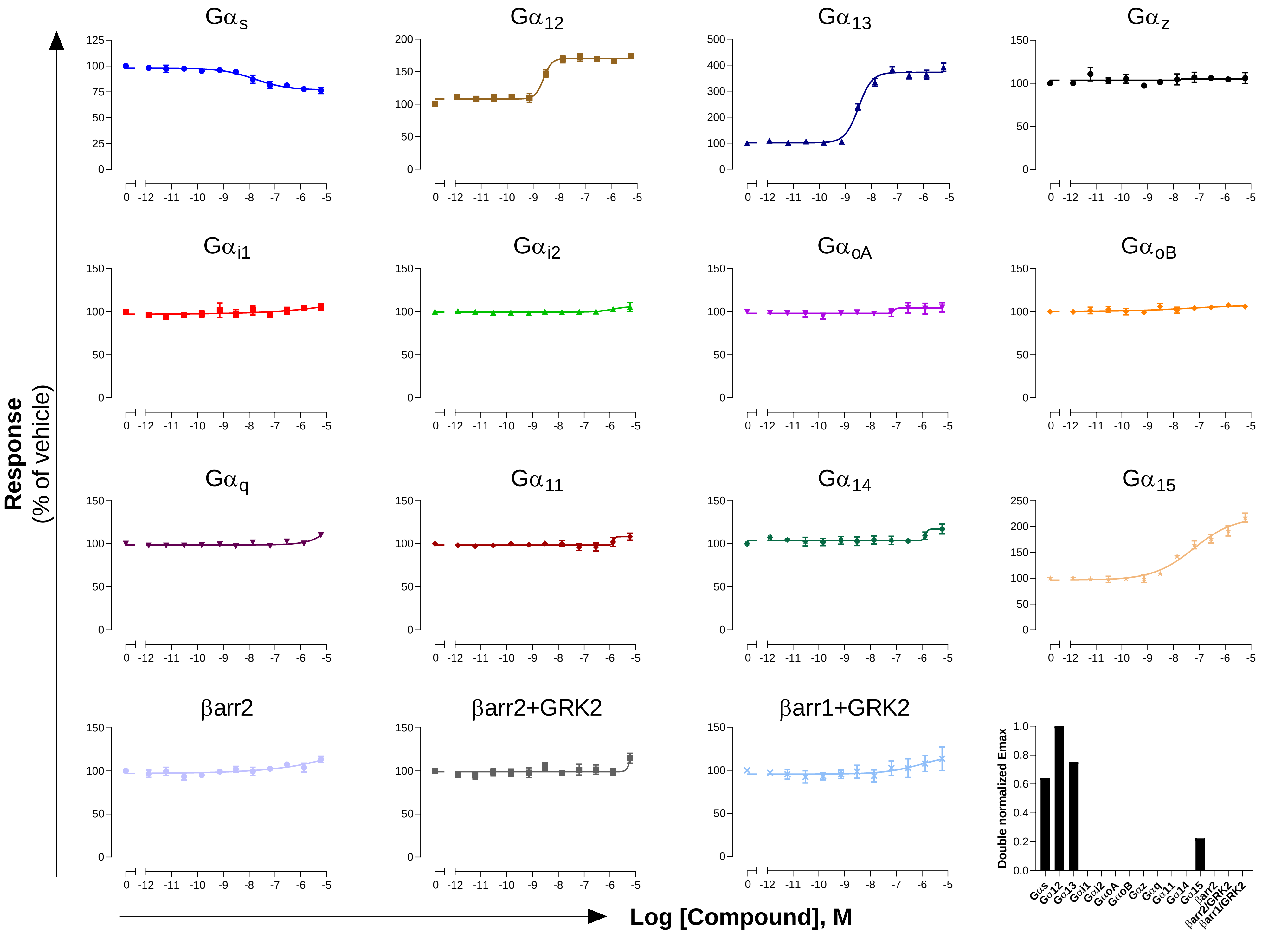

### GPCR: APJ
### Ligand: [Pyr1]-Apelin 13

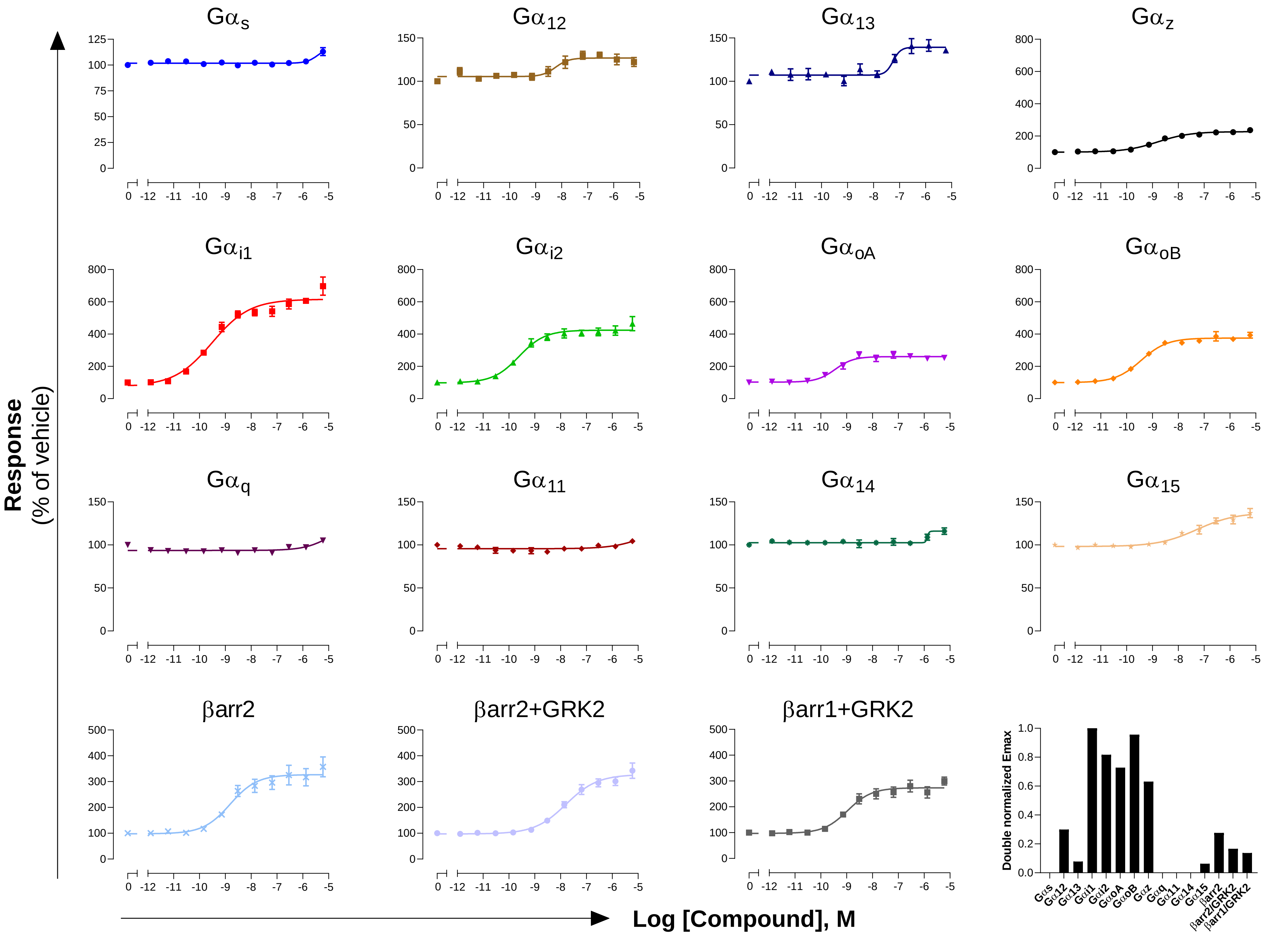

**GPCR: $AT_1$**

**Ligand: Angiotensin II**

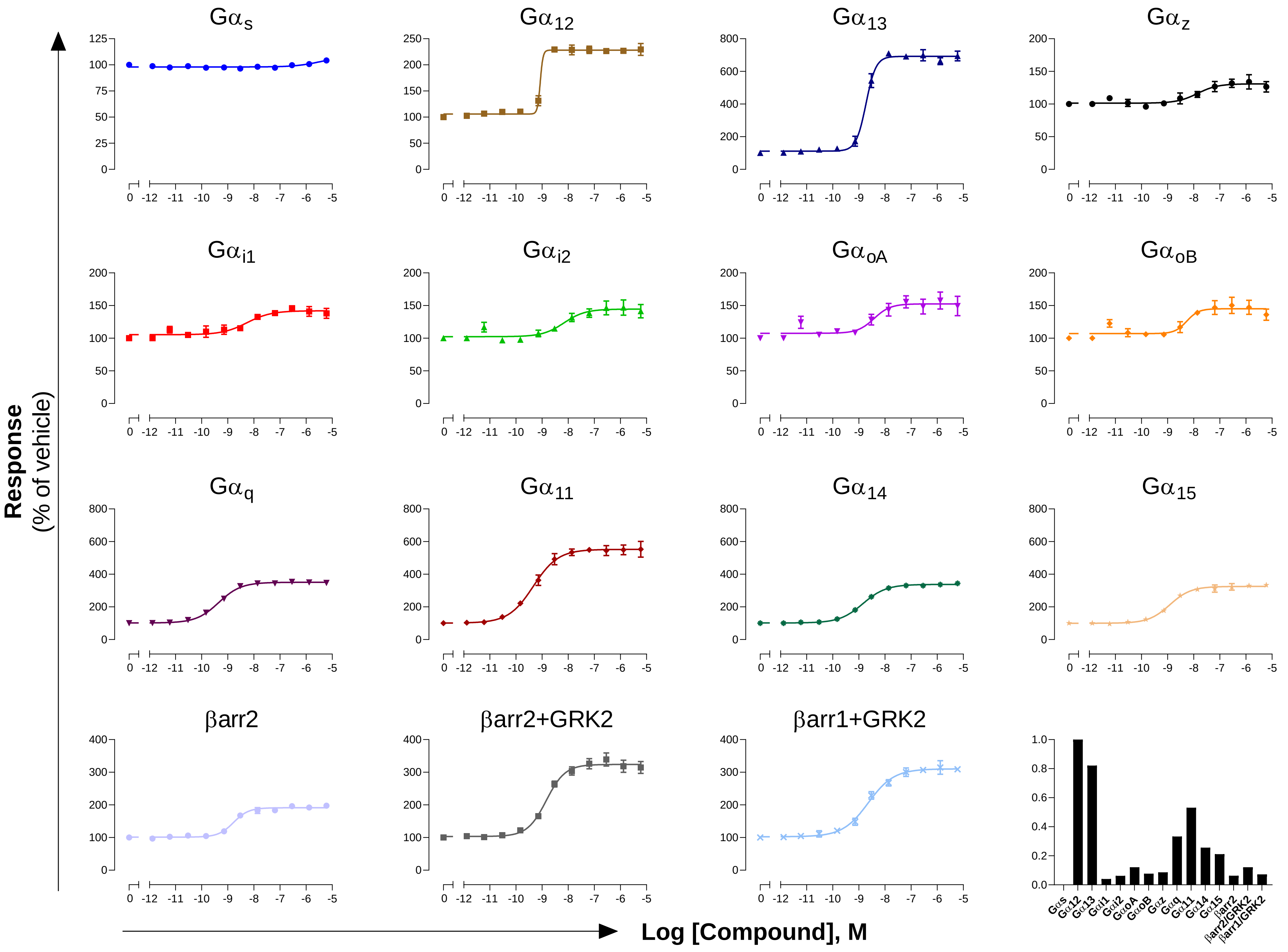

**GPCR: $B_1$**

**Ligand: Kallidin**

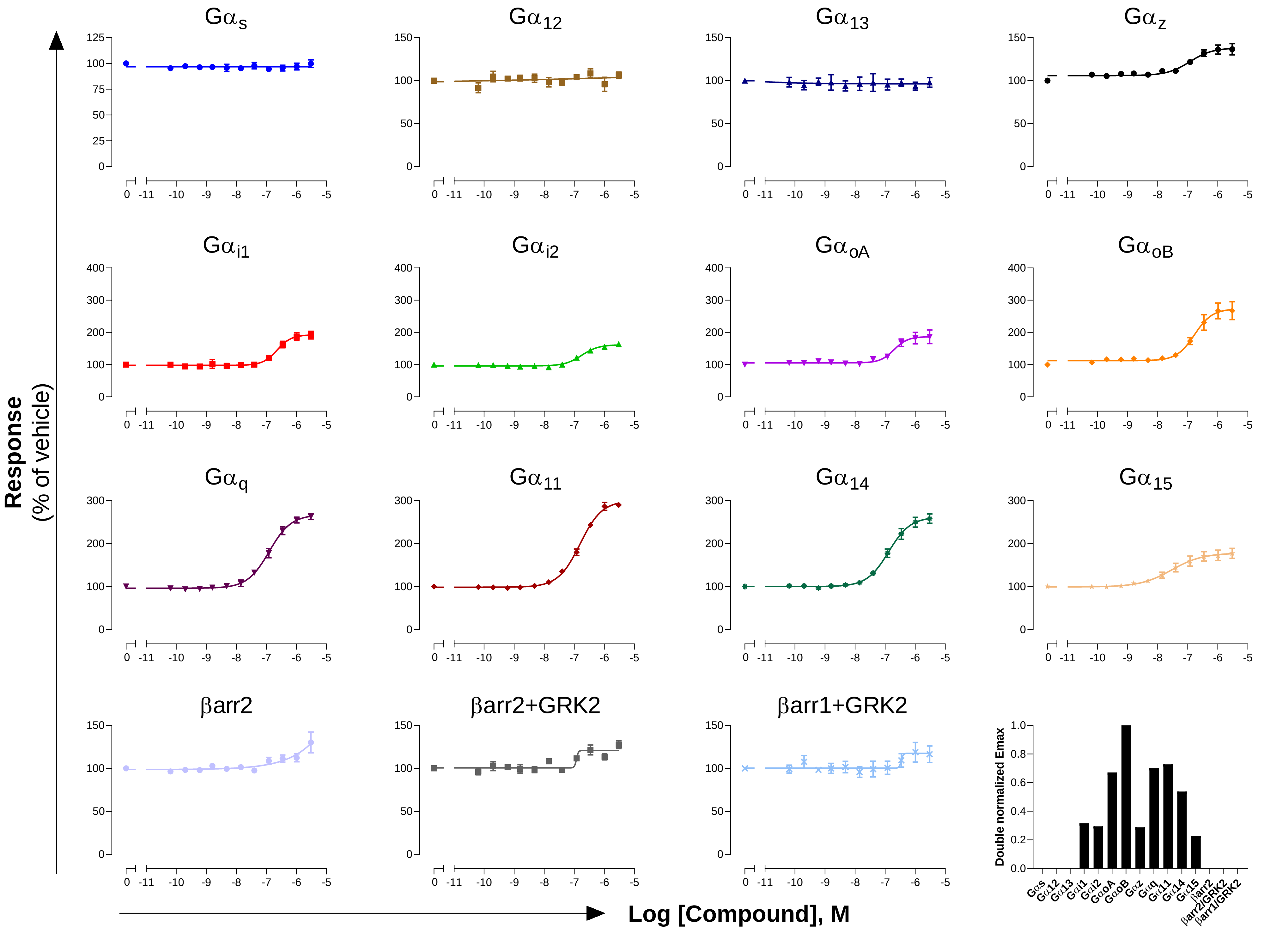

### GPCR: $B_2$
### Ligand: Kallidin

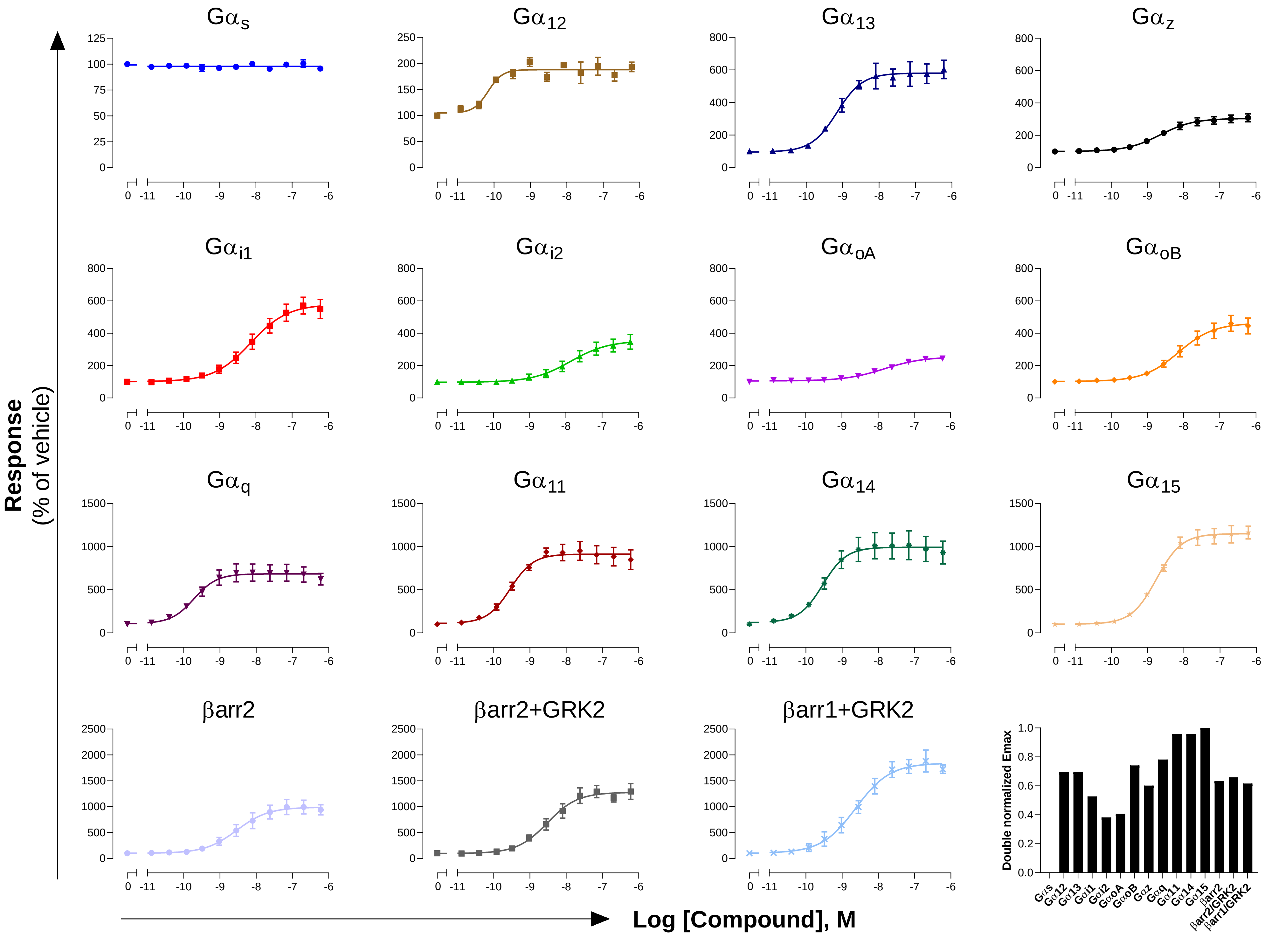

**GPCR: BLT$_1$**
**Ligand: LTB4**

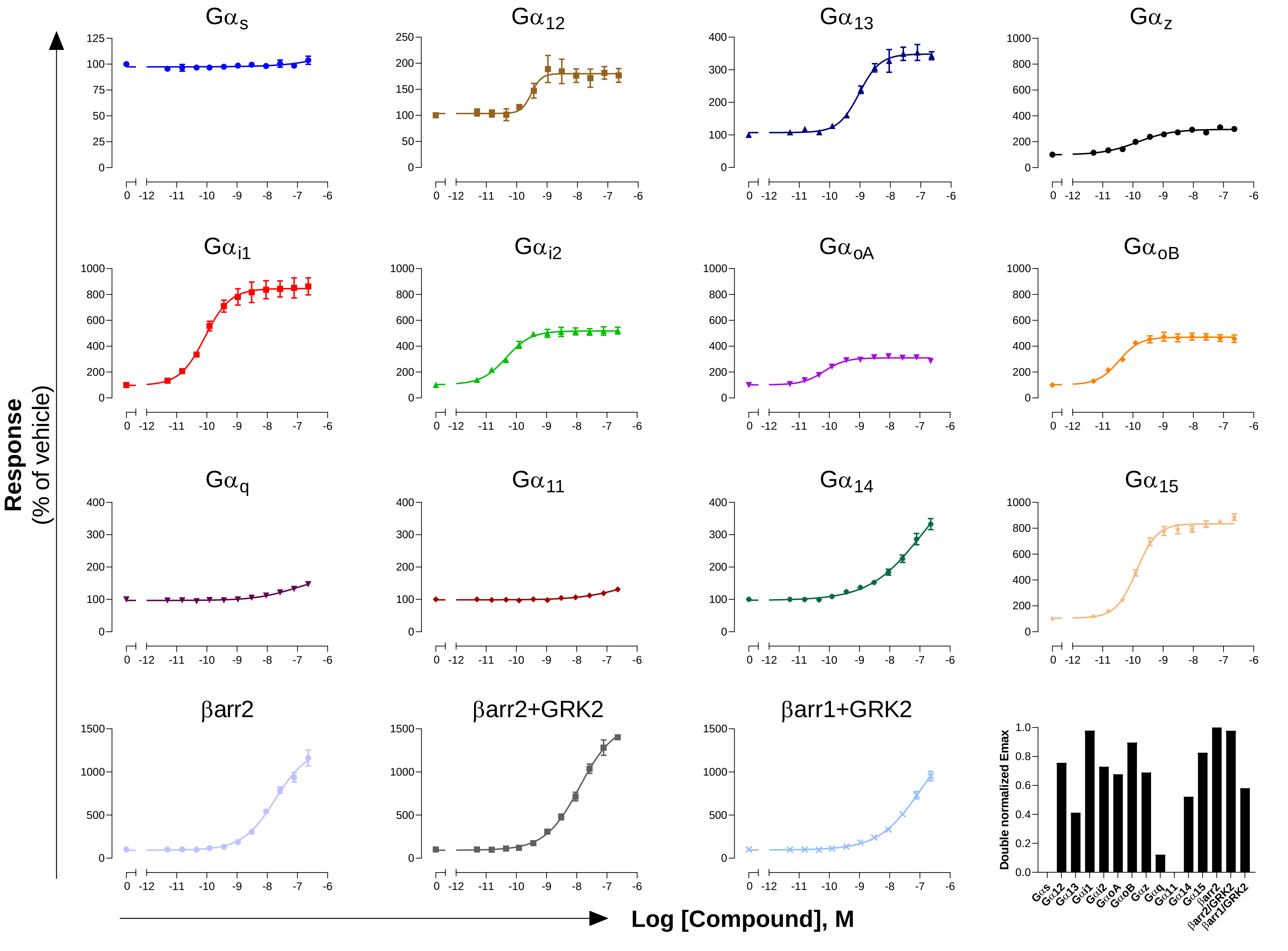

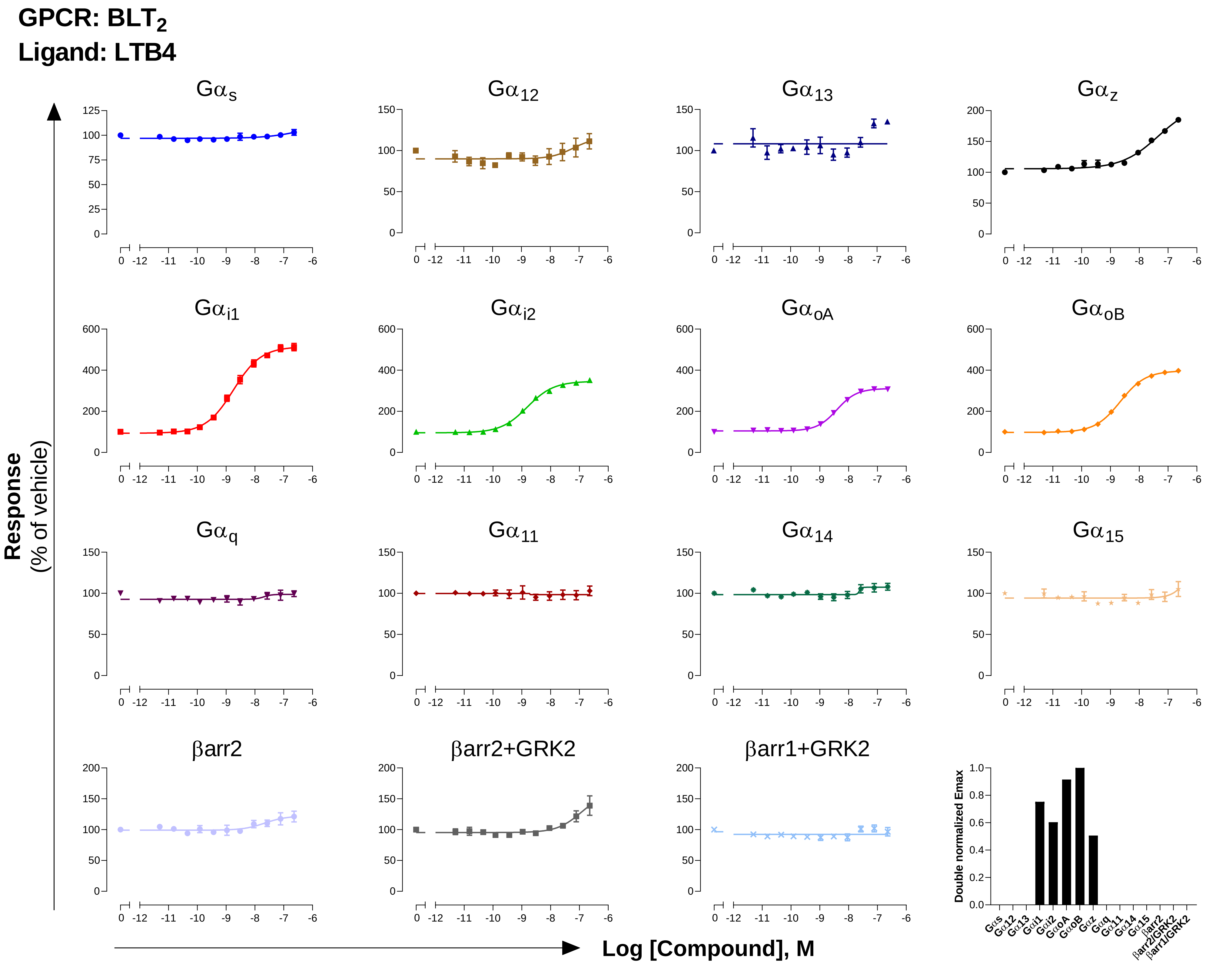

GPCR: BLT2
Ligand: LTB4
Gαs
Gα12
Gα13
Gαz
Gαi1
Gαi2
GαoA
GαoB
Gαq
Gα11
Gα14
Gα15
βarr2
βarr2+GRK2
βarr1+GRK2
Response
(% of vehicle)
Log [Compound], M
Double normalized Emax

**GPCR: C5a$_1$**

**Ligand: C5a**

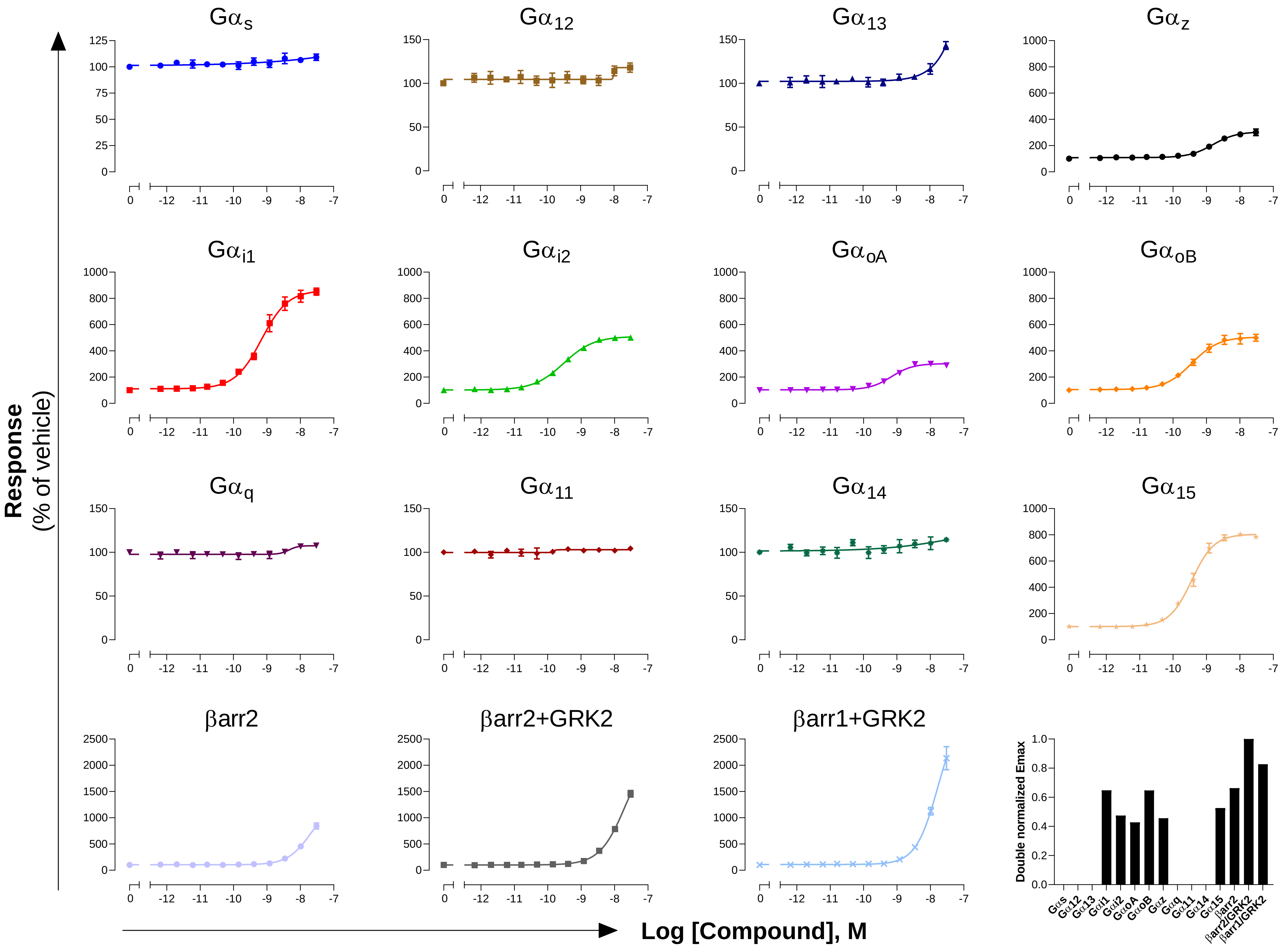

**GPCR: $CB_1$**

**Ligand: WIN55,212-2**

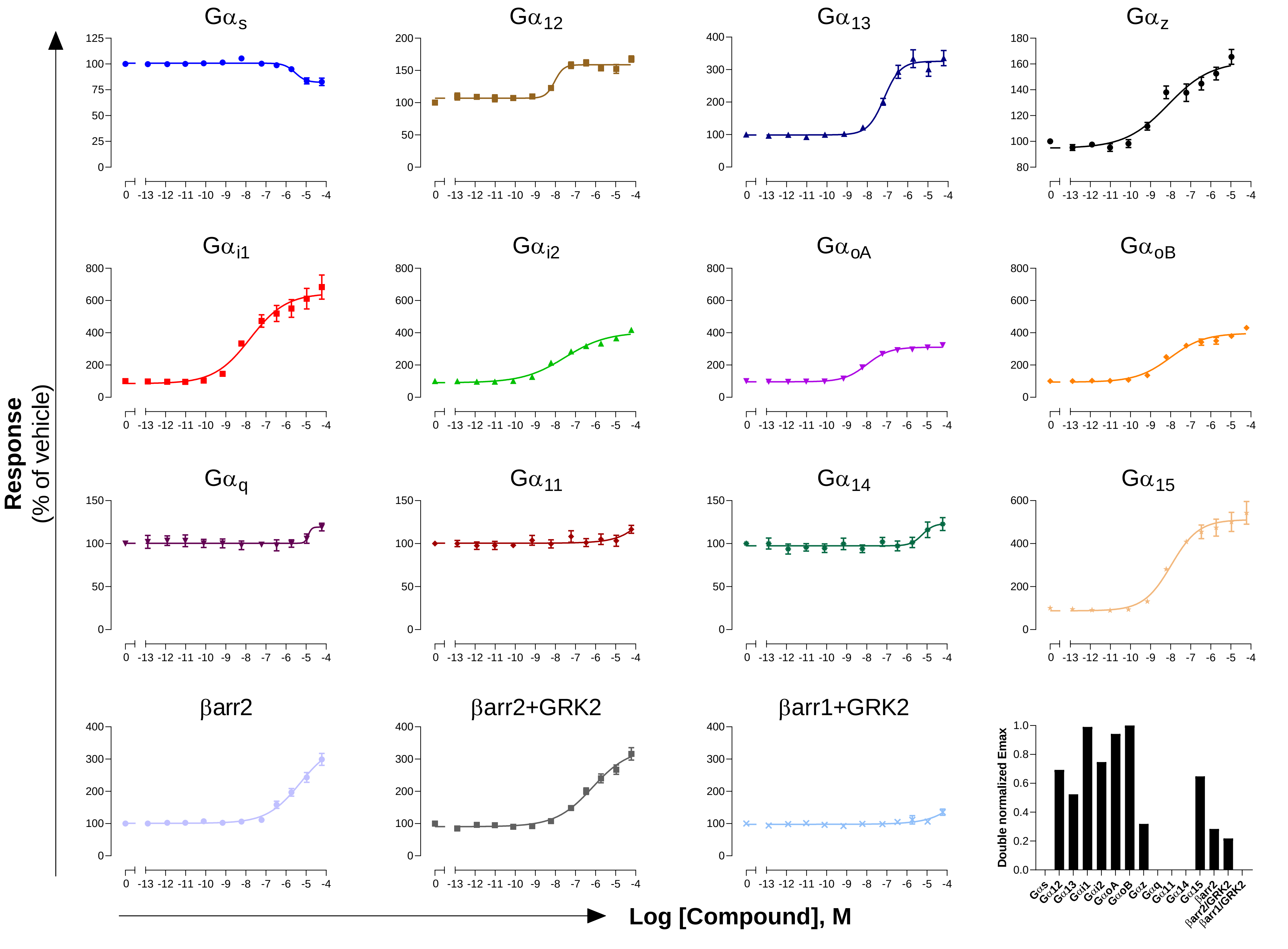

### GPCR: $CB_2$
### Ligand: WIN55,212-2

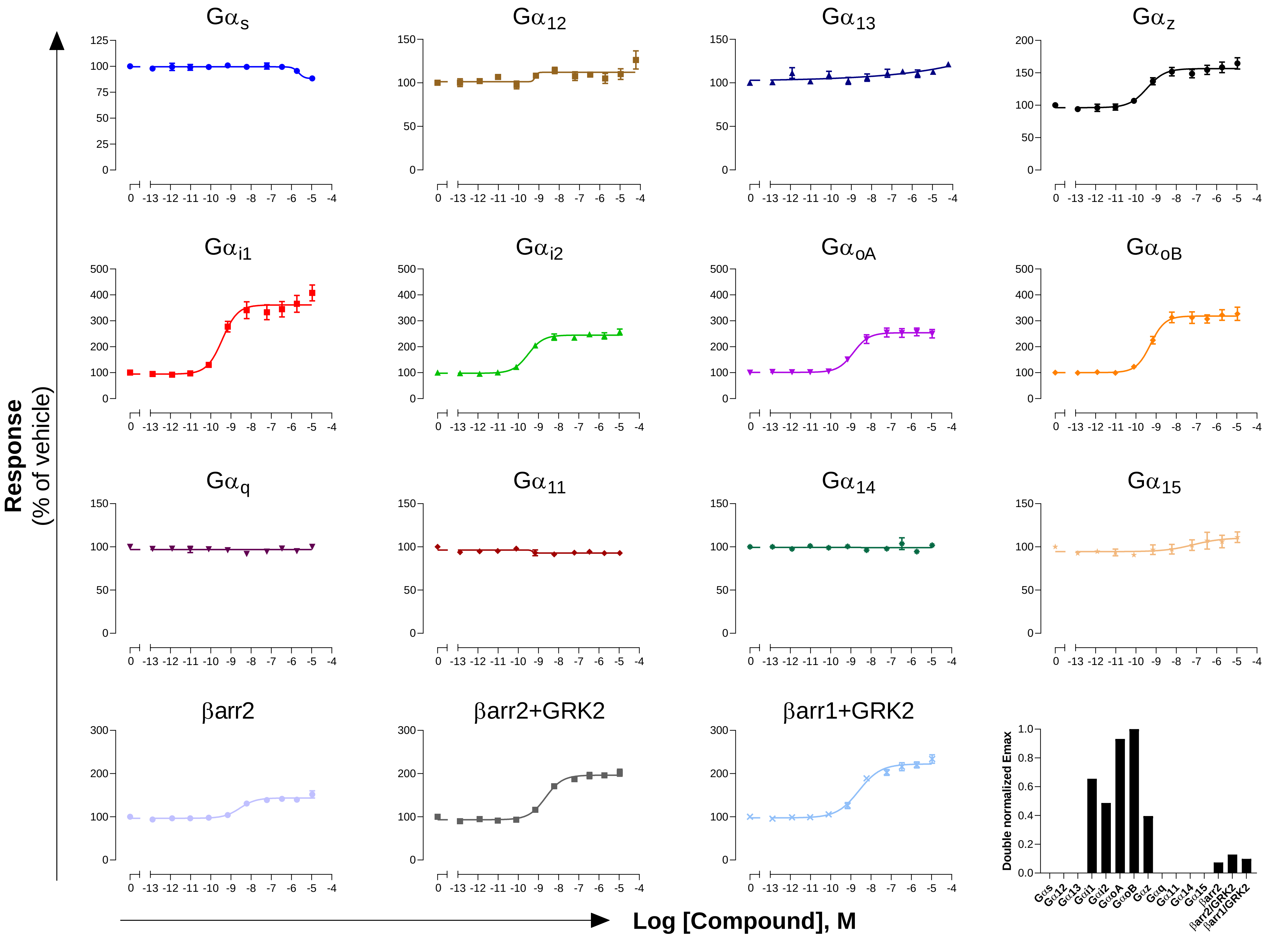

**GPCR: CCK$_1$**
**Ligand: CCK8**

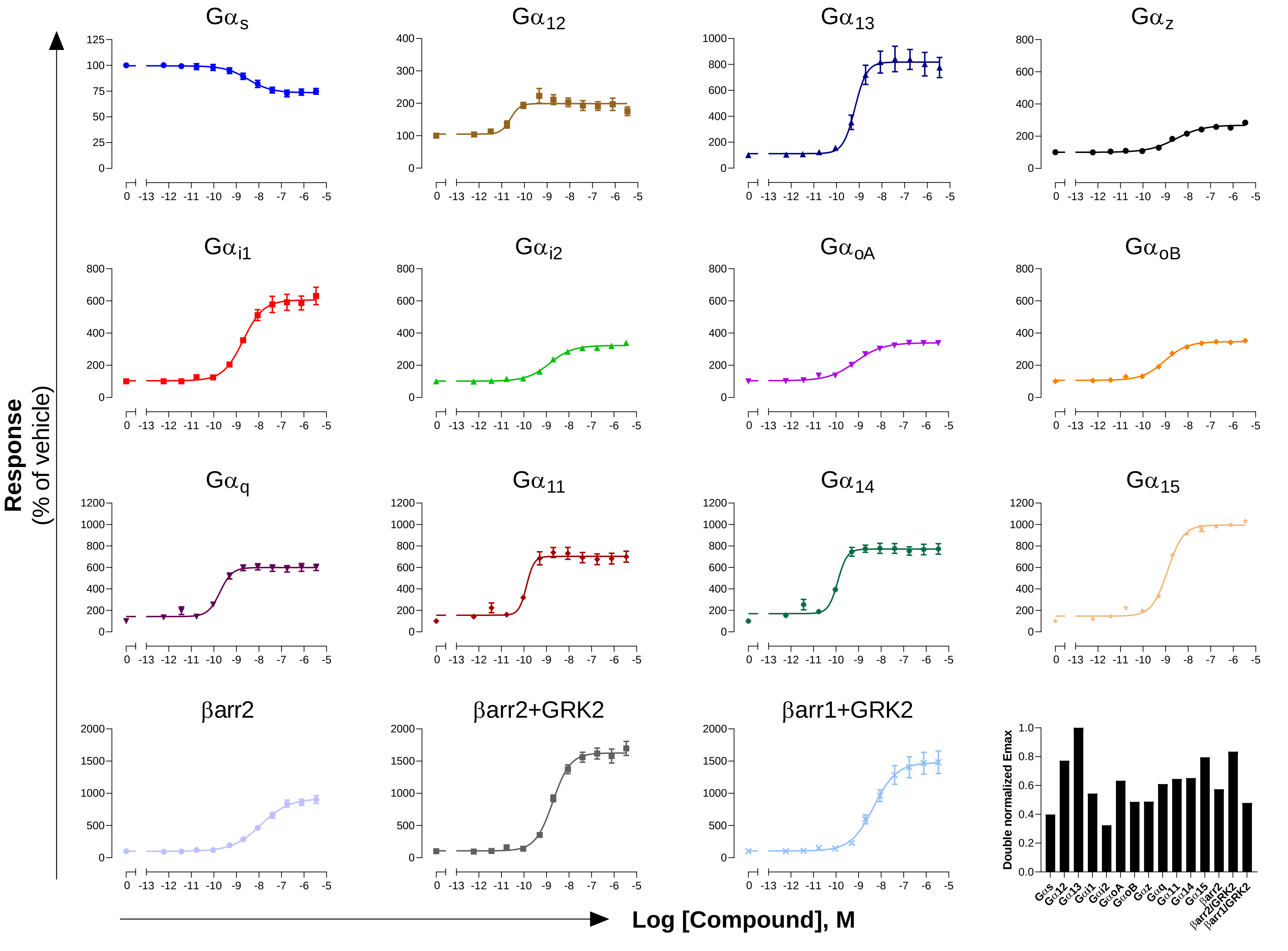

**GPCR: CCR5**
**Ligand: CCL3**

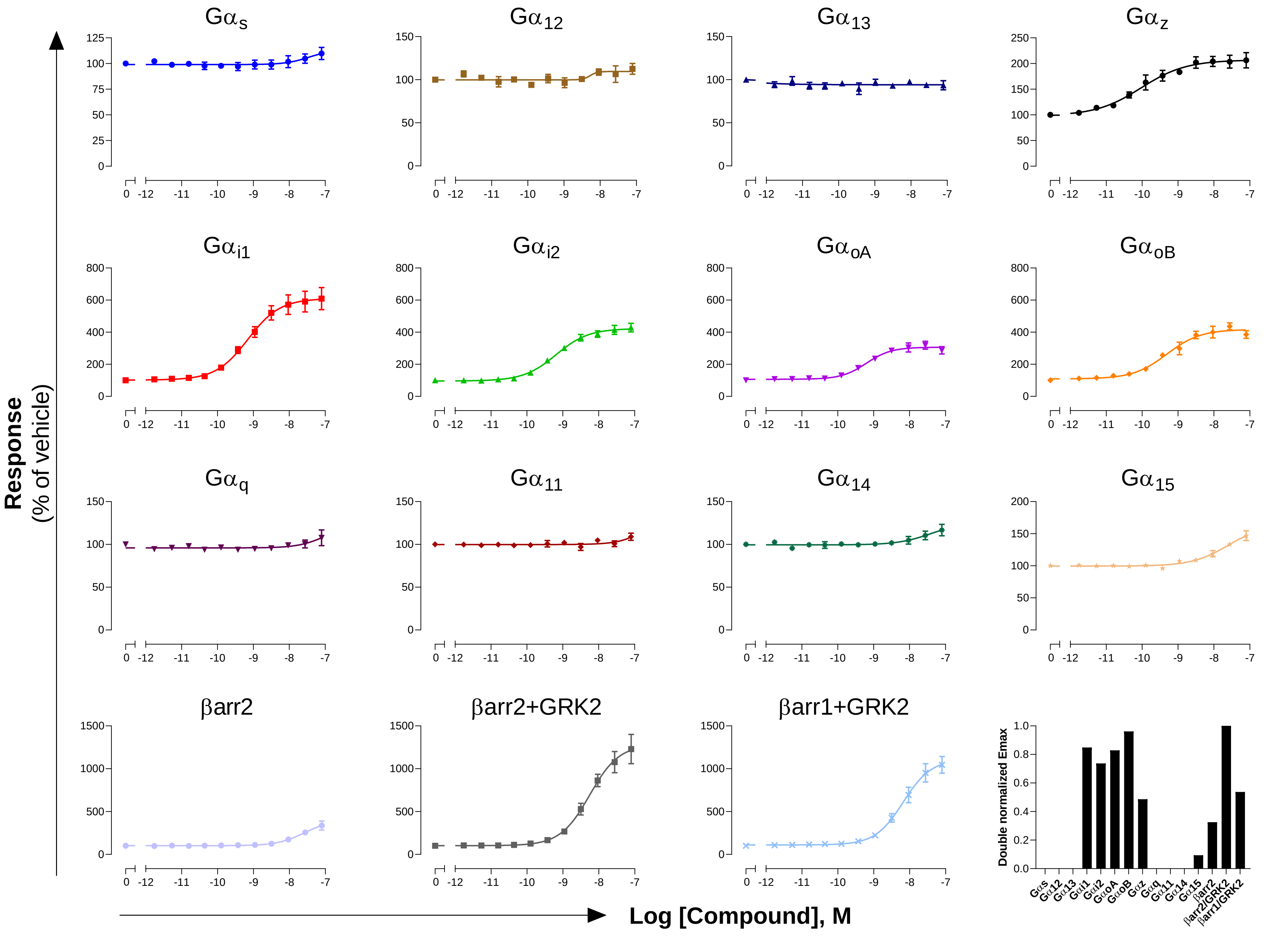

**GPCR: CCR6**
**Ligand: CCL20**

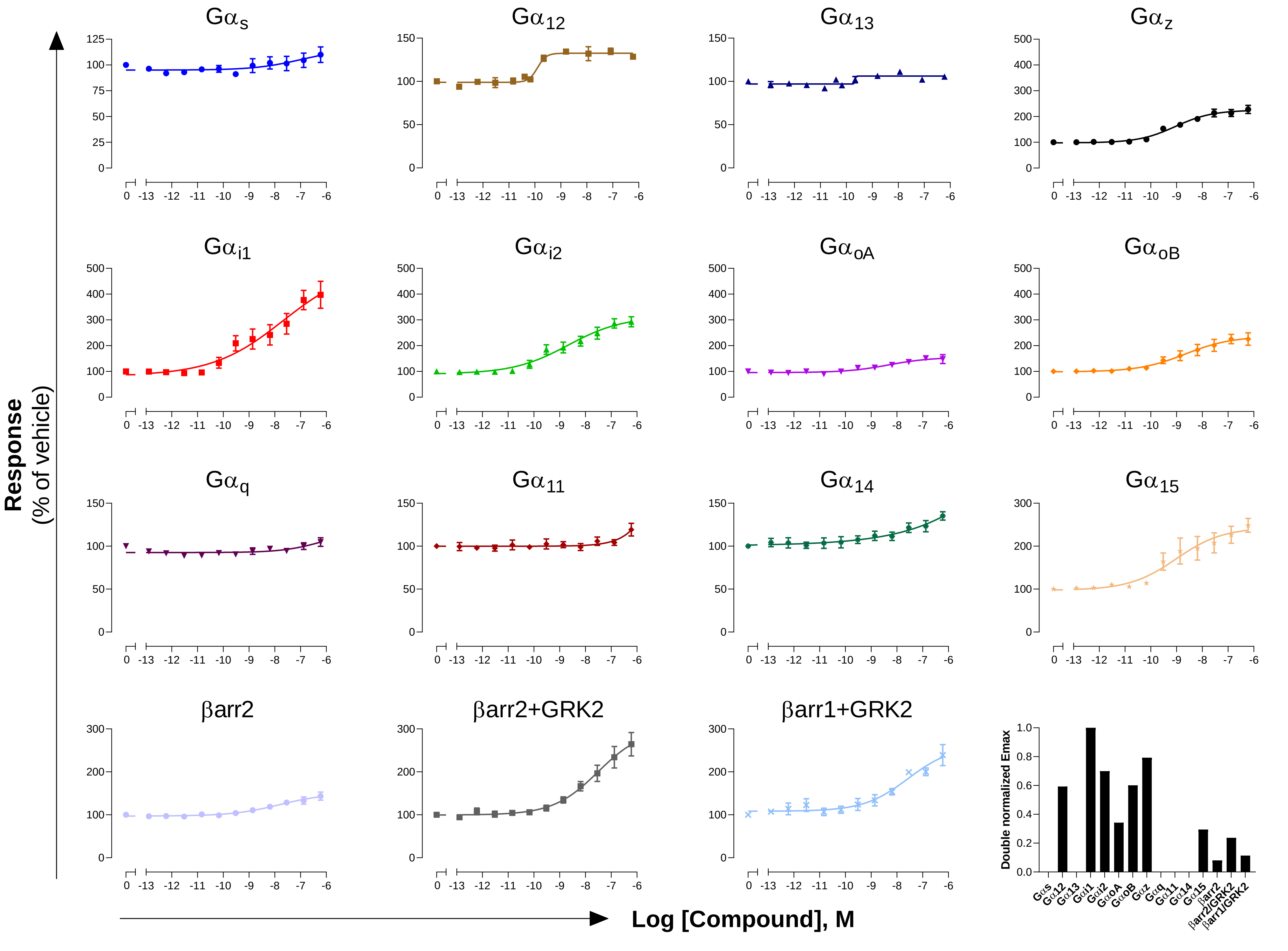

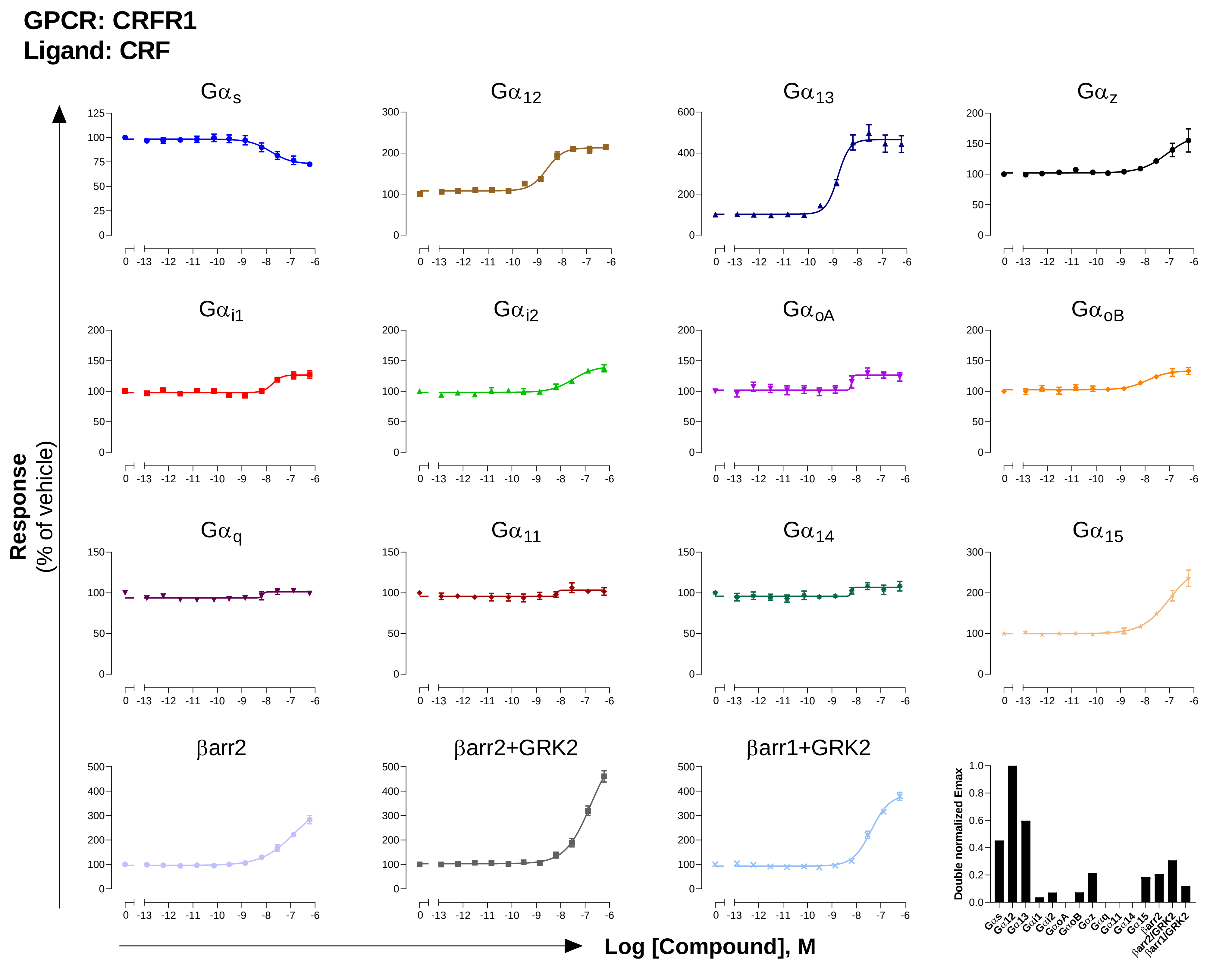
GPCR: CRFR1
Ligand: CRF
Response
(% of vehicle)
Log [Compound], M
Gαs
Gα12
Gα13
Gαz
Gαi1
Gαi2
GαoA
GαoB
Gαq
Gα11
Gα14
Gα15
βarr2
βarr2+GRK2
βarr1+GRK2
Double normalized Emax

**GPCR: CRFR2**
**Ligand: Urocortin II**

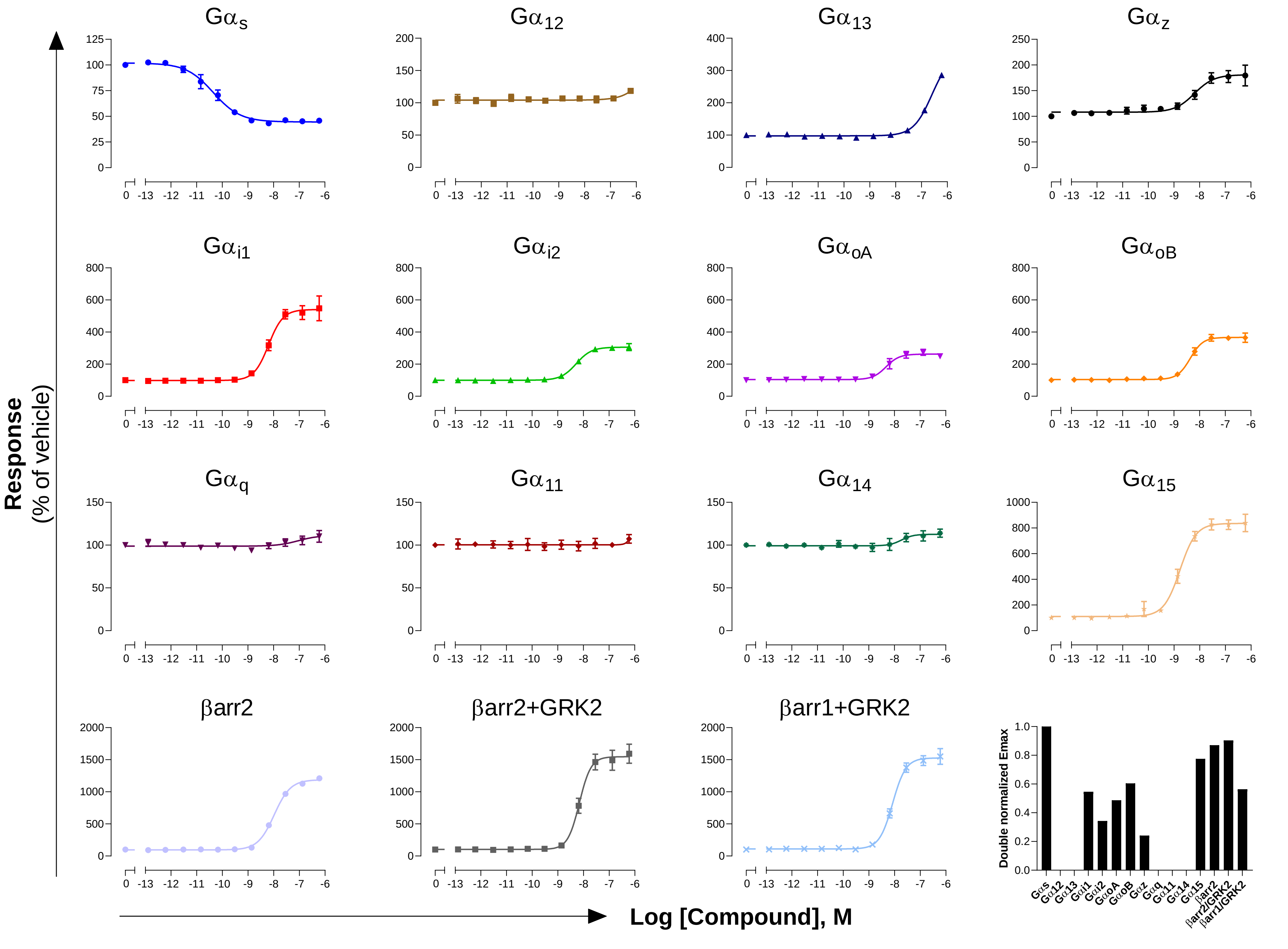

**GPCR: CTR**
**Ligand: Calcitonin**

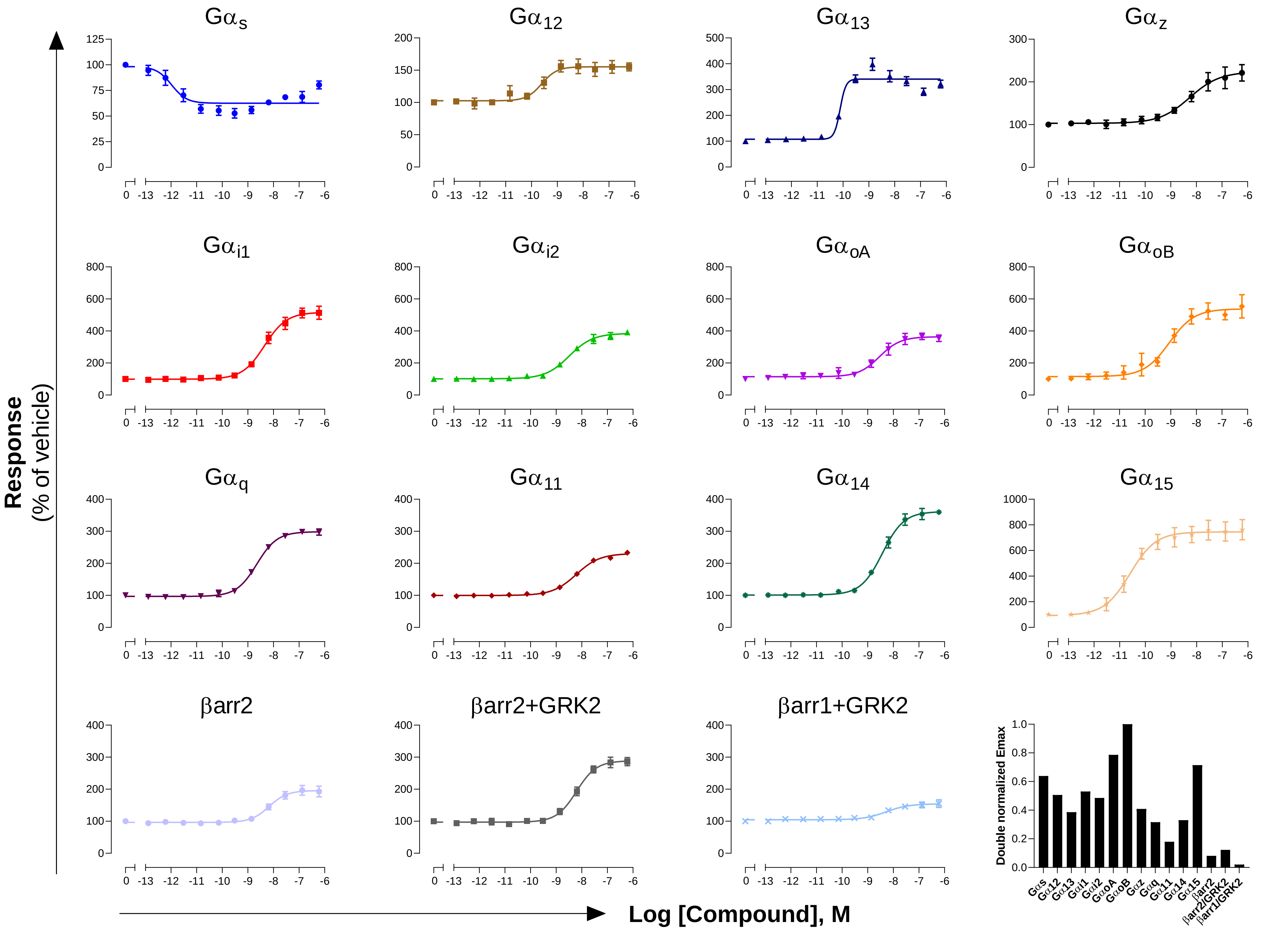

**GPCR: CXCR2**
**Ligand: CXCL8**

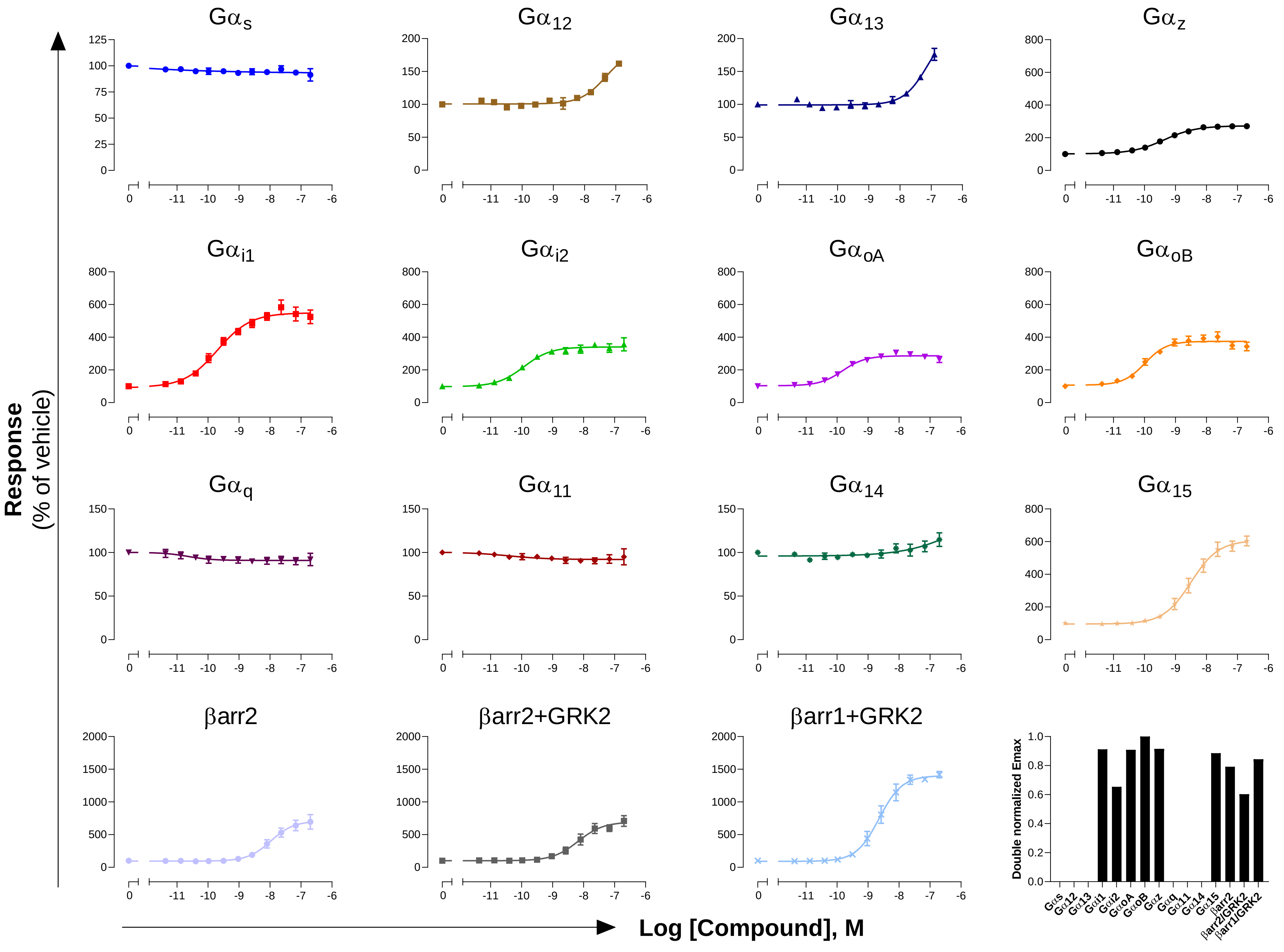

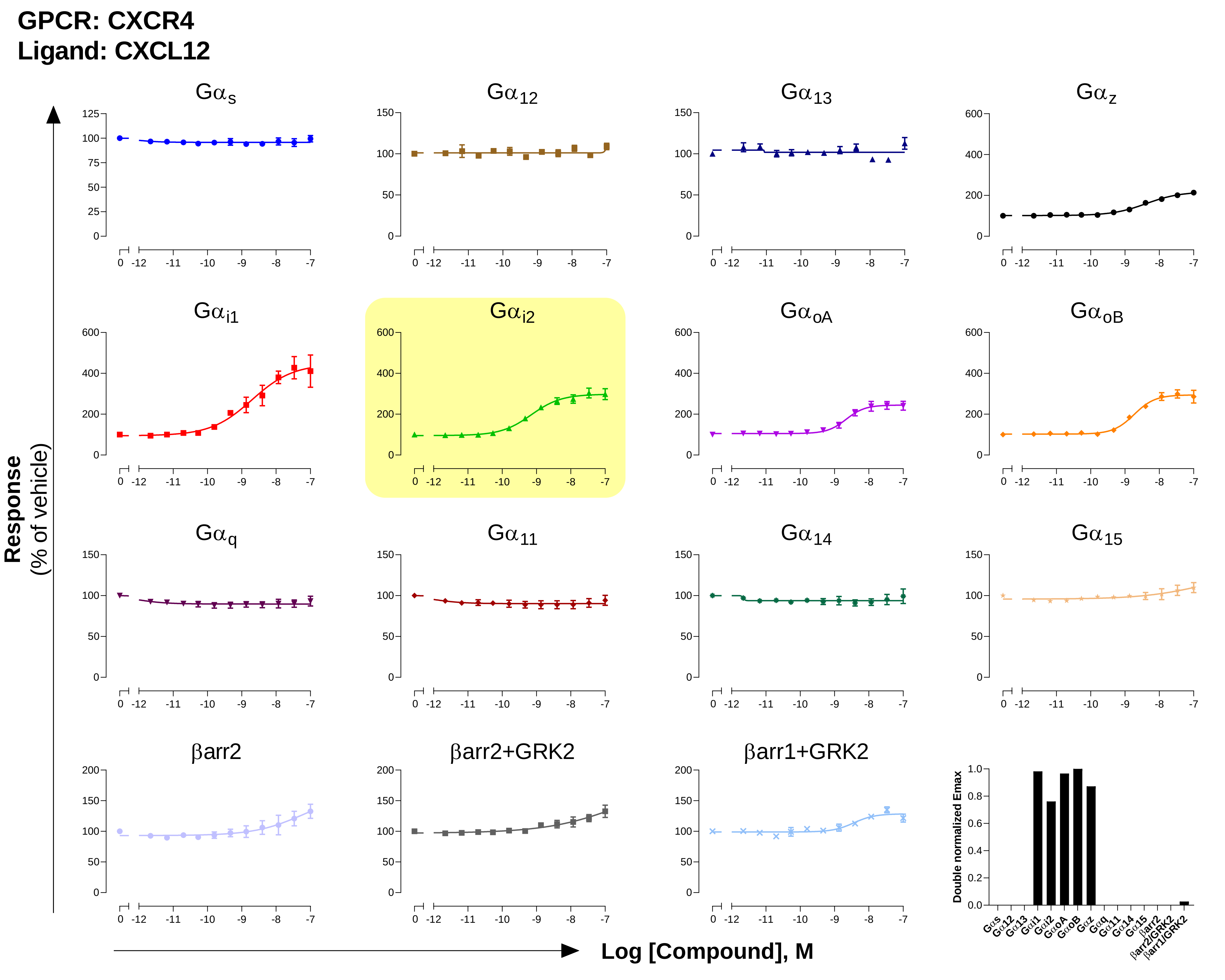
GPCR: CXCR4
Ligand: CXCL12
Gαs
Gα12
Gα13
Gαz
Gαi1
Gαi2
GαoA
GαoB
Gαq
Gα11
Gα14
Gα15
βarr2
βarr2+GRK2
βarr1+GRK2
Response
(% of vehicle)
Log [Compound], M
Double normalized Emax

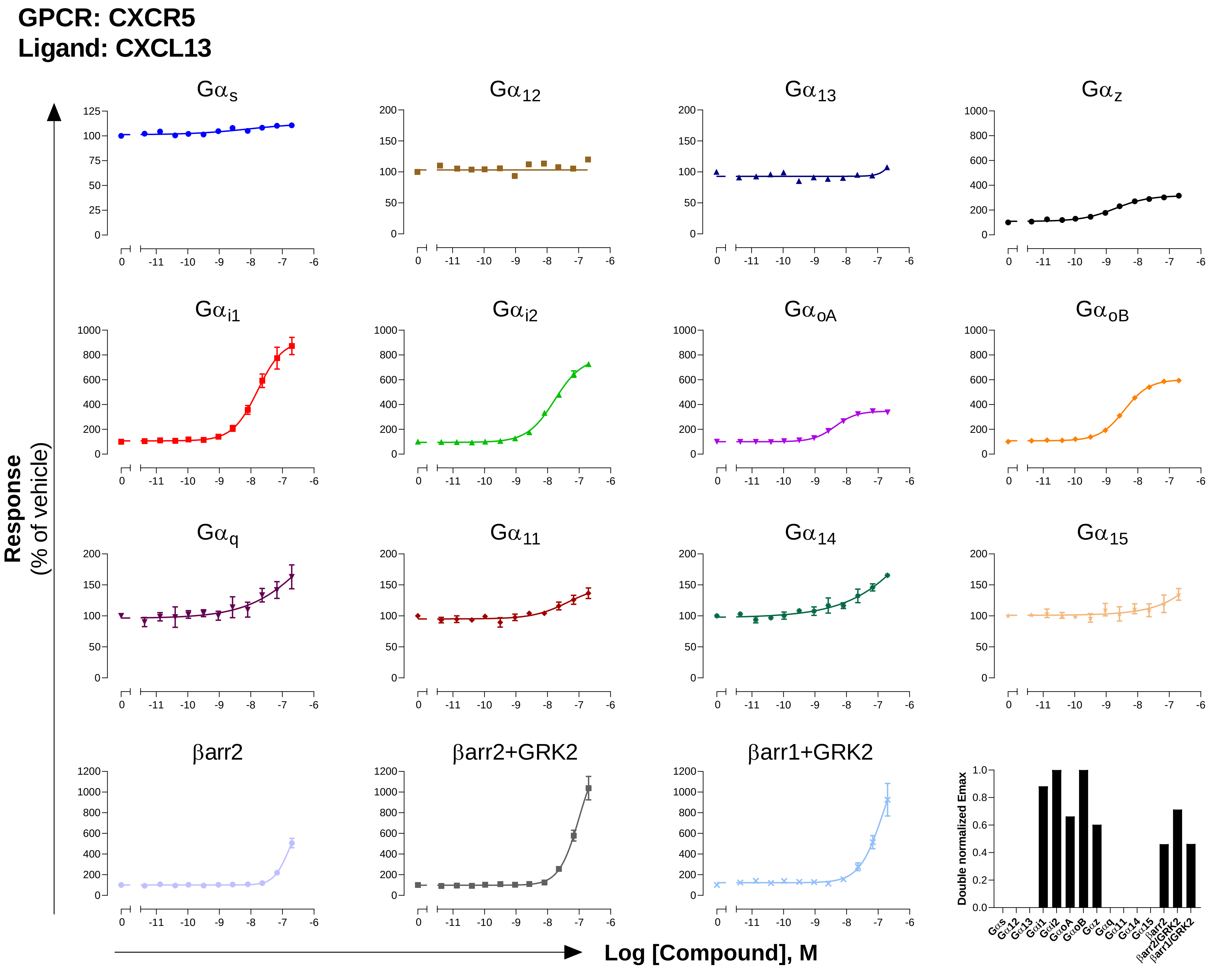

GPCR: CXCR5
Ligand: CXCL13
Response
(% of vehicle)
Log [Compound], M
Gαs
Gα12
Gα13
Gαz
Gαi1
Gαi2
GαoA
GαoB
Gαq
Gα11
Gα14
Gα15
βarr2
βarr2+GRK2
βarr1+GRK2
Double normalized Emax

**GPCR: $CysLT_1$**

**Ligand: LTD4**

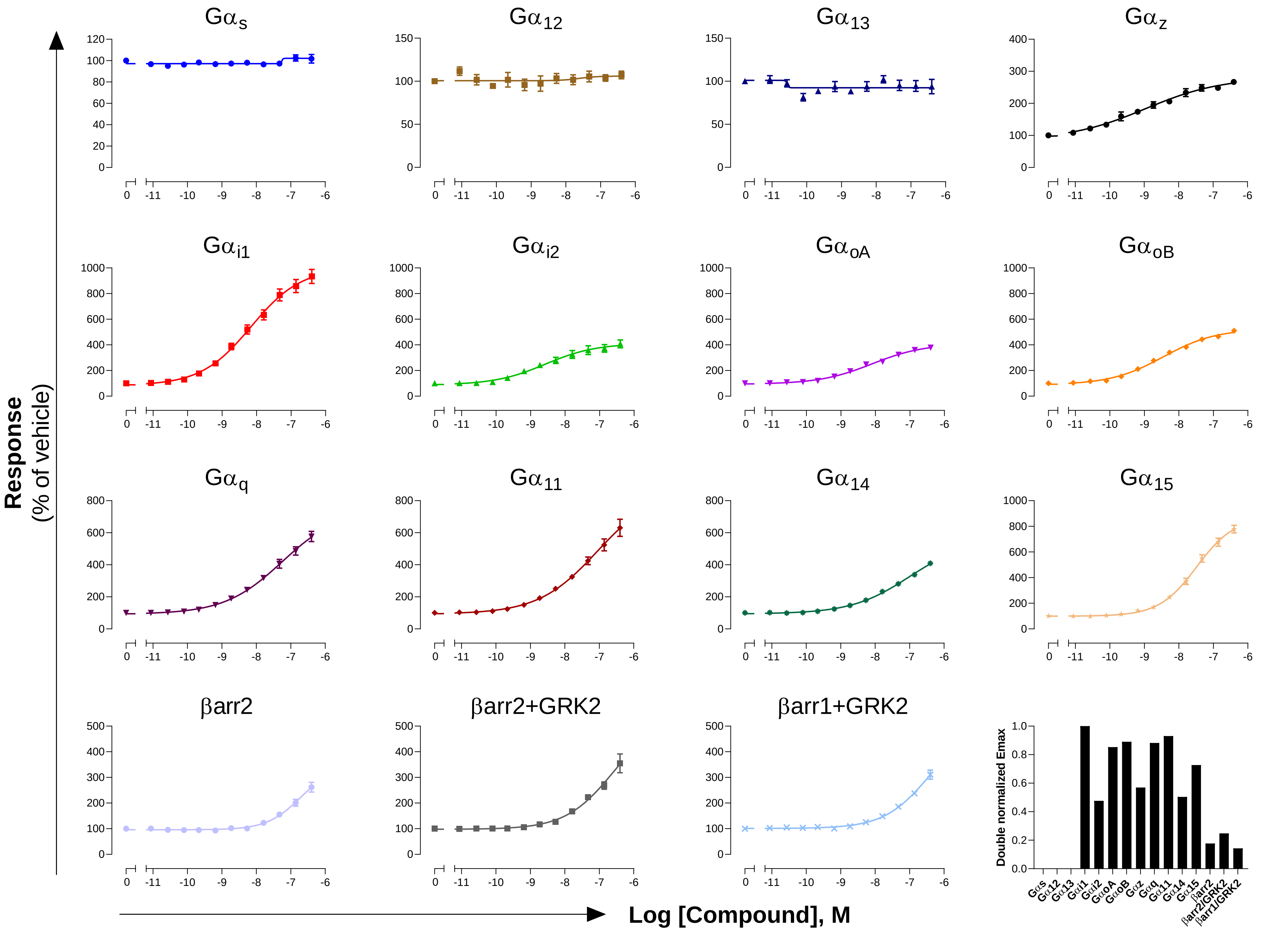

### GPCR: $CysLT_2$
### Ligand: LTD4

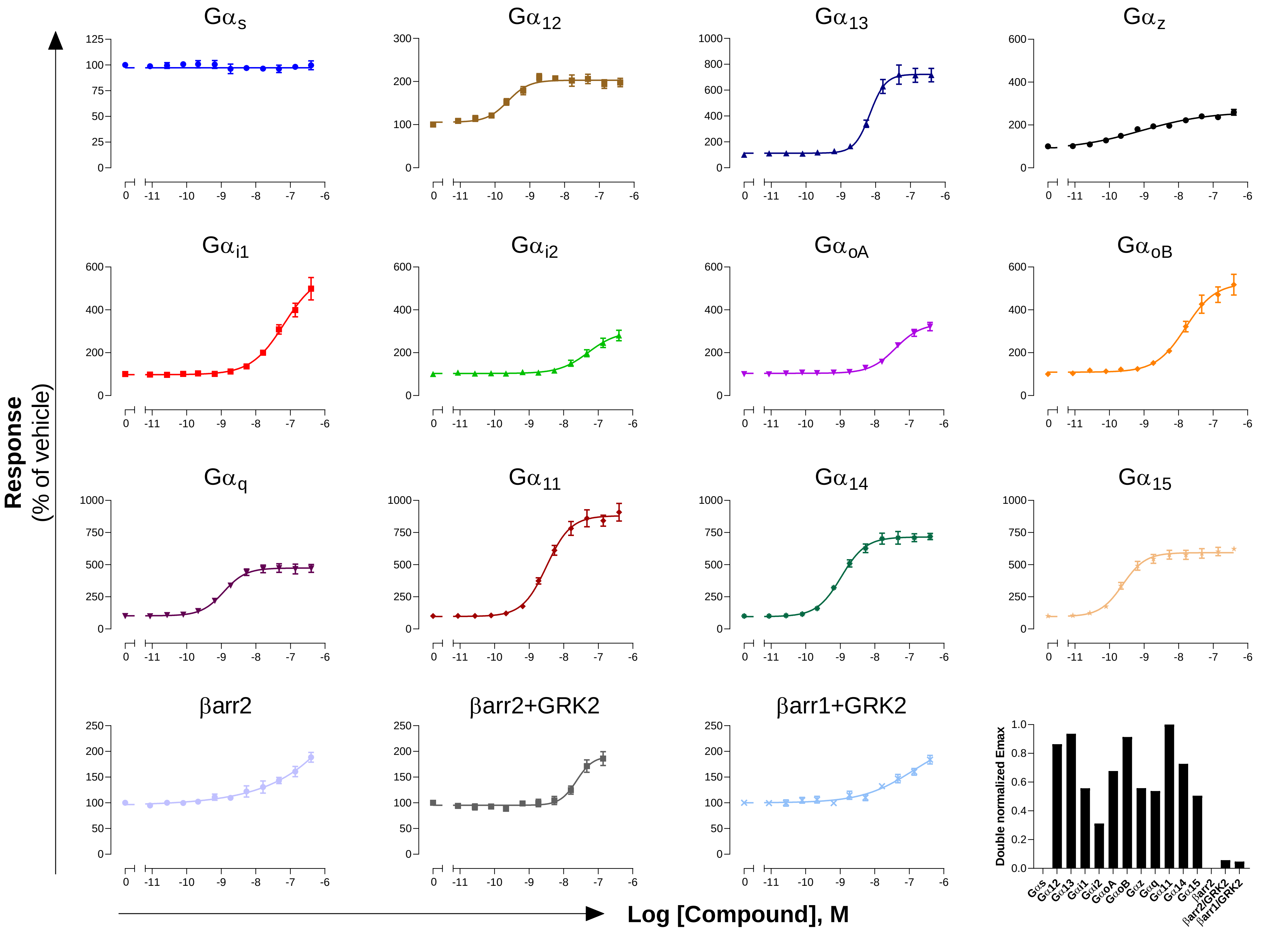

**GPCR: D$_1$**
**Ligand: Dopamine**

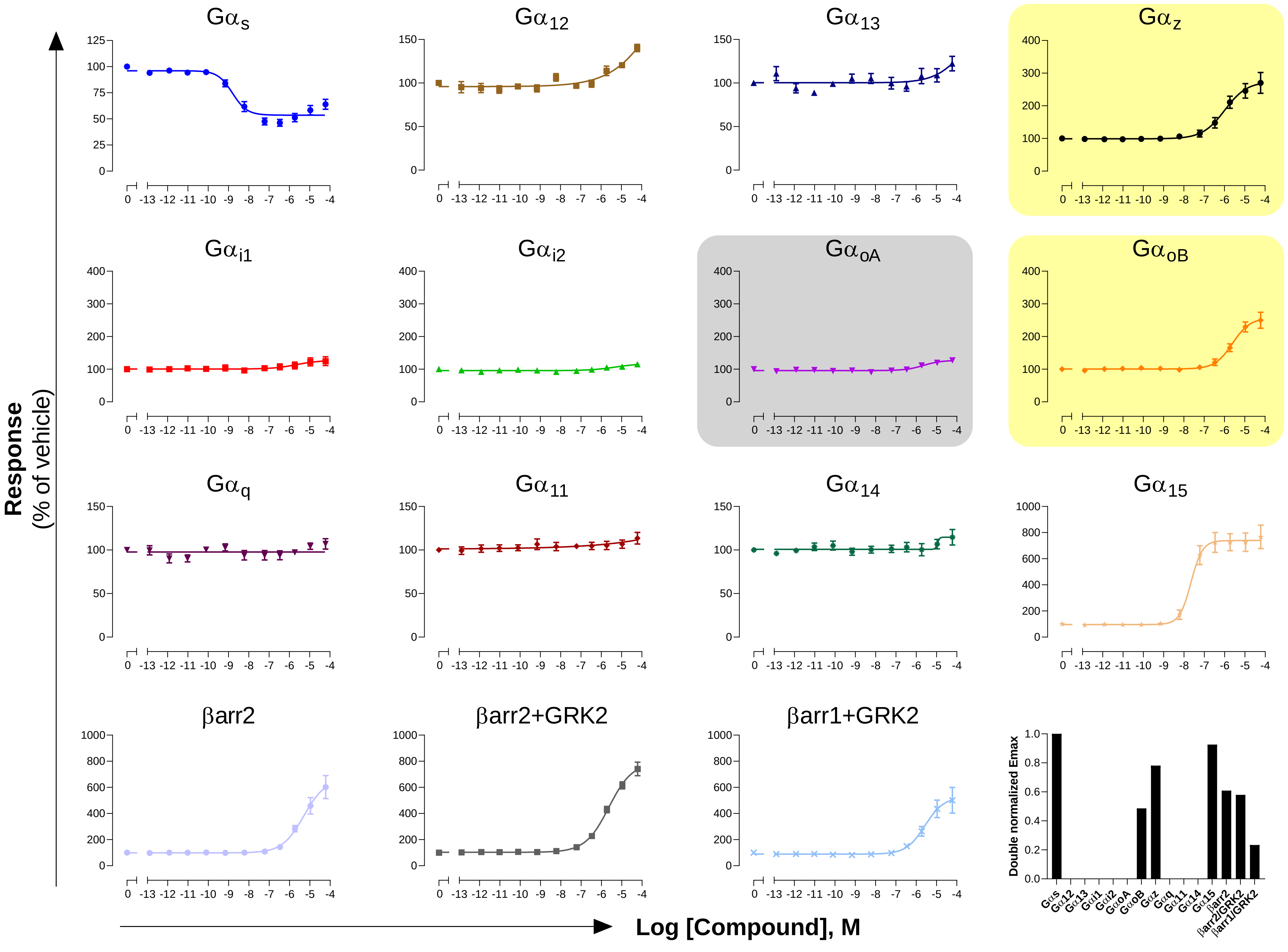

**GPCR: $D_2$**

**Ligand: Dopamine**

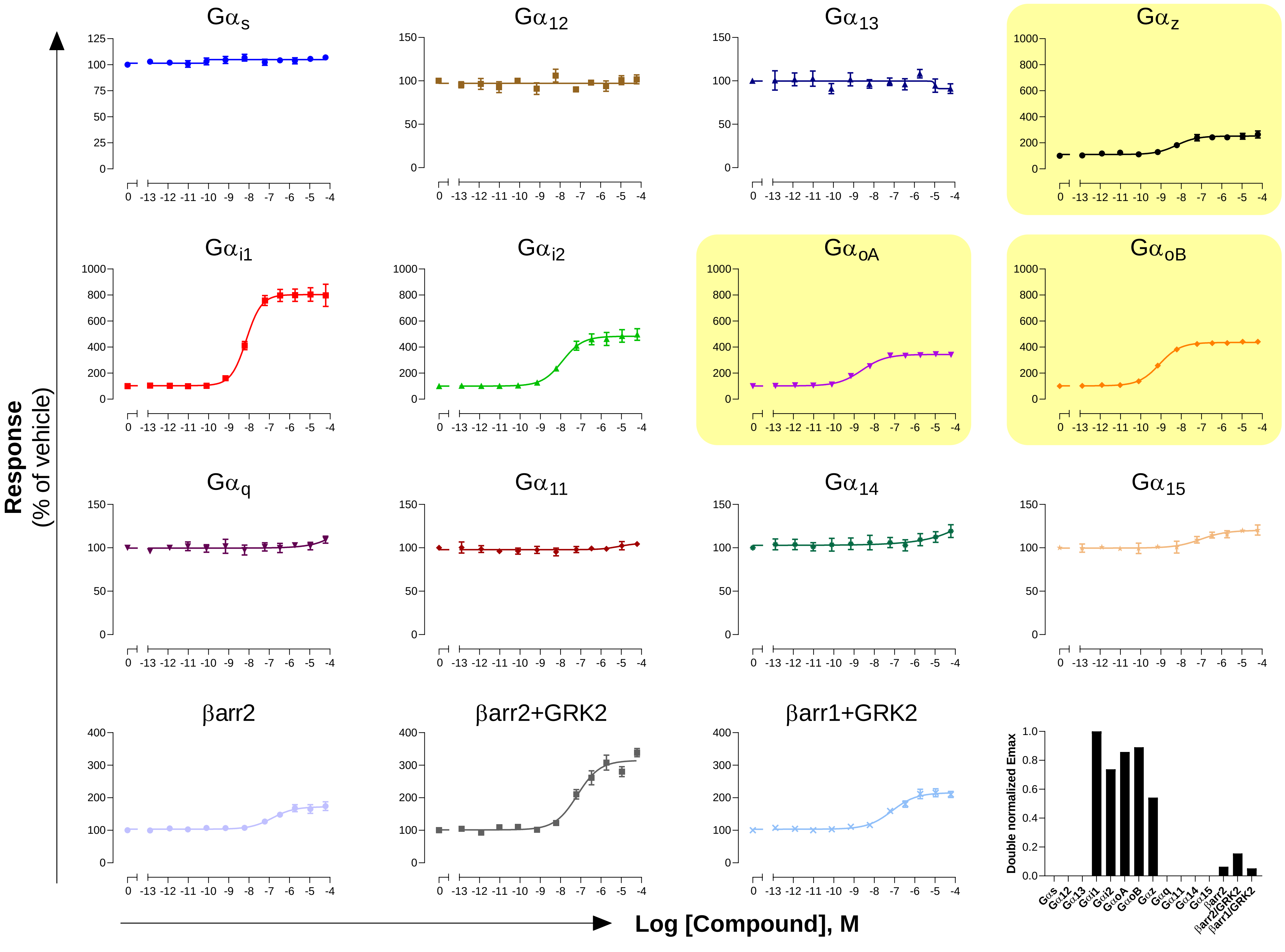

#### GPCR: $D_5$
#### Ligand: Dopamine

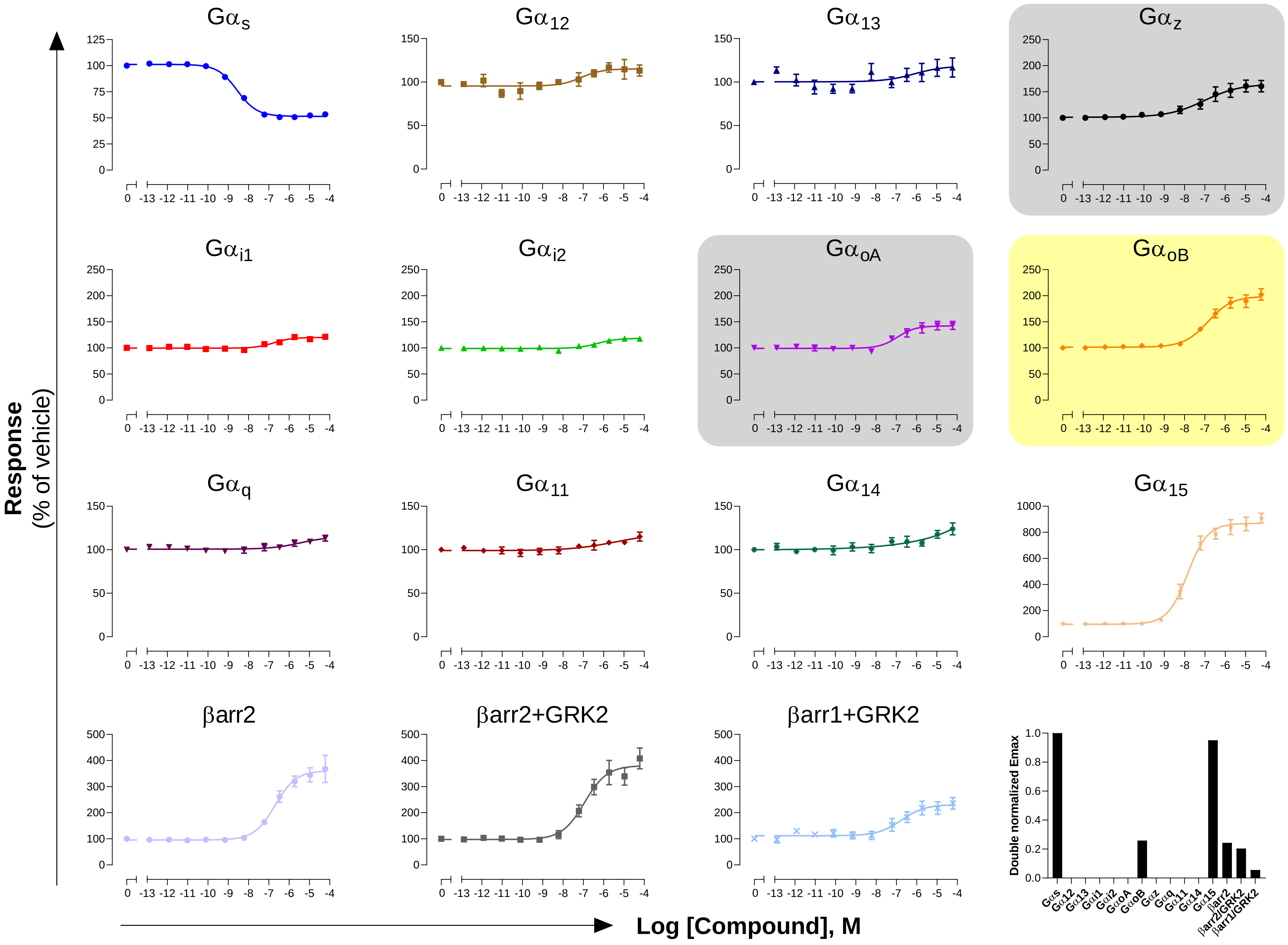

**GPCR: δOR**
**Ligand: SNC-80**

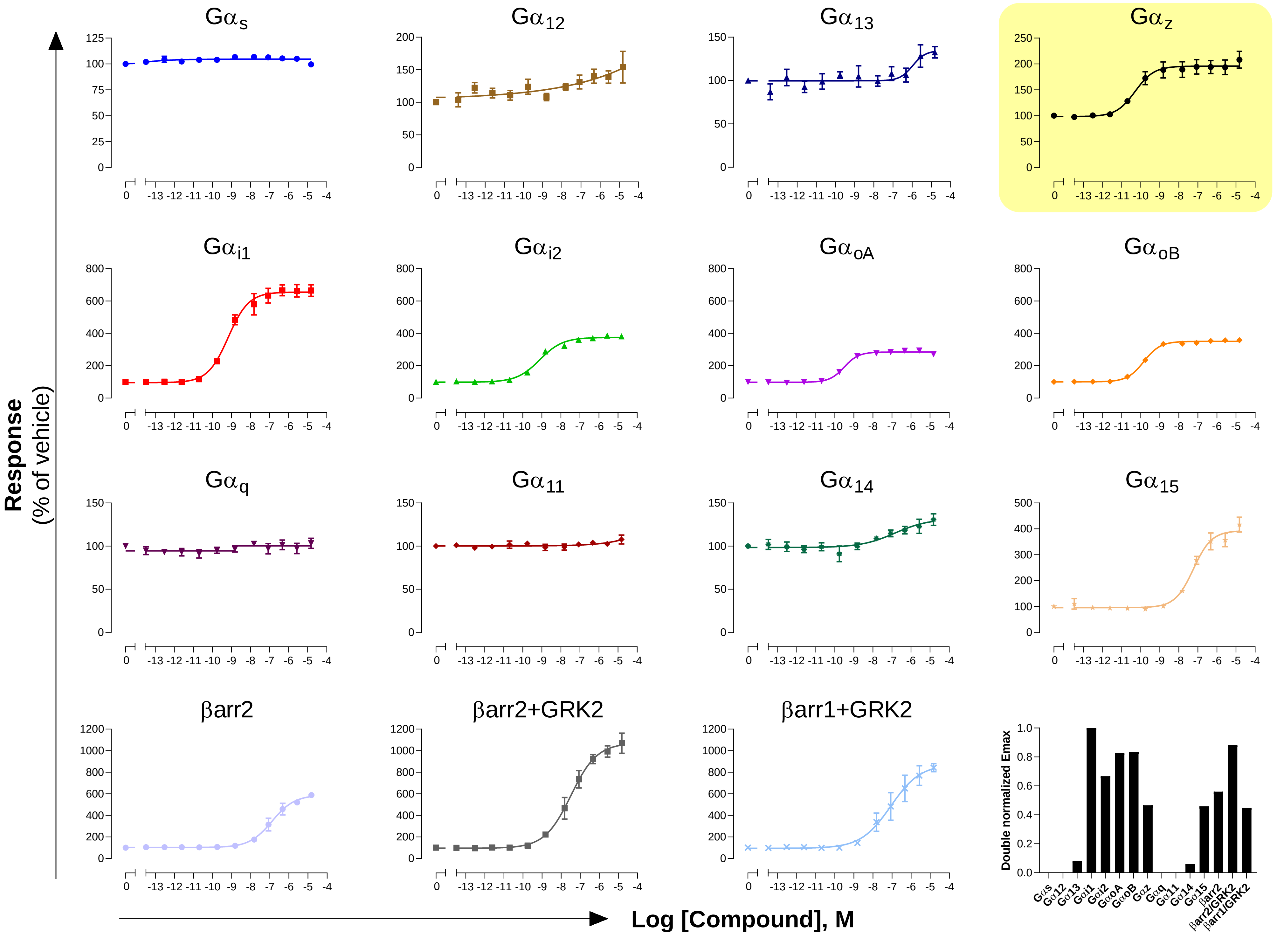

**GPCR: $EP_1$**

**Ligand: PGE2**

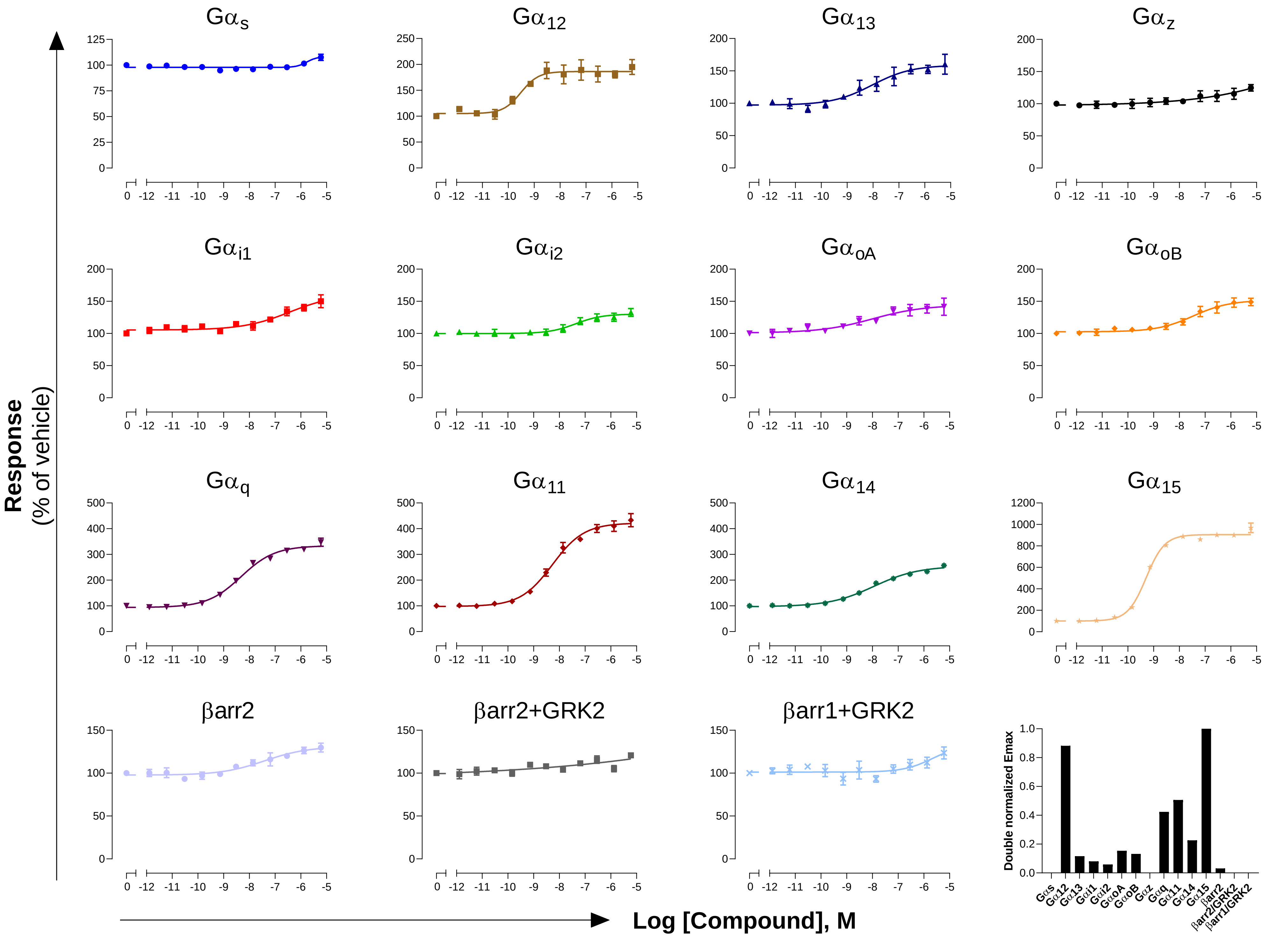

**GPCR: EP$_2$**

**Ligand: PGE2**

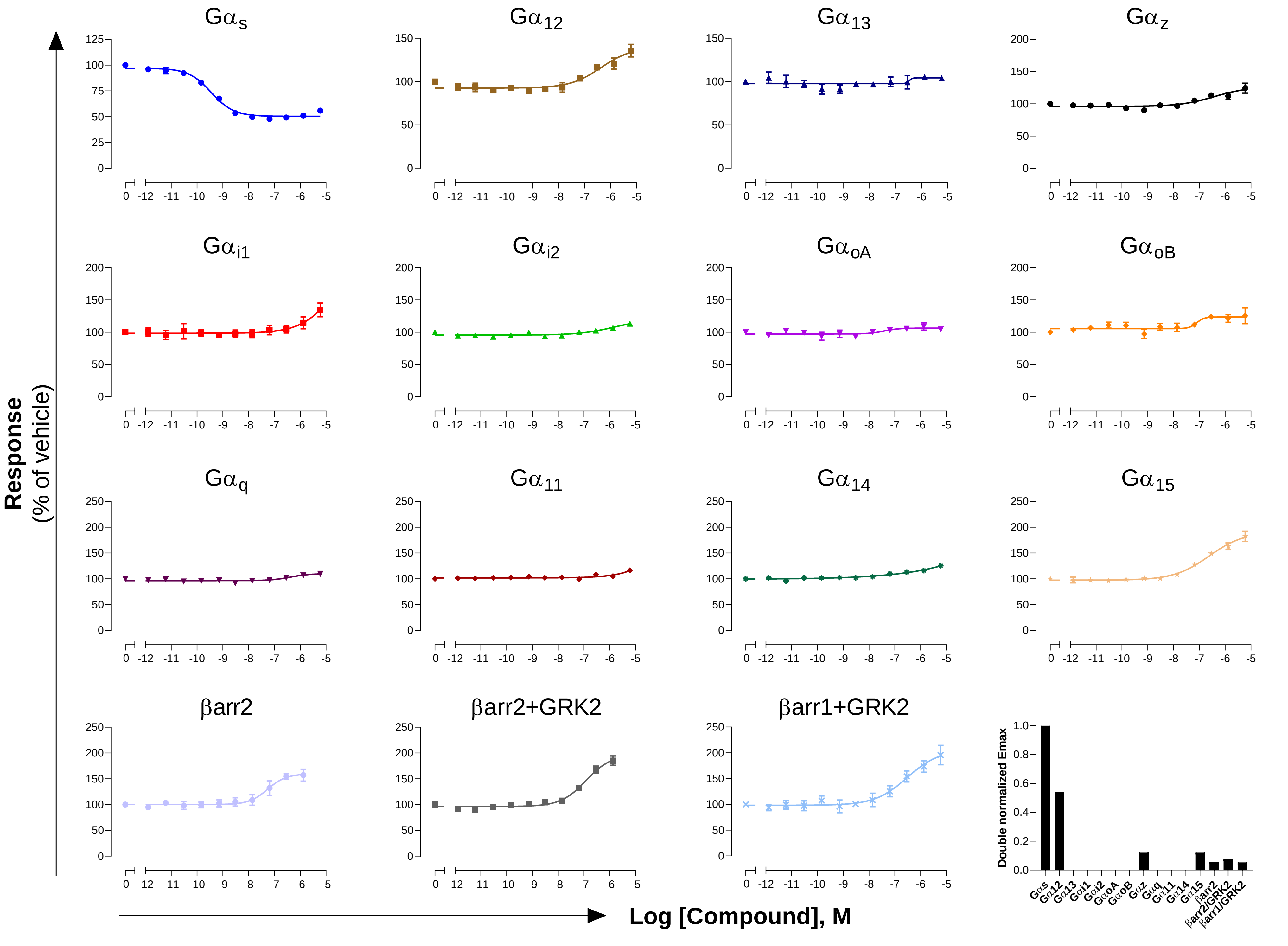

**GPCR: $EP_3$**

**Ligand: PGE2**

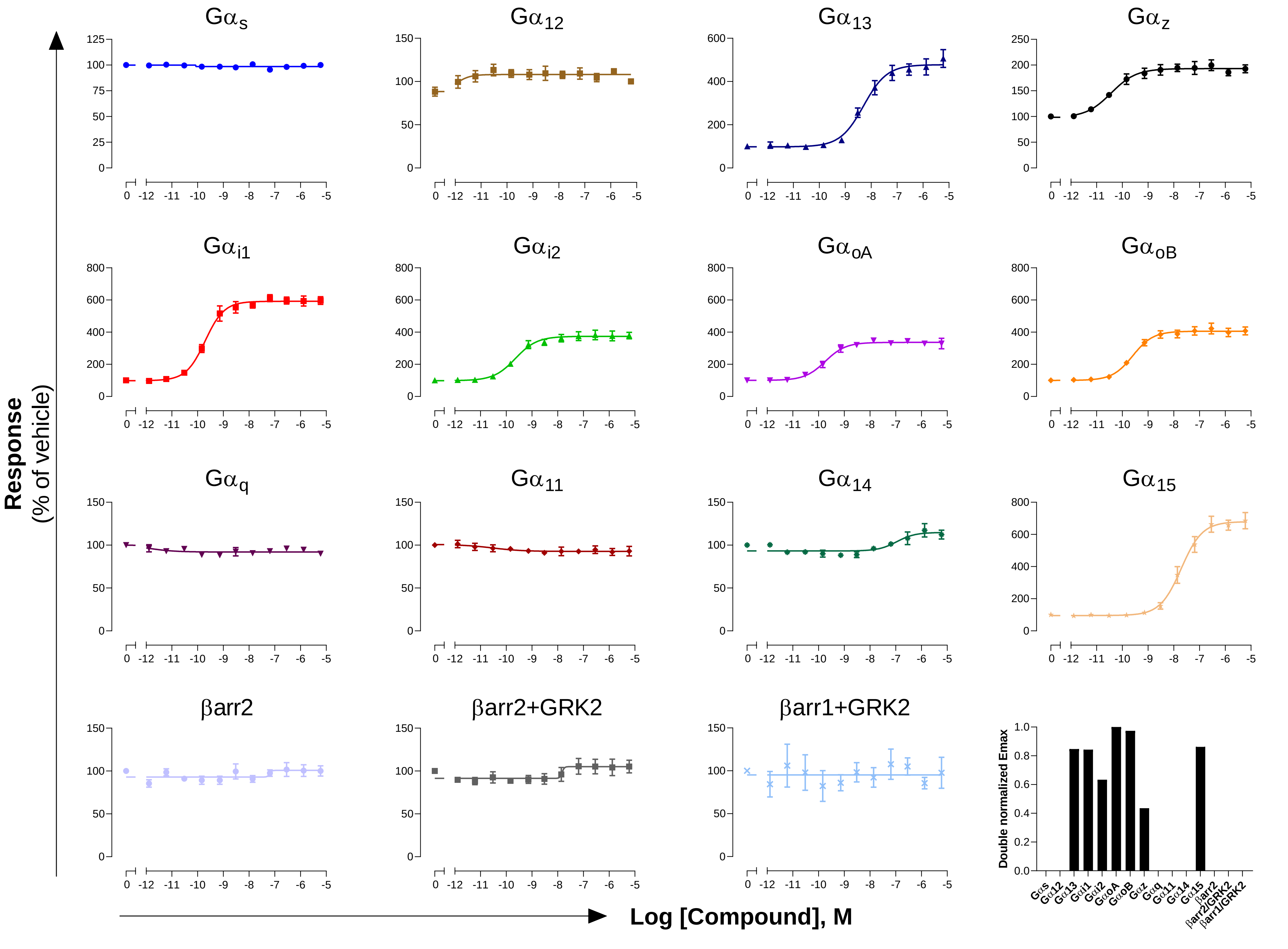

**GPCR: $EP_4$**

**Ligand: PGE2**

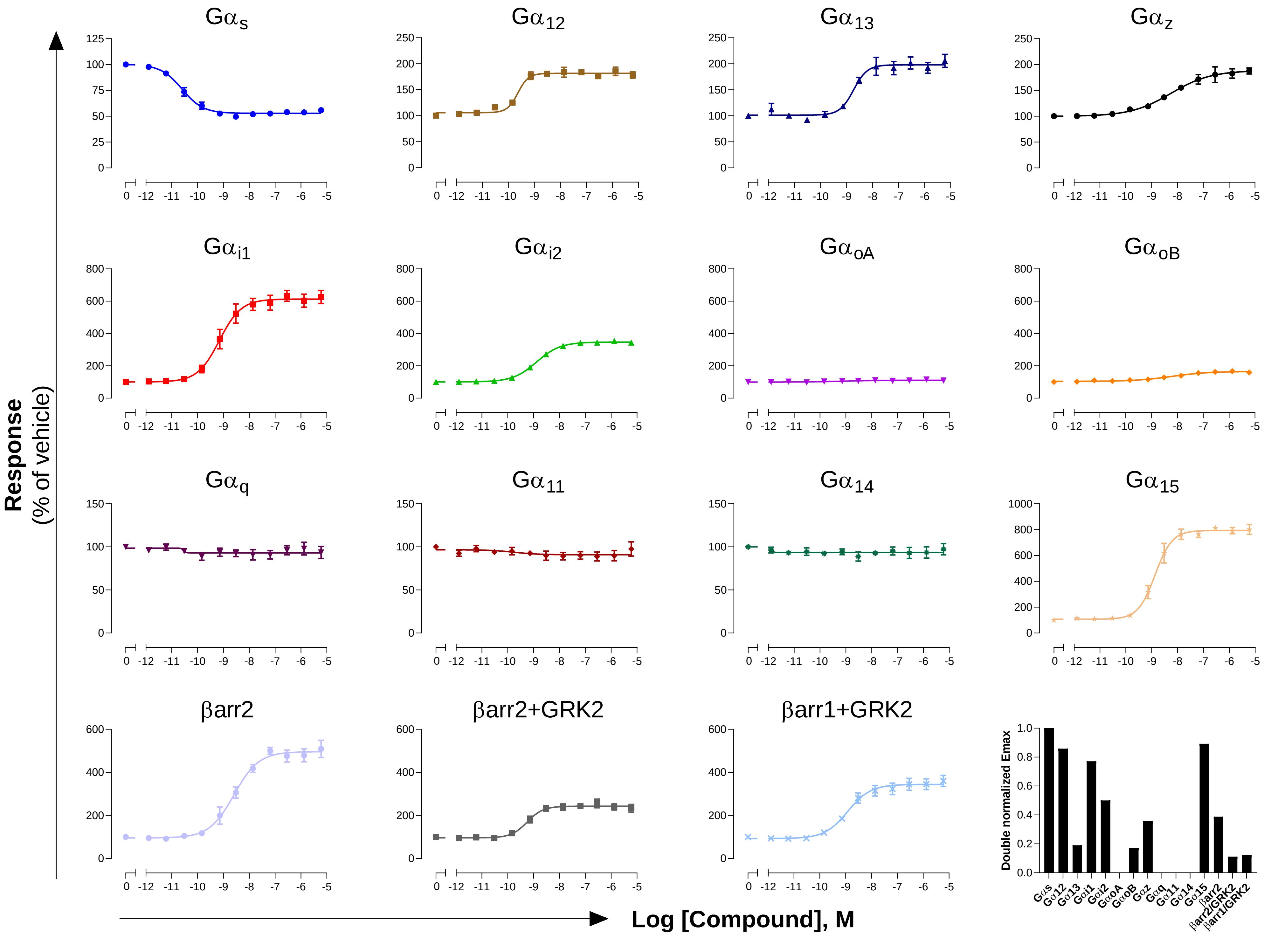

**GPCR: $ET_A$**

**Ligand: Endothelin-1**

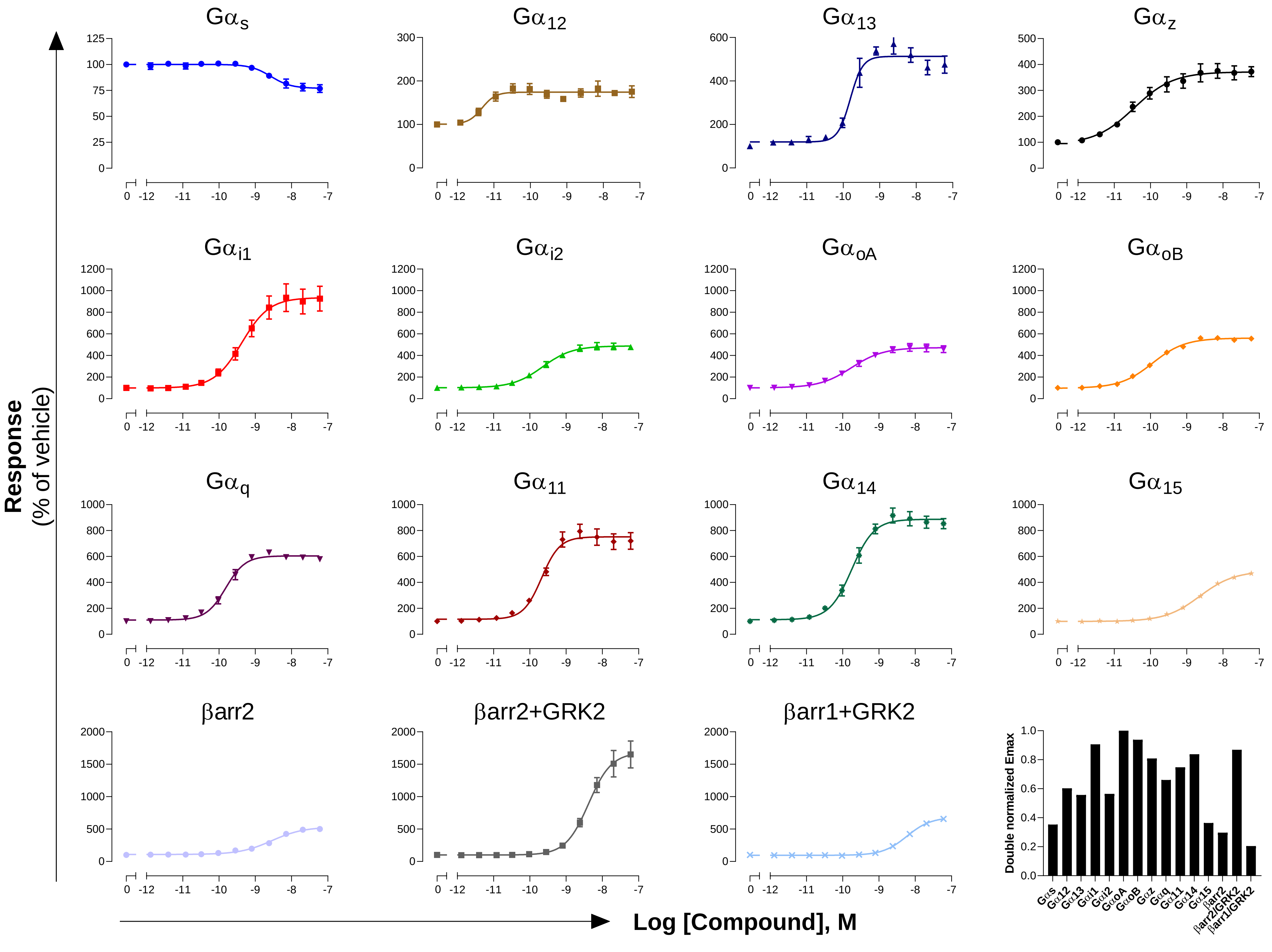

GPCR: FFA2
Ligand: Propionate

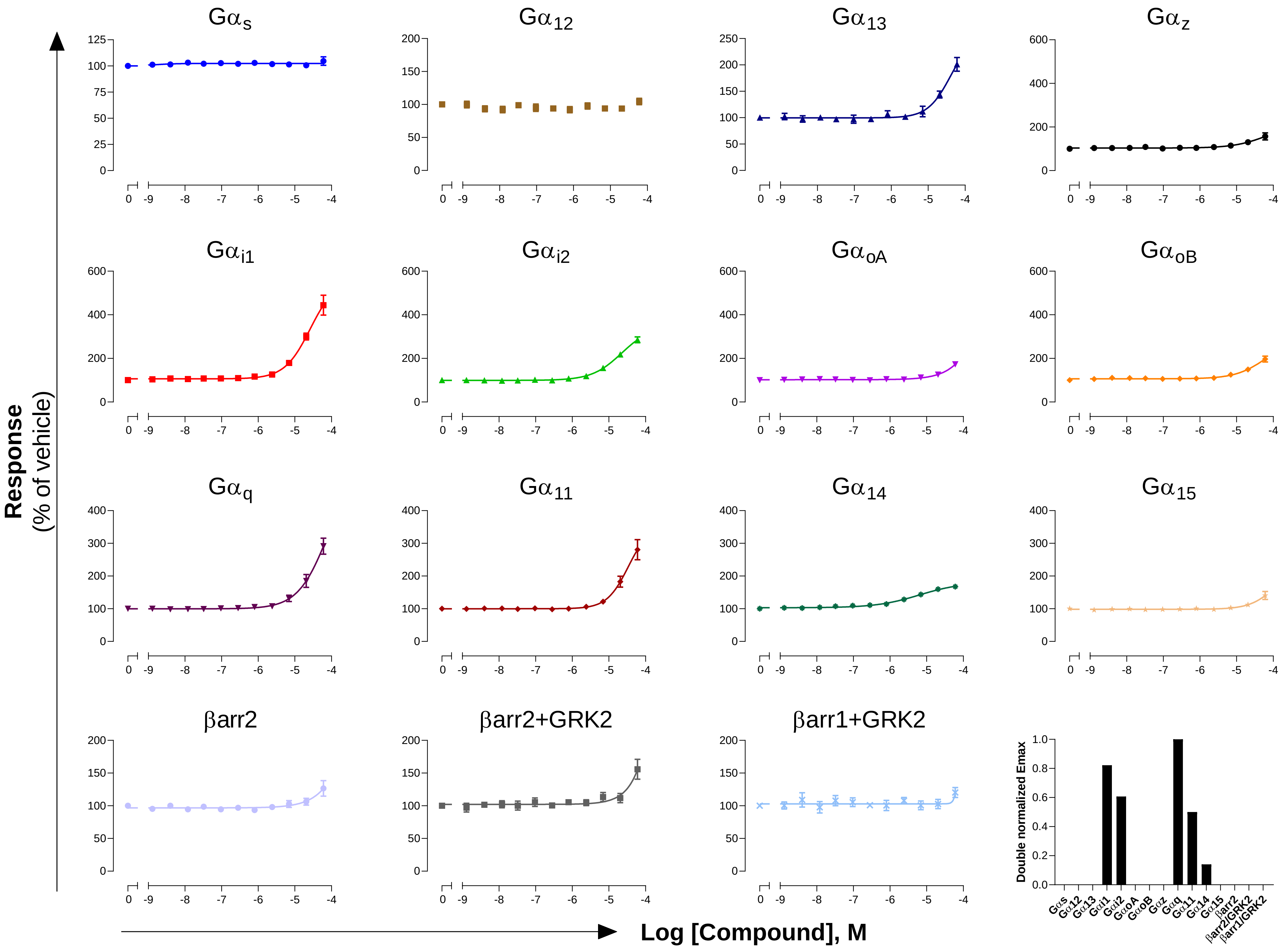

**GPCR: FFA3**
**Ligand: Propionate**

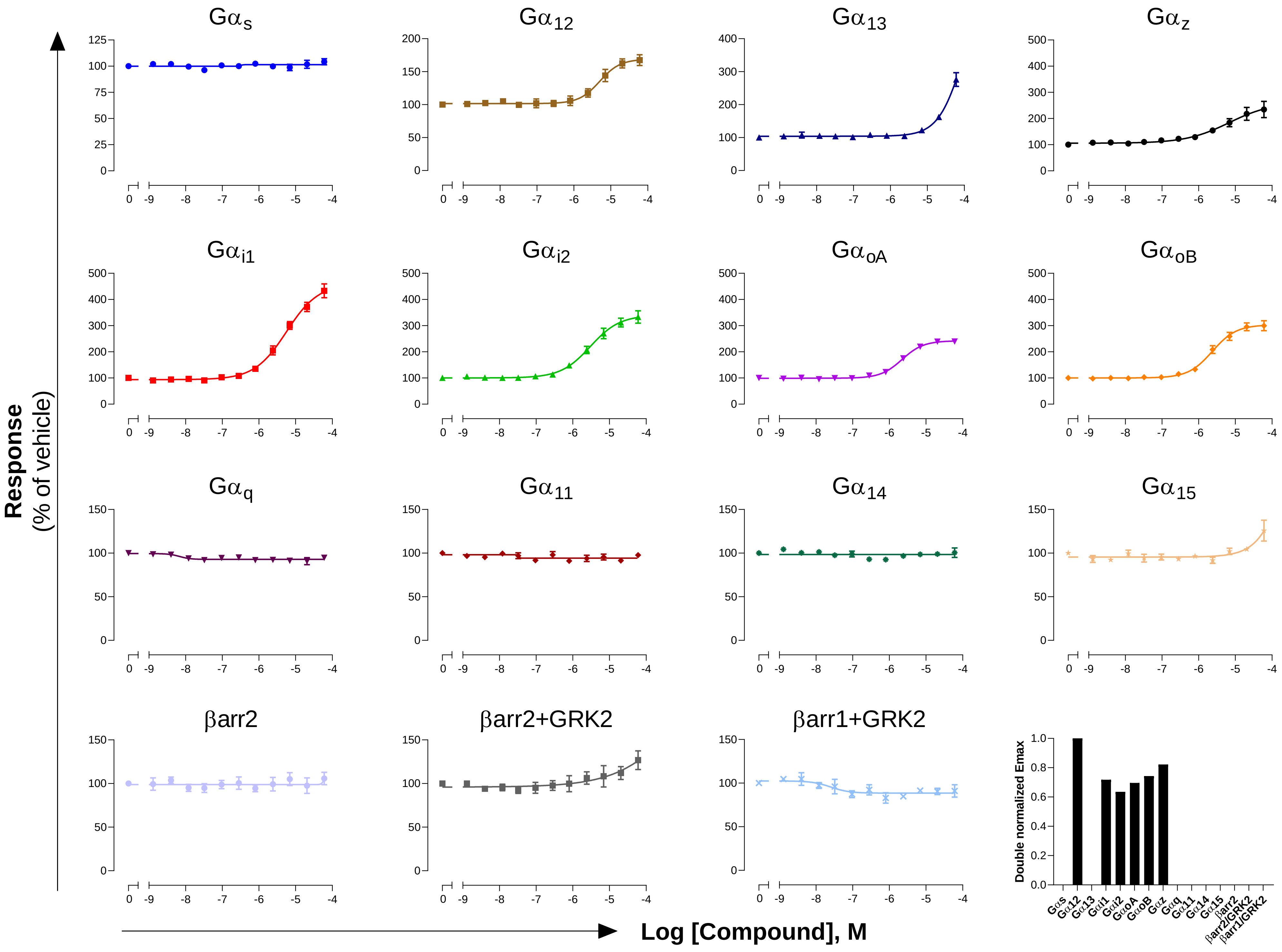

**GPCR: FFA4**
**Ligand: α-linolenic acid**

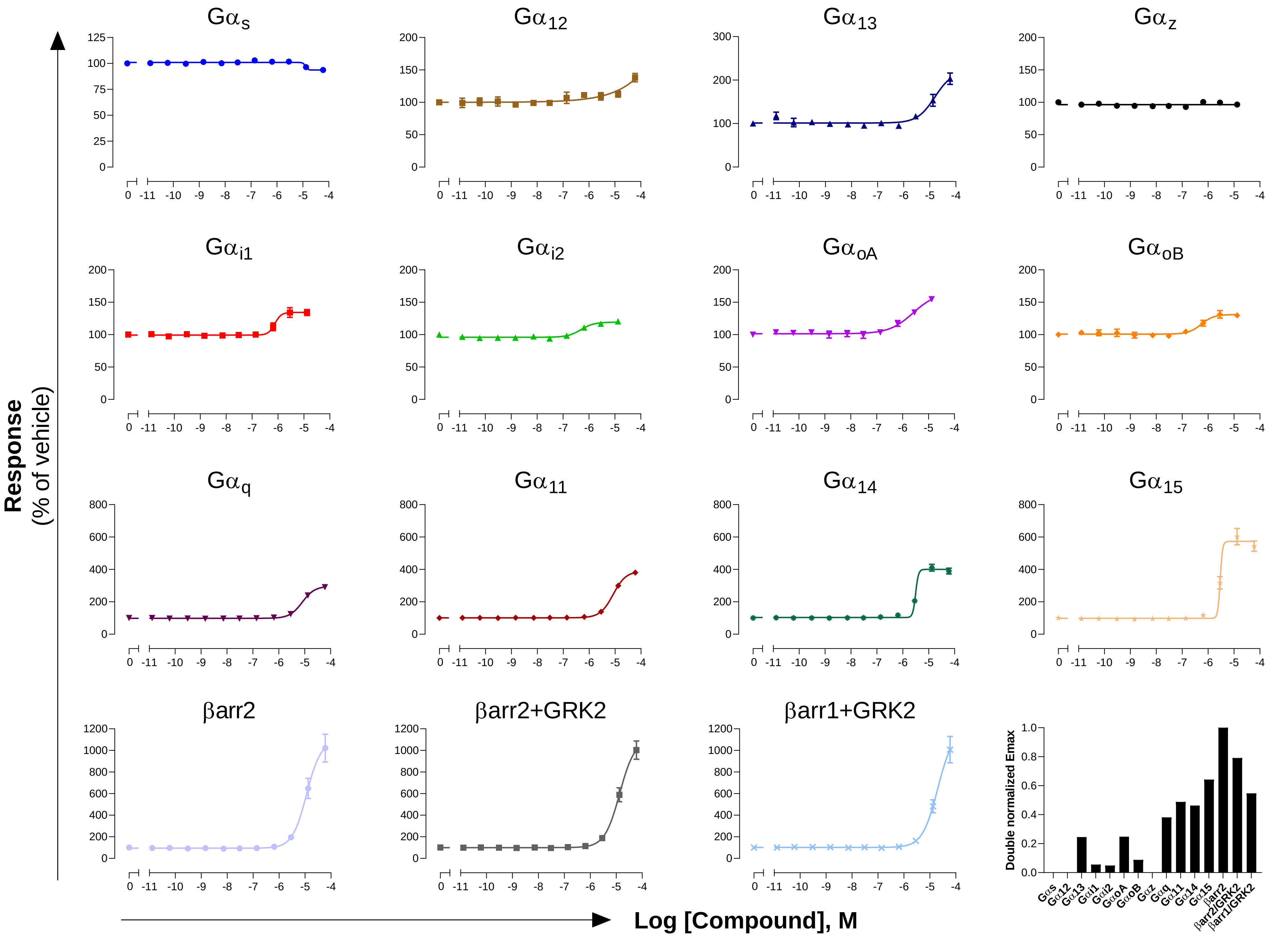

**GPCR: FP**
**Ligand: PGD2**

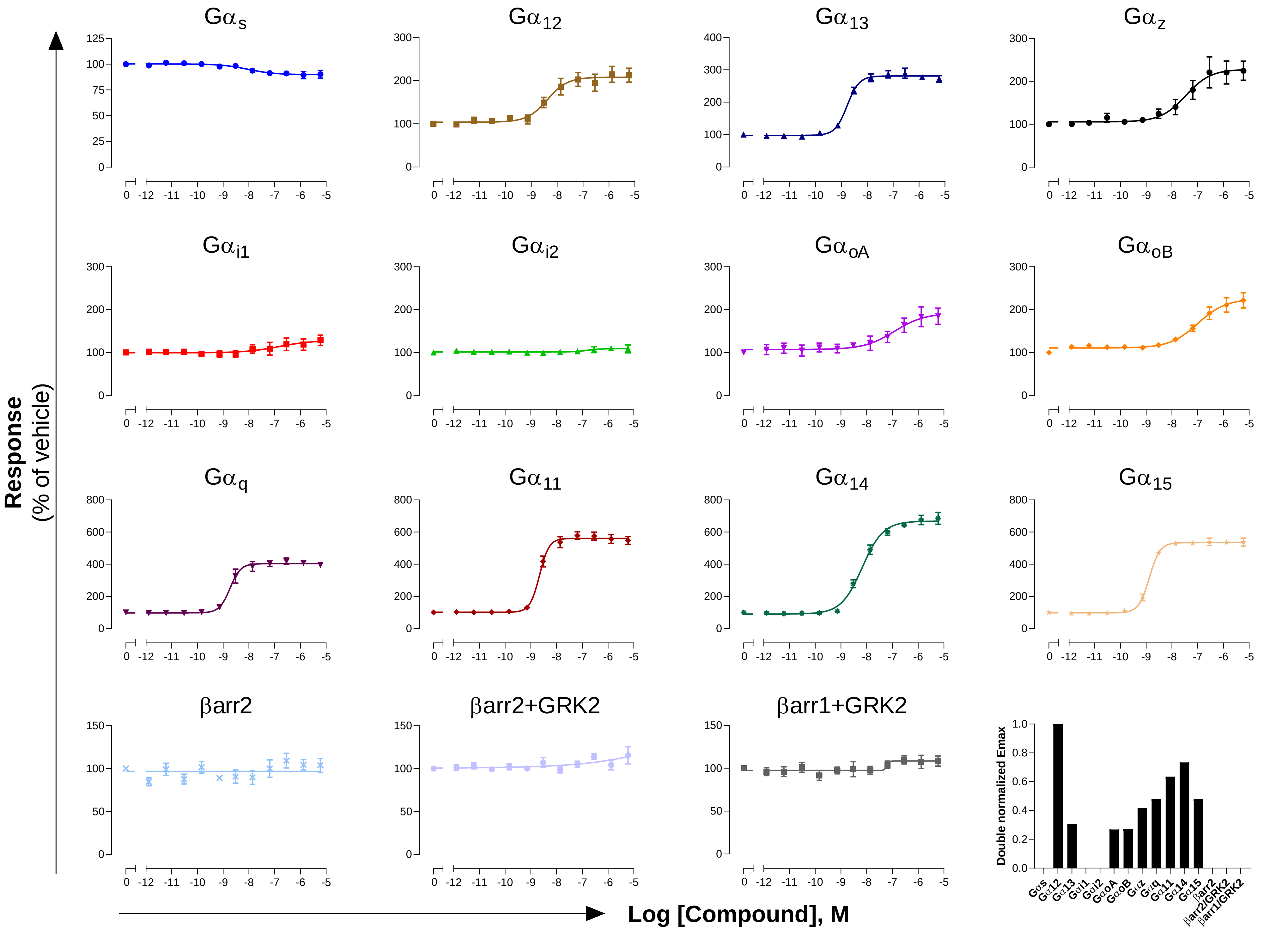

**GPCR: GCGR**
**Ligand: Glucagon**

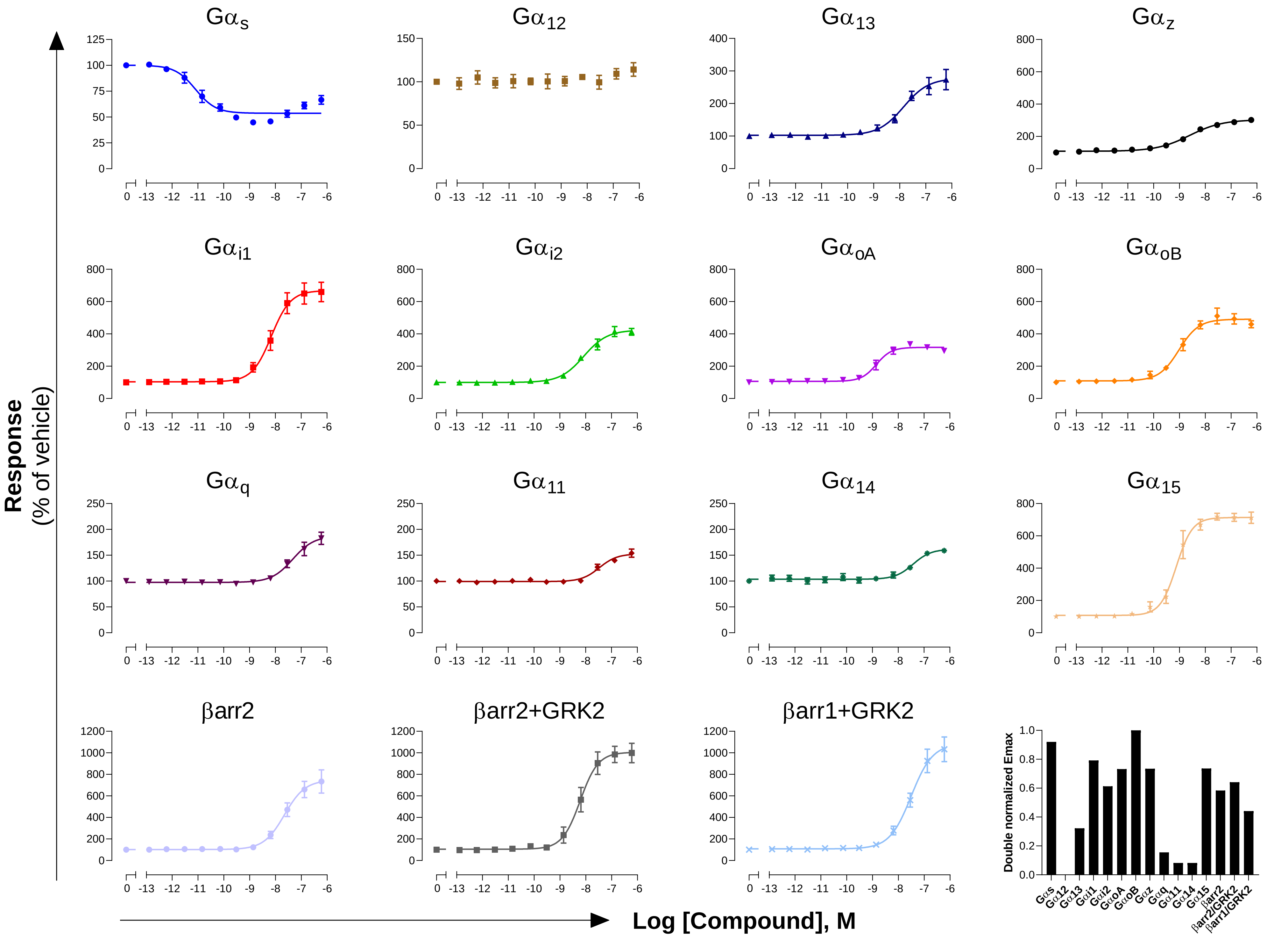

**GPCR: GHSR**
**Ligand: Ghrelin**

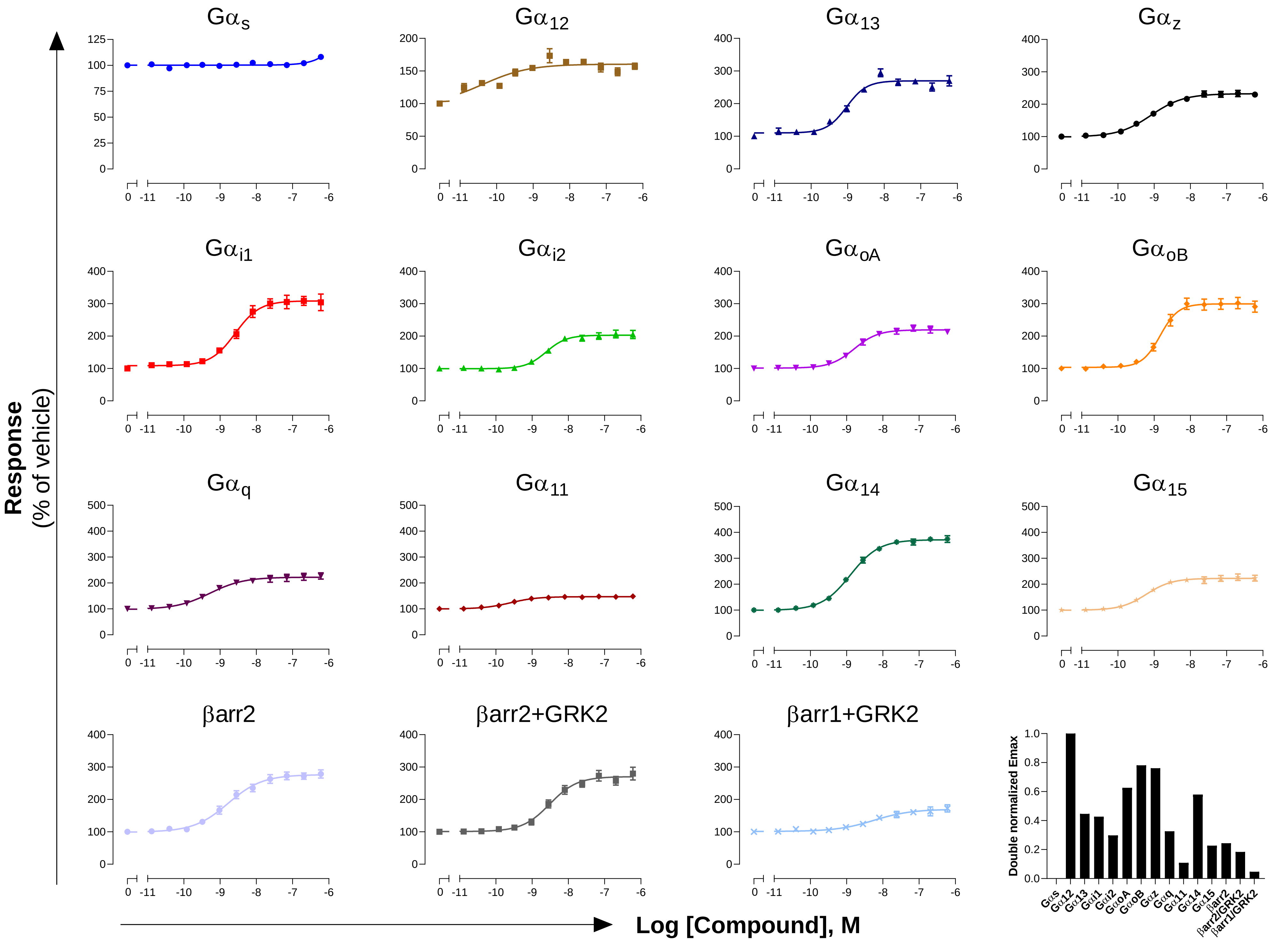

### GPCR: GIP
### Ligand: GIP

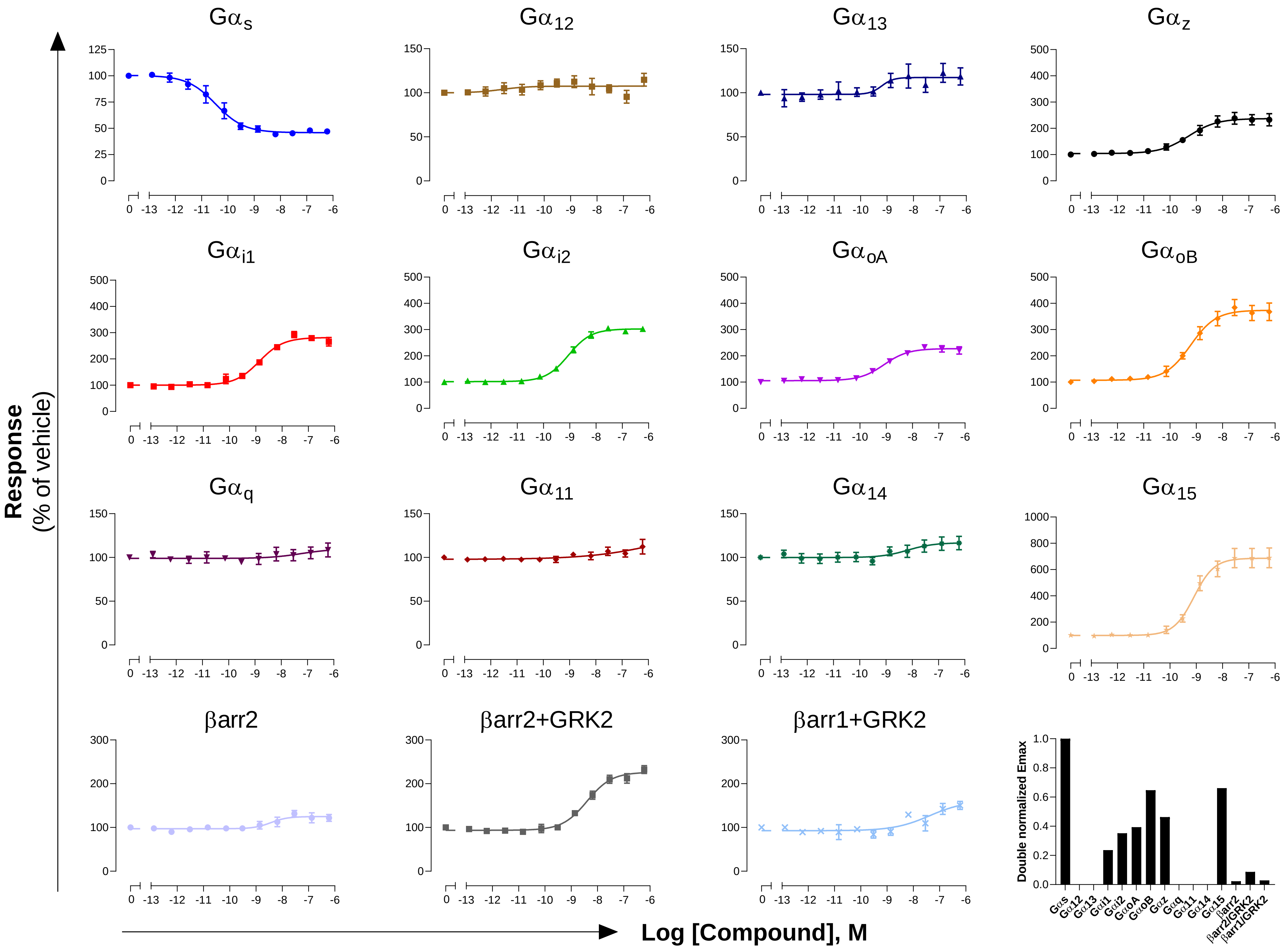

GPCR: GLP-1
Ligand: GLP-1 (7-36)

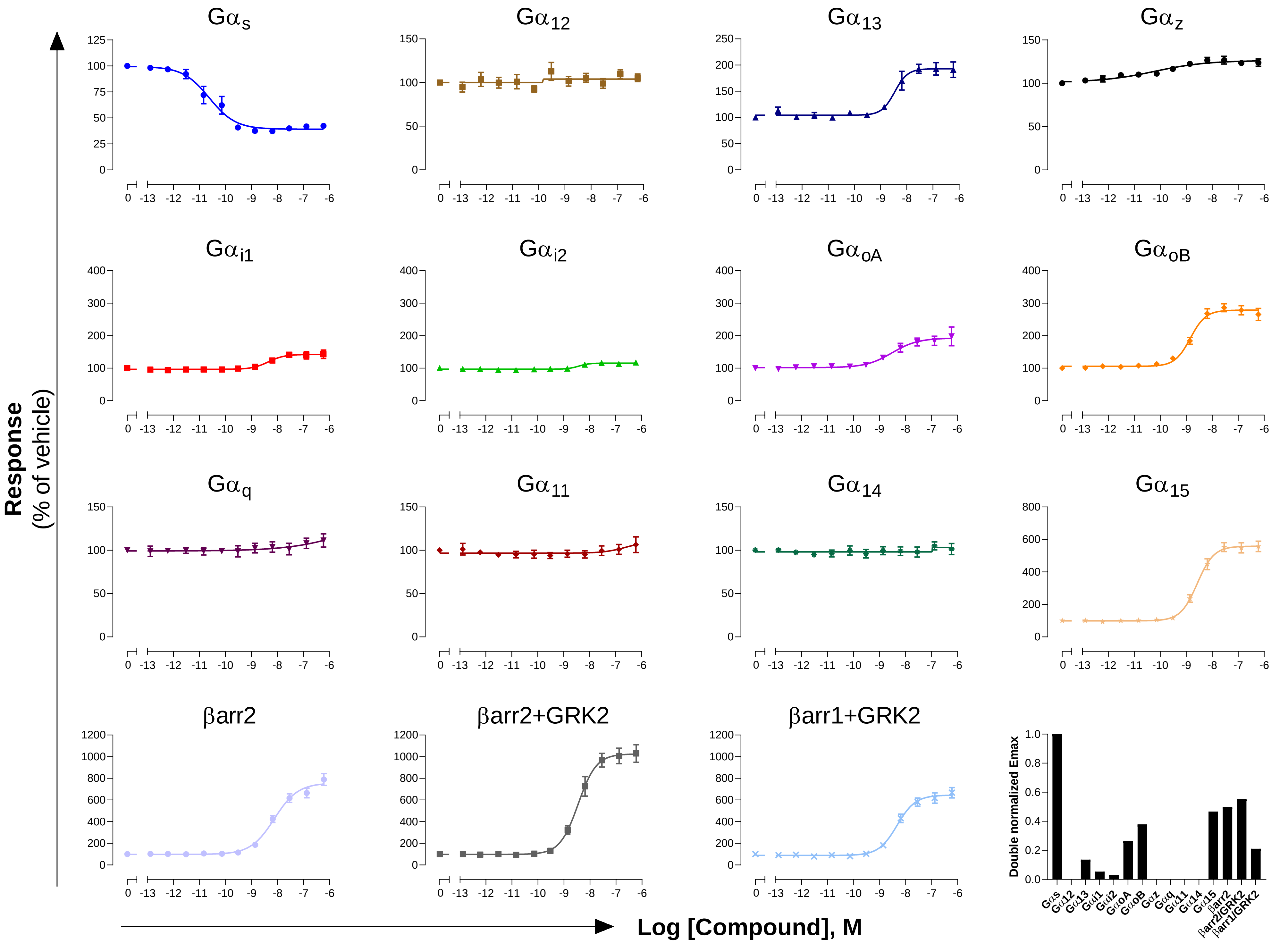

**GPCR: GLP-2**
**Ligand: GLP-2 (1-33)**

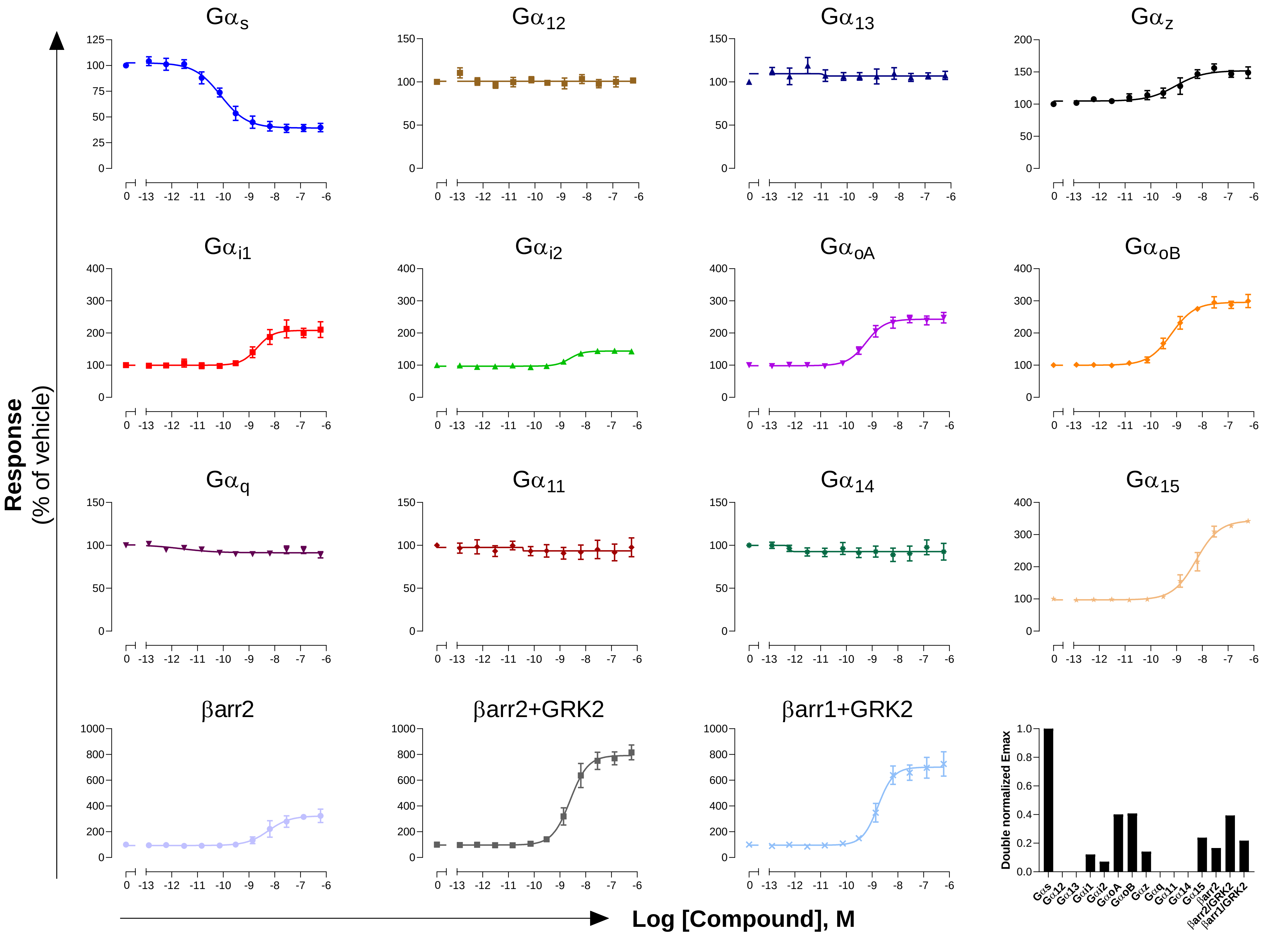

### GPCR: GnRHR
### Ligand: GnRH

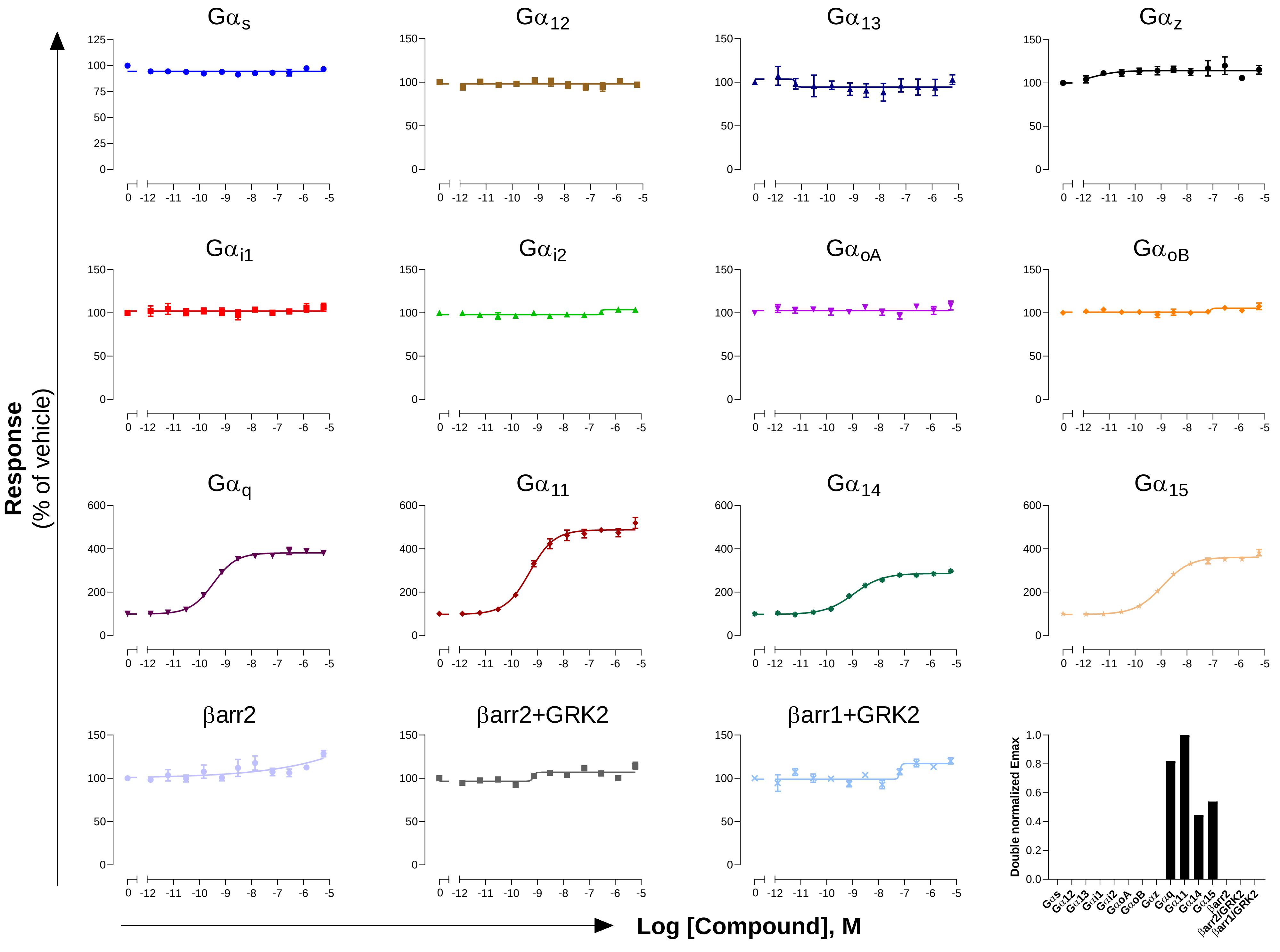

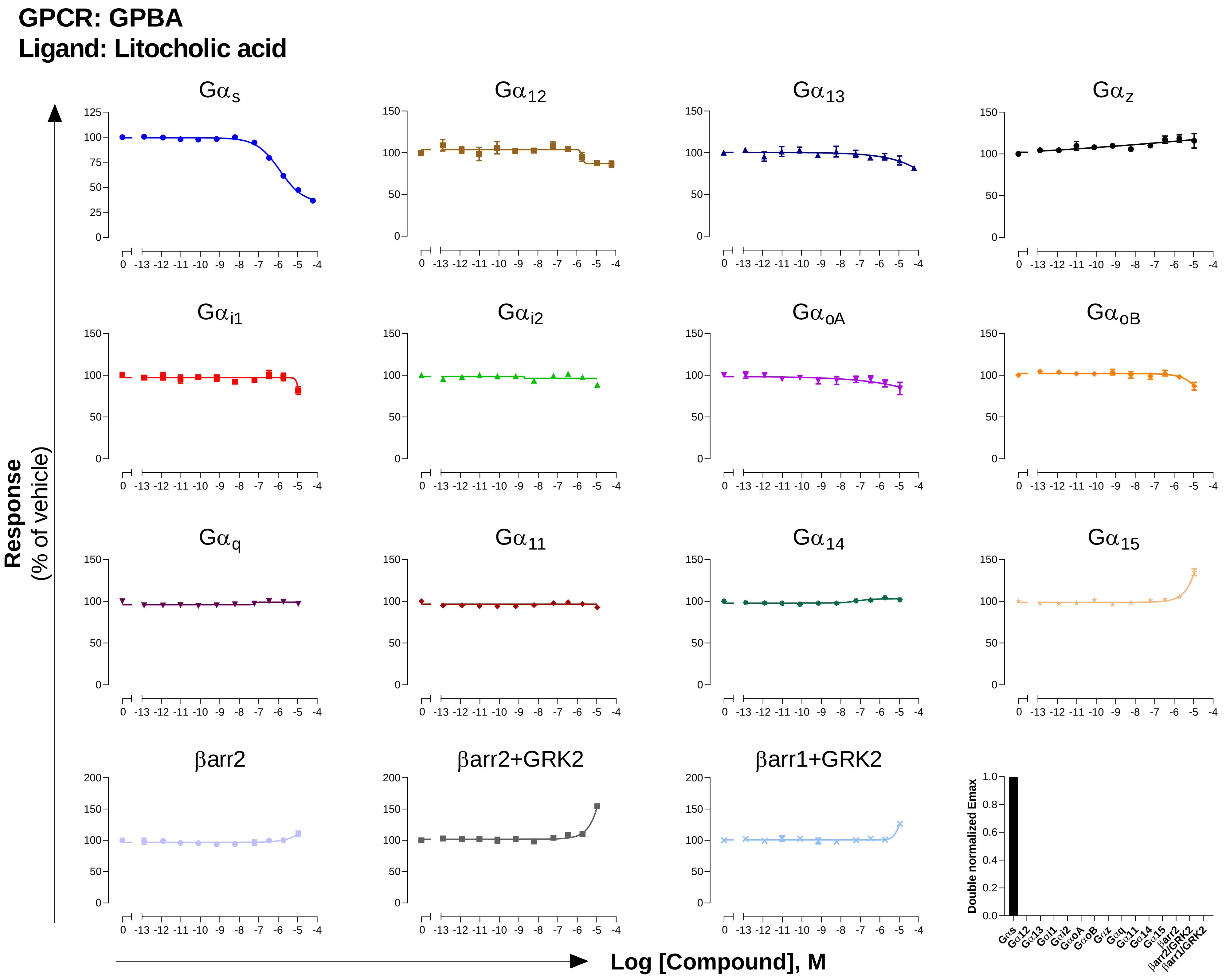

GPCR: GPBA
Ligand: Litocholic acid
Gαs
Gα12
Gα13
Gαz
Gαi1
Gαi2
GαoA
GαoB
Gαq
Gα11
Gα14
Gα15
βarr2
βarr2+GRK2
βarr1+GRK2
Response
(% of vehicle)
Log [Compound], M
Double normalized Emax

GPCR: GPR39
Ligand: $Zn^{2+}$

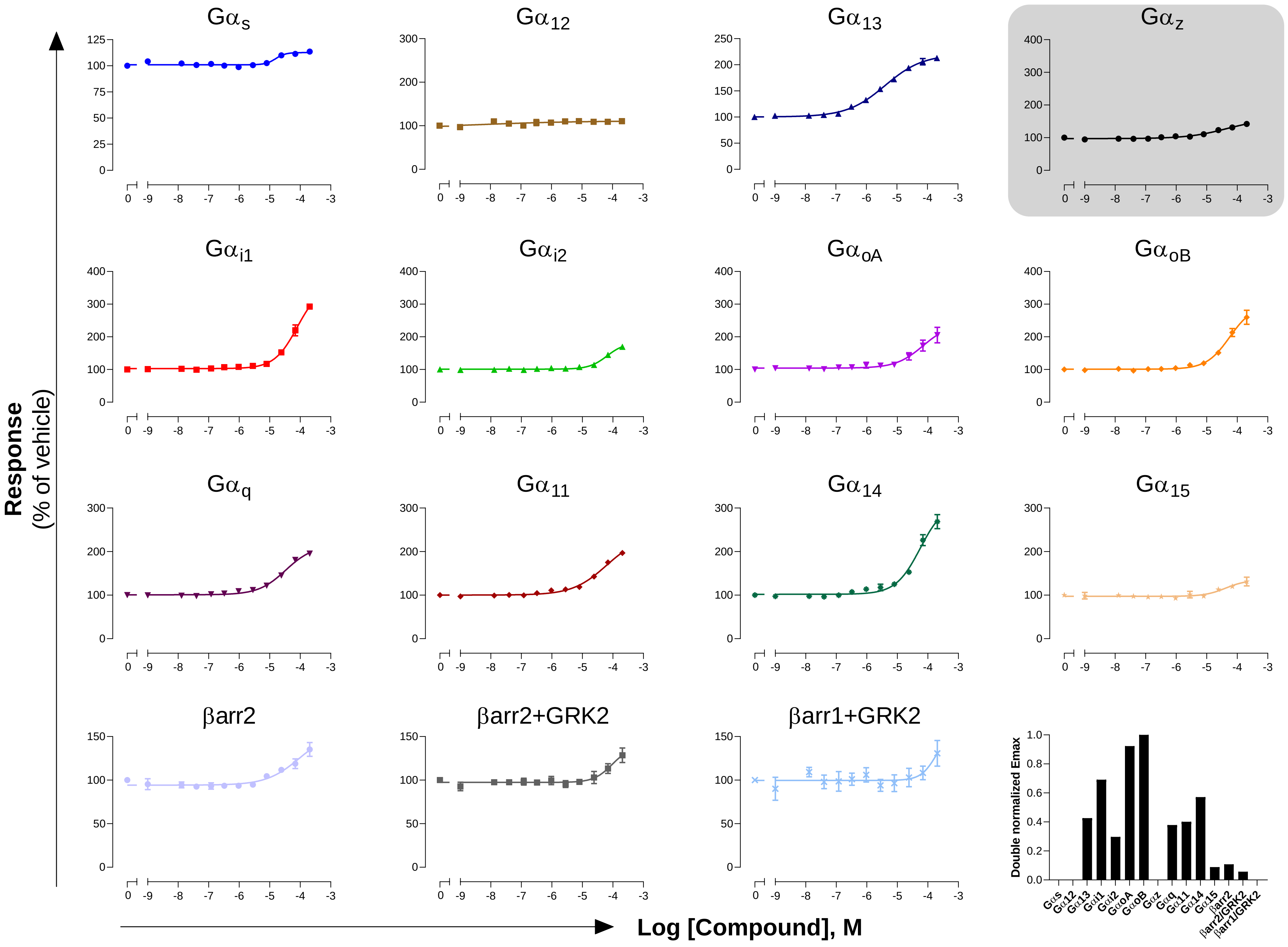

### GPCR: GPR4
### Ligand: Protons (pH)

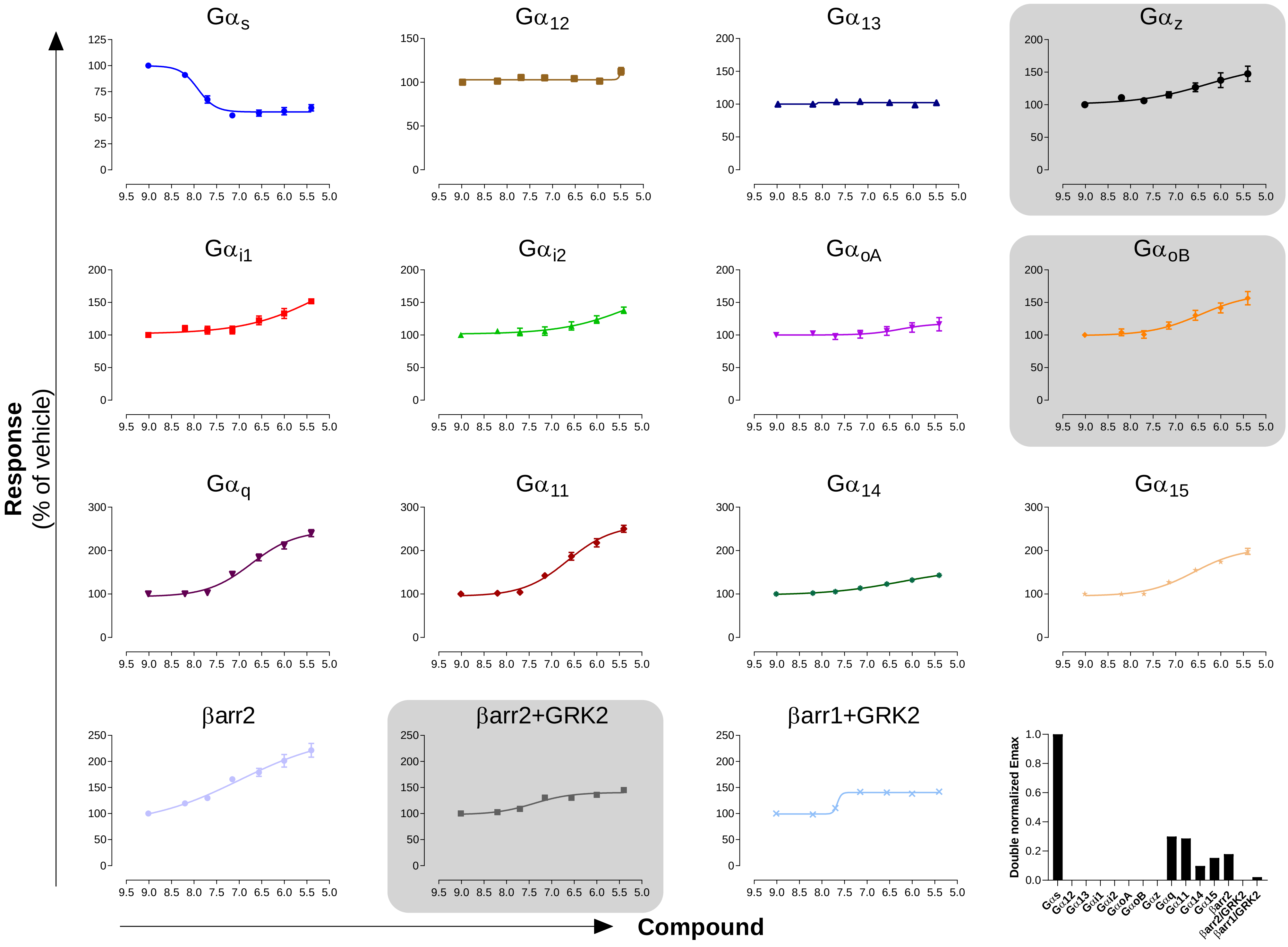

### GPCR: GPR65
### Ligand: protons (pH)

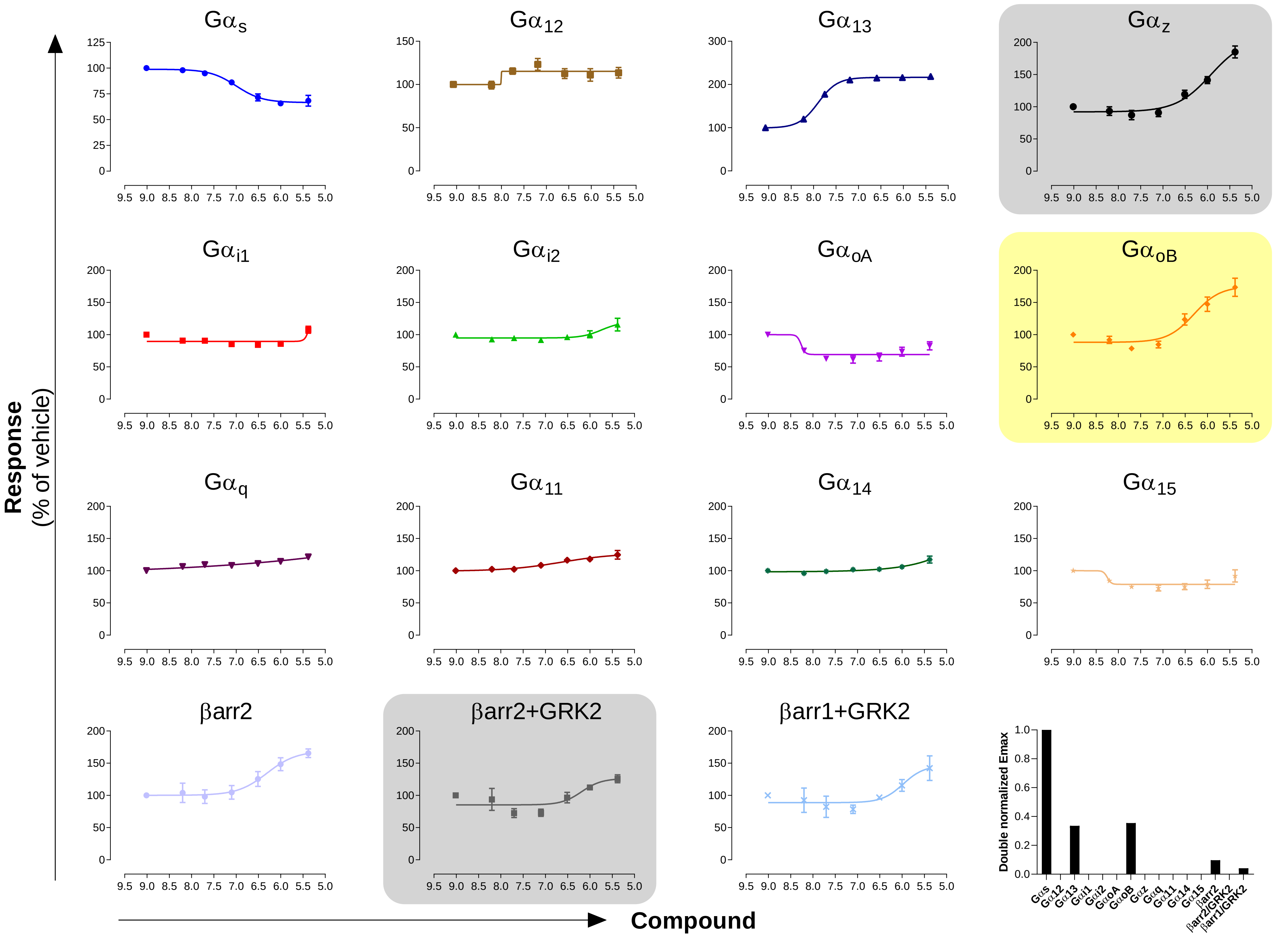

### GPCR: GPR68
### Ligand: Protons (pH)

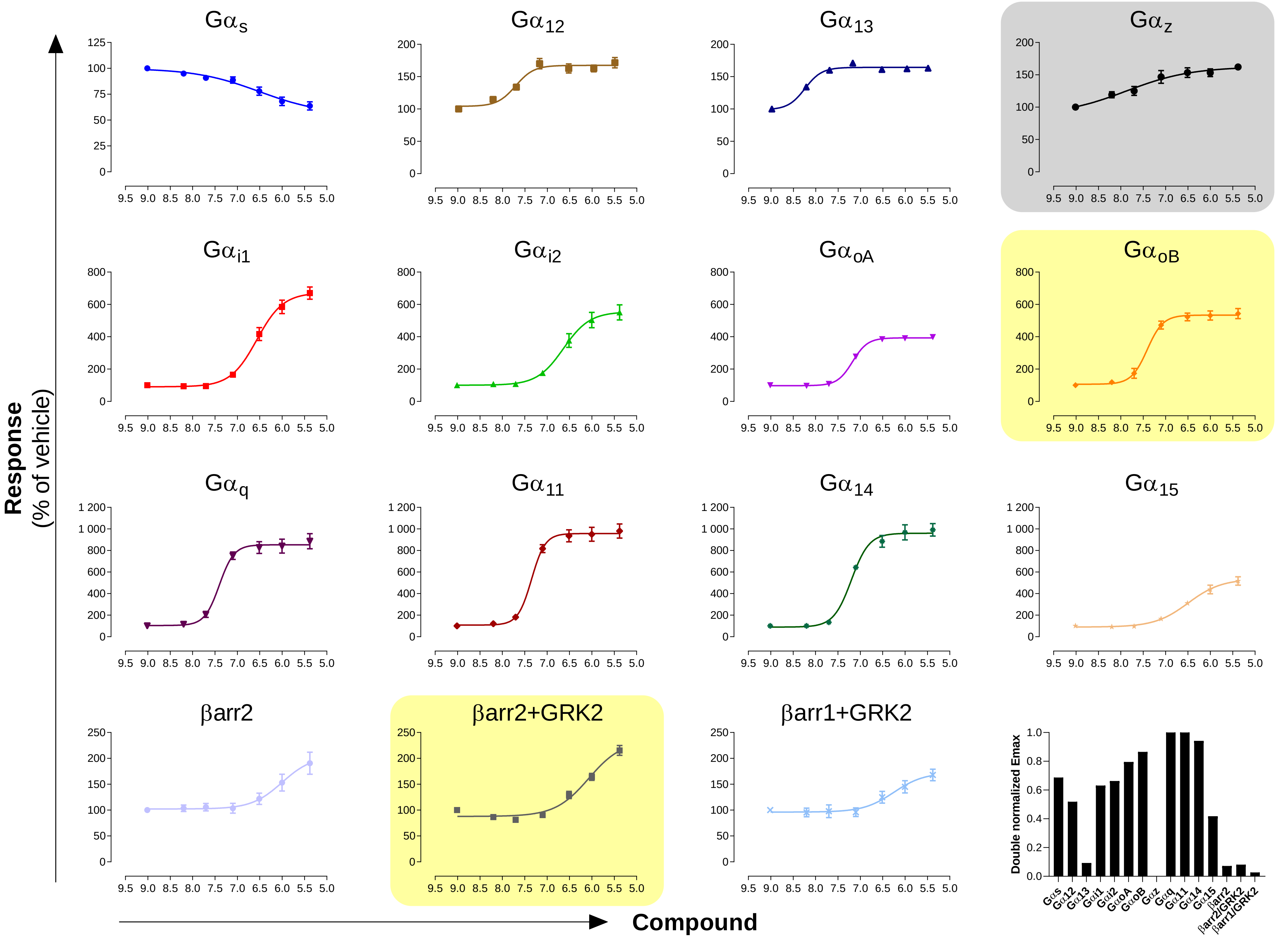

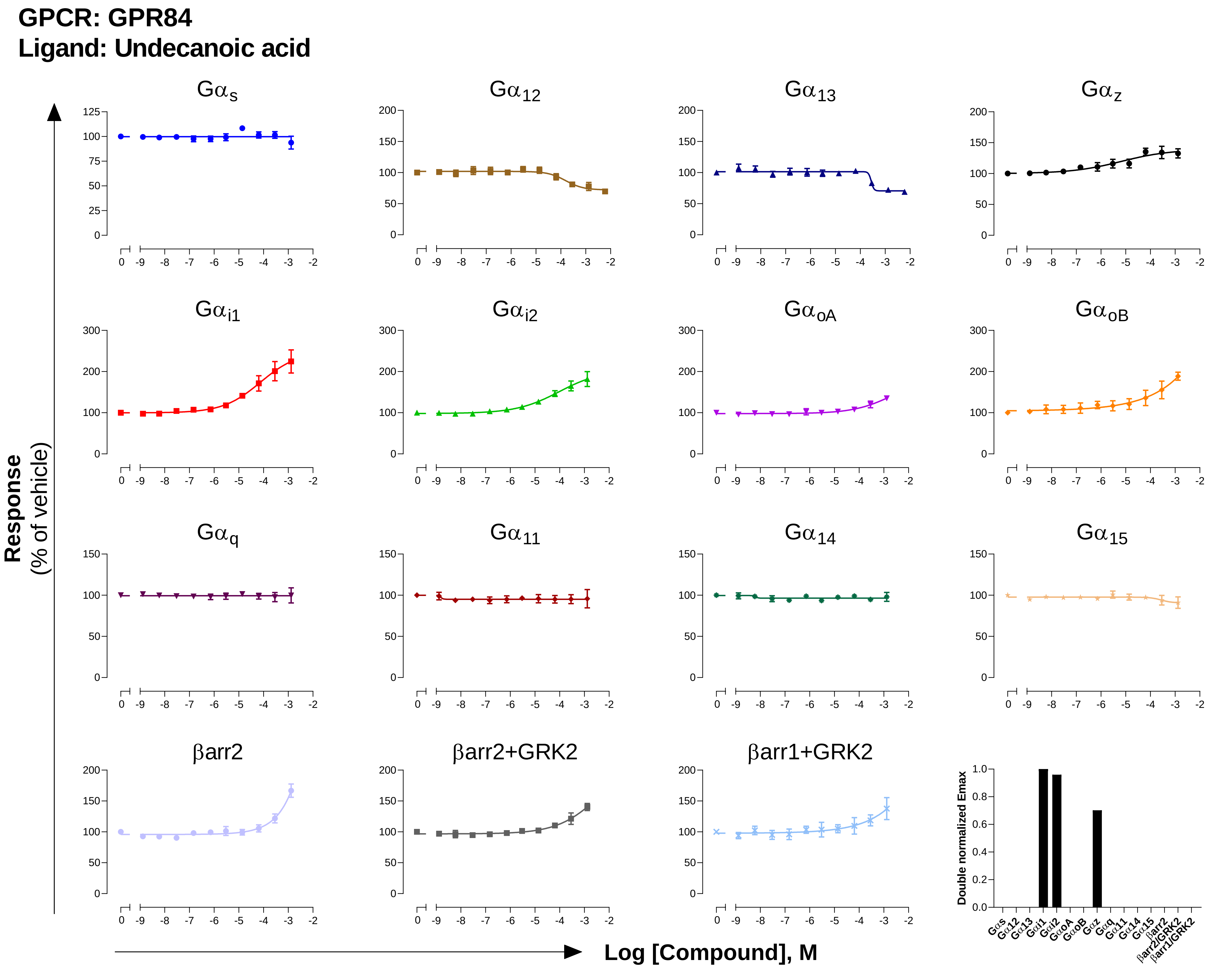
GPCR: GPR84
Ligand: Undecanoic acid
Gαs
Gα12
Gα13
Gαz
Gαi1
Gαi2
GαoA
GαoB
Gαq
Gα11
Gα14
Gα15
βarr2
βarr2+GRK2
βarr1+GRK2
Response
(% of vehicle)
Log [Compound], M
Double normalized Emax

**GPCR: GPR183**
**Ligand: 7α-25 dihydroxycholesterol**

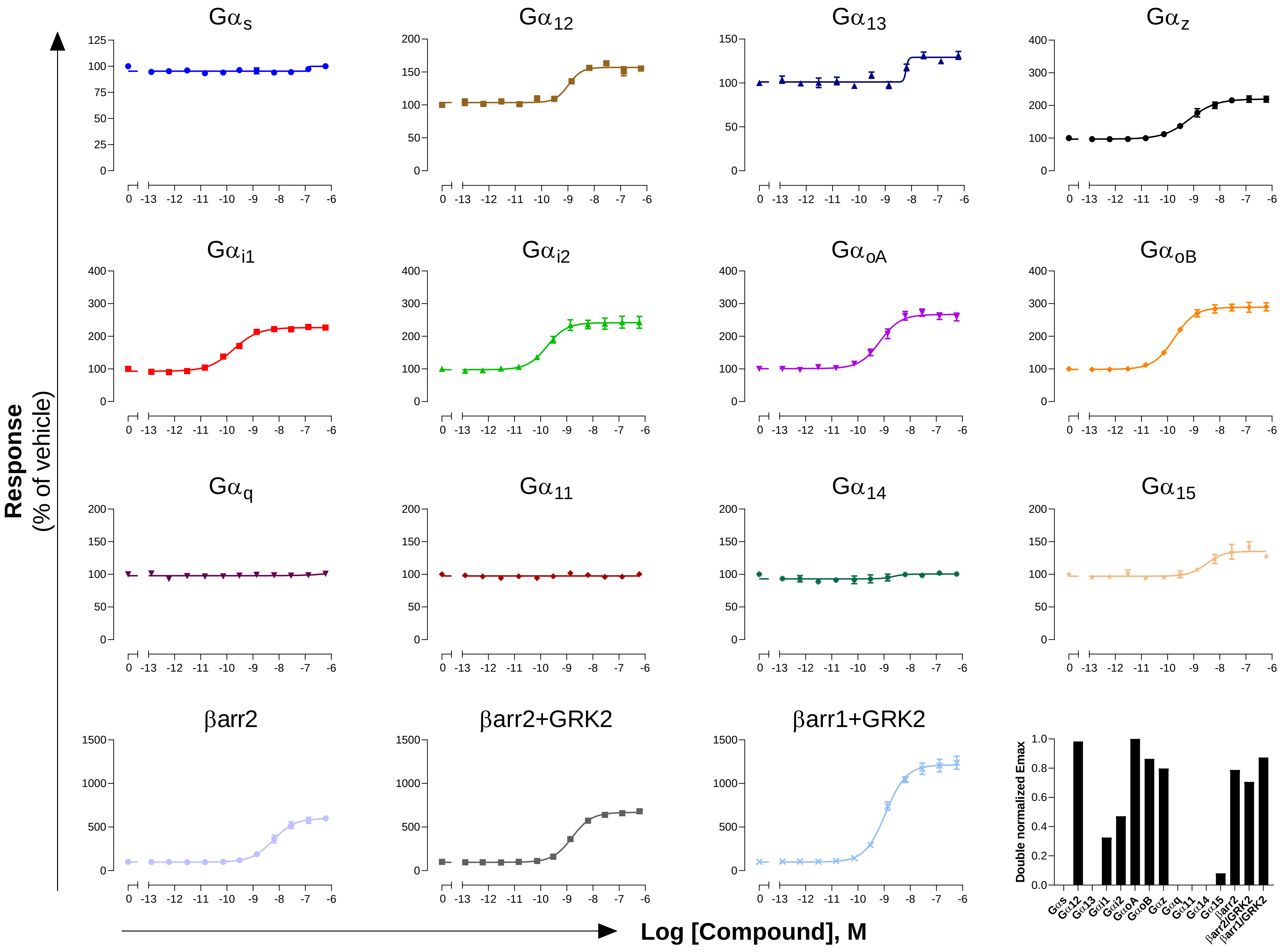

### GPCR: $H_1$
### Ligand: Histamine

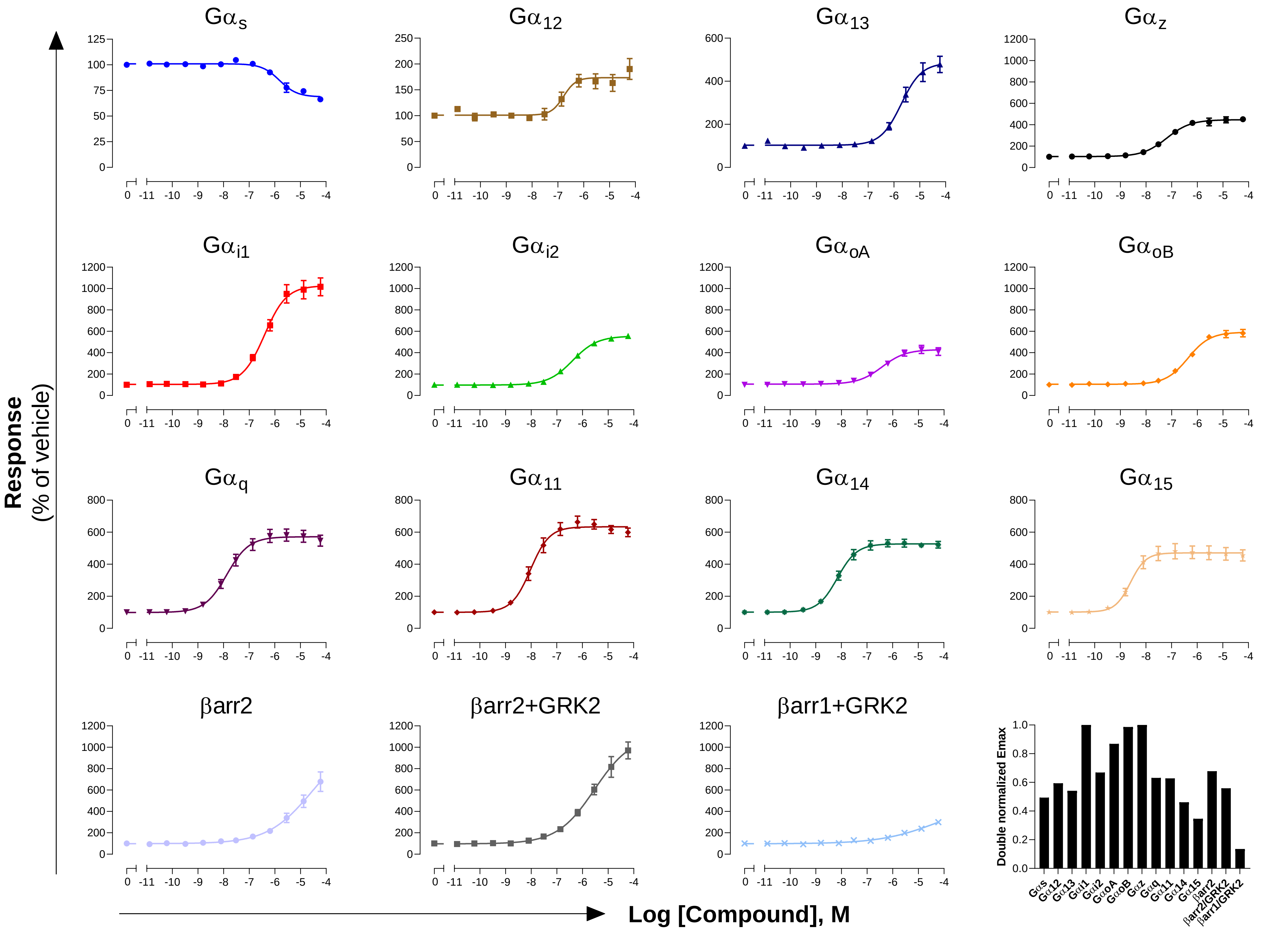

**GPCR: $H_2$**

**Ligand: Histamine**

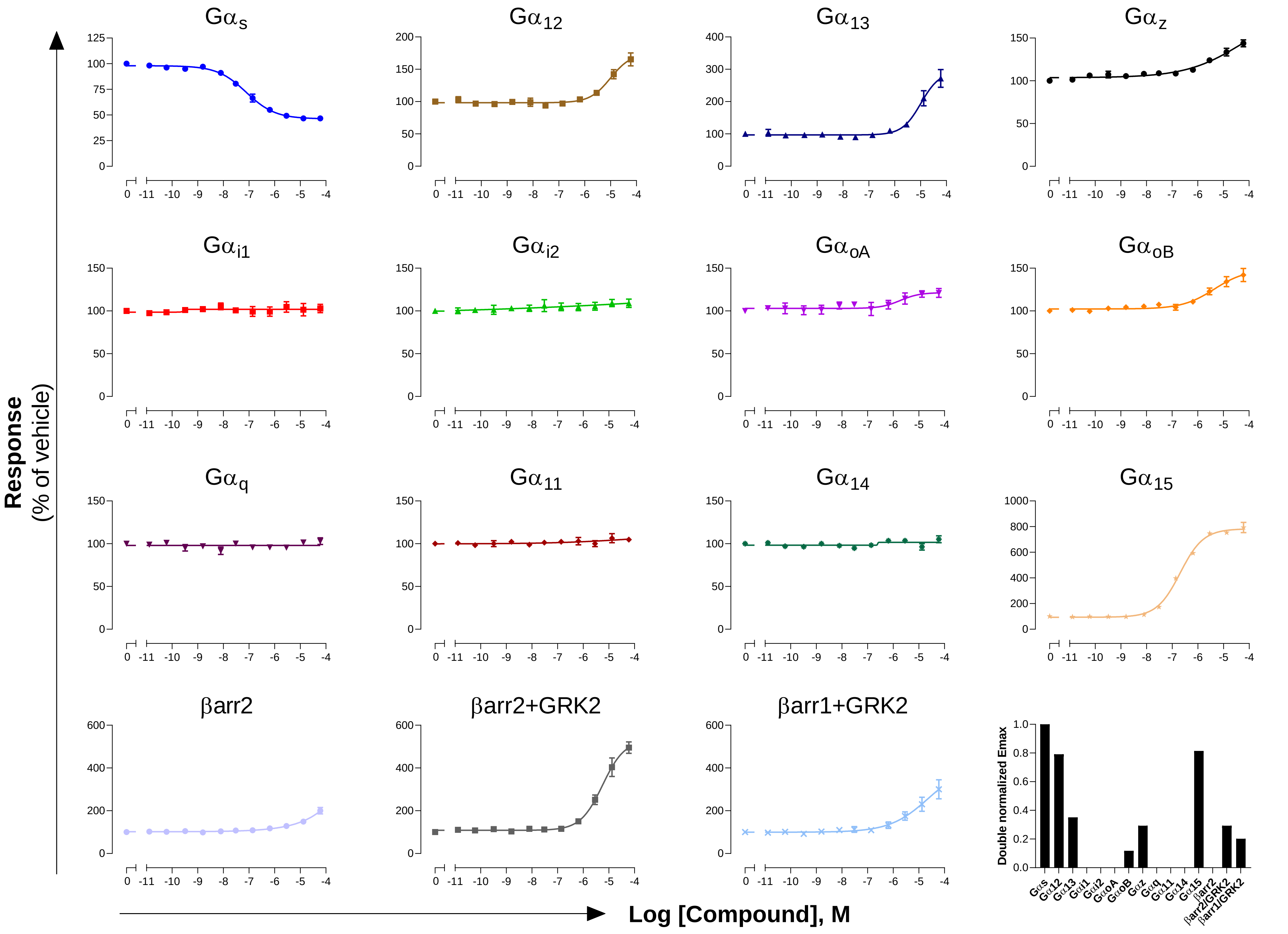

### GPCR: HCA$_2$
### Ligand: Nicotinic acid

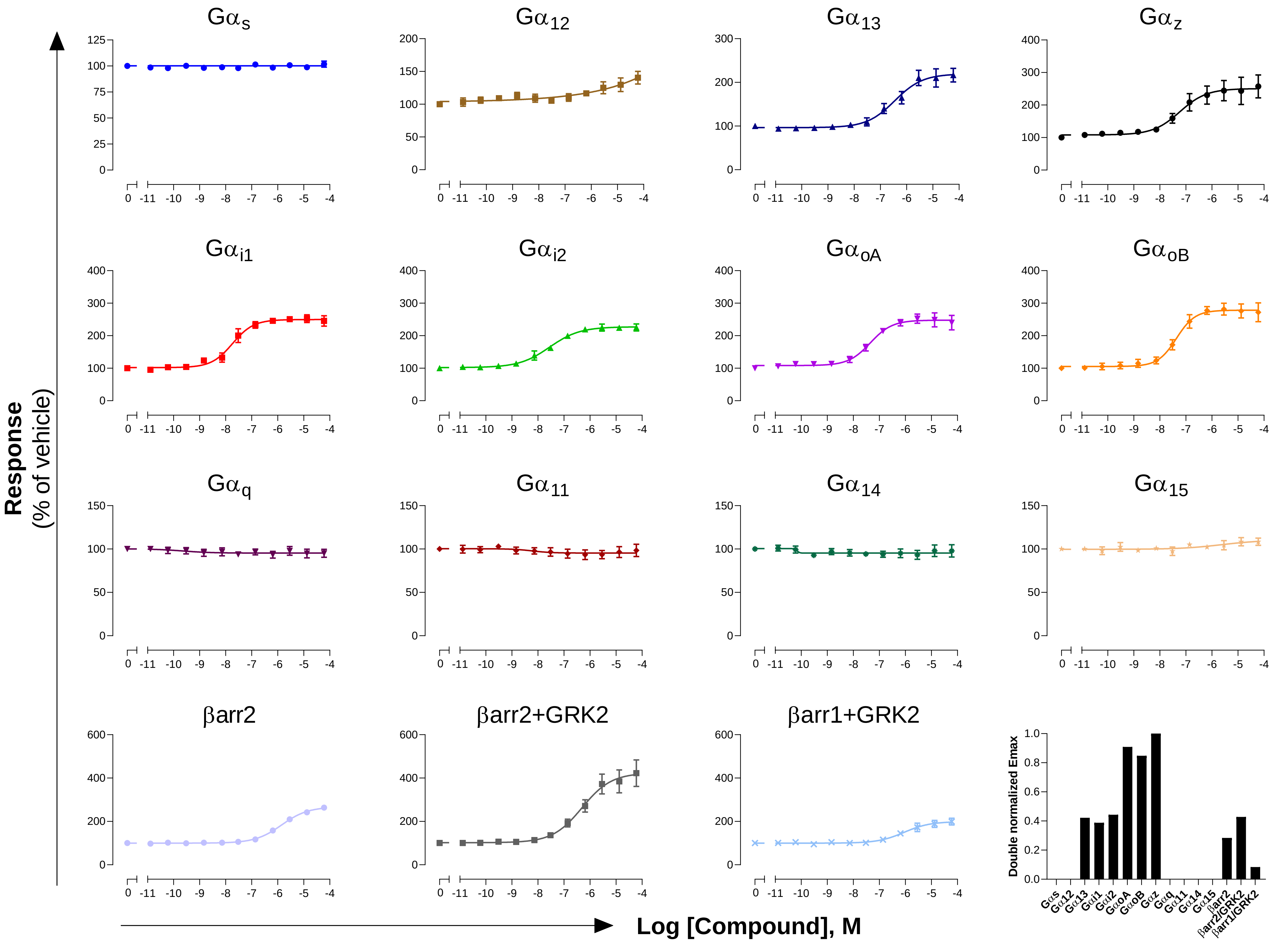

**GPCR: $HCA_3$**
**Ligand: 3-hydroxyoctanoic acid (3-HOA)**

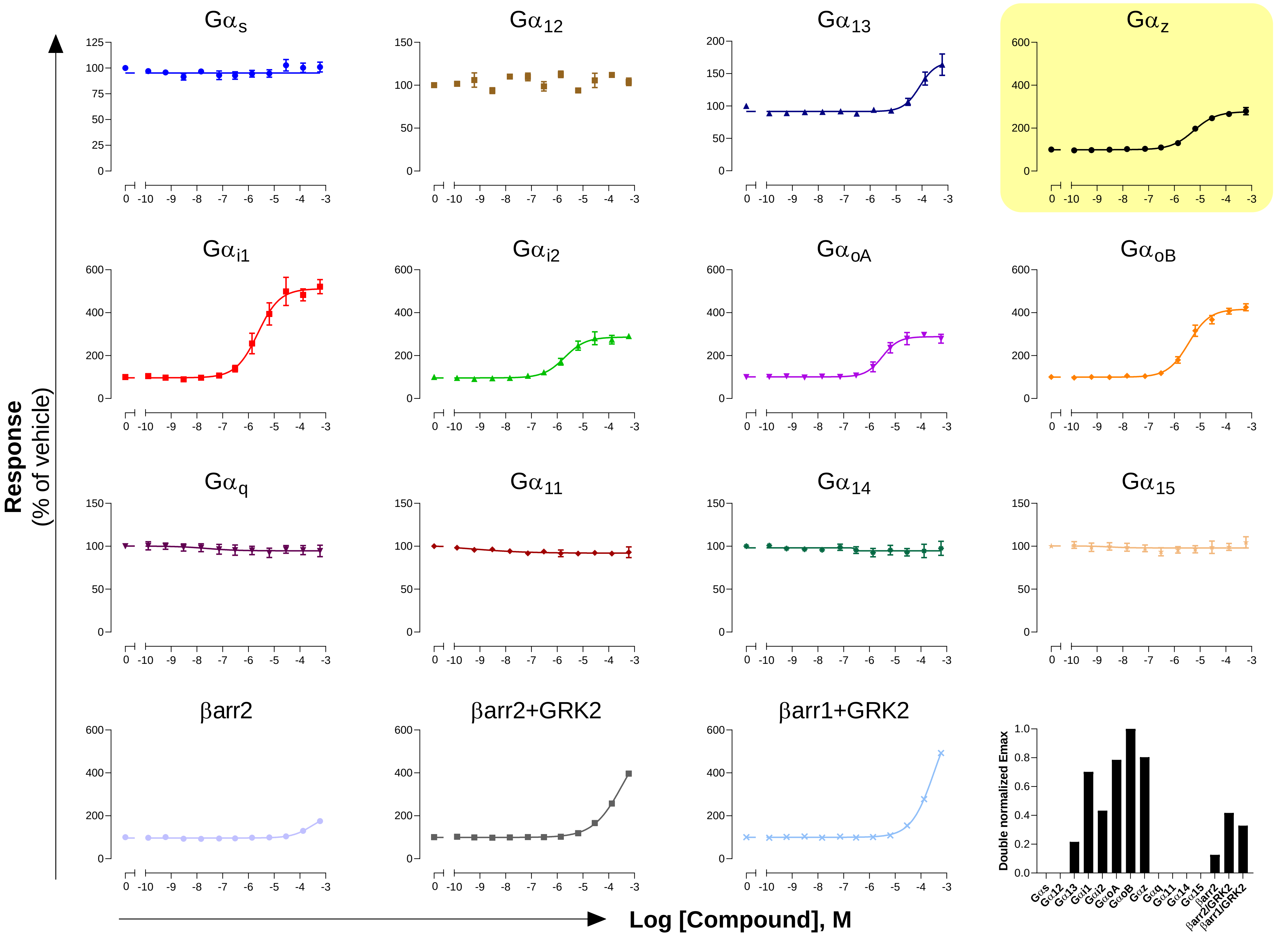

### GPCR: κOR
### Ligand: Dynorphin A

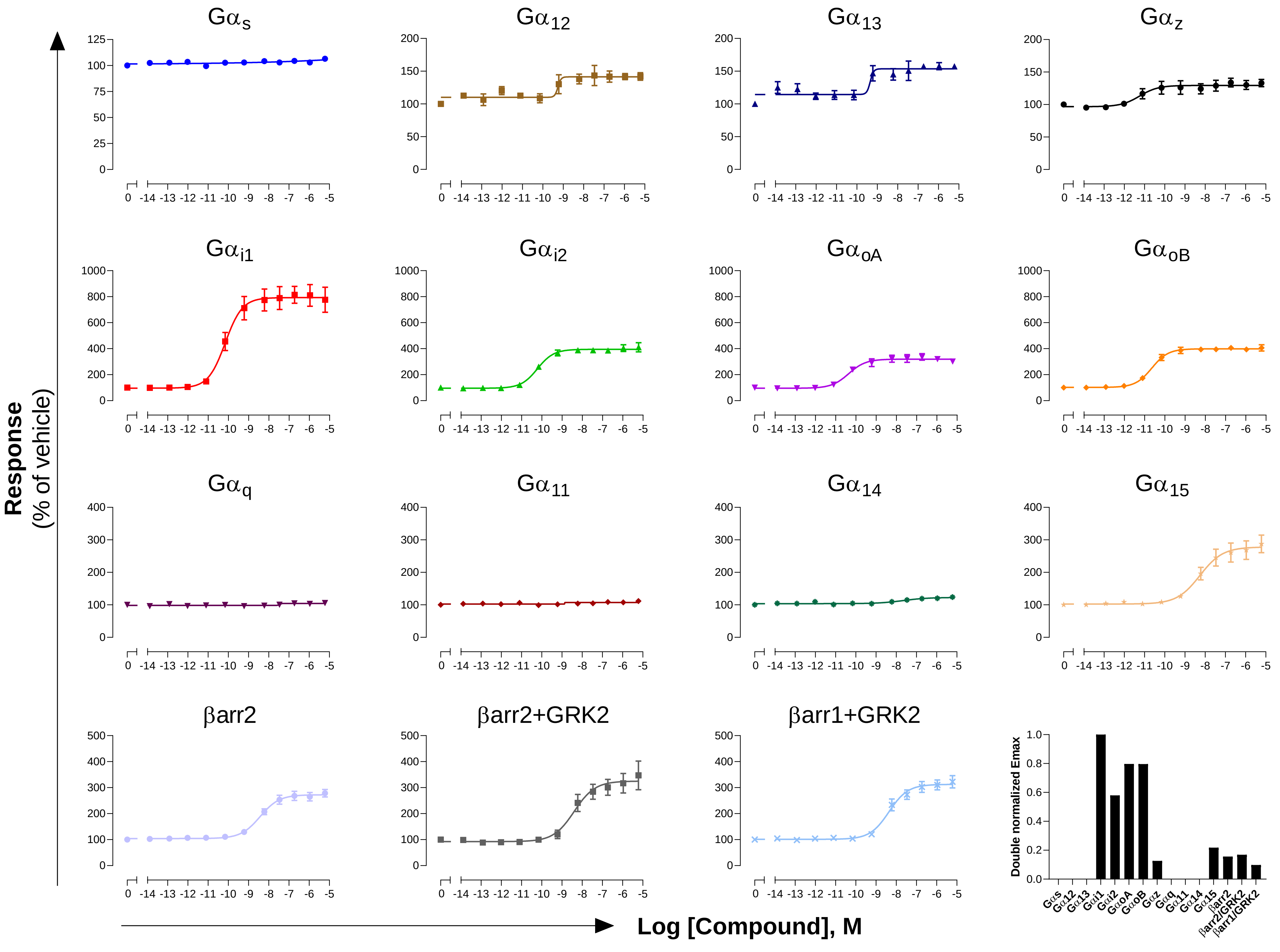

**GPCR: LPA$_1$**

**Ligand: oleoyl-LPA**

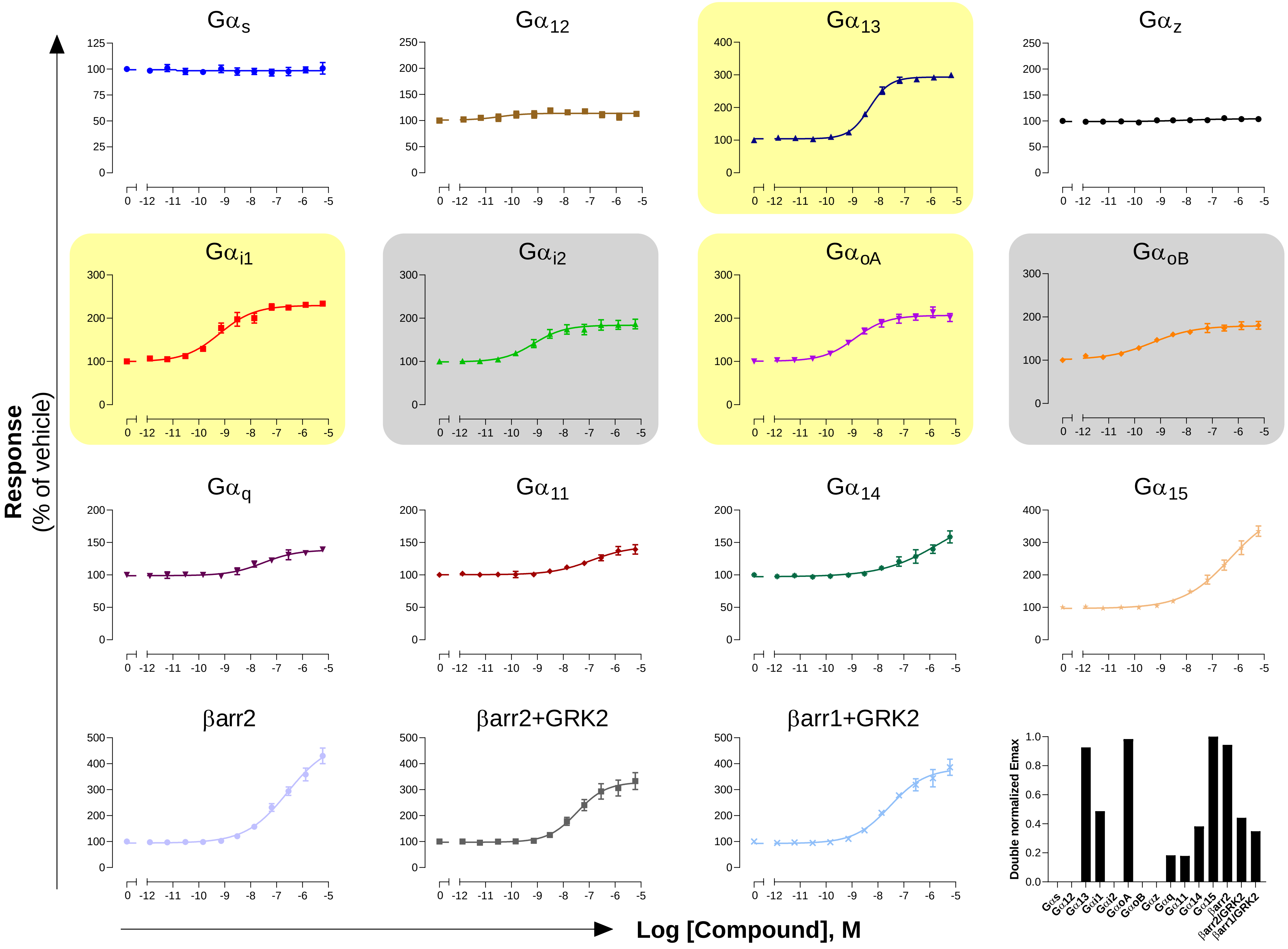

**GPCR: $LPA_2$**

**Ligand: oleoyl-LPA**

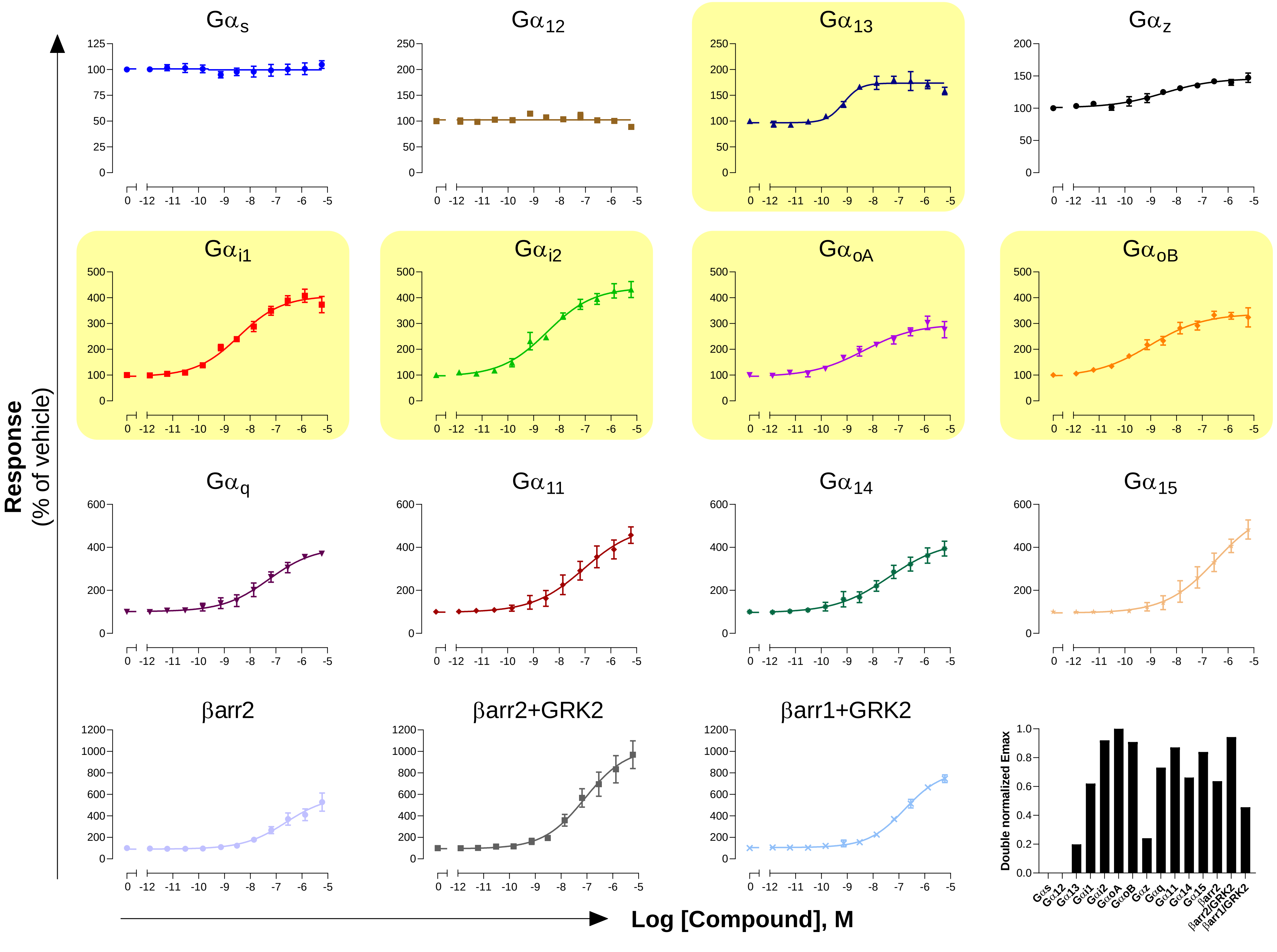

#### GPCR: $M_1$
#### Ligand: Acetylcholine

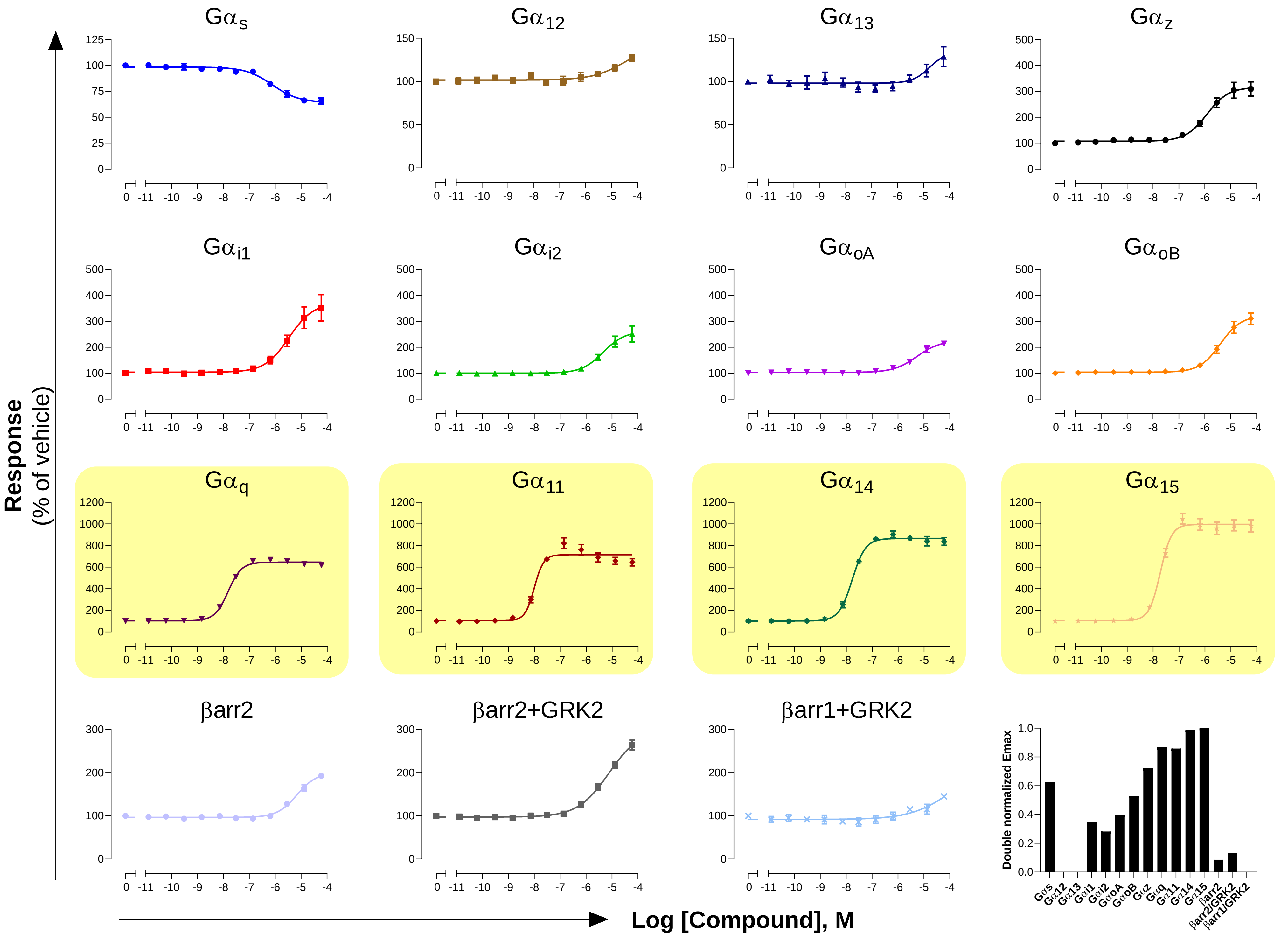

**GPCR: $M_2$**

**Ligand: Acetylcholine**

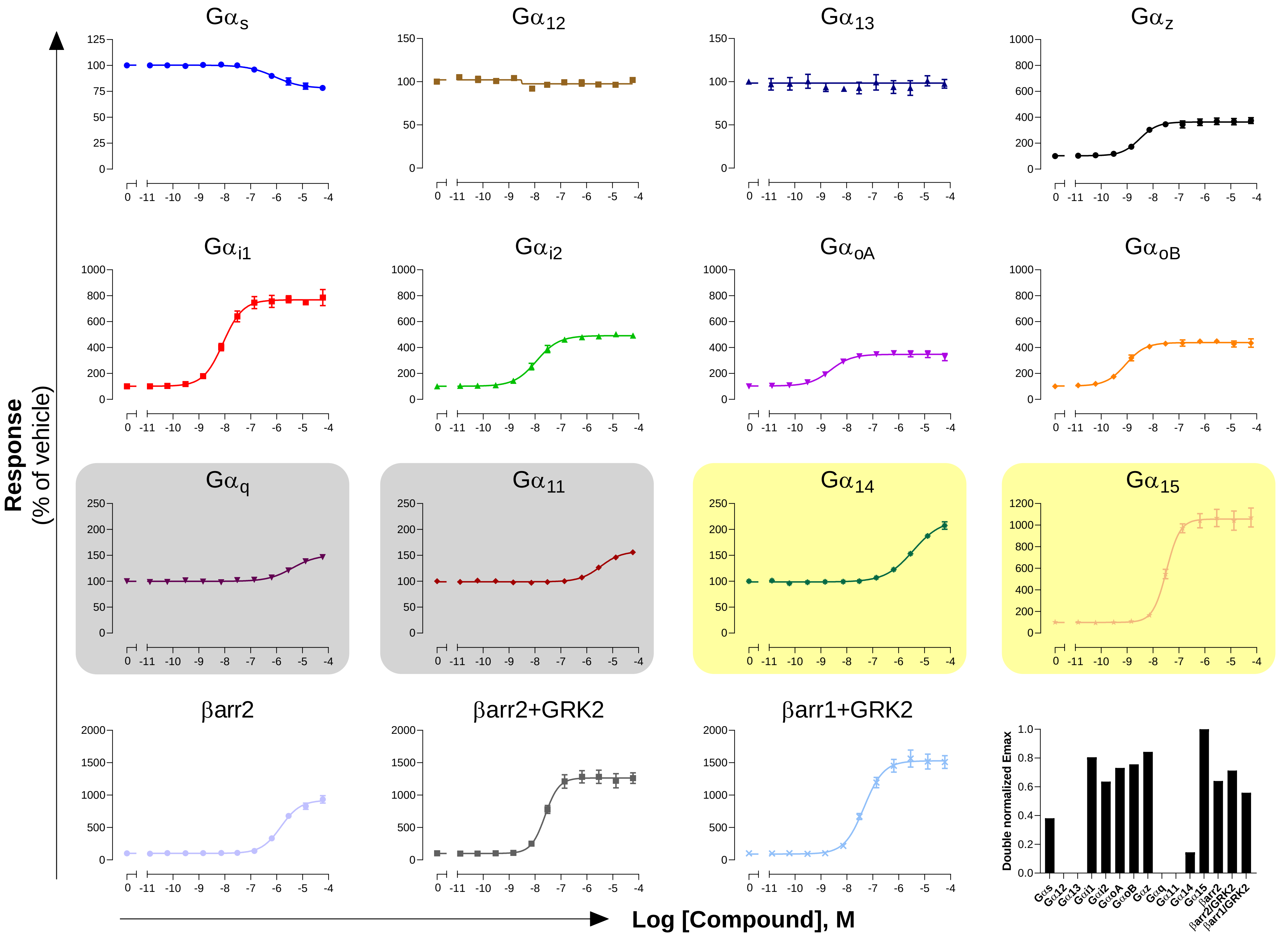

**GPCR: M$_3$**

**Ligand: Acetylcholine**

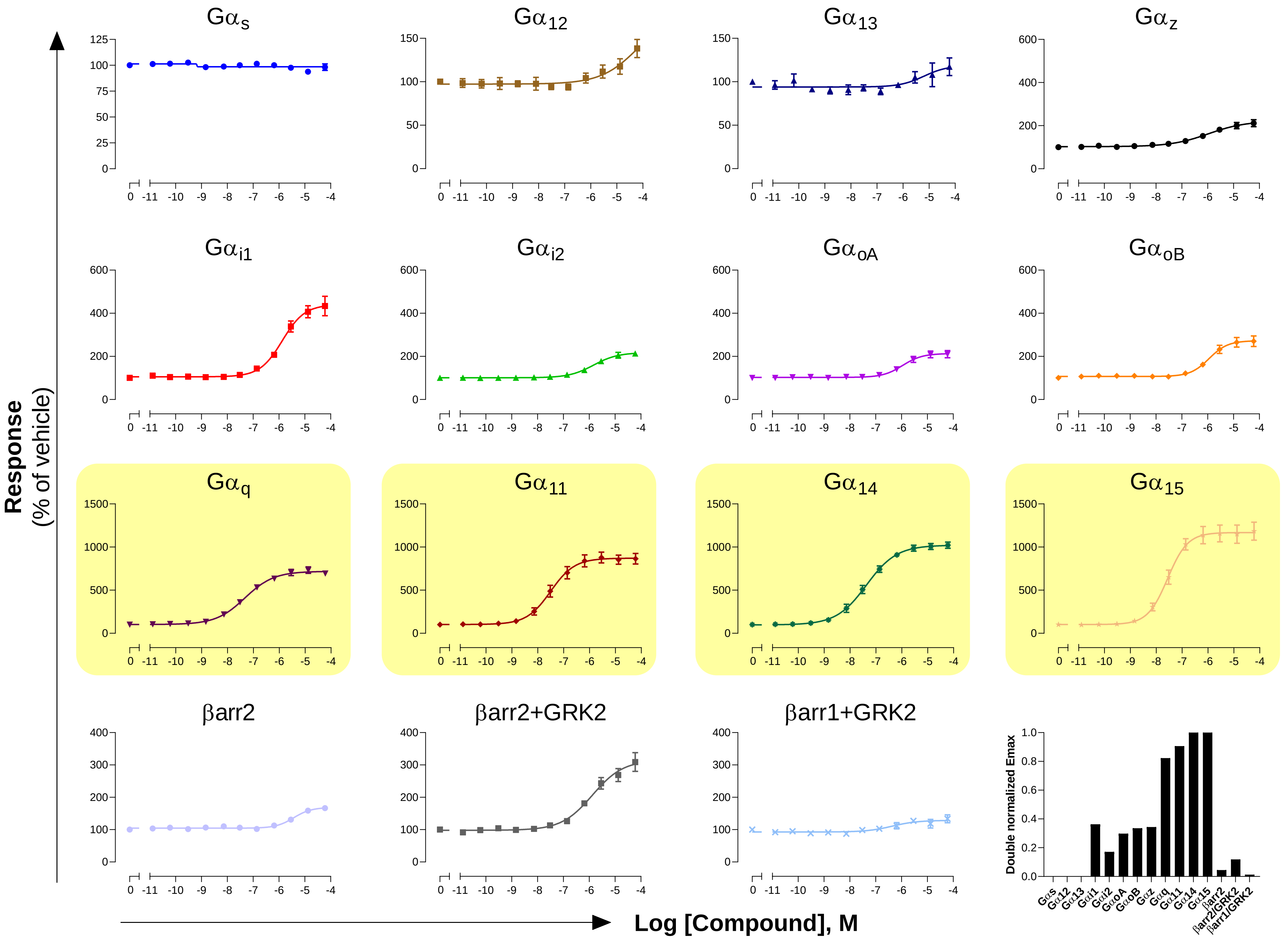

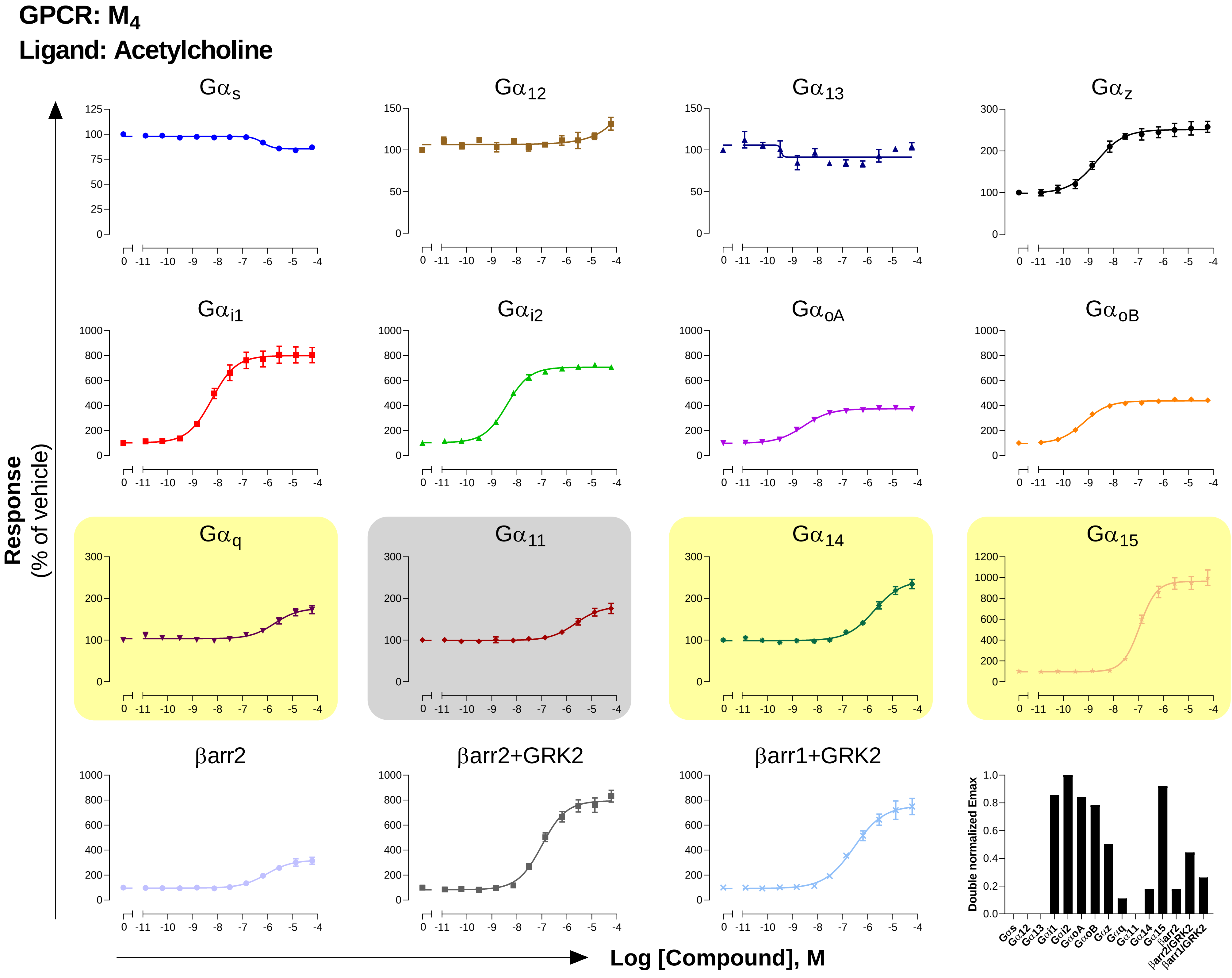
GPCR: M4
Ligand: Acetylcholine
Gαs
Gα12
Gα13
Gαz
Gαi1
Gαi2
GαoA
GαoB
Gαq
Gα11
Gα14
Gα15
βarr2
βarr2+GRK2
βarr1+GRK2
Response
(% of vehicle)
Log [Compound], M
Double normalized Emax

### GPCR: MC3R
### Ligand: γ-MSH

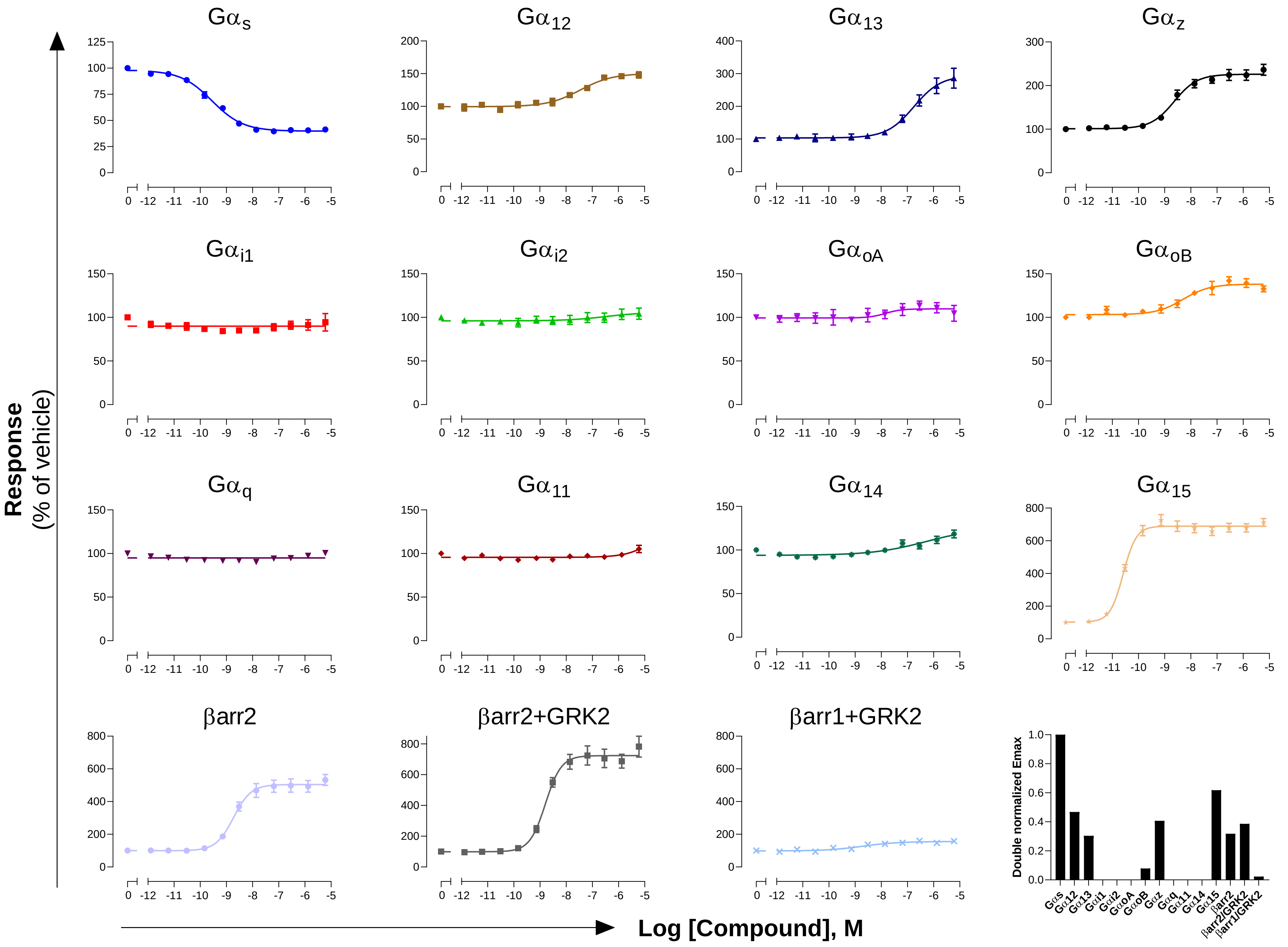

**GPCR: MC4R**
**Ligand: α-MSH**

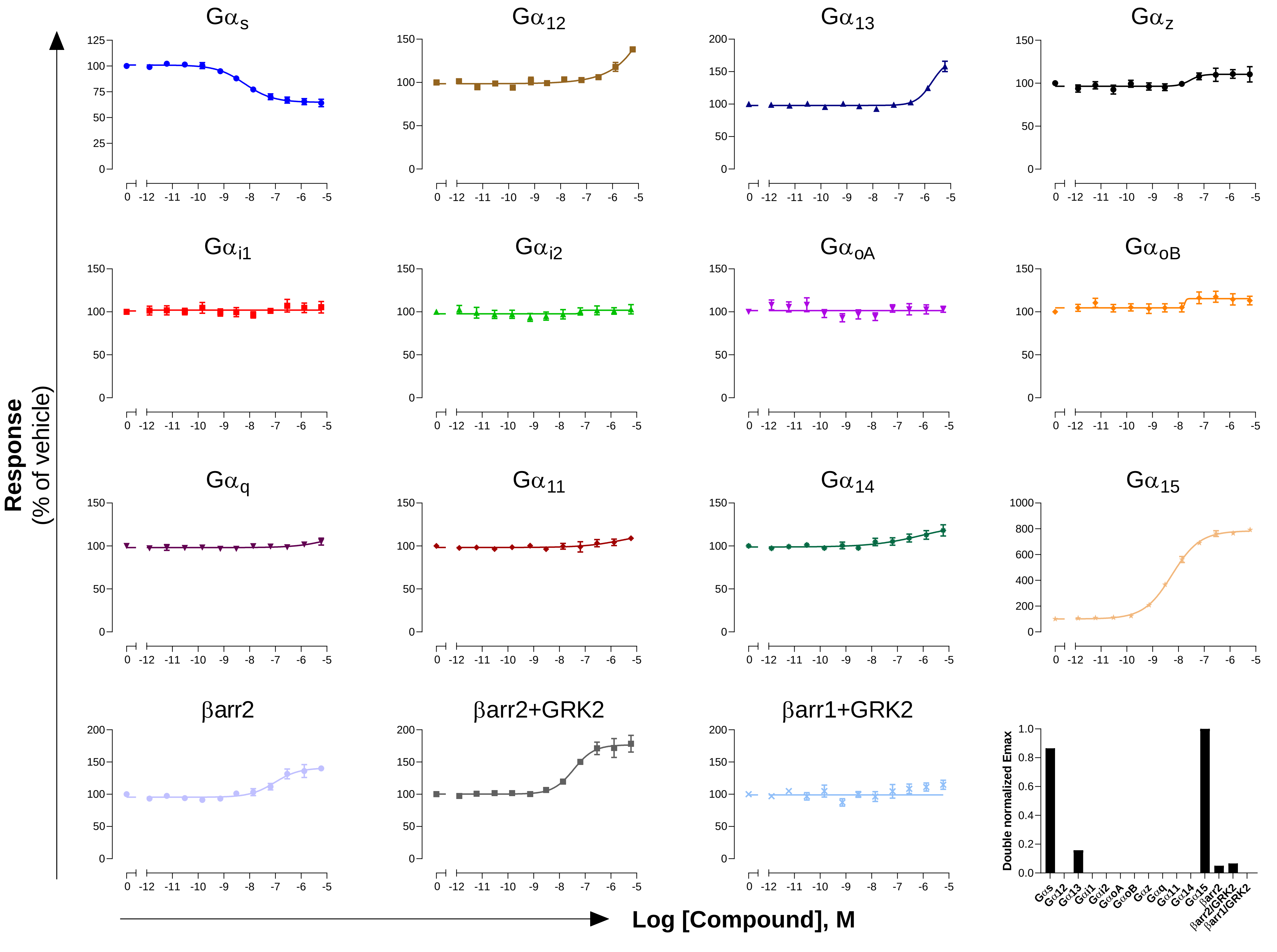

GPCR: mGluR2
Ligand: Glutamate

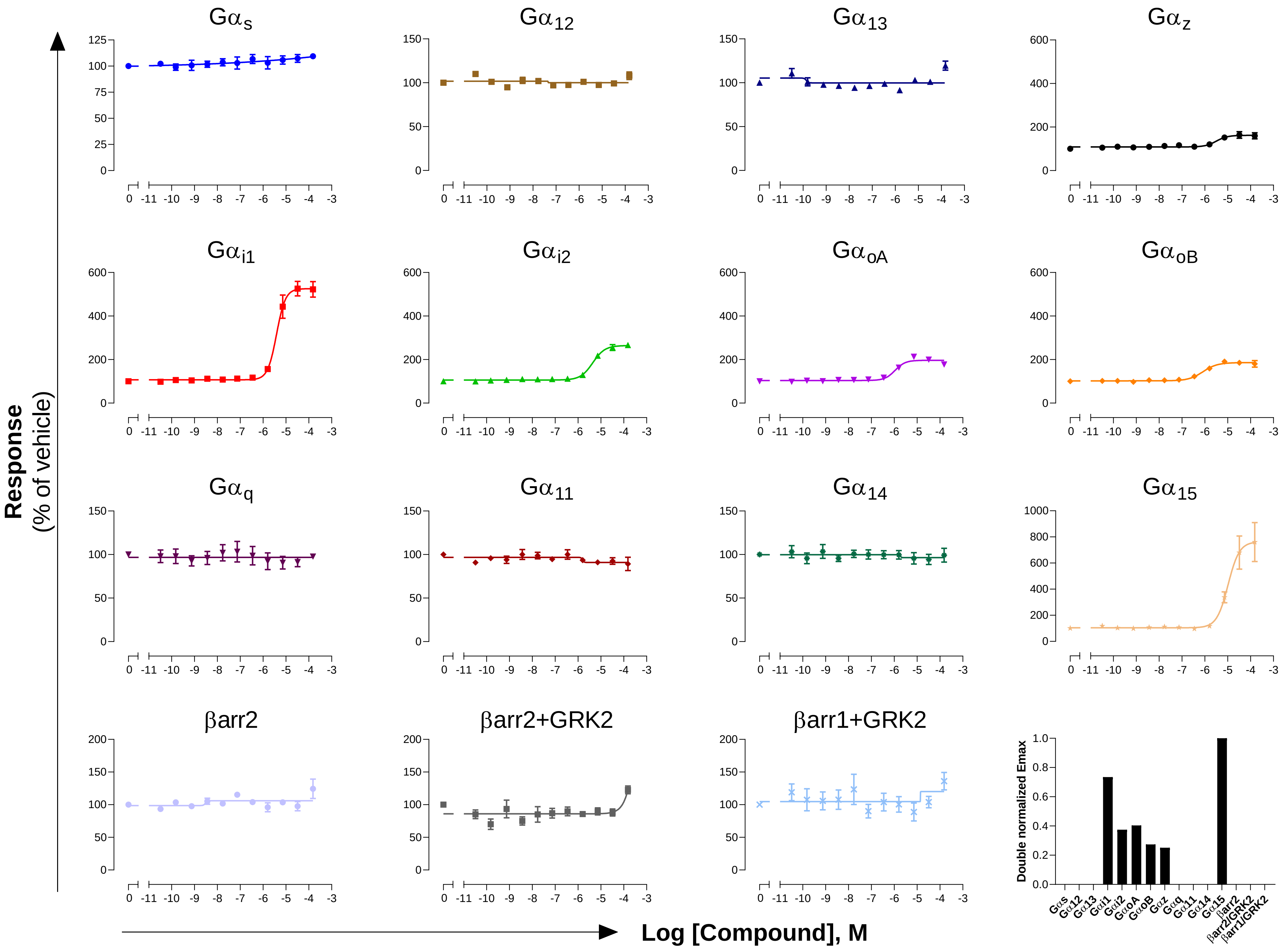

**GPCR: mGluR4**
**Ligand: Glutamate**

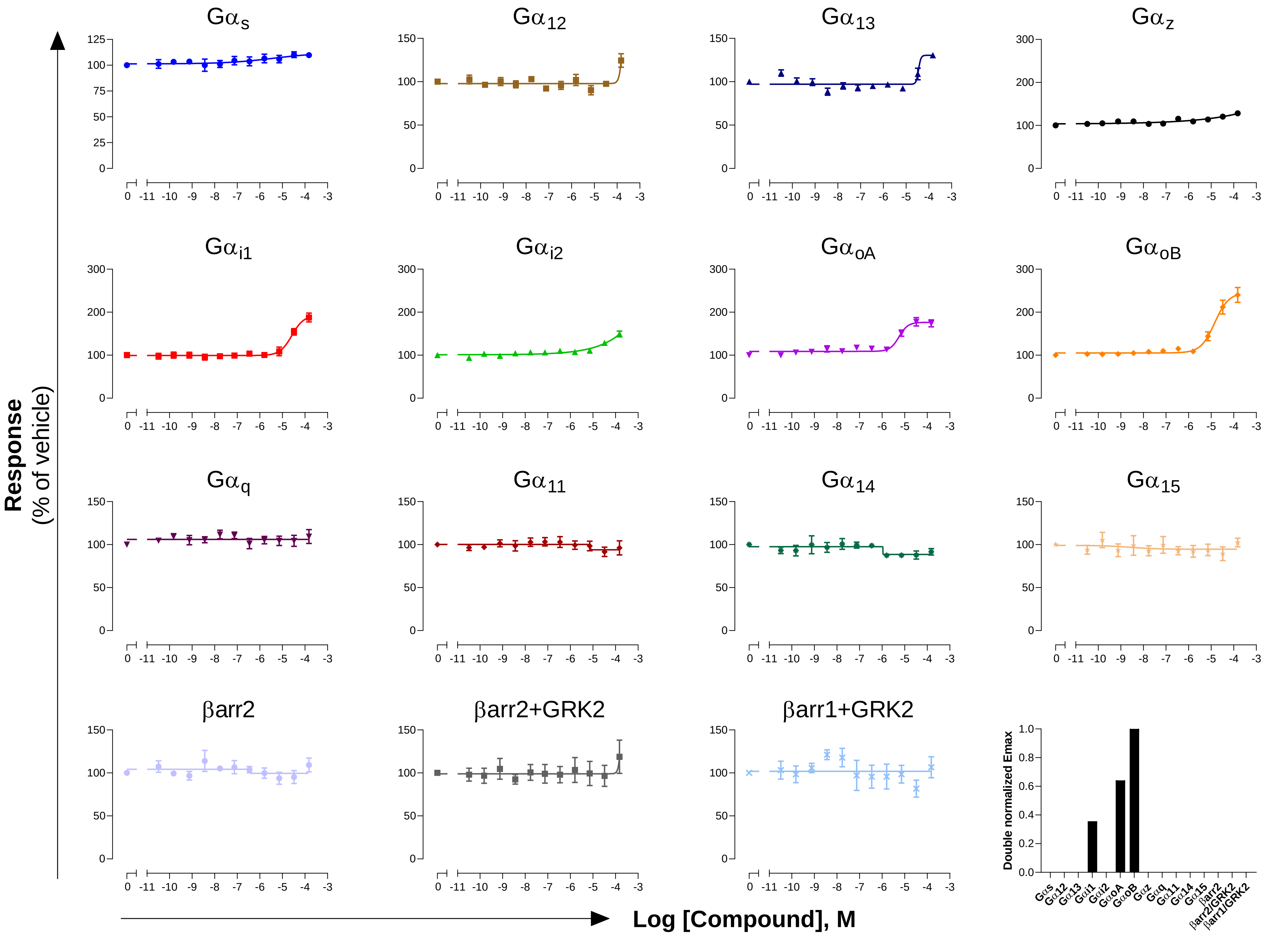

**GPCR: mGluR5**
**Ligand: Glutamate**

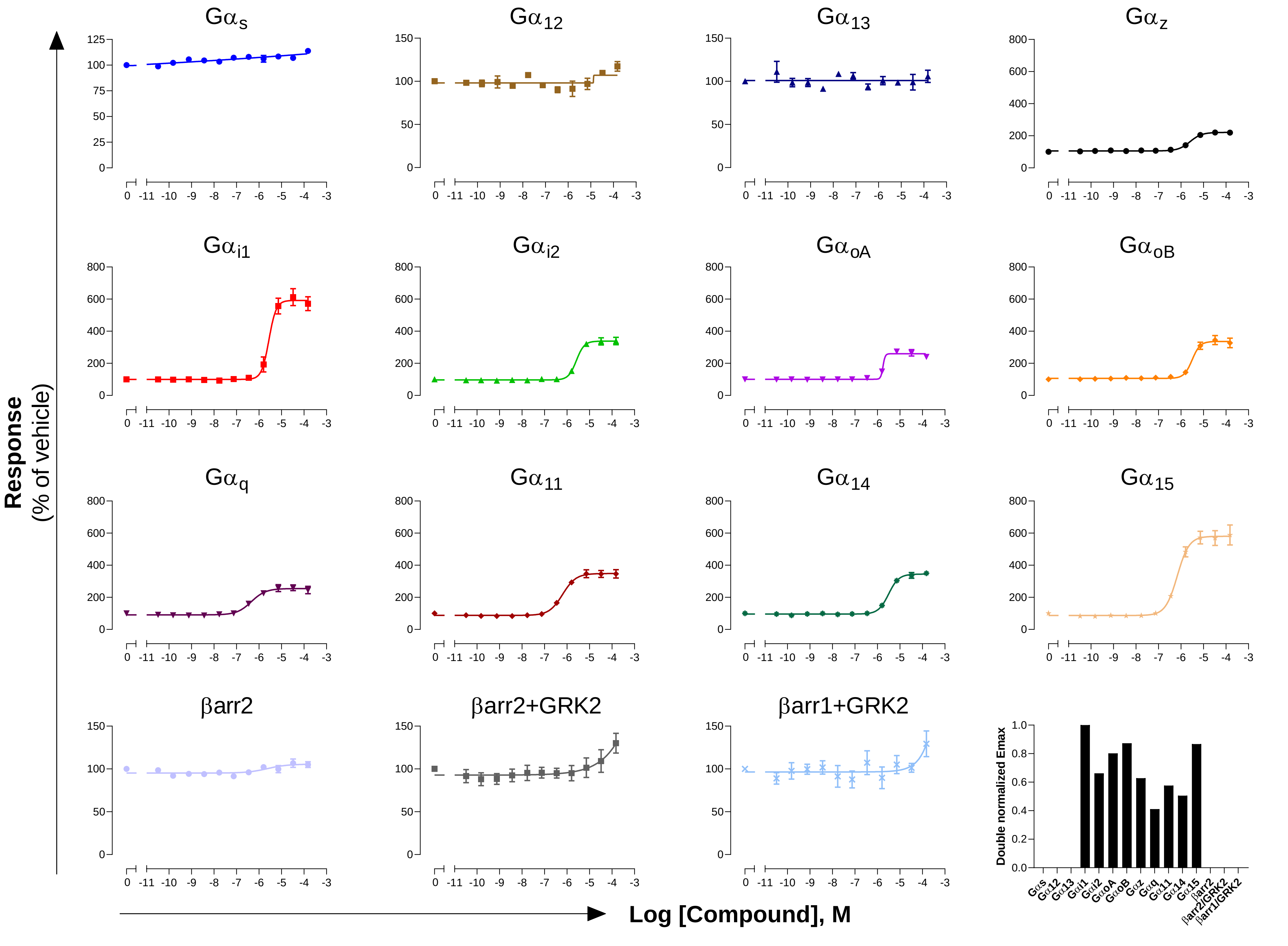

**GPCR: mGluR6**
**Ligand: Glutamate**

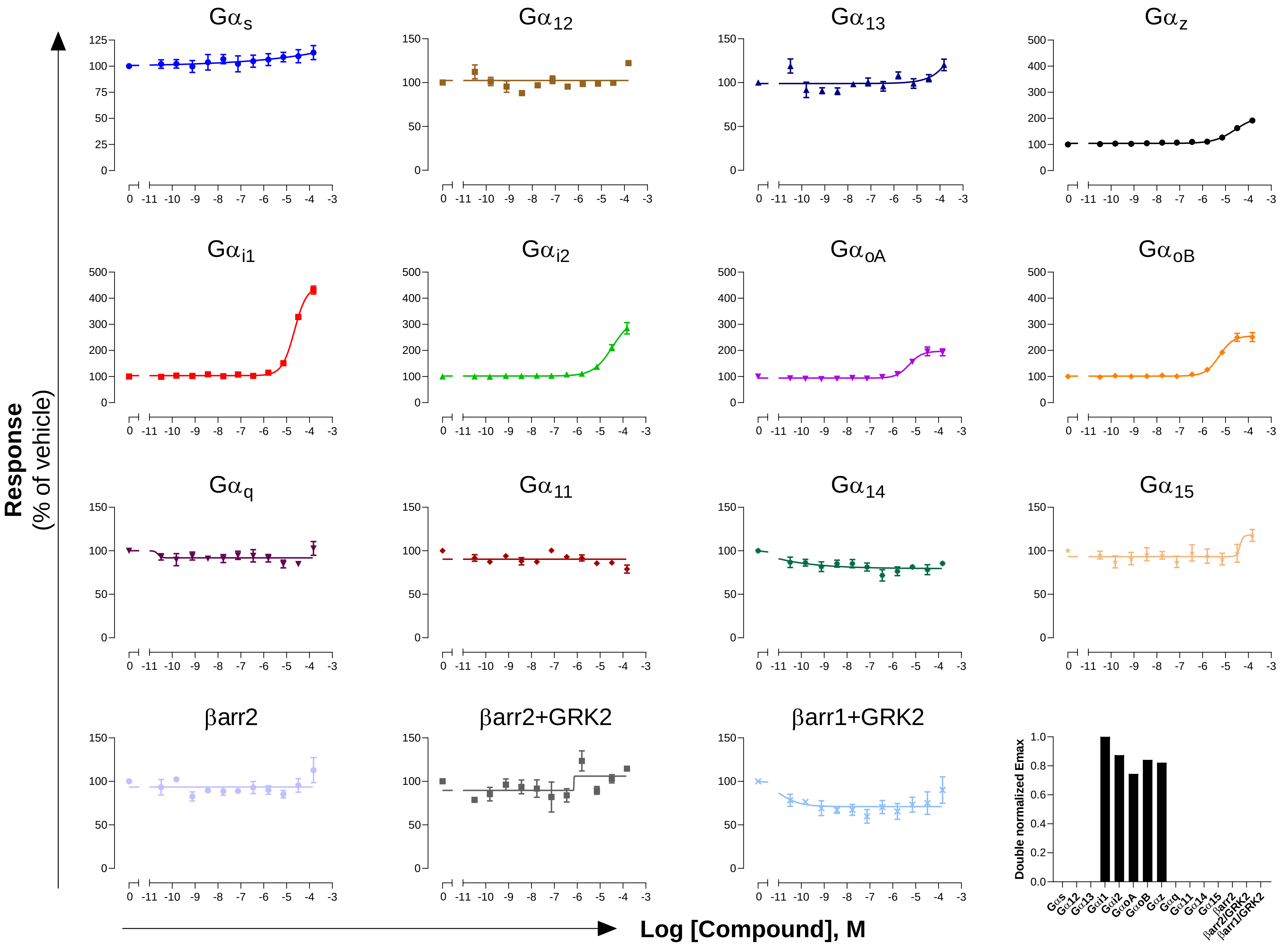

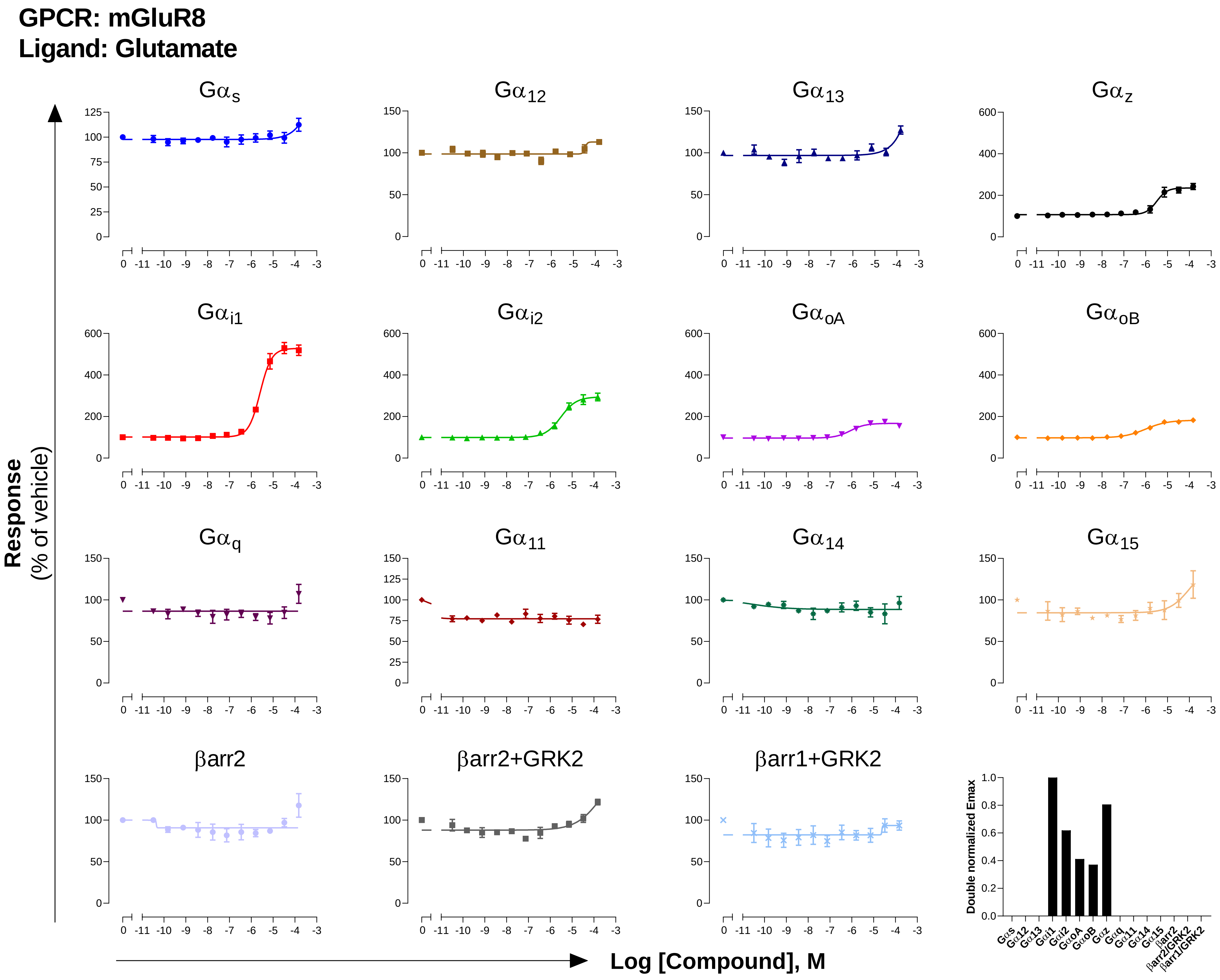

GPCR: mGluR8
Ligand: Glutamate
Gαs
Gα12
Gα13
Gαz
Gαi1
Gαi2
GαoA
GαoB
Gαq
Gα11
Gα14
Gα15
βarr2
βarr2+GRK2
βarr1+GRK2
Response
(% of vehicle)
Log [Compound], M
Double normalized Emax

**GPCR: μOR**
**Ligand: DAMGO**

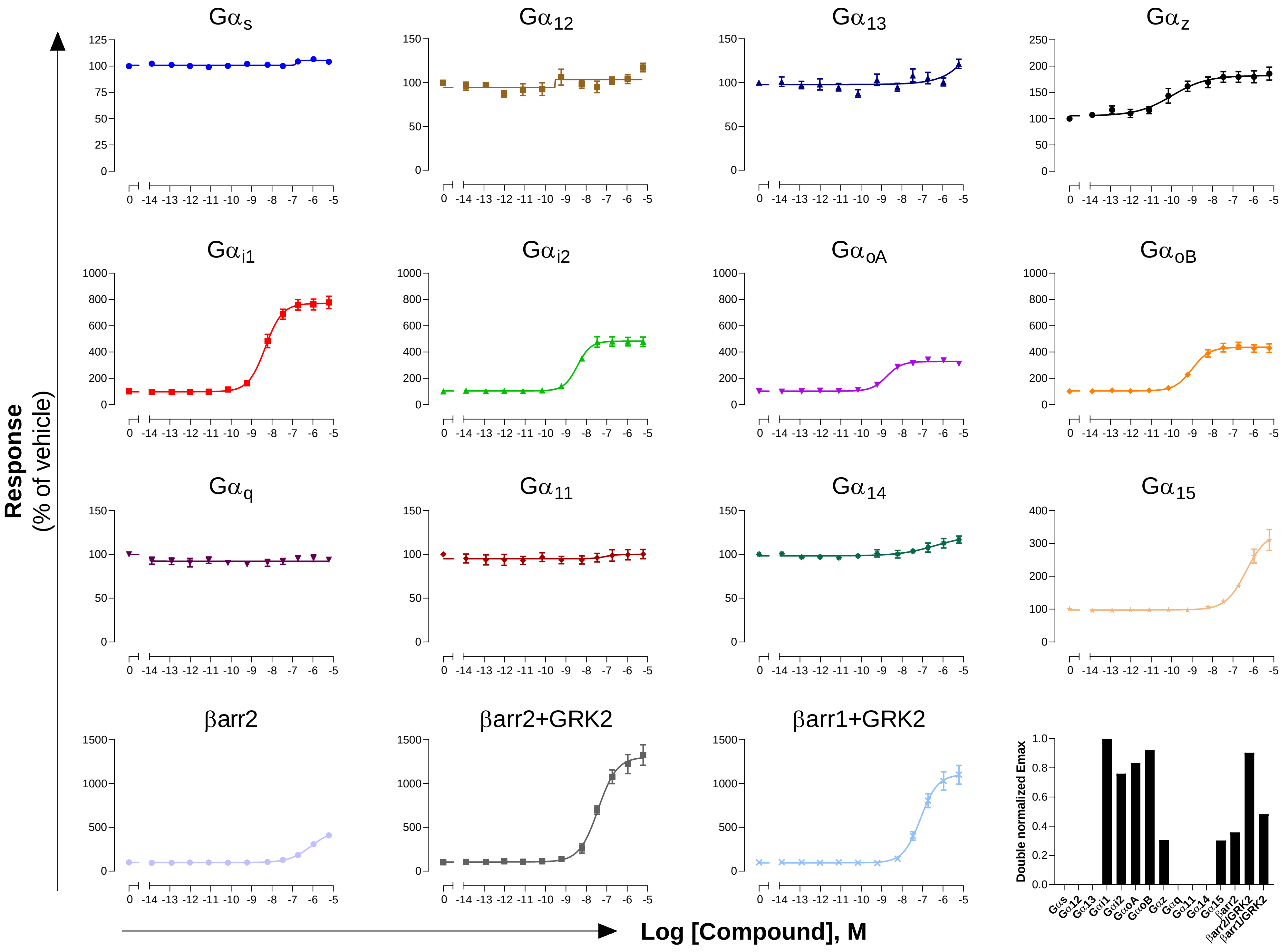

**GPCR: $MT_1$**
**Ligand: Melatonin**

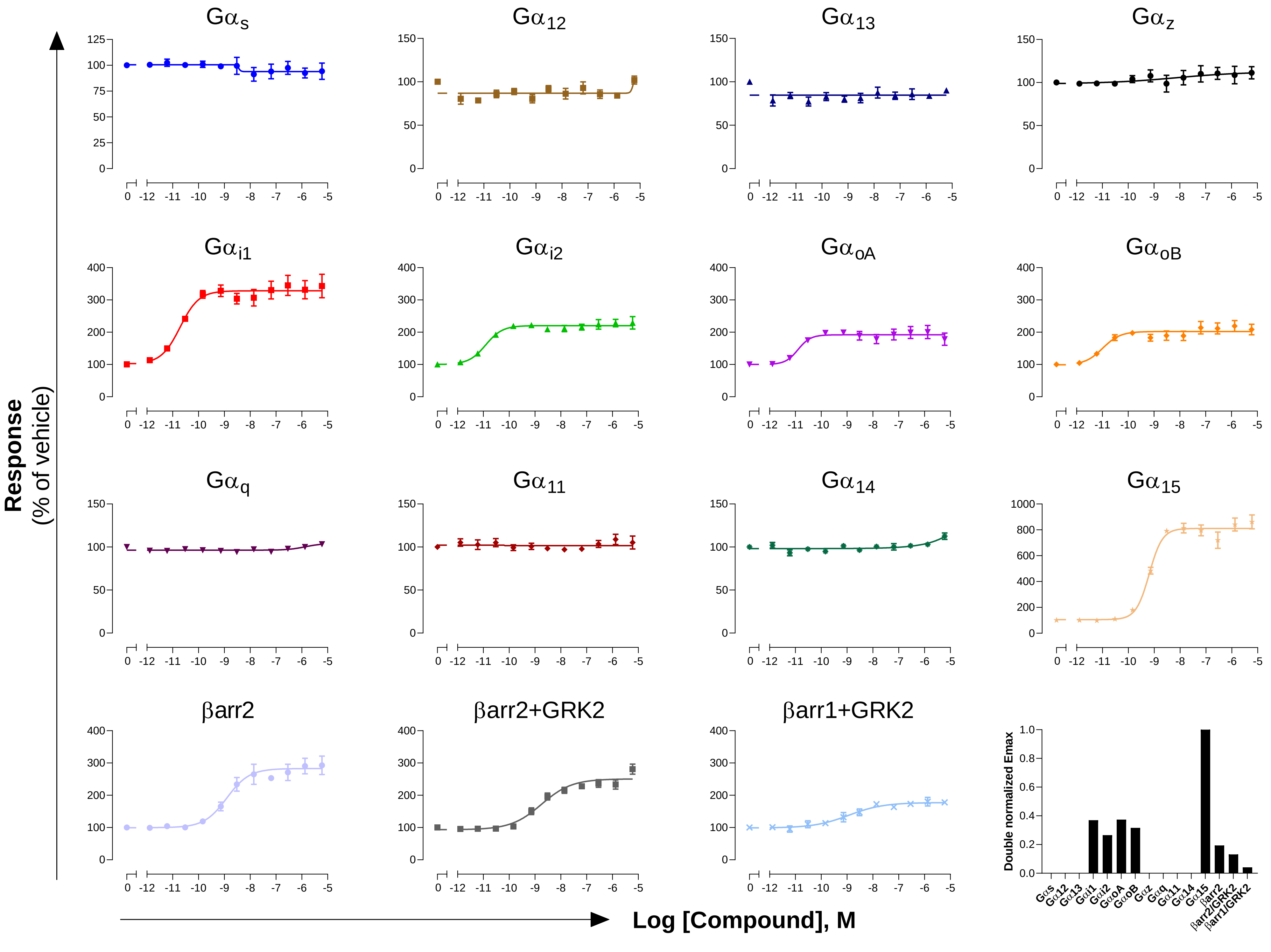

**GPCR: MT$_2$**
**Ligand: Melatonin**

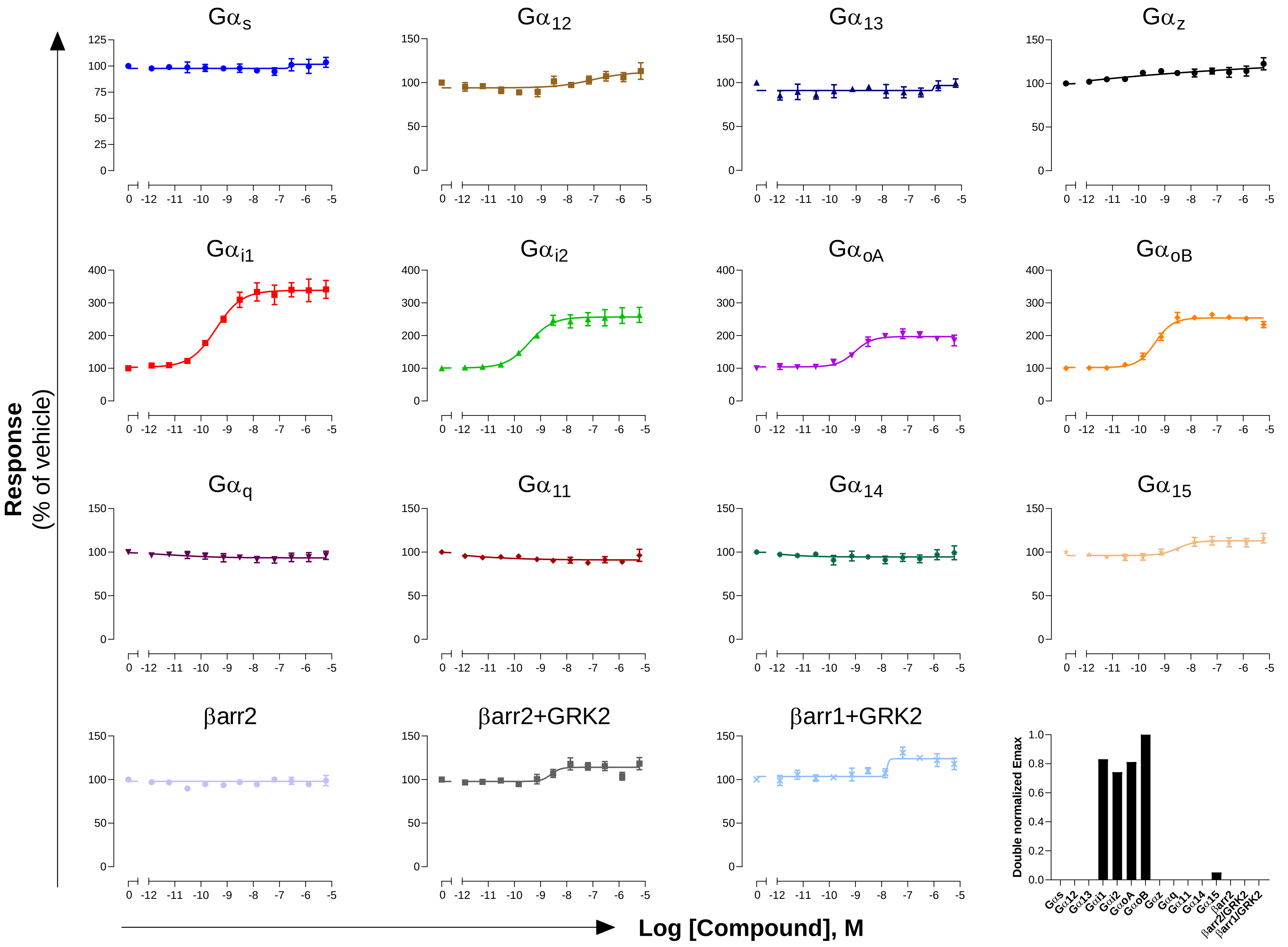

**GPCR: NOP**
**Ligand: Nociceptin**

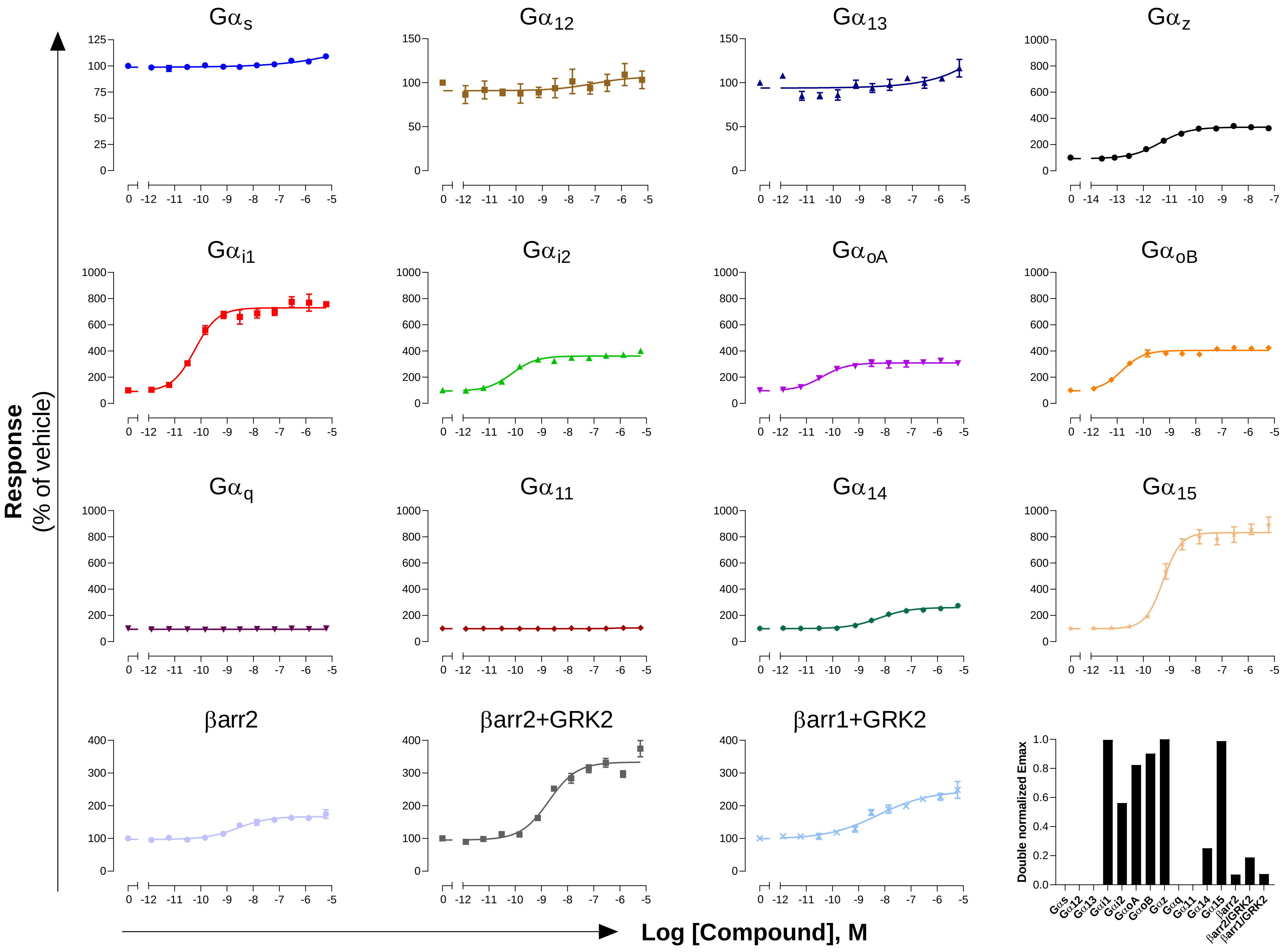

**GPCR: NPFF1**
**Ligand: RFRP3**

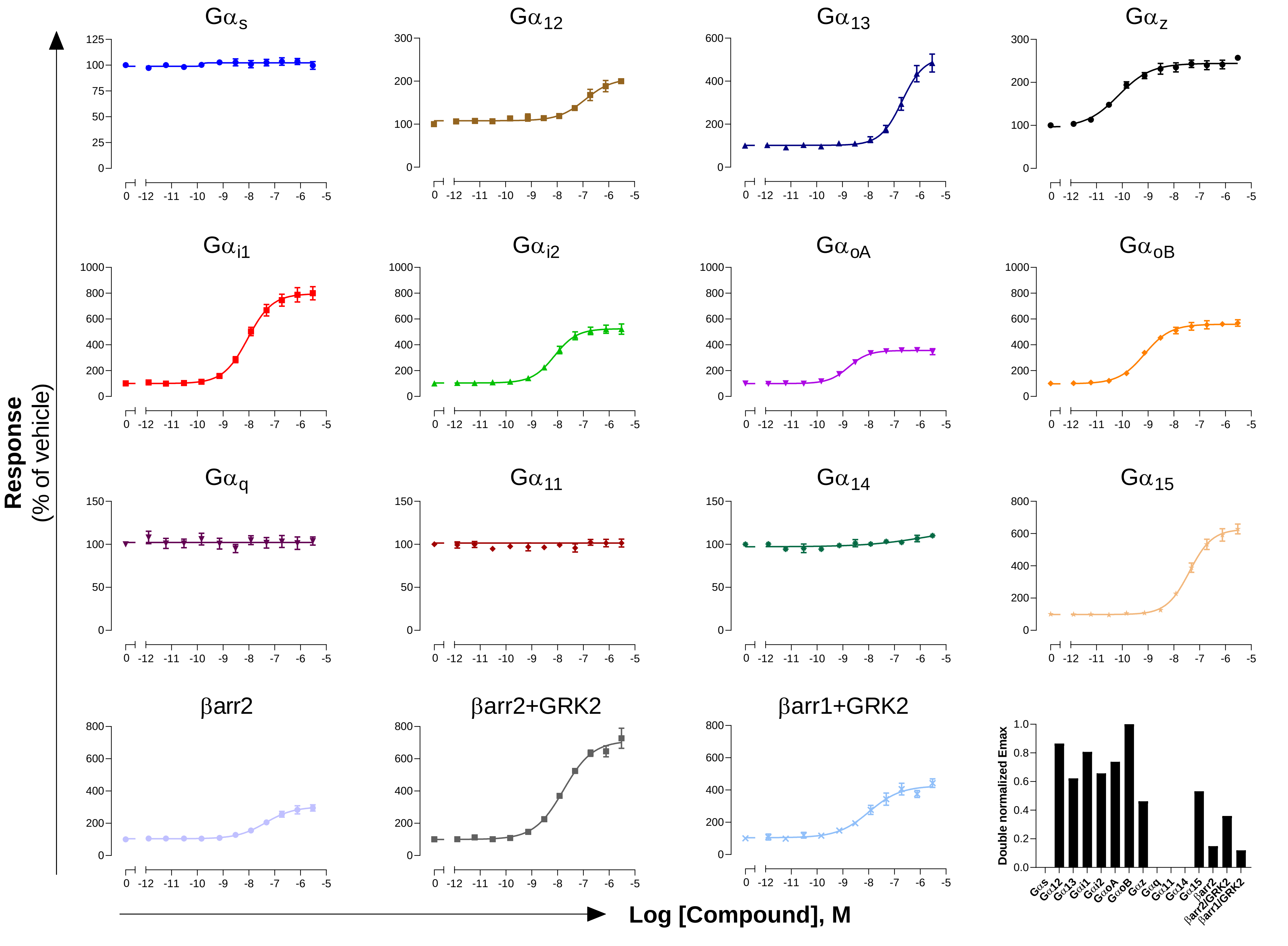

**GPCR: NPFF2**
**Ligand: NPFF**

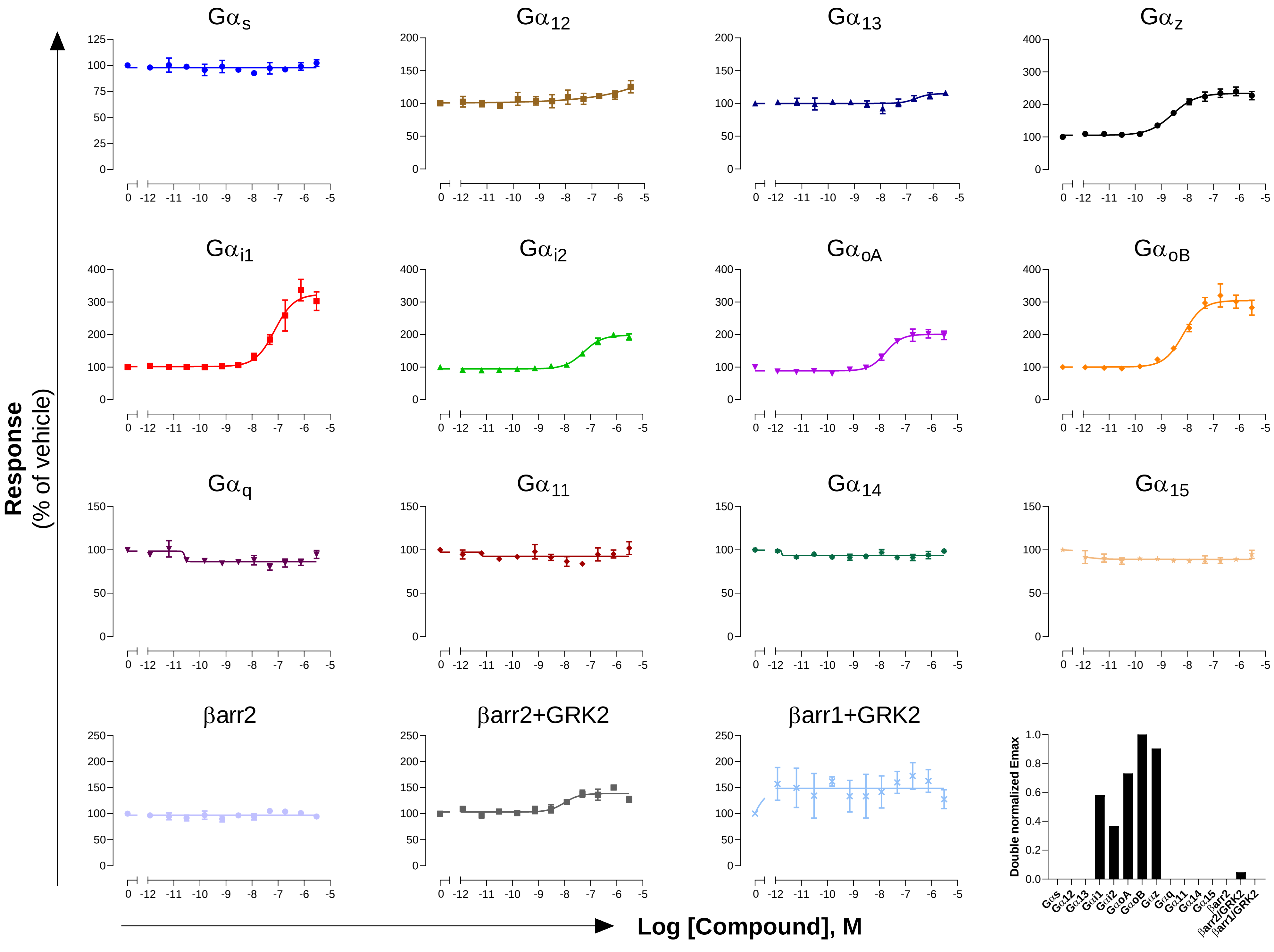

### GPCR: OT
### Ligand: Oxytocin

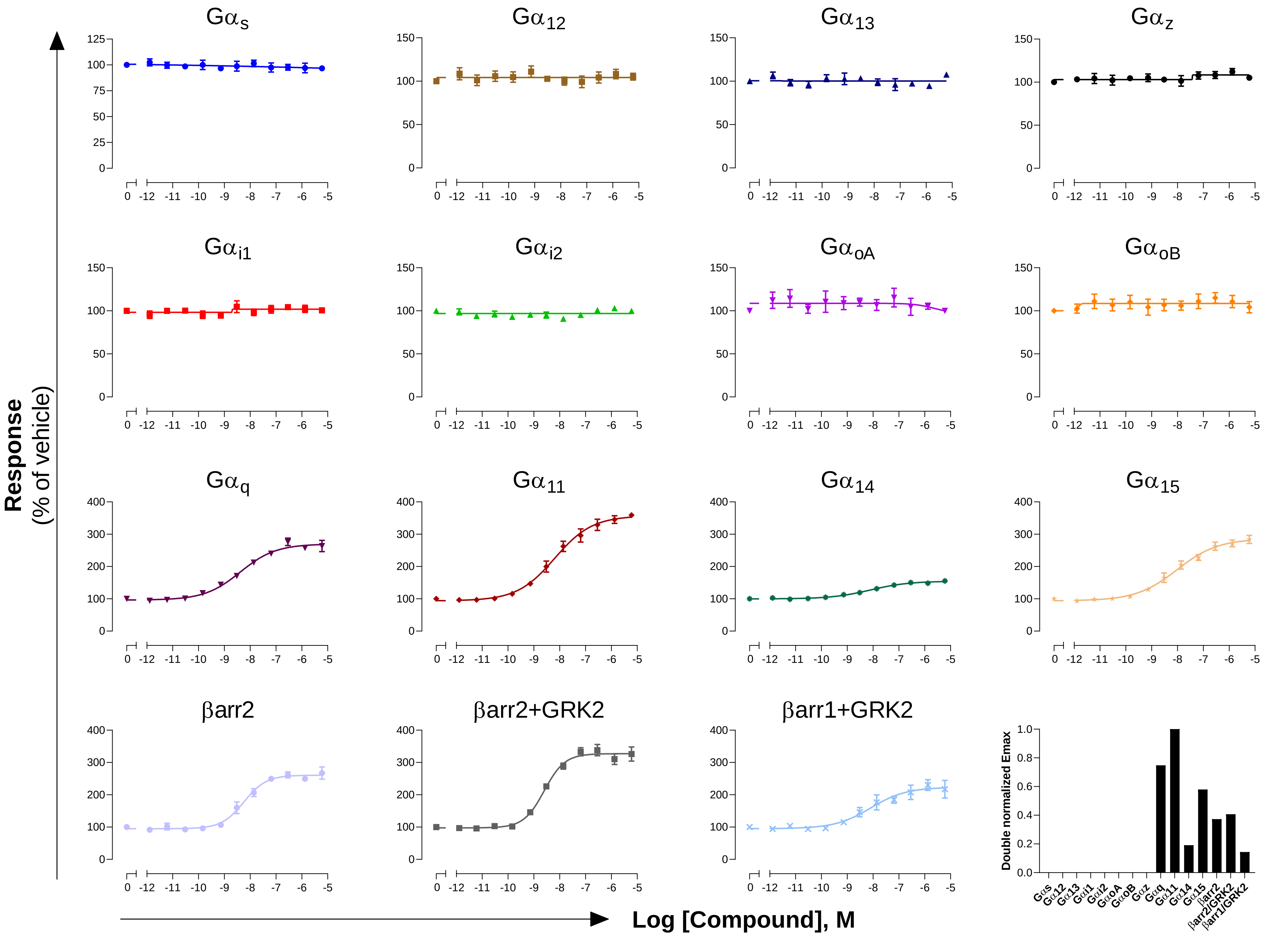

**GPCR: $OX_2$**

**Ligand: Orexin-A**

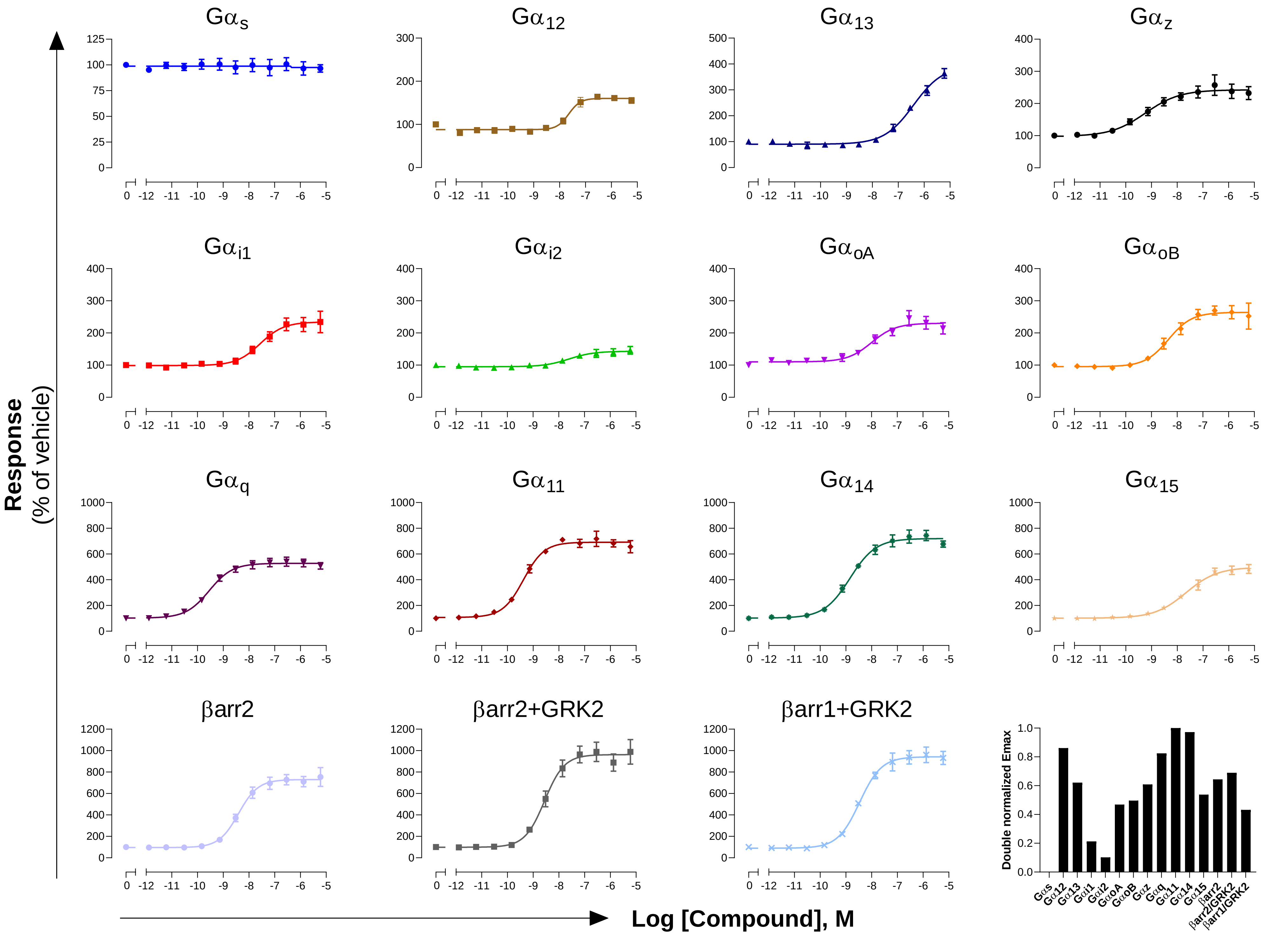

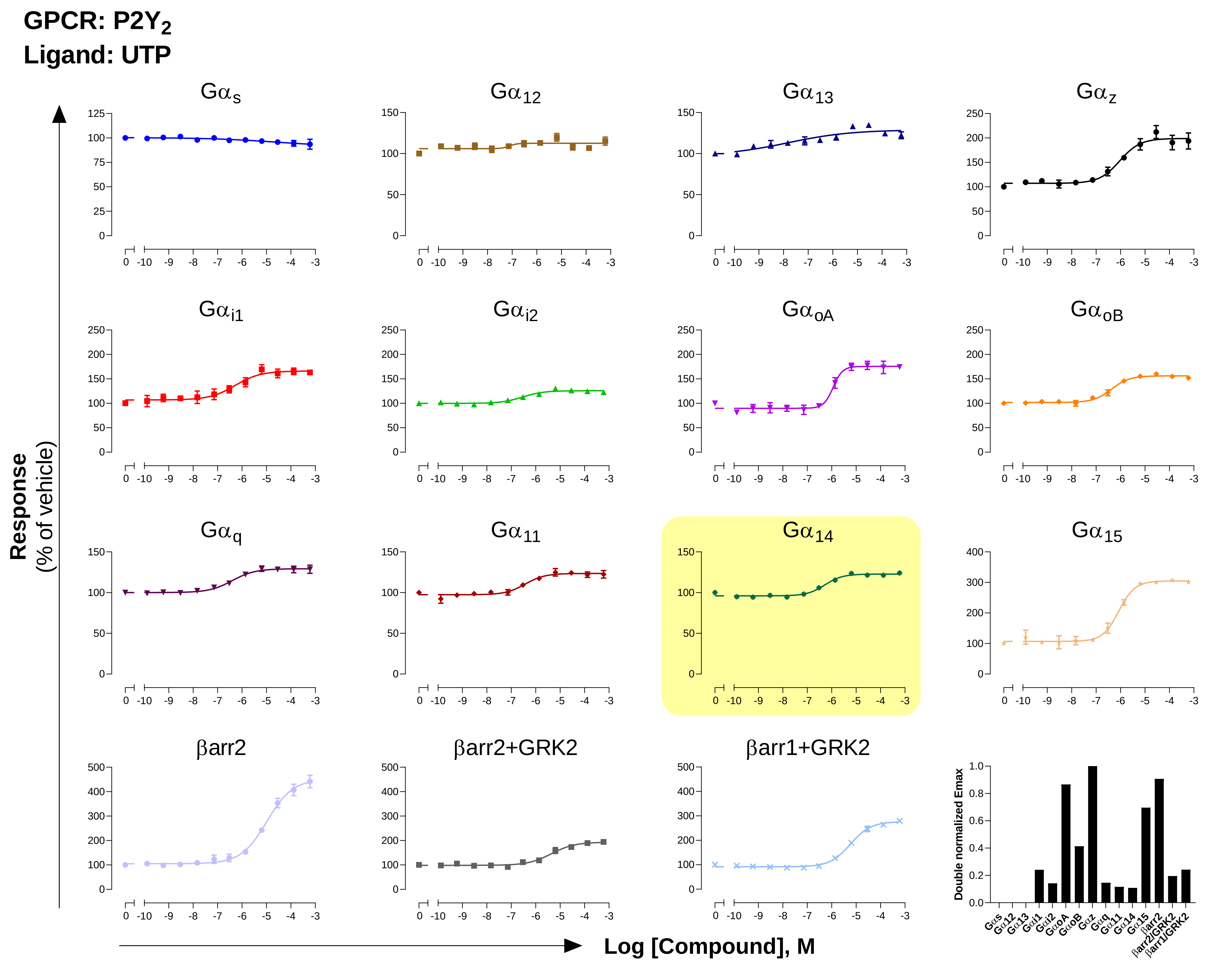
GPCR: P2Y2
Ligand: UTP
Gαs
Gα12
Gα13
Gαz
Gαi1
Gαi2
GαoA
GαoB
Gαq
Gα11
Gα14
Gα15
βarr2
βarr2+GRK2
βarr1+GRK2
Response
(% of vehicle)
Log [Compound], M
Double normalized Emax

GPCR: PAR1
Ligand: TFLLR-NH$_2$

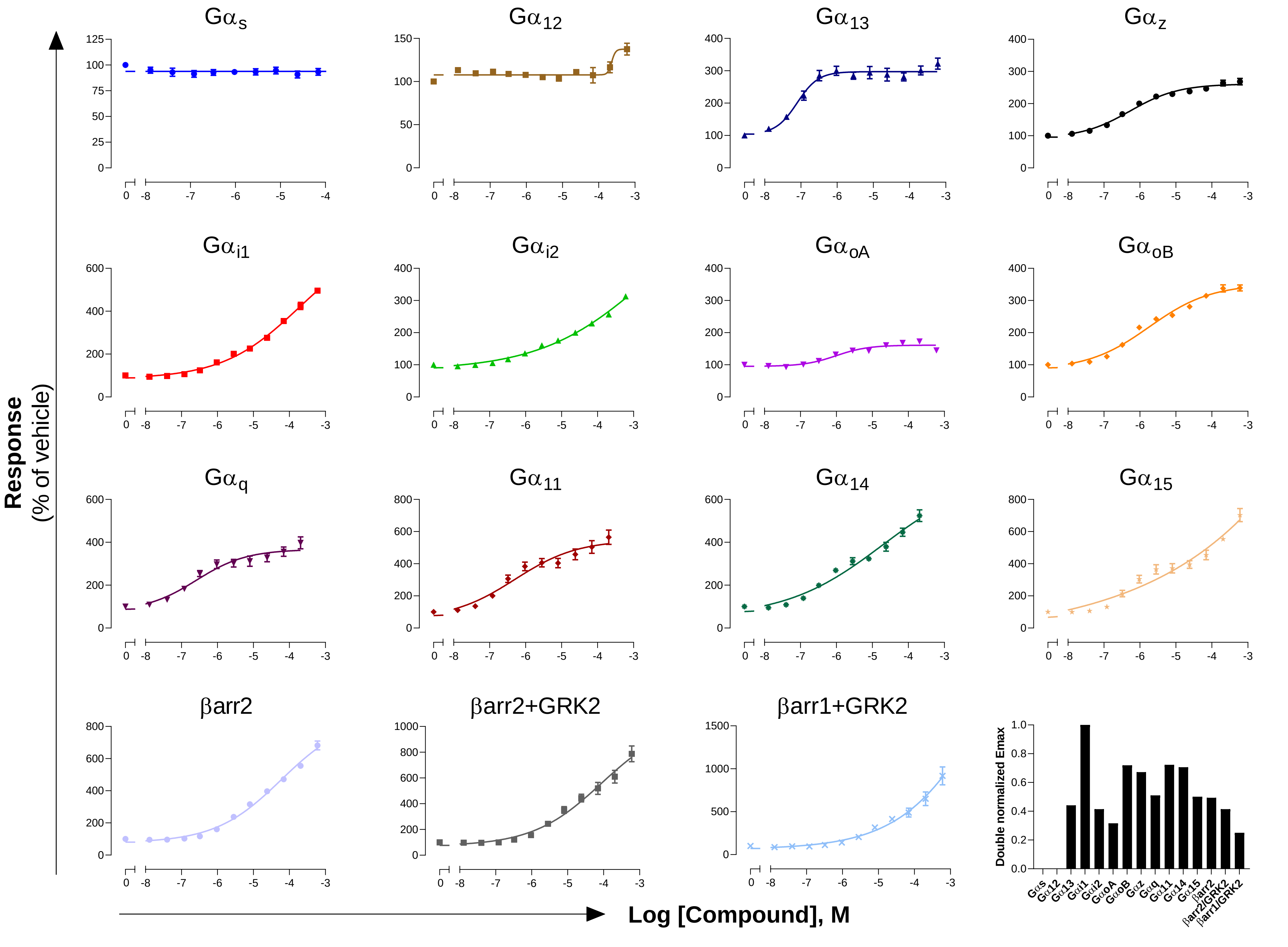

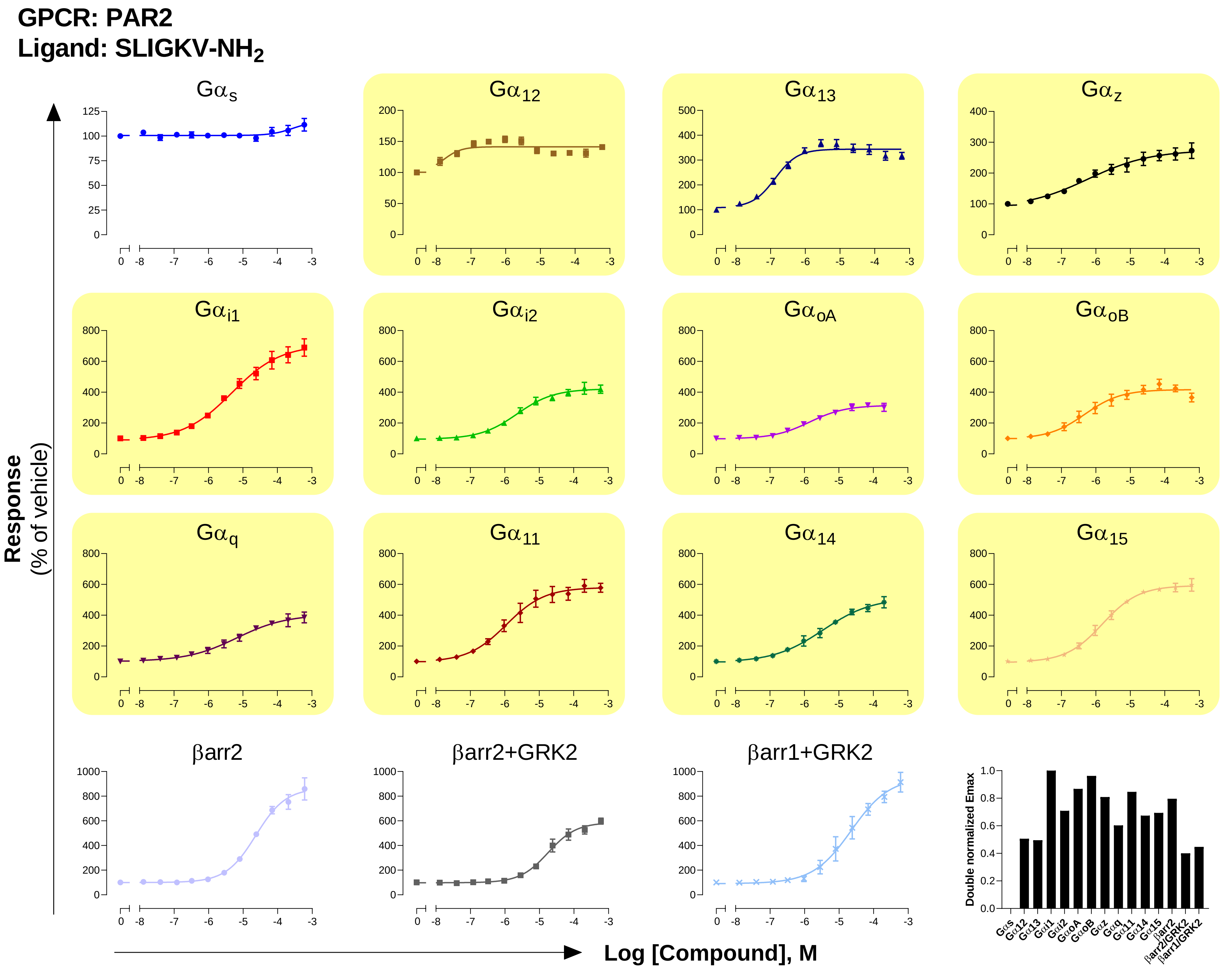

GPCR: PAR2
Ligand: SLIGKV-NH2
Gαs
Gα12
Gα13
Gαz
Gαi1
Gαi2
GαoA
GαoB
Gαq
Gα11
Gα14
Gα15
βarr2
βarr2+GRK2
βarr1+GRK2
Response
(% of vehicle)
Log [Compound], M
Double normalized Emax
Gαs
Gα12
Gα13
Gαi1
Gαi2
GαoA
GαoB
Gαz
Gαq
Gα11
Gα14
Gα15
βarr2
βarr2/GRK2
βarr1/GRK2

### GPCR: PTH1
### Ligand: PTH (1-34)

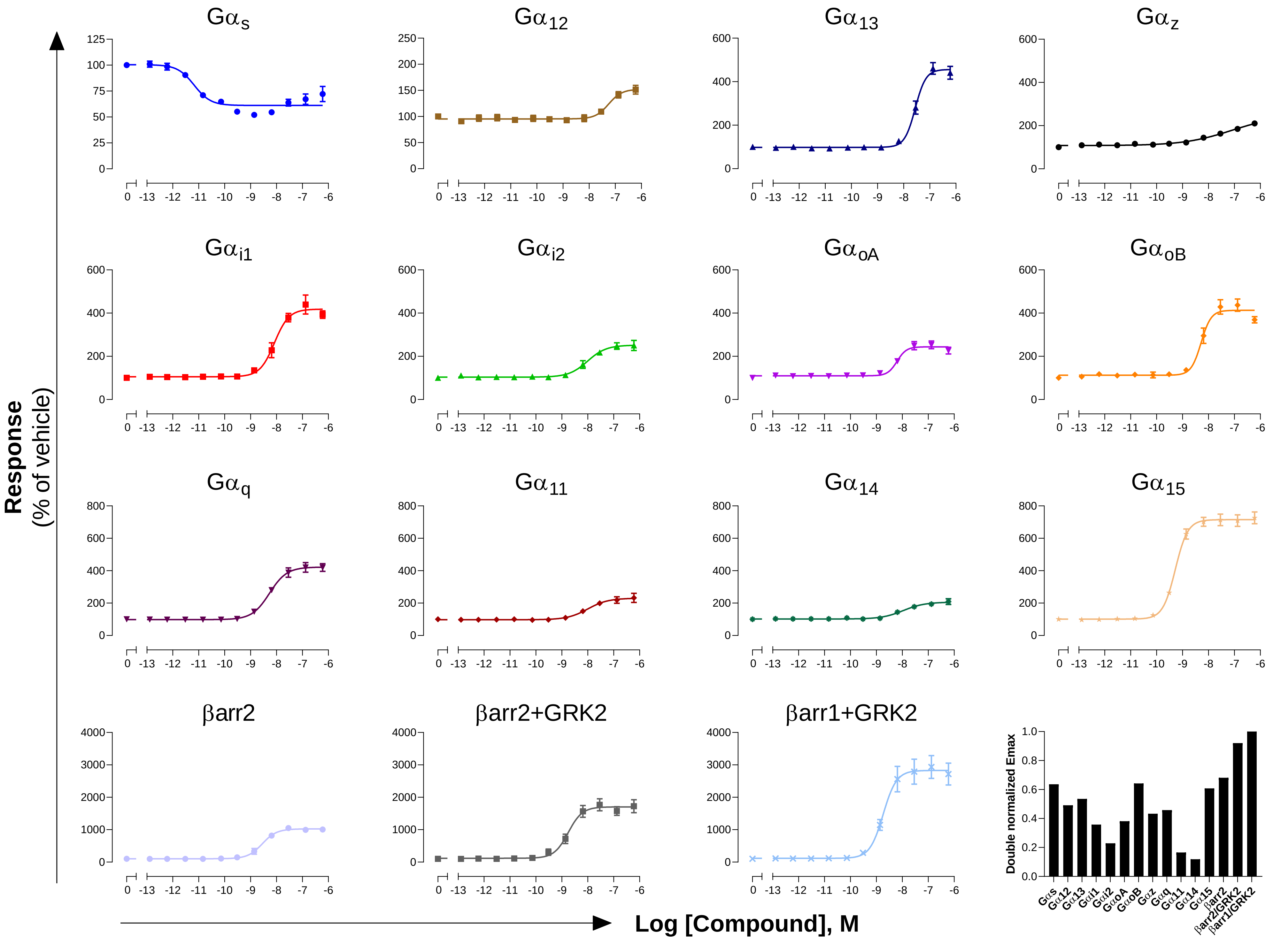

**GPCR: S1P$_1$**
**Ligand: Sphingosine 1-phosphate**

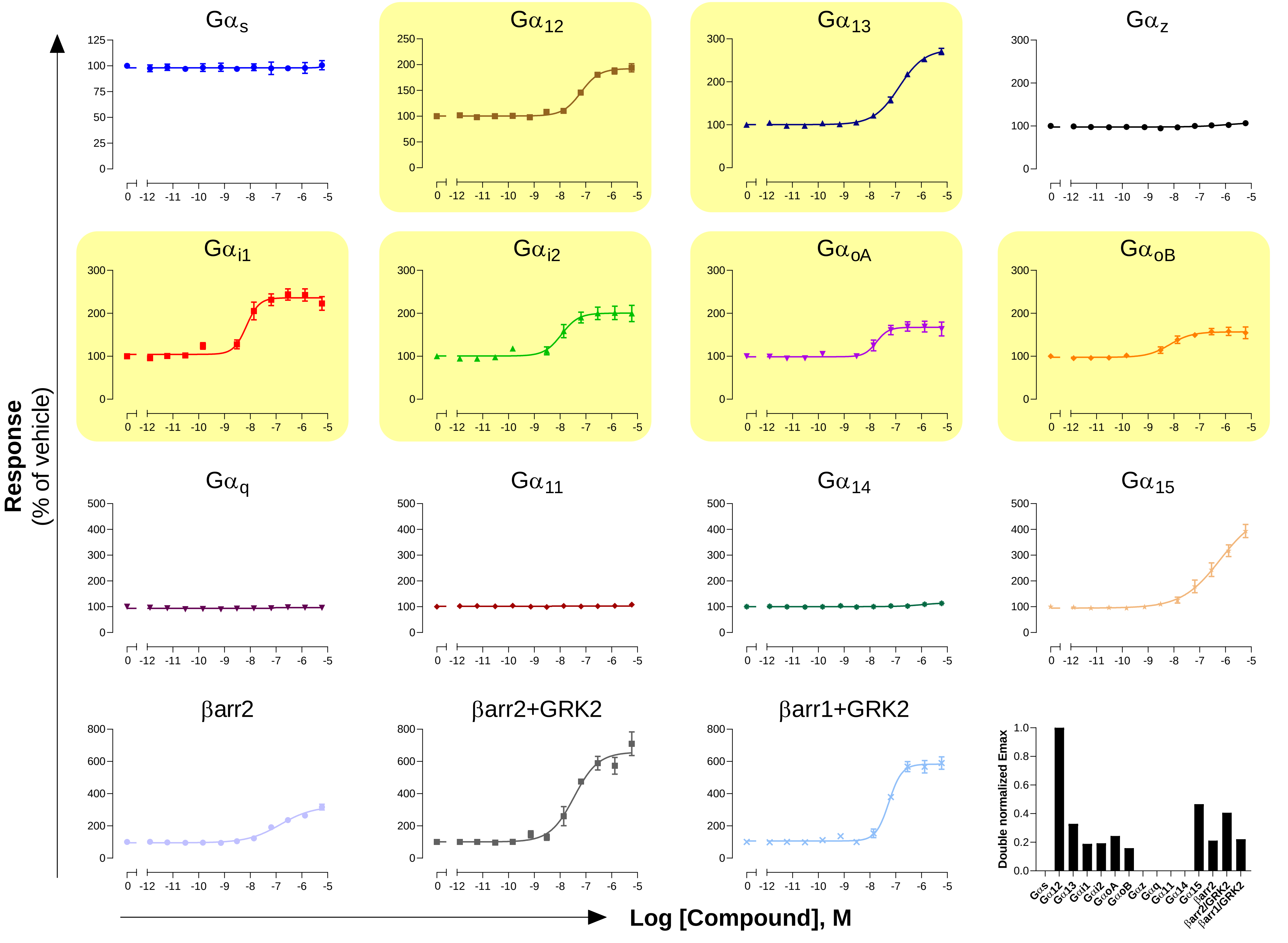

### GPCR: $SST_{2A}$
### Ligand: Somatostatin-14

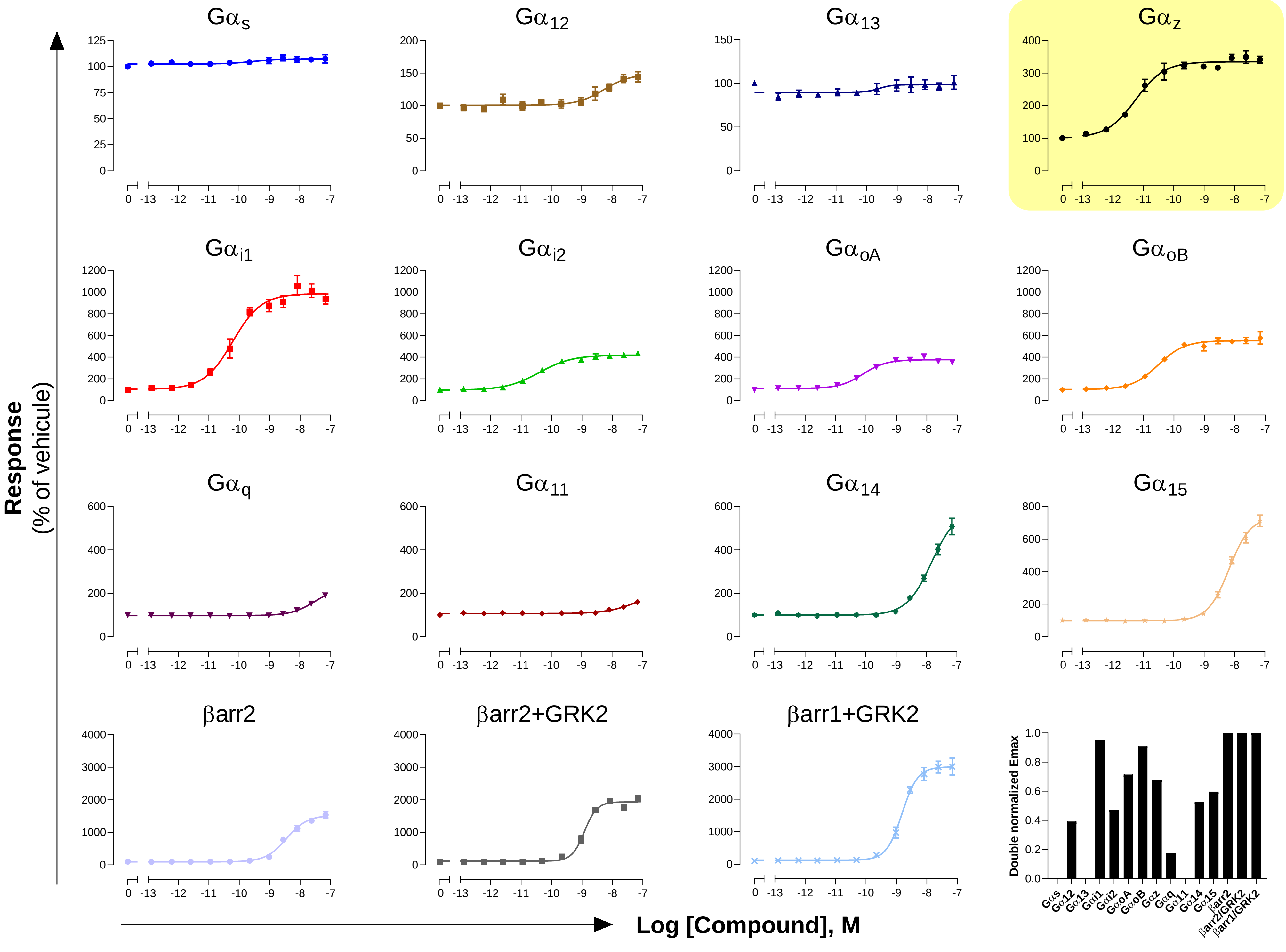

**GPCR: $V_{1A}$**
**Ligand: AVP**

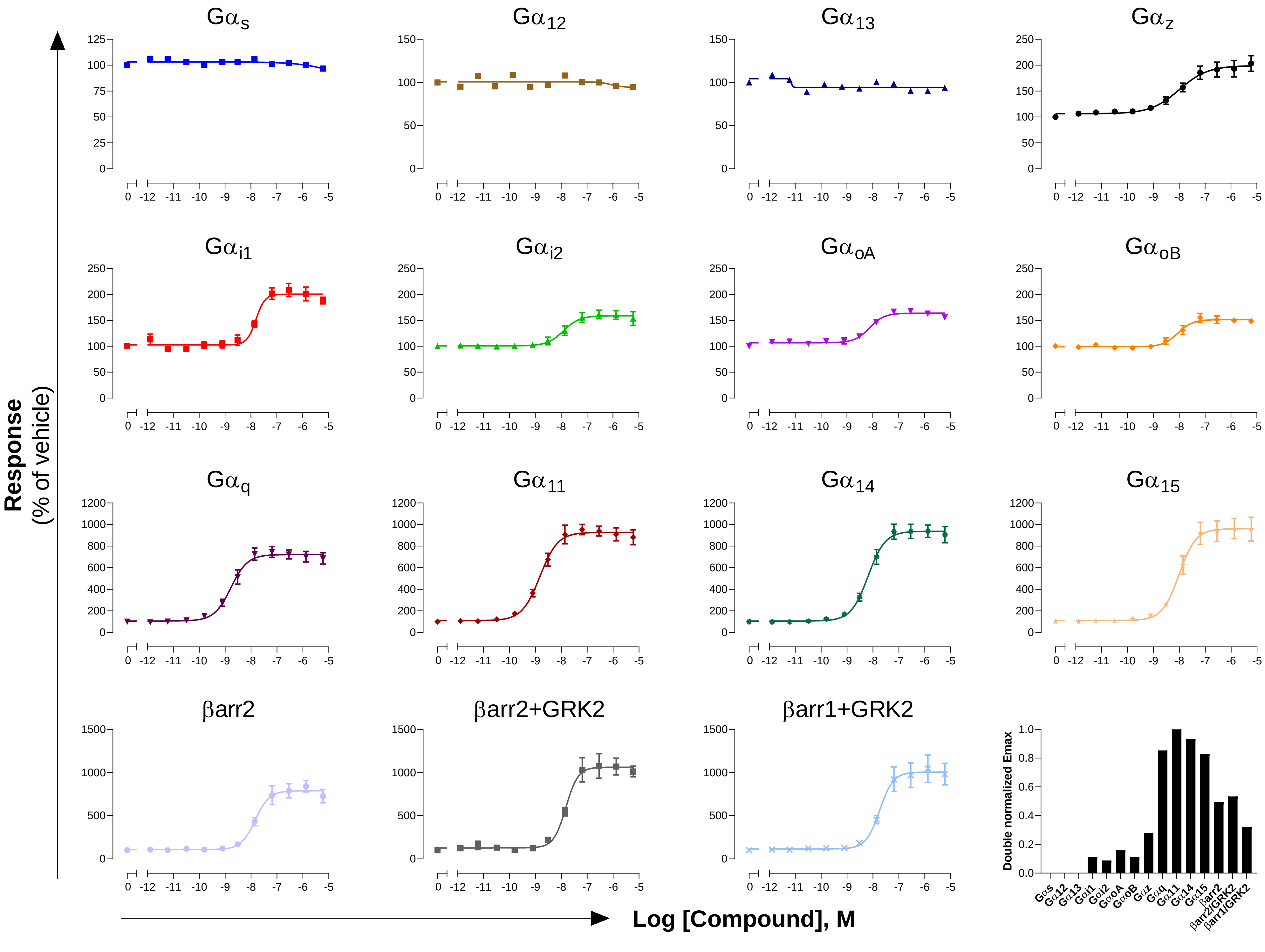

**GPCR: $V_2$**
**Ligand: AVP**

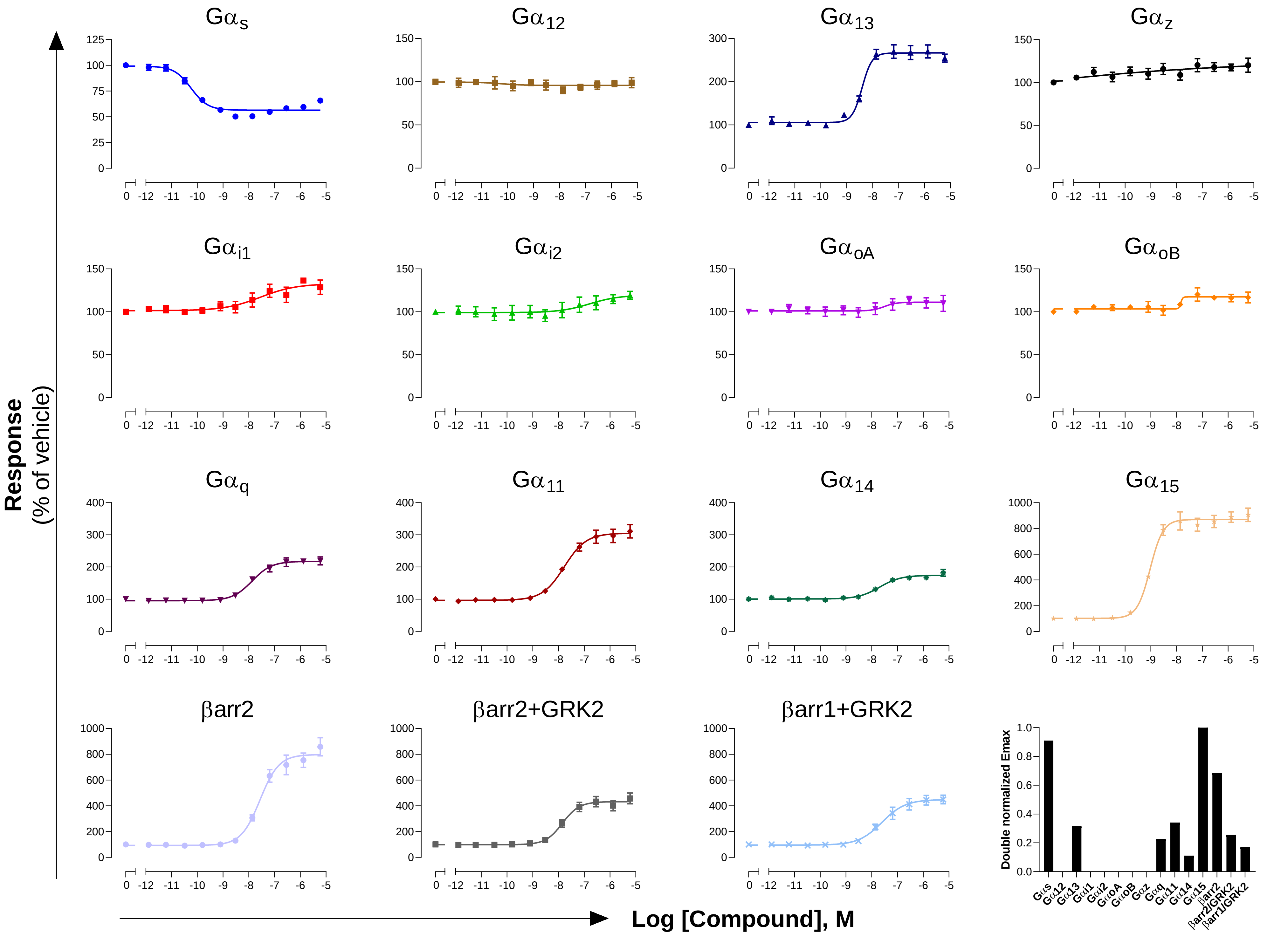

**GPCR: $VPAC_1$**

**Ligand: VIP**

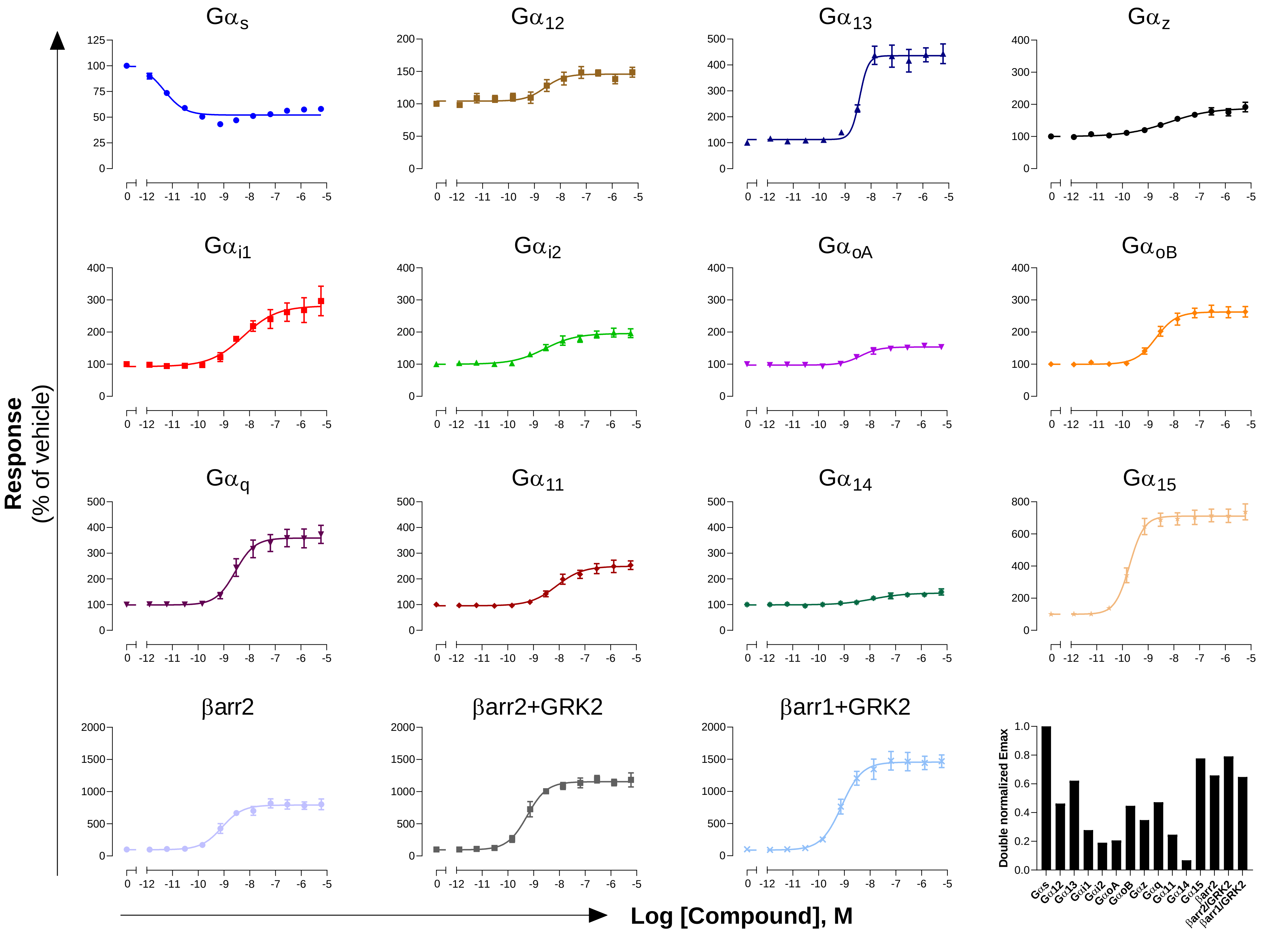

**GPCR: $Y_1$**
**Ligand: NPY**

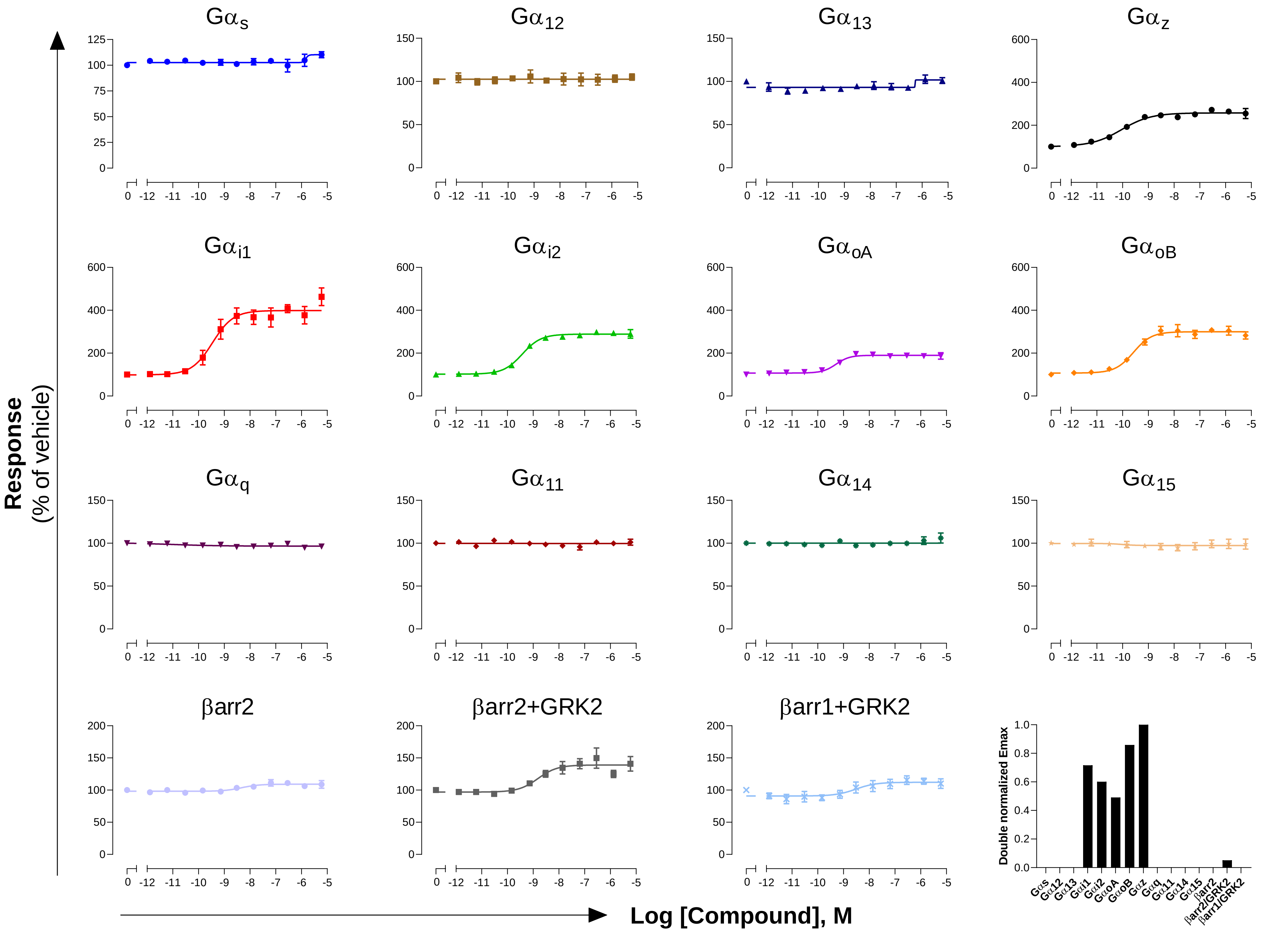

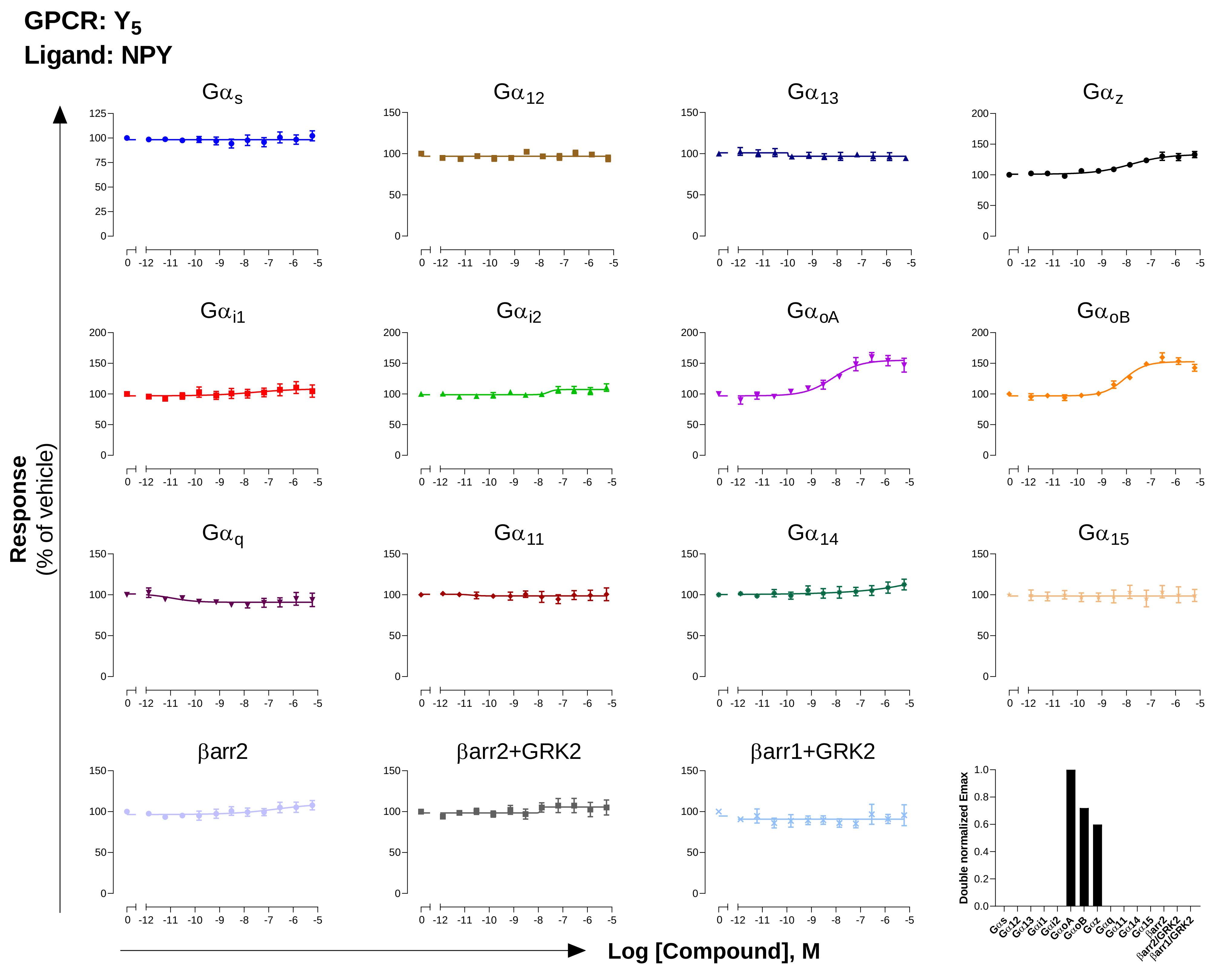

GPCR: Y5
Ligand: NPY
Gαs
Gα12
Gα13
Gαz
Gαi1
Gαi2
GαoA
GαoB
Gαq
Gα11
Gα14
Gα15
βarr2
βarr2+GRK2
βarr1+GRK2
Response
(% of vehicle)
Log [Compound], M
Double normalized Emax
